# Balancing Rigidity and Flexibility: Optimised 4-(Hexyloxy)benzoate Antagonists with Enhanced Affinity and Tuneable Duration at Muscarinic Receptors

**DOI:** 10.1101/2025.06.26.661741

**Authors:** Eva Dolejší, Eva Mezeiova, Jana Bláhová, Nikolai Chetverikov, Alena Janoušková-Randáková, Dominik Nelic, Lukas Prchal, Barbora Svobodova, John Boulos, Jan Korabecny, Jan Jakubík

**Affiliations:** Department of Neurochemistry, Institute of Physiology, Czech Academy of Sciences, Prague, Czech Republic; Biomedical Research Centre, University Hospital Hradec Kralove, Sokolska 581, 500 05 Hradec Kralove, Czech Republic; Department of Physical Sciences, Barry University, Miami Shores, FL, USA

**Keywords:** Muscarinic receptors, long-acting antagonists, ligand docking, molecular dynamics

## Abstract

Muscarinic acetylcholine receptors (mAChRs) are key regulators of diverse physiological processes and longstanding therapeutic targets. Building on the long-acting antagonist KH-5, we synthesised and evaluated a series of 4-(hexyloxy)benzoate derivatives and their quaternary N-methylated analogues to explore how structural modifications influence receptor affinity and the duration of functional antagonism. Our structure-activity analysis revealed that introducing a rigid azabicyclo[2.2.2]octan-1-ium group boosted binding affinity (up to 250-fold compared to parental compounds) yet reduced the half-life of functional antagonism. In contrast, analogues with moderate flexibility maintained high potency while preserving longer receptor residence time. Computational docking and molecular dynamics (MD) simulations demonstrated that stable hydrogen bonding with residue N6.52 and salt-bridge formation with D3.32 were critical for sustained ligand binding to the receptor, with MD-derived metrics outperforming docking energies in predicting biological activity. Crucially, a positively charged nitrogen and a 4-hexyloxy substituent are essential features for high-affinity binding and prolonged antagonism. Shortening the alkyl chain resulted in a marked loss of affinity and abolished sustained activity. These findings underscore the need to balance molecular rigidity with conformational flexibility and charge distribution in the design of long-residence mAChR antagonists, offering a framework for further development of mAChR-targeted long-acting antagonists.

**Highlights:**

- New analogues show up to 250× higher affinity at muscarinic receptors
- N6.52 H-bonding during MD predicts compound binding better than docking energies
- Charged nitrogen and 4-hexyloxy are key to high affinity and sustained action
- Rigid azabicyclo groups boost potency but shorten antagonism duration
- Flexible analogues balance potency with longer receptor residence time better

## 1. Introduction

Muscarinic acetylcholine receptors mediate a wide range of physiological functions in the CNS as well as in the periphery. Depending on the anatomic location and subtype of muscarinic receptor, abnormalities in cholinergic signalling are manifested as various diseases and conditions. M₁, M₃ and M₅ receptors couple to Gₐ/₁₁ G-proteins, activating phospholipase C (PLC) to increase intracellular calcium. For example, the M₁ receptor in the CNS modulates cognitive functions like memory, motor activity in the striatum and GABAergic transmission in the hippocampus. M₃ receptor mediates smooth muscle contraction (e.g., bronchoconstriction) and glandular secretion. M_5_ receptor mediates vasodilation of brain microvasculature and is part of the dopamine reward circuit in the substantia nigra and ventral tegmental area. M₂ and M₄ receptors couple to Gᵢ/ₒ G-proteins, inhibiting cAMP and modulating ion channels. For example, the M₂ receptor slows heart rate and presynaptically inhibits acetylcholine release. The M_4_ receptor mediates cognitive processes in the brain and modulates acetylcholine release and dopaminergic transmission in the striatum. Both M_2_ and M_4_ receptors mediate analgesia.

Muscarinic antagonists have been used in the treatment of endogenously (e.g., asthma, COPD) or exogenously (e.g., organophosphate poisoning) overstimulated muscarinic receptors. Muscarinic antagonists, which inhibit parasympathetic activity, are clinically used for respiratory disorders or an overactive bladder. Short-acting antagonists (e.g., ipratropium) are used for acute bronchospasm in chronic obstructive pulmonary disease (COPD) and asthma[1]. Long-acting muscarinic antagonists (LAMAs) like tiotropium are used for maintenance therapy[2]. Oxybutynin reduces urinary urgency by blocking M₃receptors[3]. Recently, a selective M_5_ antagonist for the treatment of opioid use disorder was developed[4]. Historically, centrally acting muscarinic antagonists are used for neurological conditions for motion sickness (scopolamine), Parkinson’s tremors (benztropine) or status epilepticus after organophosphate exposure[5].

LAMAs (e.g., tiotropium, glycopyrronium) have become the cornerstone therapy for COPD and asthma. Tiotropium exerts extremely slow dissociation from M₃ receptors (t_1/2_ ≈ 62 hours)[6]. In general, the effects of ligands with long residence time at receptors last longer, thus allowing for lower daily drug doses to reach a maximum therapeutic effect and reduce side effects like dry mouth, constipation, and blurred vision, resulting from systemic anticholinergic activity. Only several LAMAs are clinically approved, all of them for COPD treatment [7,8]. LAMAs with different selectivity profiles would be beneficial for the treatment of other conditions associated with excessive activity of muscarinic receptors. For example, M_1_ antagonists have potential in the treatment of epilepsy or seizures after organophosphate poisoning[5] or cholinergic dystonia[9]. Centrally acting M_4_ antagonists have potential in the treatment of Parkinson’s disease[10].

KH-5 (1-{2-[4-(hexyloxy)benzoyloxy]ethyl}-1-methyl-1,2,3,6-tetrahydropyridin-1-ium) is the LAMA with moderately slow dissociation (t_1/2_ up to 5 hours)[11]. The sustained activity of long-acting agonist xanomeline[12] (t_1/2_ ≈ 30 hours) is remarkably longer[13]. Similarly, to the long-acting agonist xanomeline, the hexyloxy moiety is required for sustained effects. In our recent study, we synthesised 3-hexyloxy and 2-hexyloxy analogues of KH-5 and have shown that they have shorter residence times than KH-5, proving that the 4-hexyloxy position on the benzene ring is optimal for sustained activity[14]. Therefore, in this study, we have focused on the effects of modifications of KH-5 nitrogen-containing heterocycle primarily on binding affinity (and thus potency) while preserving the extended duration of functional antagonism. In silico, we docked compounds to the orhtosteric binding site and compared computed affinity estimates with experimentally determined ones to test the docking procedure for possible future virtual screening.

## 2. Design

The sustained activity of long-acting agonist xanomeline (t_1/2_ ≈ 30 hours) is remarkably longer than the sustained antagonism of long-acting antagonist KH-5 (t_1/2_ up to 5 hours) [11,13] (Supplementary Information Table S1). It was postulated that the hexyloxy chain is responsible for wash-resistant binding and consequent sustained action of muscarinic ligands. The relative orientation of the hexyloxy moiety towards the tetrahydropyridine ring differs between xanomeline and KH-5. Therefore, in our recent study, we synthesised 3-hexyloxy and 2-hexyloxy analogues of KH-5 [14]. We have found that moving the hexyloxy chain to positions 3 and 2 shortened residence time and decreased binding affinity of these compounds. Therefore, we investigated 4-hexyloxy analogues of KH-5.

The design of novel derivatives followed the structural changes in one region of the lead compound KH-5 (Figure 1). The targeted part is highlighted in purple and orange. Firstly, we focused on the structural features in the vicinity of nitrogen, resulting either in amino-or imino-based compounds (Figure 1, purple). The 1,2,3,6-tetrahydropyridine from the KH-5 template was replaced by bicyclic scaffolds or more complex piperidine-based structures (2-methylpiperidine or 1,2,3,4-tetrahydroquinoline). We aimed to investigate the impact of replacing a six-membered heterocycle with a five-membered one on the biological activity of the compounds. To this end, we designed and synthesised derivatives incorporating pyrrolidine and 3-pyrroline, followed by the development of compounds with more structurally complex frameworks. The second modification, highlighted in orange (Figure 1), also targeted the same region, increasing the rigidity of the side linker that connects the nitrogen atom to an ester bond. Specifically, we replaced the two methylene linkers with (pyrrolidin-2(or 3)-yl)methyl or (piperidin-2(or 3 or 4)-yl)methyl moieties.

**Figure 1.**
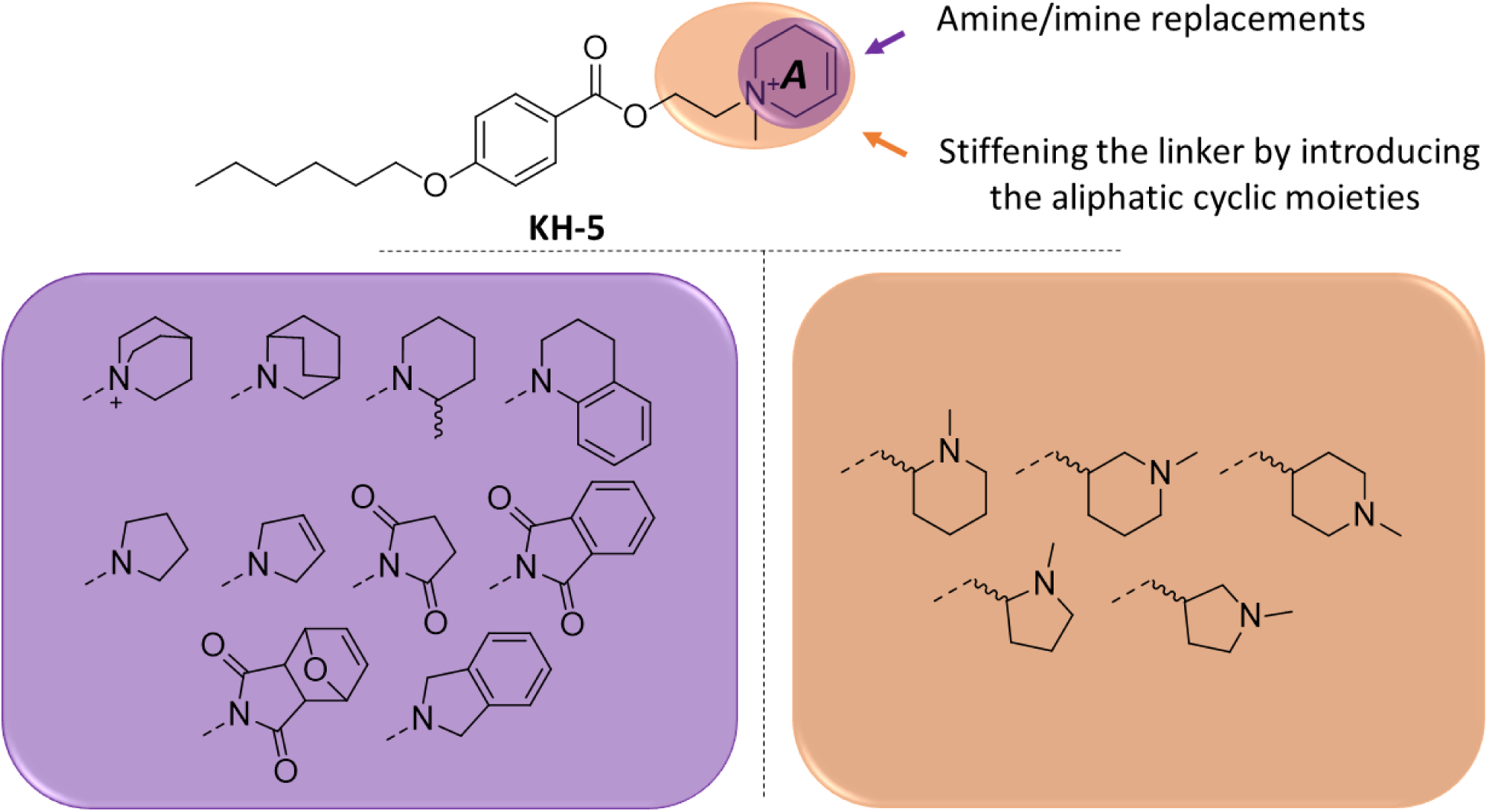
Design of novel derivatives inspired by KH-5. Two types of changes in the parent were considered: i) with compounds either containing amine or imine moieties (purple), and ii) stiffening the linker between the ester moiety and nitrogen by introducing aliphatic cyclic moieties (orange).

## 3. Results and discussion

### 3.1. Synthesis

Carboxylic acid **4** was prepared in two steps, starting from 4-hydroxybenzaldehyde (**1**) (Figure 2). *O*-Alkylation of **1** with 1-bromohexane (**2**) provided aldehyde **3**, which was then oxidised into **4** applying Pinnick oxidation (NaH_2_PO_4_, NaClO_2_, 2-methyl-2-butene, THF:H_2_O:*^t^*BuOH = 4:4:1). Final compounds **6a**-**z** were prepared by two different synthetic routes. Esterification of **4** with 2-bromoethanol generated bromide **5**. Its reaction with corresponding amines led to the desired compounds **6a**-**j** in 7–64% yields (Figure 2). The second synthetic route consisted of the esterification of **4** by various alcohols (**8k-z**), affording the formation of final compounds **6k**-**z** (Figure 3). Quaternary ammonium salts (**7e**-**z**) were synthesised by *N*-methylation of **6e**-**z** with CH_3_I (Figure 2 and Figure 3). Unfortunately, we were not able to isolate derivative **7l** in sufficient purity, exhibiting low stability and leading to gradual compound decomposition. Derivatives **12**–**14** were prepared from *p*-alkyloxybenzoyl chlorides **9**–**11** and 3-quinuclidinol (Figure 4) by a substitution reaction that provided corresponding esters, which were then transformed into hydrochloride salts **12a**, **13a**, **14a** or into quaternary ammonium salts **12b**, **13b**, **14b**.

**Figure 2.**
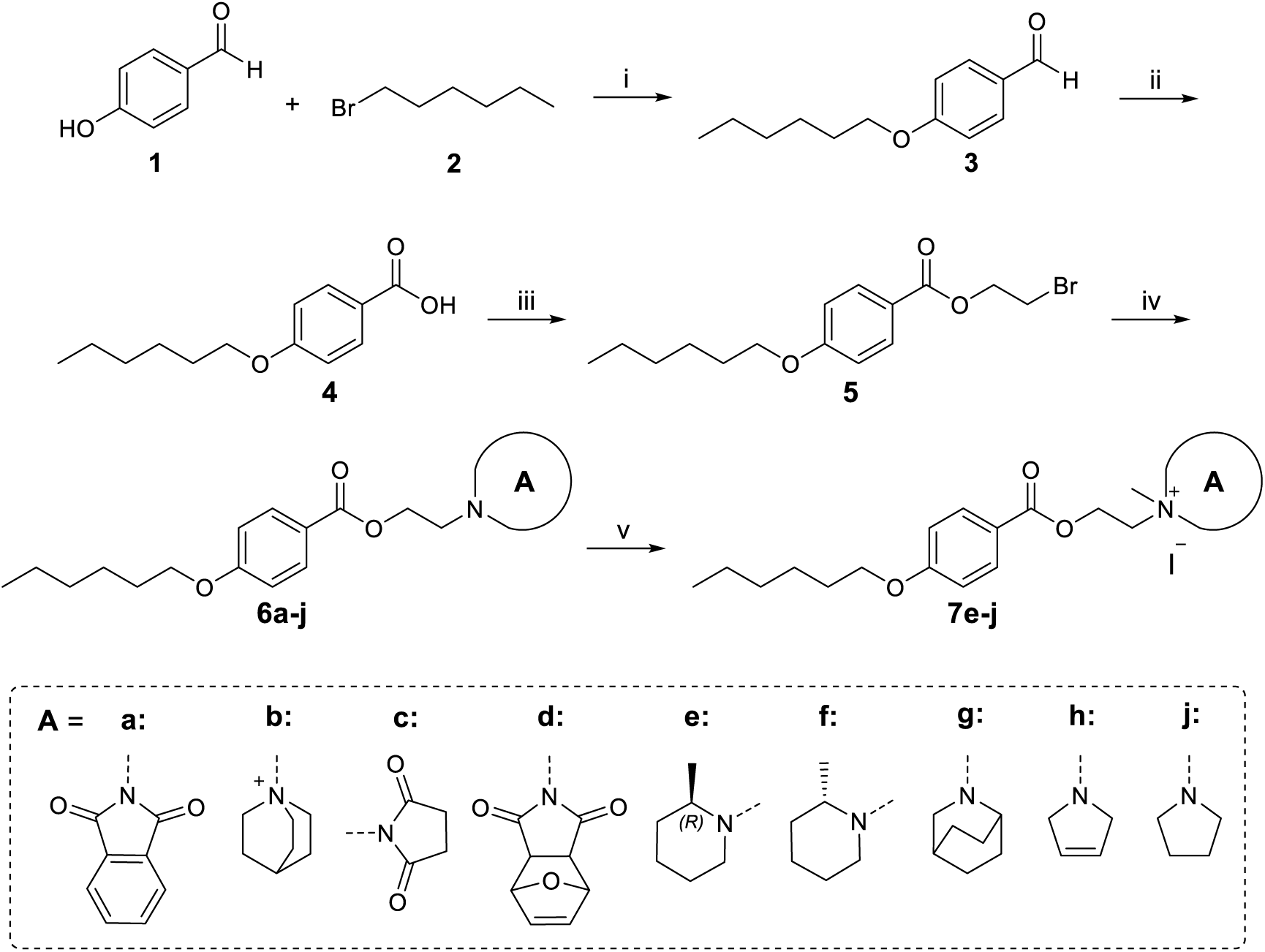
Synthesis of final compounds 6a-j and 7e-j. Reagents and conditions: i) K_2_CO_3_, CH_3_CN (dry), reflux, overnight, 96%; ii) NaH_2_PO_4_, NaClO_2_, 2-methyl-2-butene, THF:H_2_O:^t^BuOH = 4:4:1, rt, 1 h, 99%; iii) 2-bromoethanol, DCC, DMAP, CH_2_Cl_2_ (dry), rt, overnight, 65%; iv) corresponding amine, K_2_CO_3_, CH_3_CN, rt, overnight, 7–64%; v) CH_3_I, CH_3_CN, rt, overnight, 37–71%.

**Figure 3.**
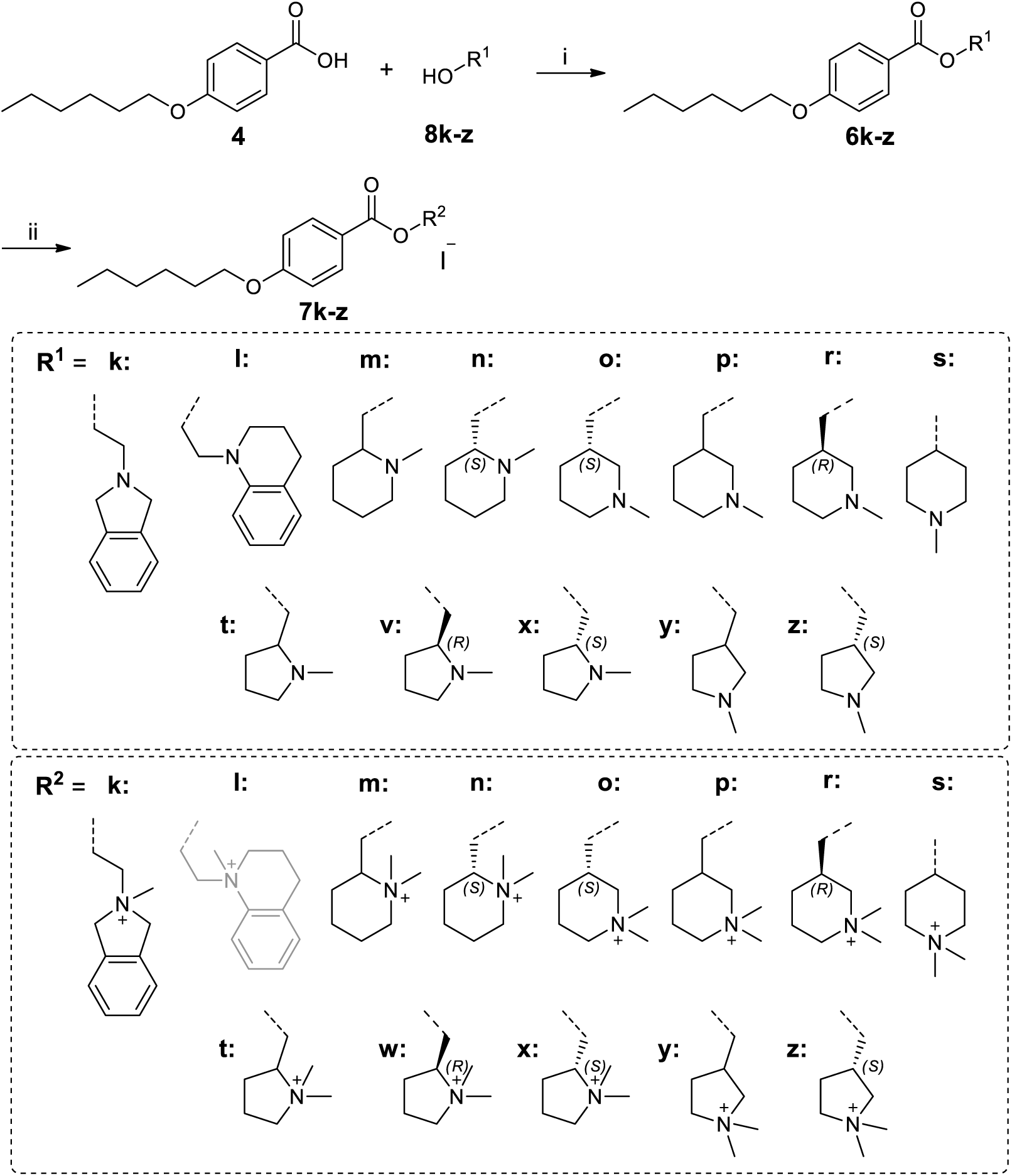
Synthesis of final compounds 6k-z and 7k-z. Reagents and conditions: i) corresponding alcohol **8k**-**z**, DCC, DMAP, CH_2_Cl_2_ (dry), rt, overnight, 18–64%; ii) CH_3_I, CH_3_CN, rt, overnight, 17–42%.

**Figure 4.**
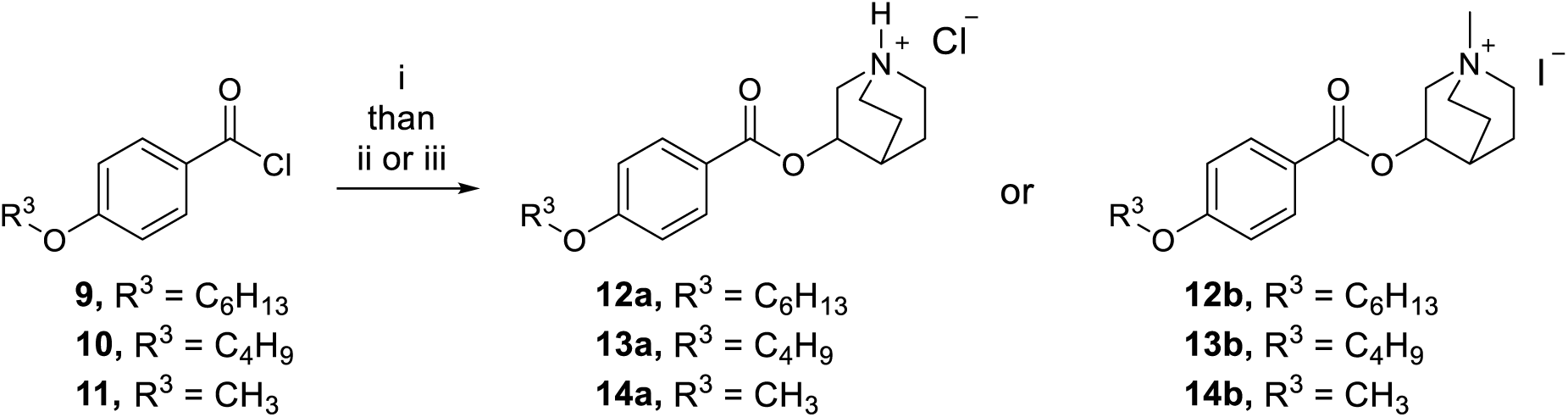
Synthesis of final compounds 12–14. Reagents and conditions: i) 3-quinuclidinol, K_2_CO_3_, CH_3_CN, rt, overnight; ii) gas HCl, dry pentane, 40–91%; iii) CH_3_I, CH_3_CN, rt, overnight, 78–89%.

The newly developed compounds were characterised using ^1^H NMR and ^13^C NMR spectroscopy (except for **7m** and **7n** due to insufficient amount for analysis) and HPLC−HRMS experiments (Supporting Information). All compounds exerted >95% purity by HPLC analysis (see Experimental section for more details), with the only exception of **12b** (92% purity).

### 3.2. Molecular Modelling

#### 3.2.1. Ligand docking

To test the suitability of the proposed compounds as muscarinic binders, KH-5 and the proposed compounds were docked to the orthosteric sites of all five muscarinic receptors. Except for the M_3_ receptor, binding energies of top-scoring poses of proposed compounds were up to 2 kcal/mol higher than the binding energies of KH-5 (Supplementary information Table S3), corresponding to about a 30-fold increase in affinity[15]. In general, the binding energy estimates were higher for compounds bearing charged nitrogen than for compounds with neutral nitrogen. The highest binding energies were estimated for compounds containing an azabicyclo[2.2.2]octan-1-ium group attached to the rest of the molecule at position 3 (compounds **12a** and **12b**) or via nitrogen (compound **6b**). Except for the M_2_ receptor, binding energy estimates for compounds containing 2-azabicyclo[2.2.2]octan-2-yl group (compounds **6g** and **7g**) were about 1 kcal/mol lower than for **12a** and **12b**, respectively. Interestingly, compounds containing phthalimide (compound **6a**), indoline (compounds **6k** and **7k**) or tetrahydroquinoline (compound **6l**) heterocycles. The lowest binding energy estimates were obtained for compounds **6j** and **7j**.

All compounds formed hydrogen bonds with N^6.52^ (Ballesteros and Weinstein numbering [16]) in TM6, which is essential for the binding of antagonists [17–20] at all receptors. Some KH-5 analogues formed additional interactions, salt bridges with D^3.32^ via nitrogen, cation-π interactions with Y^3.33^, W^6.48^ and Y^7.39^ and additional hydrogen bond with T^5.42^ via ether oxygen (Figure 5, Supplementary information Table S3). In total, six amino acid residues (D^3.32^, Y^3.33^, T^5.42^, W^6.48^, N^6.52^, Y^7.39^) principal for compounds binding were identified. Compounds **7e** to **7k** lacked salt bridges to D^3.32,^ although they possess a tertiary nitrogen. The ligand-receptor interactions were identical at all receptor subtypes, indicating similar affinities to all subtypes and a lack of subtype binding selectivity.

**Figure 5.**
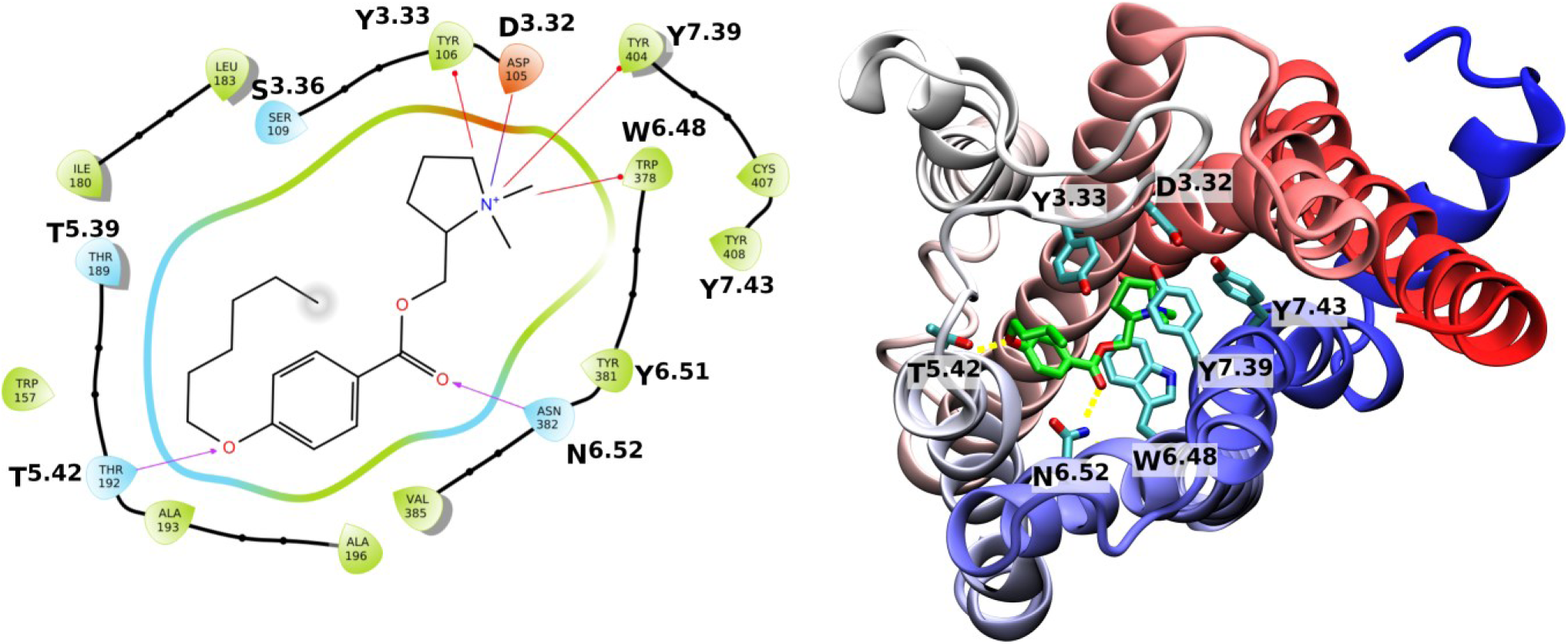
Example of KH-5 analogue (7w) binding to the orthosteric site of muscarinic receptor. 2D scheme (left) and 3D view (right) of top binding pose of **7w** in the orthosteric site of M_1_ receptor (5CXV). Compound **7w** forms hydrogen bonds (violet arrows left, dashed yellow lines right) with Asn382 (N^6.52^) via ketoxy oxygen and Thr192 (T^5.42^) via ether oxygen, salt bridge (blue-red line) with Asp105 (D^3.32^) and cation-π interactions with Tyr106 (Y^3.33^), Trp378 (W^6.48^) and Tyr404 (Y^7.39^).

#### 3.2.2. Molecular dynamics

To test the stability of compound binding and to analyse ligand-receptor interactions of top binding poses, 120 ns of conventional molecular dynamics (cMD) was simulated. Simulation of cMD revealed the instability of key interactions and subsequent binding instability for some ligands (Supplementary Information Table S4). Although binding energy estimates from docking were relatively high, hydrogen bonding of compounds **6a**, **6c**, **6d** and **6g** to **6l** with N^6.52^ was disrupted during cMD. The breakdown of the hydrogen bond to N^6.52^ of these compounds was accompanied by relatively large movement from the initial pose, indicating weak or no binding. The M_1_ receptor examples are in Figure 6, left. In contrast, salt bridges to D^3.32^ that were absent in top docking poses formed for compounds **6b**, **7e**, **7f**, **7h,** and **7j** during cMD runs (Supplementary Information Table S4). The binding of compounds **7g**, **7s**, **7w**, **12a**, and **12b,** which had high binding energy estimates, was stable with low RMDS during cMD (Figure 66, right).

**Figure 6.**
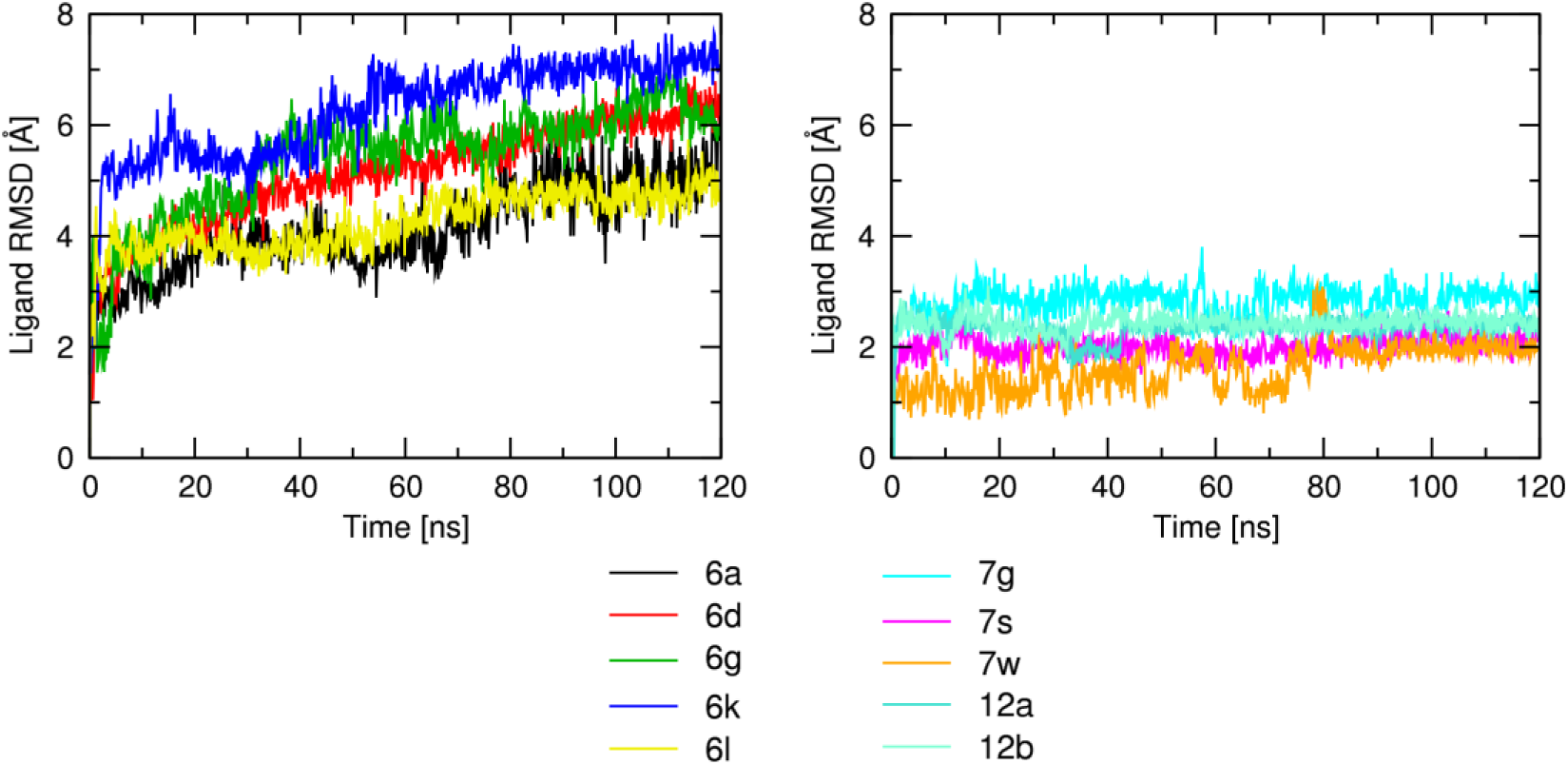
Ligand RMSDs over the course of cMD at the M_1_ receptor. Examples of the RMSDs of top poses to the orthosteric binding site from the protein-ligand complex aligned on the protein backbone of the initial frame in Å are plotted against simulation time in ns. Individual compounds are indicated in the graph legend. Traces are representative of 3 independent MD replicas.

### 3.3. Activity at muscarinic receptors

#### 3.3.1. Structure-activity relationships – basic centre

The binding affinity of a given compound was measured in competition experiments with tritiated orthosteric antagonist *N*-methyl scopolamine ([^3^H]NMS). Compounds with affinity lower than 10 μM (10 μM compounds displacing less than 50 % of [^3^H]NMS binding) were considered non-binders. Eight compounds were non-binders: **6a**, **6c**, **6d** and **6g** to **6l**. A common feature of non-binders is the lack of charged nitrogen. Analogues of non-binders with charged nitrogen bound to muscarinic receptors with an affinity of about 1 μM or higher. Overall, none of the compounds displayed substantial binding selectivity. In addition, compounds with quaternary nitrogen had higher affinity than analogues with tertiary nitrogen (e.g. **7g** vs **6g**). However, **12b** (excluding M_2_) had slightly lower affinities than **12a**. The highest binding affinity, about 30 nM, was detected for compounds **12a** and **12b**, with **12a** having a higher affinity than **12b** at all subtypes except M_2_ ^(^Table ^1^). In comparison to parental compounds, it is increased up to 250-fold for hydrogen derivates (**12a** over OT-231) and up to 8-fold for methyl derivates (**12b** over KH-5). These compounds contain an azabicyclo[2.2.2]octan-1-ium moiety attached at position 3 to the rest of the molecule. Attaching azabicyclo[2.2.2]octan-1-ium via nitrogen (compound **6b**) resulted in affinities for muscarinic receptors about 10-fold lower than **12a**, except for M_4_ and M_5_ receptors, where the affinity of **6b** was only slightly lower. Using 2-azabicyclo[2.2.2]octan-2-yl (compound **6g**) resulted in the loss of affinity for muscarinic receptors. Charged nitrogen-containing compound **7g** regained affinities to a similar extent as for compound **7b**.

**Table 1.**
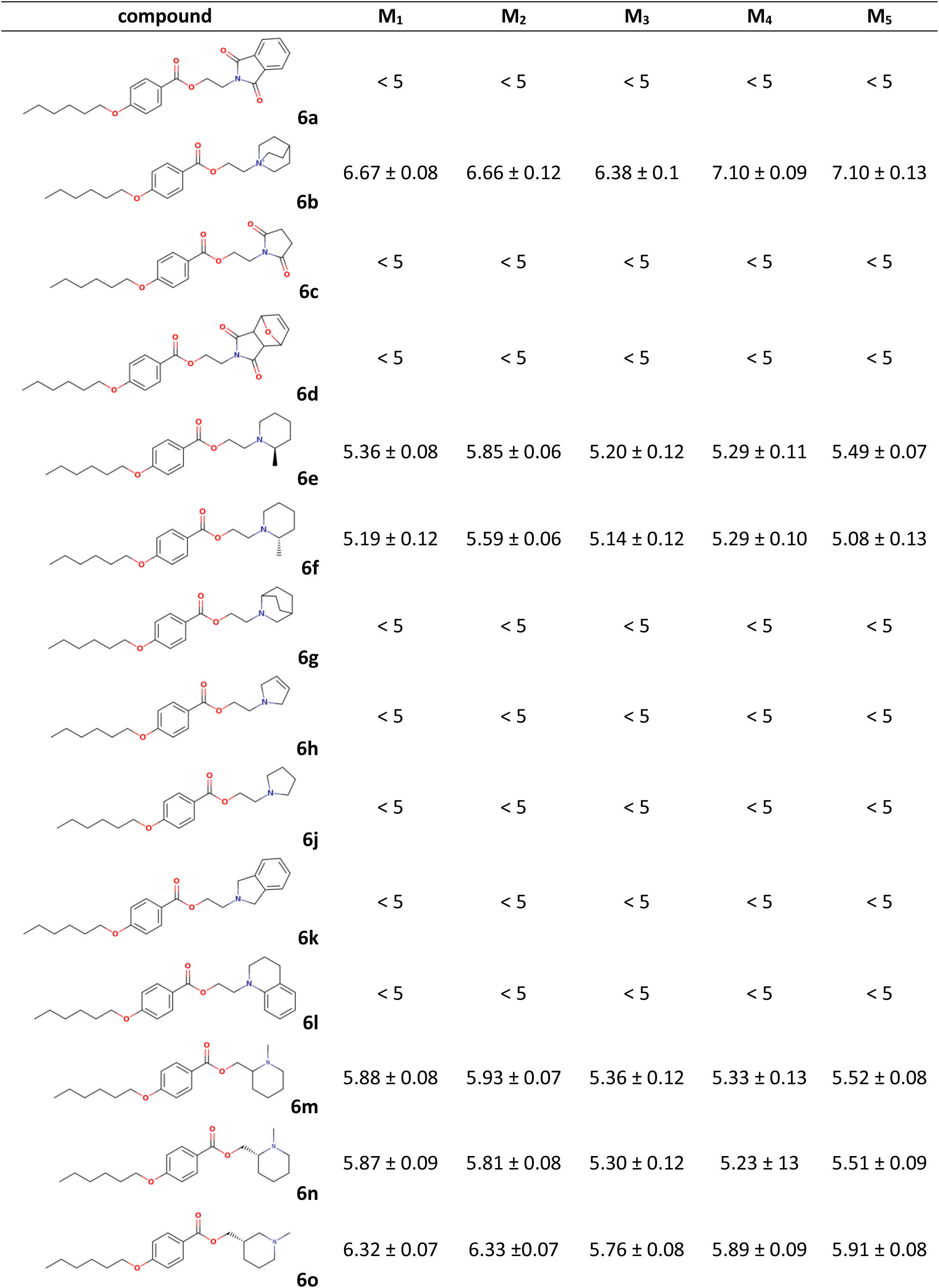
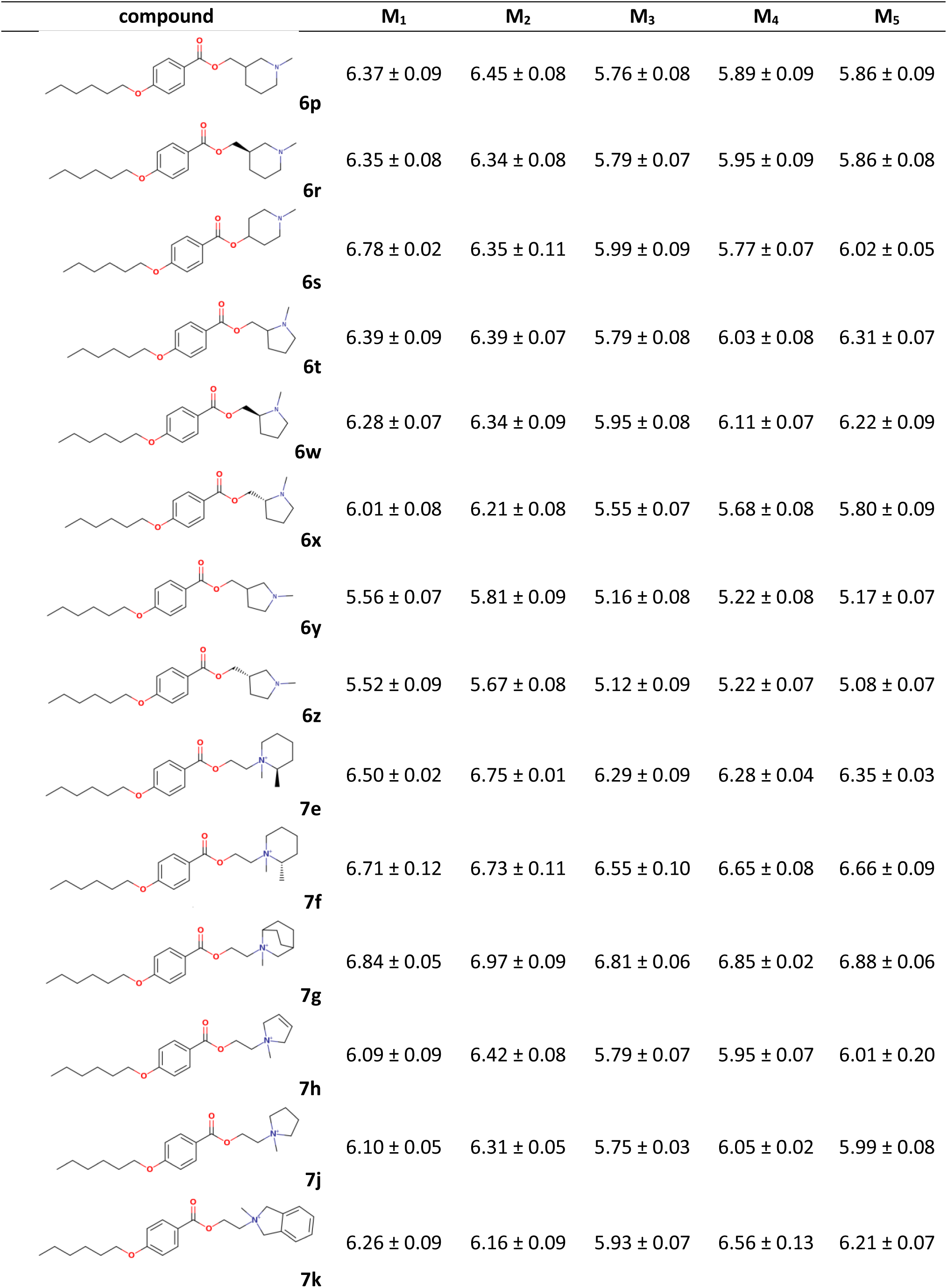
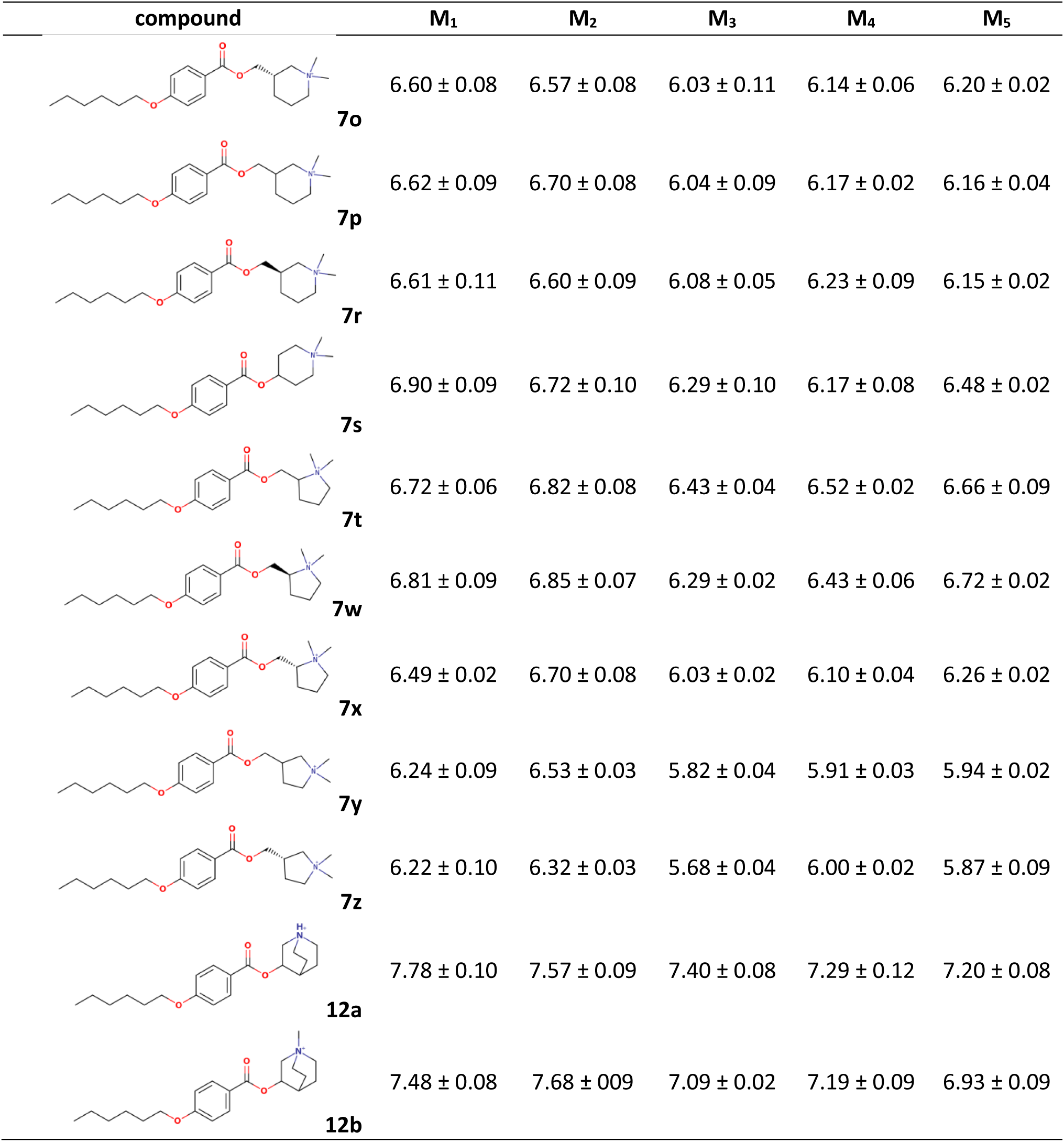
Affinities (pK_i_) of 3-[4-(hexyloxy)benzoyloxy] analogues. Affinities are expressed as the negative decadic logarithm of the inhibition constant (pK_i_) of [^3^H]NMS binding. Data are means ± SD from 3 independent experiments.

| compound | M<sub>1</sub> | M<sub>2</sub> | M<sub>3</sub> | M<sub>4</sub> | M<sub>5</sub> |
| --- | --- | --- | --- | --- | --- |
| **6a** | < 5 | < 5 | < 5 | < 5 | < 5 |
| **6b** | 6.67 ± 0.08 | 6.66 ± 0.12 | 6.38 ± 0.1 | 7.10 ± 0.09 | 7.10 ± 0.13 |
| **6c** | < 5 | < 5 | < 5 | < 5 | < 5 |
| **6d** | < 5 | < 5 | < 5 | < 5 | < 5 |
| **6e** | 5.36 ± 0.08 | 5.85 ± 0.06 | 5.20 ± 0.12 | 5.29 ± 0.11 | 5.49 ± 0.07 |
| **6f** | 5.19 ± 0.12 | 5.59 ± 0.06 | 5.14 ± 0.12 | 5.29 ± 0.10 | 5.08 ± 0.13 |
| **6g** | < 5 | < 5 | < 5 | < 5 | < 5 |
| **6h** | < 5 | < 5 | < 5 | < 5 | < 5 |
| **6j** | < 5 | < 5 | < 5 | < 5 | < 5 |
| **6k** | < 5 | < 5 | < 5 | < 5 | < 5 |
| **6l** | < 5 | < 5 | < 5 | < 5 | < 5 |
| **6m** | 5.88 ± 0.08 | 5.93 ± 0.07 | 5.36 ± 0.12 | 5.33 ± 0.13 | 5.52 ± 0.08 |
| **6n** | 5.87 ± 0.09 | 5.81 ± 0.08 | 5.30 ± 0.12 | 5.23 ± 13 | 5.51 ± 0.09 |
| **6o** | 6.32 ± 0.07 | 6.33 ± 0.07 | 5.76 ± 0.08 | 5.89 ± 0.09 | 5.91 ± 0.08 |
| 6p | 6.37 ± 0.09 | 6.45 ± 0.08 | 5.76 ± 0.08 | 5.89 ± 0.09 | 5.86 ± 0.09 |
| 6r | 6.35 ± 0.08 | 6.34 ± 0.08 | 5.79 ± 0.07 | 5.95 ± 0.09 | 5.86 ± 0.08 |
| 6s | 6.78 ± 0.02 | 6.35 ± 0.11 | 5.99 ± 0.09 | 5.77 ± 0.07 | 6.02 ± 0.05 |
| 6t | 6.39 ± 0.09 | 6.39 ± 0.07 | 5.79 ± 0.08 | 6.03 ± 0.08 | 6.31 ± 0.07 |
| 6w | 6.28 ± 0.07 | 6.34 ± 0.09 | 5.95 ± 0.08 | 6.11 ± 0.07 | 6.22 ± 0.09 |
| 6x | 6.01 ± 0.08 | 6.21 ± 0.08 | 5.55 ± 0.07 | 5.68 ± 0.08 | 5.80 ± 0.09 |
| 6y | 5.56 ± 0.07 | 5.81 ± 0.09 | 5.16 ± 0.08 | 5.22 ± 0.08 | 5.17 ± 0.07 |
| 6z | 5.52 ± 0.09 | 5.67 ± 0.08 | 5.12 ± 0.09 | 5.22 ± 0.07 | 5.08 ± 0.07 |
| 7e | 6.50 ± 0.02 | 6.75 ± 0.01 | 6.29 ± 0.09 | 6.28 ± 0.04 | 6.35 ± 0.03 |
| 7f | 6.71 ± 0.12 | 6.73 ± 0.11 | 6.55 ± 0.10 | 6.65 ± 0.08 | 6.66 ± 0.09 |
| 7g | 6.84 ± 0.05 | 6.97 ± 0.09 | 6.81 ± 0.06 | 6.85 ± 0.02 | 6.88 ± 0.06 |
| 7h | 6.09 ± 0.09 | 6.42 ± 0.08 | 5.79 ± 0.07 | 5.95 ± 0.07 | 6.01 ± 0.20 |
| 7j | 6.10 ± 0.05 | 6.31 ± 0.05 | 5.75 ± 0.03 | 6.05 ± 0.02 | 5.99 ± 0.08 |
| 7k | 6.26 ± 0.09 | 6.16 ± 0.09 | 5.93 ± 0.07 | 6.56 ± 0.13 | 6.21 ± 0.07 |
| **7o** | 6.60 ± 0.08 | 6.57 ± 0.08 | 6.03 ± 0.11 | 6.14 ± 0.06 | 6.20 ± 0.02 |
| **7p** | 6.62 ± 0.09 | 6.70 ± 0.08 | 6.04 ± 0.09 | 6.17 ± 0.02 | 6.16 ± 0.04 |
| **7r** | 6.61 ± 0.11 | 6.60 ± 0.09 | 6.08 ± 0.05 | 6.23 ± 0.09 | 6.15 ± 0.02 |
| **7s** | 6.90 ± 0.09 | 6.72 ± 0.10 | 6.29 ± 0.10 | 6.17 ± 0.08 | 6.48 ± 0.02 |
| **7t** | 6.72 ± 0.06 | 6.82 ± 0.08 | 6.43 ± 0.04 | 6.52 ± 0.02 | 6.66 ± 0.09 |
| **7w** | 6.81 ± 0.09 | 6.85 ± 0.07 | 6.29 ± 0.02 | 6.43 ± 0.06 | 6.72 ± 0.02 |
| **7x** | 6.49 ± 0.02 | 6.70 ± 0.08 | 6.03 ± 0.02 | 6.10 ± 0.04 | 6.26 ± 0.02 |
| **7y** | 6.24 ± 0.09 | 6.53 ± 0.03 | 5.82 ± 0.04 | 5.91 ± 0.03 | 5.94 ± 0.02 |
| **7z** | 6.22 ± 0.10 | 6.32 ± 0.03 | 5.68 ± 0.04 | 6.00 ± 0.02 | 5.87 ± 0.09 |
| **12a** | 7.78 ± 0.10 | 7.57 ± 0.09 | 7.40 ± 0.08 | 7.29 ± 0.12 | 7.20 ± 0.08 |
| **12b** | 7.48 ± 0.08 | 7.68 ± 0.09 | 7.09 ± 0.02 | 7.19 ± 0.09 | 6.93 ± 0.09 |

Compound **6a** is a non-binder. It contains a phthalimide moiety, which is part of many synthetic allosteric modulators of muscarinic receptors. Therefore, compound **6a** was tested for possible allosteric effects. The hallmark of allosteric interaction is a change in the binding kinetics of the tracer. However, compound **6a** did not affect the dissociation rate of [^3^H]NMS binding, rendering it a pure non-binder. Except for the M_2_ receptor, two *S*-stereoisomers (**6w** and **7w**) exerted about a two-fold higher affinity than corresponding *R*-stereoisomers (**6x** and **7x**), respectively (Supplementary Information Table S2). For example, at the M_2_ receptor, 7w had 140 nM affinity and **7x** had 200 nM (only 1.4-fold lower), while at the M_5_ receptor, **7w** had 190 nM and **7x** had 550 nM (2.9-fold lower). On the other hand, affinities of stereoisomers **6o**/**6r** and **7o**/**7r** were the same at all receptor subtypes.

Predicted binding energies from the docking study (Supplementary information Table S3) do not correlate with experimentally determined affinities at any receptor subtype (“Binding energy” column). Importantly, the docking study also failed to rule out non-binders based on analysis of principal interactions with amino acid residues in the orthosteric binding site (Supporting information Table S3). Notably, all compounds, including non-binders **6a**, **6c**, **6d** and **6g** to **6l**, formed hydrogen bonds with N^6.52^.

Three replicas of short (120 ns) cMD were simulated to accommodate ligands in the orthosteric binding site and study ligand-receptor interactions. Simulations revealed profound differences in the orthosteric binding among compounds (Supplementary Information Table S4). The frequency of ligand hydrogen bonding to N^6.52^ correlated with binding affinity (Table 2) as hydrogen bonding of non-binders to N^6.52^ disappeared during cMD simulation. Except for the M_1_ receptor, experimentally determined binding affinity also correlated with the frequency of ionic binding to D^3.32^ and hydrogen bonding to T^5.42^. The multiple regression model (MRM) gave good estimates of binding affinities. The decision tree model set the threshold value of frequency of hydrogen bonding to N^6.52^ to about 15 % for all receptor subtypes. Thus, the absence or low frequency of hydrogen bonding to N^6.52^ identifies non-binders.

**Table 2.** Correlation of ligand docking and cMD trajectory analysis with experimentally determined affinities. Spearman rank correlation factors of experimentally determined pKi and predicted binding energy of top pose from ligand docking (Binding Energy), frequency of ligand interactions with individual key residues (D^3.32^ to Y^7.43^) or all key residues (MRM – multiple regression model). Last column, the decision tree threshold value of hydrogen bonding to N6.52 during cMD for binders. *, P<0.05, **, P<0.01, ***, P<0.001.

|  | Binding Energy | D<sup>3.32</sup> | Y<sup>3.33</sup> | T<sup>5.42</sup> | W<sup>6.48</sup> | Y<sup>6.51</sup> | N<sup>6.52</sup> | Y<sup>7.39</sup> | Y<sup>7.43</sup> | MRM | N<sup>6.52</sup> threshold |
| --- | --- | --- | --- | --- | --- | --- | --- | --- | --- | --- | --- |
| M<sub>1</sub> | -0.058 | 0.158 | -0.187 | 0.285 | -0.052 | 0.192 | 0.599*** | -0.074 | -0.018 | 0.566*** | 0.154 |
| M<sub>2</sub> | -0.133 | 0.311* | -0.131 | 0.298* | 0.116 | 0.234 | 0.597*** | 0.017 | 0.044 | 0.554*** | 0.157 |
| M<sub>3</sub> | 0.135 | 0.341* | -0.090 | 0.310* | 0.032 | 0.208 | 0.529*** | 0.040 | 0.051 | 0.545*** | 0.141 |
| M<sub>4</sub> | -0.201 | 0.545*** | 0.010 | 0.349* | 0.094 | 0.262 | 0.536*** | 0.063 | 0.106 | 0.539*** | 0.157 |
| M<sub>5</sub> | -0.093 | 0.292* | -0.082 | 0.285 | 0.059 | 0.190 | 0.491*** | 0.026 | 0.009 | 0.503*** | 0.161 |

Root mean square fluctuations (RMSF) of individual receptor residues during MD were calculated to analyse putative effects of ligand binding on the receptor dynamics (Supplementary Information Figure S1 and Table S6). Except for the M_5_ receptor, ligand binding did not change the RMSFs of receptors. At the M_5_ receptor, ligand binding only marginally reduced the RMSF of some parts of the receptor (Supplementary Information Table S6). These data indicate that ligand binding has no effect on receptor dynamics during MD.

The following eight descriptors were calculated. Five molecular descriptors: The distance between ketoxy oxygen (forming a hydrogen bond to N^6.52^) and nitrogen (forming an ionic bond to D^3.32^), the improper dihedral angle ketoxy oxygen – ketoxy carbon – ester oxygen – nitrogen, the molecular volume, the electrostatic potential (ESP) and the net charge of the molecule according to the NOVA force field. And three descriptors of the basic centre (residue without ethyl 4-(hexyloxy)benzoate): the volume, radius and surface area (Supplementary Information Table S5). The distance between ketoxy oxygen and nitrogen varied from 4.9 to 6.7 Å, and their improper angle varies from –77° to +70°. Except for compounds **13a,b** and **14a,b**, the molecular volume is similar, ranging from 358 to 444 Å^3^. The volume, radius and surface of the basic centre vary substantially from 104 to 185 Å^3^, 3.7 to 5.8 Å, and 327 to 339 Å^2^, respectively. The compounds ESP vary from –2.7 to +2.5 kcal/mol. The net charge of compounds is either 0 or 0.205 for compounds with charged nitrogen. The net charge of the compound molecule is the only descriptor displaying a strong correlation with compound affinity (Table 3). The compounds ESP correlated with affinity at M_4_ and M_5_ receptors and weakly correlated at M_2_ and M_3_ receptors. The distance between ketoxy oxygen and nitrogen weakly correlated with affinity only at M_1_ and M_2_ receptors. The multiple regression model (MRM) gave good estimates of binding affinities. Analysis of molecular descriptors suggests that nitrogen charge has a key role in compound affinity, as charged compounds are all binders with about 10-fold higher affinity than their uncharged analogues. The positive ESP contributes to compound affinity. Even uncharged compounds with positive ESP (**6e** and **6f**) display micromolar affinity. The optimal distance between ketoxy oxygen and nitrogen is 5.3 to 5.5 Å (compounds **7g**, **7s**, **7w**, **12a** and **12b**).

**Table 3.** Correlations of molecular descriptors with experimentally determined affinities. Spearman rank correlation factors of experimentally determined pKi and the distance between ketoxy oxygen and nitrogen (Distance), the improper dihedral angle (Angle), molecular volume (Mol. Volume), basic centre (BC) volume, radius and surface area and electrostatic potential (ESP) and net charge of the molecule. (MRM – multiple regression model). *, P<0.05, **, P<0.01, ***, P<0.001.

|  | Distance | Angle | Mol. volume | BC volume | BC radius | BC surface | ESP | Net charge | MRM |
| --- | --- | --- | --- | --- | --- | --- | --- | --- | --- |
| **M<sub>1</sub>** | 0.361* | 0.135 | 0.053 | 0.123 | -0.182 | -0.032 | 0.246 | 0.544*** | 0.677*** |
| **M<sub>2</sub>** | 0.317* | 0.199 | 0.066 | 0.160 | -0.201 | 0.011 | 0.371* | 0.613*** | 0.751*** |
| **M<sub>3</sub>** | 0.198 | 0.294 | 0.098 | 0.153 | -0.156 | 0.047 | 0.398* | 0.643*** | 0.787*** |
| **M<sub>4</sub>** | 0.282 | 0.187 | 0.159 | 0.218 | -0.140 | 0.113 | 0.464** | 0.694*** | 0.787*** |
| **M<sub>5</sub>** | 0.198 | 0.117 | 0.098 | 0.153 | -0.156 | 0.047 | 0.393** | 0.617*** | 0.729*** |

Except for the M_2_ receptor, two *S*-stereoisomers (**6w** and **7w**) exerted about a two-fold higher affinity than corresponding *R*-stereoisomers (**6x** and **7x**), respectively (Supplementary Information Table S2). In the top docking poses, both stereo isomers make ionic interaction with D^3.32^ and hydrogen bonds with N^6.52^ via ketoxy oxygen and T^5.42^ via ether oxygen (Figure 7). *S*-stereoisomers make π-π interaction with Y^3.33^, W^6.48^ and F^45.52^ in the ECL2, while *R*-stereoisomers make π-π interaction only with Y^3.33^. *S*-stereoisomers make cation-π interaction with Y^7.39^ and Y^7.43,^ while *R*-stereoisomers make cation-π interaction with Y^7.39^ and W^6.48^. On the other hand, affinities of stereoisomers **6o**/**6r** and **7o**/**7r** were the same at all receptor subtypes. The molecular descriptors are the same or similar (ketoxy oxygen to nitrogen distance, improper angle) for all pairs of stereoisomers (Supplementary Information Table S5). The distance is similar in compounds with ternary and quaternary nitrogen. The angle varies between compounds with ternary and quaternary nitrogen, as well as both selective and non-selective stereoisomers. Thus, stereoselectivity cannot be attributed to any of the descriptors.

**Figure 7.**
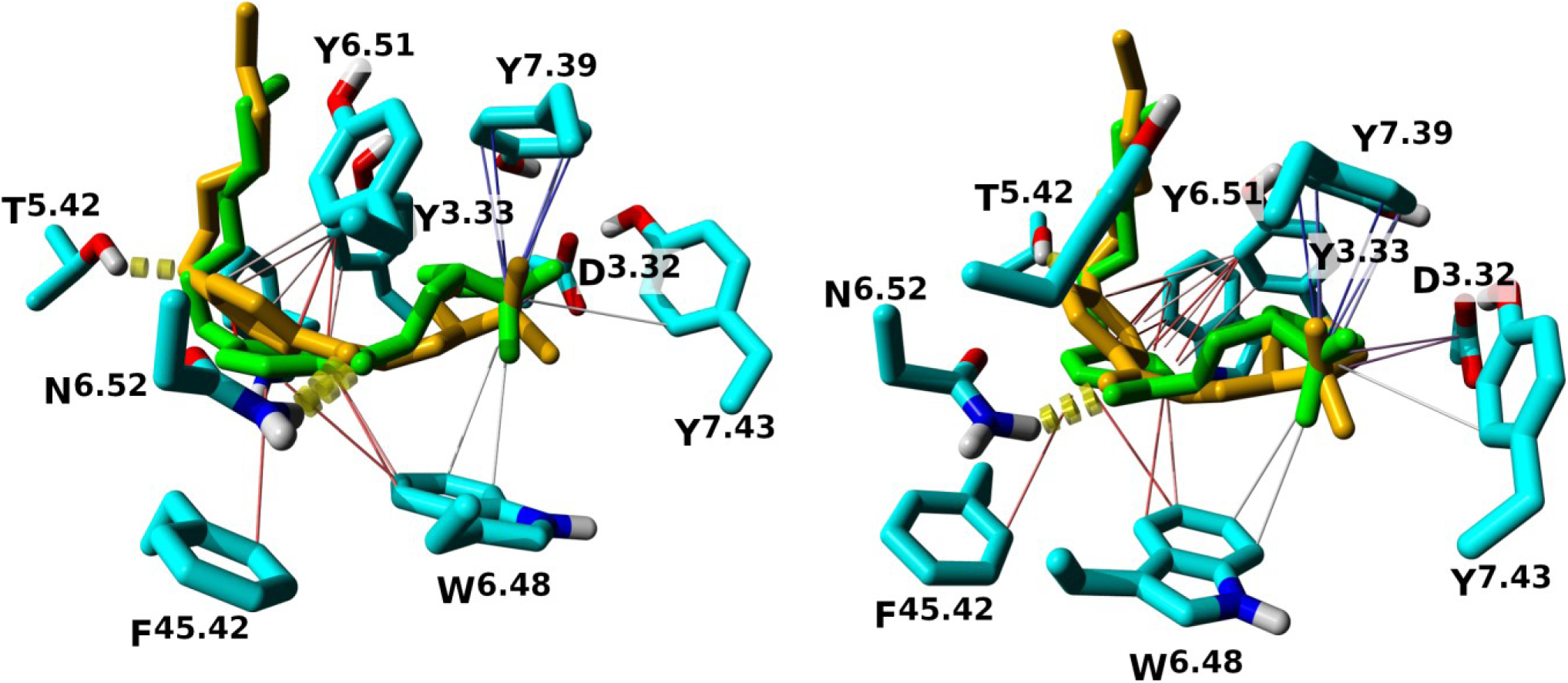
Interactions of 7w and 7x in top binding poses at the M_1_ receptor. Side (left) and extracellular (right) views of 7w (green) and 7x (gold) binding to the orhosteric binding site of the M_1_ receptor. Atoms: cyan – carbon; blue – nitrogen; red – oxygen; white – hydrogen. Interactions: dashed yellow – hydrogen bonds; purple – ionic; red – π-π; blue or white – cation-π.

#### 3.3.2. Structure-activity relationships – alkyloxy chain

It has been shown that, similarly to xanomeline[21], the hexyloxy chain contributes to the affinity of paternal compound KH-5, as a shortening of the hexyloxy chain led to a decrease in affinity[11]. Therefore, analogues of the azabicyclo[2.2.2]octan-1-ium compounds, exerting the highest affinity (**9b** and **9c**) with butyloxy (**13a** and **13b**) or methyloxy chain (**14a** and **14b**) were synthesised. Similarly, to tetrahydropyridine analogues, shortening of the hexyloxy chain results in a profound (up to 100-fold) decrease in affinity for all muscarinic receptors (Table 4).

**Table 4.**
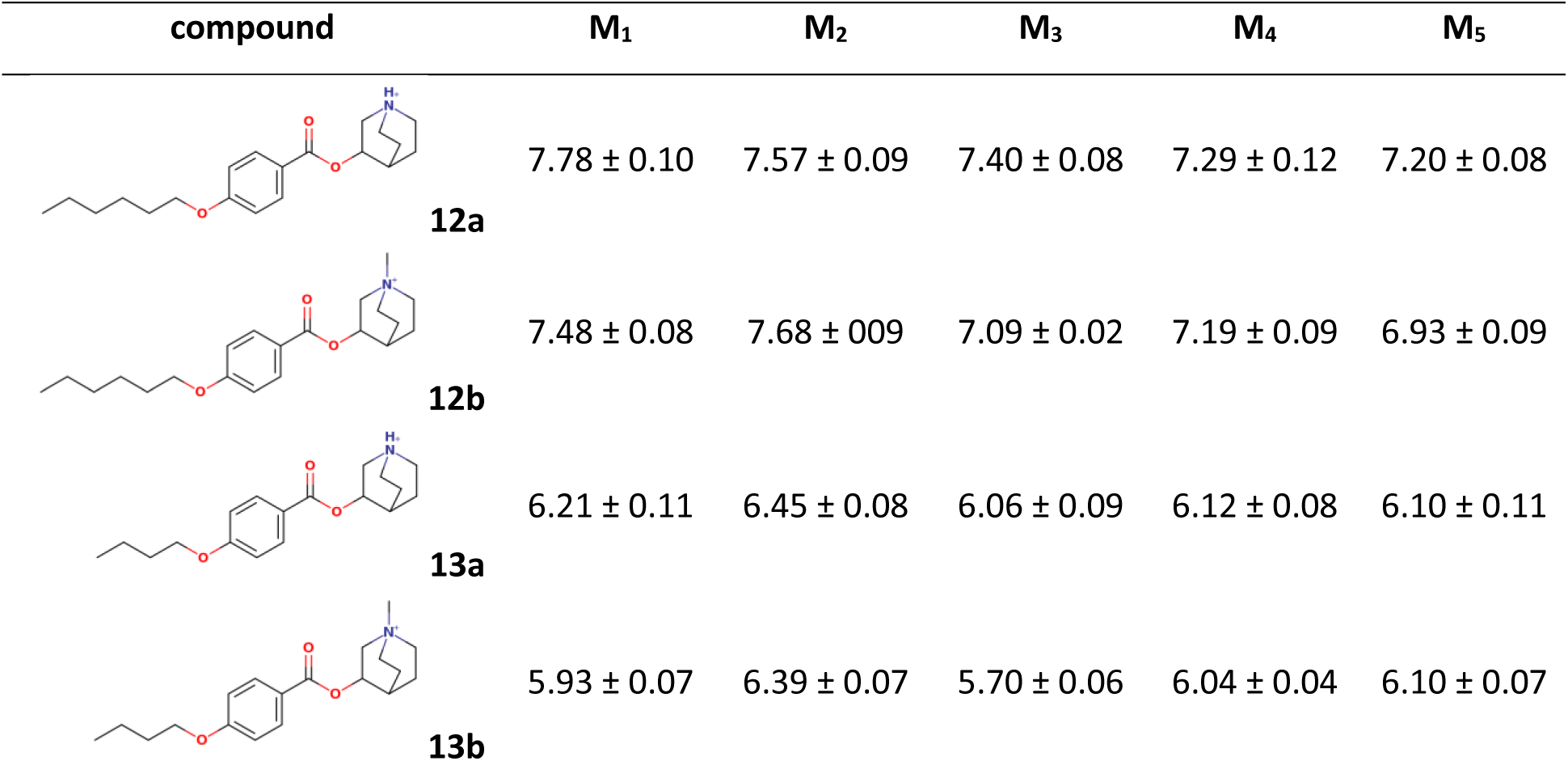
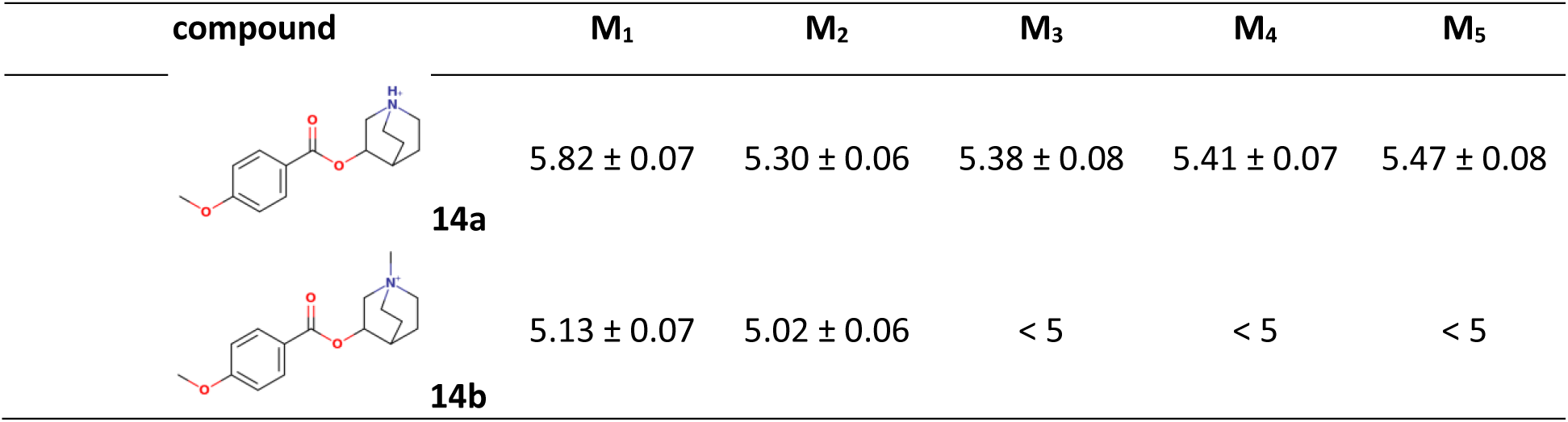
Effect of length of alkyloxy chain on affinity of azabicyclo[2.2.2]octan-1-ium analogues. Affinities are expressed as the negative decadic logarithm of the inhibition constant (pK_i_) of [^3^H]NMS binding. Data are means ± SD from 3 independent experiments.

| compound | $M_1$ | $M_2$ | $M_3$ | $M_4$ | $M_5$ |
| --- | --- | --- | --- | --- | --- |
| **12a** | $7.78 \pm 0.10$ | $7.57 \pm 0.09$ | $7.40 \pm 0.08$ | $7.29 \pm 0.12$ | $7.20 \pm 0.08$ |
| **12b** | $7.48 \pm 0.08$ | $7.68 \pm 0.09$ | $7.09 \pm 0.02$ | $7.19 \pm 0.09$ | $6.93 \pm 0.09$ |
| **13a** | $6.21 \pm 0.11$ | $6.45 \pm 0.08$ | $6.06 \pm 0.09$ | $6.12 \pm 0.08$ | $6.10 \pm 0.11$ |
| **13b** | $5.93 \pm 0.07$ | $6.39 \pm 0.07$ | $5.70 \pm 0.06$ | $6.04 \pm 0.04$ | $6.10 \pm 0.07$ |

| compound | M<sub>1</sub> | M<sub>2</sub> | M<sub>3</sub> | M<sub>4</sub> | M<sub>5</sub> |
| --- | --- | --- | --- | --- | --- |
| 14a | 5.82 ± 0.07 | 5.30 ± 0.06 | 5.38 ± 0.08 | 5.41 ± 0.07 | 5.47 ± 0.08 |
| 14b | 5.13 ± 0.07 | 5.02 ± 0.06 | < 5 | < 5 | < 5 |

#### 3.3.3. Antagonistic potency and long-term antagonism

In a previous study, KH-5 exerted sustained functional antagonism after washing with a half-life of up to 5 hours (Supplementary Information Table S1). The potency of functional antagonism and functional antagonism half-life after washing was measured for five with the highest binding affinity: **7g**, **7s**, **7w**, **12a** and **12b** (Table 5 and Figure 87). The potency of functional antagonism was determined from the inhibition of carbachol-induced accumulation of inositol phosphates by the tested compounds. The functional antagonism half-life was determined by comparison of the antagonistic effects of 10 μM compound before and after a 1-hour washing. The substitution of the tetrahydropyridine ring with azabicyclo[2.2.2]octan-1-ium yielded compounds **12a** and **12b**, and resulted in an increase in potency at all subtypes that was most profound at M_2_ (145-fold and 160-fold increase, respectively) (Table 5). However, the half-life of antagonism was severely shortened (t_1/2_ from 4 – 5 hours (KH-5) to 1 – 3 hours (**12a**)), except for the M_2_ receptor, where it was prolonged from 1.7 hours to 2.8 hours. The affinities and potencies of **12a** *N*-methyl analogue (**12b**) were lower than the affinities and potencies of **12a** at respective subtypes, except for M_2_ receptors, where they were slightly higher, correlating with lower affinities. Compound **12b** did not show long-term antagonism at any subtype.

**Table 5.**
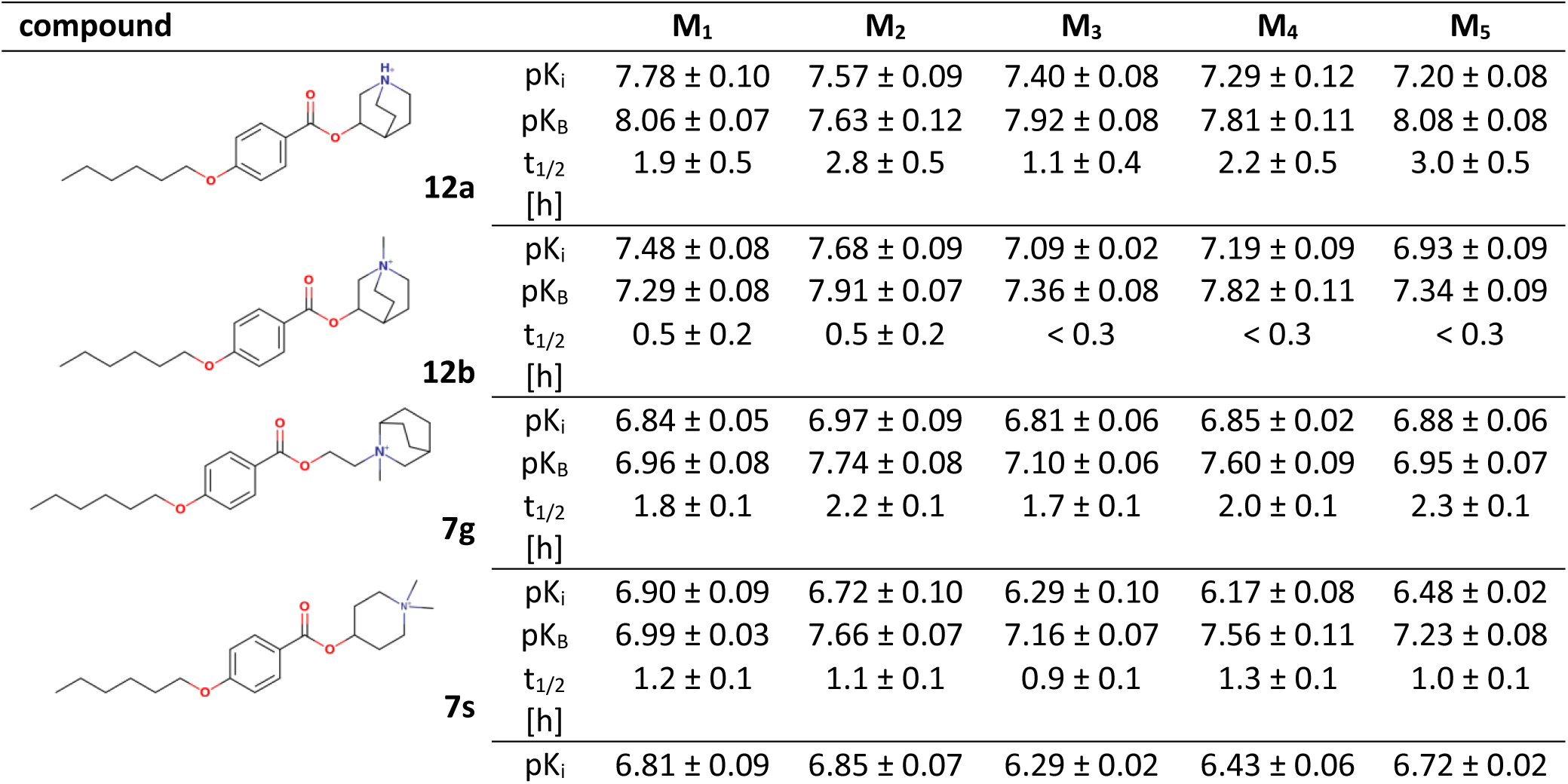
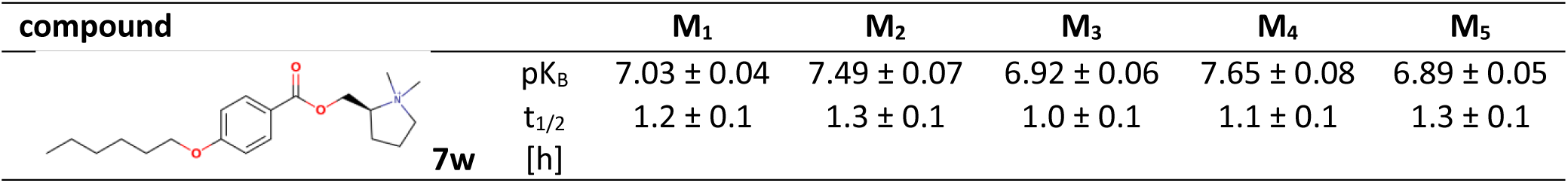
Affinity (pKi), potency (pK_B_) and antagonism half-life after washing (t_1/2_) of KH-5 analogues. Affinities are expressed as the negative decadic logarithm of the inhibition constant (pK_i_) of 1 nM [^3^H]NMS binding. Potencies are expressed as negative decadic logarithms inhibition constant (pK_B_) of antagonising carbachol-induced functional response. Residence times are expressed as half-life (t_1/2_) of antagonism of functional response induced by one-hour exposure to 10 μM compound, followed by washing. Data are means ± SD from 3 to 5 independent experiments.

Compound **7g** exerted up to 3-fold lower affinities for individual subtypes of muscarinic receptors than **12b** (Table 5). Potencies of **7g** to antagonise functional responses to carbachol were about 2-fold lower than potencies of **12b**. In comparison to KH-5, the half-life of antagonism of **7g** was also severely shortened to about 2 hours (ranging from 1.7 hours at M_3_ to 2.3 hours at M_5_ receptor). With the exception of the M_1_ receptor, affinities of **7s** and **7w** were slightly lower than affinities of **7g,** while the potencies were the same. The half-life of antagonism of compounds **7s** and **7w** was about 1 hour (ranging from 0.9 hours for **7s** at M_3_ to 1.3 hours for **7w** at M_3_ and M_5_ receptors).

## 4. Discussion and Conclusions

KH-5 is the LAMA with moderately slow dissociation (t_1/2_ up to 5 hours)[11]. The sustained activity of long-acting agonist xanomeline (t_1/2_ ≈ 30 hours) is remarkably longer[13]. Similarly, to the long-acting agonist xanomeline, the hexyloxy moiety is required for sustained effects[11]. Our recent study shows that the 4-hexyloxy position on the benzene ring is optimal for sustained activity[14]. In this study, we investigated the effects of modifications of KH-5 nitrogen heterocycle on binding affinity, potency and duration of functional antagonism. Regarding binding affinity, we show that replacing the tetrahydropyridine ring with rigid azabicyclo[2.2.2]octan-1-ium (compounds **12a** and **12b**) increased affinity (up to 250-fold) (Table 1) and potency (up to 160-fold) but substantially shortened antagonism half-life (Figure 8, Table 5). Compounds with shorter (more rigid) linkers (e.g., **7s**, **7w**) also showed reduced duration of functional antagonism, suggesting that certain flexibility is vital for prolonged activity.

**Figure 8.**
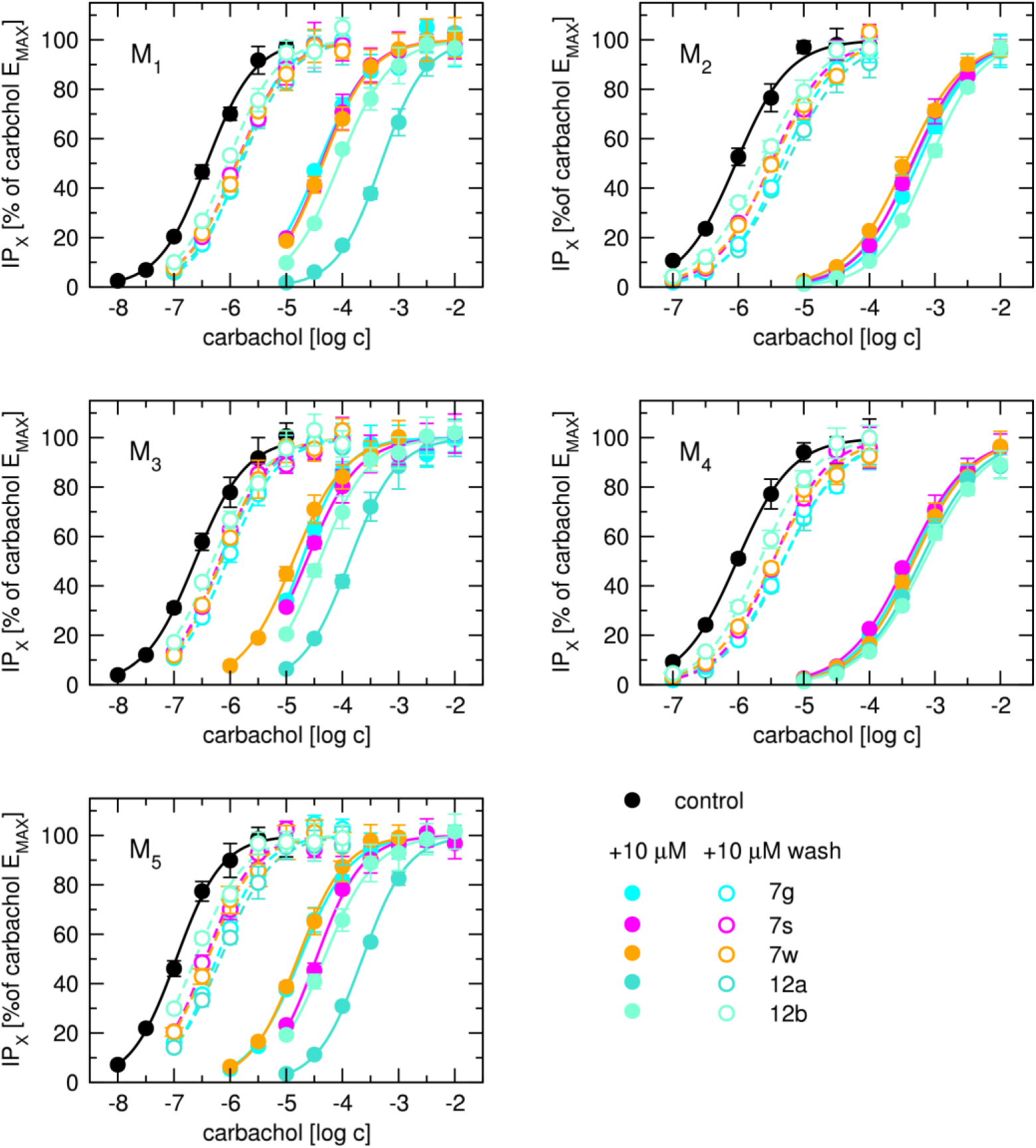
Antagonism of functional response to carbachol by compounds 7g, 7s, 7w, 12a, and 12b. The level of inositol phosphates (IP_X_) stimulated by carbachol alone (control) or in the presence (solid symbols) of 10 μM compound indicated in the legend or after incubation with compounds followed by standard washing (open symbols) is expressed as per cents of maximal response to carbachol (E_MAX_) and is plotted against the concentration of carbachol used to induce a functional response. Data are means ± SD from three independent experiments performed in quadruplicates.

We docked compounds to the orhtosteric binding site and compared computed affinity estimates with experimentally determined ones to test the docking procedure for possible future virtual screening. While docking of classical orthosteric antagonists successfully ranks affinities[22–24], docking of proposed compounds to the orthosteric binding site fails to distinguish binders from non-binders based on predicted binding energy (Table 2), indicating that some parts of the receptor outside the orthosteric binding site are involved in the binding of tested compounds. Simulations of MD provided better insights into compound binding, however, they are computationally expensive and thus unsuitable for future virtual screening. Therefore, suitable means (e.g., AI-based predictions like Boltz-2) need to be found.

The potency of functional antagonism of tested compounds (**7g**, **7s**, **7w**, **12a** and **12b**) is greater than their binding affinity. We speculated that this discrepancy may indicate that these compounds are PAM-antagonists[25]. If true, then these compounds also bind to the allosteric site. And that could be the reason why binding energy prediction from docking to the orthosteric binding site fails. However, allosteric antagonism is insurmountable and causes a decrease in maximal response to an agonist. No decrease in E_MAX_ was observed for these compounds. Thus, an alternative explanation is needed. The binding equilibrium of antagonists with slow kinetics may not be reached in functional assays. In such a case, however, antagonist potency would be underestimated. Delineation of the mechanism of action of these compounds will require detailed studies that are beyond the scope of this article. Anyway, in accordance with the current pharmacology of muscarinic receptors[26], molecular dynamics (MD) simulations revealed that hydrogen bonding to N^6.52^ and salt bridges to D^3.32^ stabilised ligand-receptor interactions in binders but not in non-binders. Further analysis of MD trajectories identified hydrogen bond stability with N^6.52^ may serve as a discriminator of binders and non-binders (Table 2).

Regarding structure-activity relationships, the study reveals that charged nitrogen is essential for compound affinity, as quaternary ammonium derivatives showed about 10-fold higher affinities than tertiary amines (e.g., **7g** vs. **6g**) (Table 1 Overall, none of the compounds displayed substantial binding selectivity. In addition, compounds with quaternary nitrogen had higher affinity than analogues with tertiary nitrogen (e.g. **7g** vs **6g**). ^Ho^wever, **12b** (excluding M_2_) had slightly lower affinities than **12a**. The highest binding affinity, about 30 nM, was detected for compounds **12a** and **12b**, with **12a** having a higher affinity than **12b** at all subtypes except M^2^ (Table 1). In comparison to parental compounds, it is increased up to 250-fold for hydrogen derivates (**12a** over OT-231) and up to 8-fold for methyl derivates (**12b** over KH-5). These compounds contain an azabicyclo[2.2.2]octan-1-ium moiety attached at position 3 to the rest of the molecule. Attaching azabicyclo[2.2.2]octan-1-ium via nitrogen (compound **6b**) resulted in affinities for muscarinic receptors about ^1^0-fold lower than **12a**, except for M_4_ and M_5_ receptors, where the affinity of **6b** was only slightly lower. Usin^g^ 2-azabicyclo[2.2.2]octan-2-yl (compound **6g**) resulted in the loss of affinity for muscarinic receptors. Charged nitrogen-containing compound **7g** regained affinities to a similar extent as for compound **7b**.

Table 1). Analysis of molecular descriptors shows that net charge and electrostatic potential strongly correlate with affinity, emphasising the importance of cationic interactions in the orthosteric site (Table 3). Shortening the hexyloxy chain to butyloxy or methoxy reduced affinity by up to 100-fold and abolished sustained antagonism, confirming hexyloxy necessity for sustained action (Table 4).

Only five LAMAs are clinically approved (tiotropium, glycopyrronium, aclidinium, umeclidinium and revefenacin)[7,8]. All of them display M_3_ kinetic selectivity and are suitable for the treatment of COPD. Therefore, novel LAMAs with different selectivity profiles are desired for treatment of other conditions, like epilepsy seizures, cholinergic dystonia or Parkinson’s disease[5,9,10].

Muscarinic antagonists with long residence times rely on slow dissociation from receptors that is warranted by several features/mechanisms. All LAMAs share a common quaternary ammonium structure, which is essential for ionic interaction with the conserved D^3.32^ in the orthosteric binding site that anchors the molecule within the site[27]. In tiotropium, the α-hydroxy group is probably the most important structural feature for achieving extended residence times. It forms hydrogen bonding to a key residue, N^6.52^, resulting in a snap-lock that prevents rapid dissociation[28]. The presence of aromatic rings (e.g. thiophene in tiotropium, thiazole in aclidinium, benzyl carbamate in umeclidinium) contributes to slow dissociation through π-π stacking interactions with the tyrosine cage surrounding the orthosteric site. These interactions help stabilise binding and create additional energy barriers that must be overcome during dissociation[8,29].

The investigation of the antagonist unbinding from muscarinic receptors showed an important role of the exit channel and binding pocket vestibule in antagonist residence time. Extracellular vestibule electrostatics and flexibility co-determine binding kinetics[30]. Tiotropium binding to the extracellular vestibule can be considered as binding to the allosteric site, as tiotropium antagonism is insurmountable [31]. These secondary binding sites may create additional energy barriers that slow dissociation. Overall, binding affinities predicted from docking are underestimated, indicating a substantial contribution of interactions in the extracellular vestibule to the affinity of synthesised compounds. In turn lack of correlation between predicted and measured affinities suggests these contributions vary among tested compounds. Unfortunately, simulation of extended MD allowing for ligand dissociation is computationally infeasible for more than several ligands.

Slow dissociation may also be mediated by other hydrophobic interactions[2,30]. Our previous studies confirmed that the hydrophobic hexyloxy chain in KH-5 analogues contributes to sustained functional antagonism, akin to the wash-resistant agonism of xanomeline[12]. However, rigidifying the linker or altering the heterocycle disrupted this balance, shortening duration despite higher affinity. The reliance on a charged nitrogen aligns with known muscarinic receptor pharmacology, where antagonists typically engage D^3.32^ via ionic bonds[32]. While compounds **12a** and **12b** achieved high affinity, their shorter half-life highlights a design challenge: optimising heterocyclic rigidity improves binding but may hinder the flexibility needed for prolonged action. This aligns with tiotropium, where a flexible linker and hydrophobic 2-(thiophen-2-ylmethyl)thiophene group synergise for long residency[30].

In summary, this study demonstrates that structural modifications to heterocycle KH-5 and the linker can enhance affinity but often at the cost of sustained antagonism. The findings underscore that i) the hexyloxy chain is a prerequisite for prolonged activity. ii) The charged nitrogen is essential for high-affinity binding. iii) The balance in the rigidity of the basic centre and flexibility of the molecule is needed in future compound design. Future work could explore hybrid structures combining a hexyloxy moiety with bicyclic heterocycles optimised for both affinity and kinetic retention. Another approach may be functional modifications of the basic centre and benzene ring aimed at strengthening hydrogen bonding to N^6.52^ to simulate snap-lock of tiotropium and cationic interaction with D^3.32^.

## 5. Materials and Methods

### 5.1. Synthesis

#### 5.1.1. General information

All solvents and chemical reagents were used in the highest available purity without further purification, and they were purchased from Sigma-Aldrich (Prague, Czech Republic). The reactions were monitored by thin-layer chromatography (TLC) on silica gel plates (60 F254, Merck, Prague, Czech Republic), and the spots were visualised by ultraviolet light (254 nm). Purification of crude products was carried out using columns of silica gel (silica gel 100, 0.063-0.200 mm, 70-230 mesh ASTM, Fluka, Prague, Czech Republic). NMR spectra were recorded in deuterated chloroform (CDCl_3_), deuterated methanol (CD_3_OD) and deuterated dimethyl sulfoxide (DMSO-*d*_6_) on a Varian S500 spectrometer. Chemical shifts (*δ*) are reported in parts per million (ppm), and spin multiplicities are given as first-order multiplets [singlet (s), doublet (d), triplet (t), quartet (q), quintet or pentet (p), septet or heptet (hept), or more complex patterns involving different coupling constants [multiplet (m), dd, dt, dq, ddd, ddt, dtt, td, tt, tdd, qd, etc.]. Coupling constants (*J*) are reported in Hz. The synthesised compounds were analysed by the LC-MS system consisting of UHPLC Dionex Ultimate 3000 coupled with Q Exactive™ Plus mass spectrometer to obtain high-resolution mass spectra (Thermo Fisher Scientific, Bremen, Germany). All the final compounds exerted purity higher than 95% (UV, LC-MS, uncalibrated) except for **12b** (92% purity). Syntheses of intermediates **3**, **4** and **8k**, **l** are listed in the Supplementary Information.

**<u>General procedure A:</u>** To a solution of carboxylic acid **4** (1.0 eq) in dry DCM (0.05 M), corresponding alcohol (2-bromoethanol or **8k**-**z**; 1.1 eq) was added, DMAP (0.1 eq) and a solution of DCC (1.2 eq) in dry DCM (1.0 M). The resulting mixture was stirred overnight at RT. The solid was filtered off, and the solvent was evaporated under reduced pressure. The crude product was purified by column chromatography, affording **5** and **6k-z**.

**<u>General procedure B:</u>**To a solution of bromide **5** (1.0 eq) and corresponding amine (1.1 eq) in dry CH_3_CN (0.2 M), K_2_CO_3_ (1.2 eq) was added. The resulting mixture was stirred at 70 °C overnight. The solid was filtered off, and the solvent was evaporated under reduced pressure. The crude product was purified by column chromatography, affording **6a-j**.

**<u>General procedure C:</u>** To a solution of the corresponding starting compound **6a**-**z** (1.0 eq) in CH_3_CN (0.2 M), CH_3_I (1.5 eq) was added. The mixture was stirred overnight at RT. The solvent was evaporated under reduced pressure, affording **7e-z** and **9c**, **10c** and **11c**. The crude products **7e-z** were purified by crystallisation from acetonitrile.

**<u>General procedure D</u>:** To a solution of 3-quinuclidinol (1.0 eq) in acetonitrile (0.05 M), K_2_CO_3_ (2.0 eq) and corresponding alkyloxybenzoyl chloride (1.0 eq) were added. The mixture was stirred overnight at RT. The solid was filtered off, and the solvent was evaporated under reduced pressure. The crude product was dissolved in dimethyl ether and a small amount of water. The resulting solution was made alkaline with 3M NaOH and extracted with ether. The organic layers were combined and dried over anhydrous MgSO_4_. The solid was filtered off, and the solvent was evaporated under reduced pressure. The crude product (**9a**, **10a**, **11a**) was used in the following reaction without any purification.

**General procedure E**: HCl gas was bubbled through a solution of the corresponding base in anhydrous pentane. The solvent was evaporated under reduced pressure. The residue was washed with anhydrous ether and dried, affording **9b**, **10b** and **11b**.

#### 5.1.2. Synthesis of intermediate 5

*2-bromoethyl 4-(hexyloxy)benzoate (**5**)* was prepared by general procedure **A** from acid **4** and 2-bromoethanol in 65% yield as a white amorphous solid. ^1^H NMR (500 MHz, CDCl_3_): *δ* 8.04–7.97 (m, 2H), 6.94–6.89 (m, 2H), 4.59 (t, *J* = 6.1 Hz, 2H), 4.01 (t, *J* = 6.6 Hz, 2H), 3.63 (t, *J* = 6.2 Hz, 2H), 1.84–1.76 (m, 2H), 1.51–1.42 (m, 2H), 1.38–1.31 (m, 4H), 0.94–0.88 (m, 3H) ppm. ^13^C NMR (126 MHz, CDCl_3_): *δ* 165.9, 163.3, 131.8, 121.7, 114.2, 68.3, 63.9, 31.6, 29.1, 29.0, 25.7, 22.6, 14.0 ppm.

#### 5.1.3. Synthesis of final compounds

*2-(1,3-Dioxo-2,3-dihydro-1H-isoindol-2-yl)ethyl 4-(pentyloxy)benzoate (**6a**)*: Prepared from bromide **5** and corresponding amines **A** by general procedure **B**. The column chromatography used a mobile phase PE/EA (1/1). ^1^H NMR (500 MHz, CDCl_3_): *δ* 7.94–7.90 (m, 2H), 7.88–7.83 (m, 2H), 7.73–7.70 (m, 2H), 6.89–6.85 (m, 2H), 4.50 (t, *J* = 5.8, 4.9 Hz, 2H), 4.10 (t, *J* = 5.3 Hz, 2H), 3.98 (t, *J* = 6.6 Hz, 2H), 1.82–1.74 (m, 2H), 1.49–1.41 (m, 2H), 1.36–1.31 (m, 4H), 0.92–0.88 (m, 3H) ppm. ^13^C NMR (126 MHz, CDCl_3_): *δ* 168.2, 167.9, 166.3, 163.2, 134.5, 134.1, 132.8, 132.2, 131.9, 131.9, 123.6, 123.5, 122.0, 114.3, 68.3, 62.1, 37.2, 31.7, 29.2, 25.8, 22.7, 14.2 ppm. HRMS [M+H]^+^: 396.17996 (calc. for [C_23_H_26_NO_5_]^+^: 396.18055), 98 % purity. **6a** was isolated as a white solid (melting point 88–90 °C) in 64% yield (77 mg).

*1-{2-[4-(Hexyloxy)benzoyloxy]ethyl}-1-azabicyclo[2.2.2]octan-1-ium (**6b**)*: Prepared from bromide **5** and corresponding amines **A** by general procedure **B**. The column chromatography used a mobile phase PE/EA (1/1). ^1^H NMR (500 MHz, DMSO-*d*_6_): *δ* 7.96–7.90 (m, 2H), 7.08–7.03 (m, 2H), 4.68–4.61 (m, 2H), 4.05 (t, *J* = 6.5 Hz, 2H), 3.66–3.61 (m, 2H), 3.58–3.53 (m, 6H), 2.10–2.04 (m, 1H), 1.88 (ddt, *J* = 8.4, 5.3, 3.1 Hz, 6H), 1.76–1.68 (m, 2H), 1.44–1.37 (m, 2H), 1.34–1.25 (m, 4H), 0.91–0.84 (m, 3H) ppm. ^13^C NMR (126 MHz, DMSO-*d*_6_): *δ* 164.8, 162.9, 131.5, 121.1, 114.6, 67.9, 62.0, 57.7, 54.3, 31.0, 28.4, 25.1, 23.4, 22.1, 19.0, 13.9 ppm. HRMS [M+H]^+^: 360.25299 (calc. for [C_22_H_34_NO_3_]^+^: 360.25284), 99 % purity. **6b** was isolated as a white solid (melting point 182– 183 °C) in 51% yield (56 mg).

*2-(2,5-Dioxopyrrolidin-1-yl)ethyl 4-(hexyloxy)benzoate (**6c**)*: Prepared from bromide **5** and corresponding amines **A** by general procedure **B**. The column chromatography used a mobile phase PE/EA (1/1). ^1^H NMR (500 MHz, CDCl_3_): *δ* 7.94–7.89 (m, 2H), 6.91–6.86 (m, 2H), 4.42 (t, *J* = 5.3 Hz, 2H), 3.99 (t, *J* = 6.6 Hz, 2H), 3.91 (t, *J* = 5.3 Hz, 2H), 1.83–1.74 (m, 2H), 1.49–1.42 (m, 2H), 1.36–1.31 (m, 4H), 0.93–0.87 (m, 3H) ppm. ^13^C NMR (126 MHz, CDCl_3_): *δ* 177.1, 166.3, 163.3, 131.9, 121.9, 114.3, 68.3, 61.4, 38.1, 31.7, 29.2, 28.3, 25.8, 22.7, 14.1 ppm. HRMS [M+H]^+^: 348.17957 (calc. for [C_19_H_26_NO_5_]^+^: 348.18055), 98 % purity. **6c** was isolated as an oil in 45% yield (45 mg).

*2-{3,5-Dioxo-10-oxa-4-azatricyclo[5.2.1.0^2,6^]dec-8-en-4-yl}ethyl 4-(hexyloxy)benzoate (**6d**)*: Prepared from bromide **5** and corresponding amines **A** by general procedure **B**. The column chromatography used a mobile phase PE/EA (1/1). ^1^H NMR (500 MHz, CDCl_3_): *δ* 7.94–7.90 (m, 2H), 6.90–6.86 (m, 2H), 6.50 (d, *J* = 1.0 Hz, 2H), 5.25 (d, *J* = 0.9 Hz, 2H), 4.41 (t, *J* = 5.3 Hz, 2H), 3.99 (t, *J* = 6.6 Hz, 2H), 3.89 (t, *J* = 5.4 Hz, 2H), 1.82–1.75 (m, 2H), 1.61–1.52 (m, 2H), 1.50–1.41 (m, 2H), 1.36–1.32 (m, 4H), 0.93–0.89 (m, 3H) ppm. ^13^C NMR (126 MHz, CDCl_3_): *δ* 176.1, 166.2, 163.2, 136.7, 131.9, 122.0, 114.2, 81.1, 68.3, 61.1, 47.6, 31.7, 29.8, 29.2, 25.8, 22.7, 14.2 ppm. HRMS [M+H]^+^: 414.18958 (calc. for [C_23_H_27_NO_6_]^+^: 414.19111), 95 % purity. **6d** was isolated as an oil in 7% yield (8 mg).

*2-[(2R)-2-Methylpiperidin-1-yl]ethyl 4-(hexyloxy)benzoate (**6e**)*: Prepared from bromide **5** and corresponding amines **A** by general procedure **B**. The column chromatography used a mobile phase PE/EA (1/1). ^1^H NMR (500 MHz, CDCl_3_): *δ* 7.92–7.88 (m, 2H), 6.85–6.81 (m, 2H), 4.33 (t, *J* = 6.1 Hz, 2H), 3.93 (t, 2H), 3.00–2.93 (m, 1H), 2.91–2.85 (m, 1H), 2.81–2.73 (m, 1H), 2.37–2.28 (m, 2H), 1.76–1.69 (m, 2H), 1.60 (t, *J* = 10.2, 3.8, 2.0 Hz, 1H), 1.57–1.51 (m, 2H), 1.50–1.44 (m, 1H), 1.42–1.36 (m, 2H), 1.29–1.25 (m, 4H), 1.24–1.16 (m, 2H), 1.06 (d, *J* = 6.2 Hz, 3H), 0.86–0.81 (m, 3H) ppm. ^13^C NMR (126 MHz, CDCl_3_): *δ* 166.5, 163.1, 131.7, 122.6, 114.2, 68.4, 62.3, 56.1, 53.5, 52.2, 34.9, 31.7, 29.2, 26.3, 25.8, 24.2, 22.7, 19.8, 14.2 ppm. HRMS [M+H]^+^: 348.25320 (calc. for [C_21_H_34_NO_3_]^+^: 348.25332), 97 % purity. **6e** was isolated as an oil in 42% yield (111 mg).

*2-[(2S)-2-Methylpiperidin-1-yl]ethyl 4-(hexyloxy)benzoate (**6f**)*: Prepared from bromide **5** and corresponding amines **A** by general procedure **B**. The column chromatography used a mobile phase PE/EA (1/1). ^1^H NMR (500 MHz, CDCl_3_): *δ* 7.92–7.87 (m, 2H), 6.86–6.80 (m, 2H), 4.33 (t, *J* = 6.2 Hz, 2H), 3.93 (t, *J* = 6.6 Hz, 2H), 3.01–2.95 (m, 1H), 2.92–2.85 (m, 1H), 2.82–2.74 (m, 1H), 2.41–2.28 (m, 2H), 1.76–1.68 (m, 2H), 1.64–1.58 (m, 1H), 1.58–1.51 (m, 2H), 1.42–1.36 (m, 2H), 1.30–1.25 (m, 4H), 1.25–1.17 (m, 3H), 1.06 (d, *J* = 6.2 Hz, 3H), 0.86–0.82 (m, 3H) ppm. ^13^C NMR (126 MHz, CDCl_3_): *δ* 166.4, 163.0, 131.6, 122.4, 114.1, 68.2, 62.1, 56.0, 53.4, 52.1, 34.7, 31.6, 31.6, 29.7, 29.1, 26.1, 25.7, 24.0, 22.6, 19.6, 14.0 ppm. HRMS [M+H]^+^: 348.25336 (calc. for [C_21_H_34_NO_3_]^+^: 348.25332), 97 % purity. **6f** was isolated as an oil in 60% yield (159 mg).

*2-{2-Azabicyclo[2.2.2]octan-2-yl}ethyl 4-(hexyloxy)benzoate (**6g**)*: Prepared from bromide **5** and corresponding amines **A** by general procedure **B**. The column chromatography used a mobile phase PE/EA (1/1). ^1^H NMR (500 MHz, CDCl_3_): *δ* 7.92–7.87 (m, 2H), 6.86–6.80 (m, 2H), 4.33 (t, *J* = 6.2 Hz, 2H), 3.93 (t, *J* = 6.6 Hz, 2H), 3.01–2.95 (m, 1H), 2.92–2.85 (m, 1H), 2.82–2.74 (m, 1H), 2.41–2.28 (m, 2H), 1.76–1.68 (m, 2H), 1.64–1.58 (m, 1H), 1.58–1.51 (m, 2H), 1.42–1.36 (m, 2H), 1.30–1.25 (m, 4H), 1.25–1.17 (m, 3H), 1.06 (d, *J* = 6.2 Hz, 3H), 0.86–0.82 (m, 3H) ppm. ^13^C NMR (126 MHz, CDCl_3_): *δ* 166.4, 163.2, 131.8, 131.7, 122.3, 114.3, 114.2, 68.4, 62.8, 56.9, 55.1, 51.9, 50.8, 31.7, 29.8, 29.2, 25.8, 25.4, 24.4, 23.9, 22.7, 14.1 ppm. HRMS [M+H]^+^: 360.25342 (calc. for [C_22_H_34_NO_3_]^+^: 360.25332), 95 % purity. **6g** was isolated as an oil in 35% yield (97 mg).

*2-(2,5-Dihydro-1H-pyrrol-1-yl)ethyl 4-(hexyloxy)benzoate (**6h**)*: Prepared from bromide **5** and corresponding amines **A** by general procedure **B**. The column chromatography used a mobile phase PE/EA (1/1). ^1^H NMR (500 MHz, CDCl_3_): *δ* 8.00–7.95 (m, 2H), 6.92–6.87 (m, 2H), 5.79 (s, 2H), 4.44 (t, *J* = 5.9 Hz, 2H), 4.00 (t, *J* = 6.6 Hz, 2H), 3.65 (s, 4H), 3.07 (t, *J* = 5.9 Hz, 2H), 1.82–1.74 (m, 2H), 1.50–1.42 (m, 2H), 1.36–1.30 (m, 4H), 0.92–0.88 (m, 3H) ppm. ^13^C NMR (126 MHz, CDCl_3_): *δ* 166.3, 163.1, 131.7, 127.4, 122.3, 114.1, 108.6, 68.2, 63.9, 61.9, 60.4, 54.4, 31.7, 31.6, 29.7, 29.1, 25.7, 22.6, 14.0 ppm. HRMS [M+H]^+^: 318.20624 (calc. for [C_19_H_28_NO_3_]^+^: 318.20637), 97 % purity. **6h** was isolated as an oil in 31% yield (75 mg).

*2-(Pyrrolidin-1-yl)ethyl 4-(hexyloxy)benzoate (**6j**)*: Prepared from bromide **5** and corresponding amines **A** by general procedure **B**. The column chromatography used a mobile phase PE/EA (1/1). ^1^H NMR (500 MHz, DMSO-*d*_6_): *δ* 7.91–7.86 (m, 2H), 7.05–7.01 (m, 2H), 4.32 (t, *J* = 5.8 Hz, 2H), 4.04 (t, *J* = 6.5 Hz, 2H), 2.77 (t, *J* = 5.8 Hz, 2H), 2.56–2.50 (m, 3H), 1.76–1.65 (m, 6H), 1.44–1.37 (m, 2H), 1.33–1.27 (m, 4H), 1.23 (s, 1H), 0.90–0.85 (m, 3H) ppm. ^13^C NMR (126 MHz, DMSO-*d*_6_): *δ* 165.8, 163.1, 131.7, 122.3, 114.9, 68.3, 63.9, 54.5, 54.4, 31.4, 29.0, 25.56, 23.7, 22.5, 14.4 ppm. HRMS [M+H]^+^: 320.22247 (calc. for [C_19_H_26_NO_5_]^+^: 320.22202), 98 % purity. **6j** was isolated as an oil in 16% yield (39 mg).

*2-(2,3-Dihydro-1H-isoindol-2-yl)ethyl 4-(hexyloxy)benzoate (**6k**)*: Prepared from carboxylic acid **4** and alcohol **8k** by general procedure **A**. The column chromatography used a mobile phase PE/EA (1/1). ^1^H NMR (500 MHz, CDCl_3_): *δ* 8.03–8.00 (m, 2H), 7.22–7.17 (m, 4H), 6.94–6.89 (m, 2H), 4.51 (t, *J* = 5.9 Hz, 2H), 4.07 (s, 4H), 4.01 (t, *J* = 6.6 Hz, 2H), 3.15 (t, *J* = 5.9 Hz, 2H), 1.83–1.77 (m, 2H), 1.50–1.43 (m, 2H), 1.38–1.33 (m, 4H), 0.93–0.89 (m, 3H) ppm. ^13^C NMR (126 MHz, CDCl_3_) δ 166.5, 163.2, 140.0, 131.8, 126.9, 122.5, 122.4, 114.2, 68.4, 63.9, 59.7, 54.5, 31.7, 29.2, 25.8, 22.7, 14.2 ppm. HRMS [M+H]^+^: 368.22144 (calc. for [C_23_H_30_NO_3_]^+^: 368.22202), 95 % purity. **6k** was isolated as an oil in 64% yield (472 mg).

*2-(1,2,3,4-Tetrahydroquinolin-1-yl)ethyl 4-(hexyloxy)benzoate (**6l**)*: Prepared from the carboxylic acid **4** and alcohol **8l** by general procedure **A**. The column chromatography used a mobile phase PE/EA (1/1). ^1^H NMR (500 MHz, CDCl_3_): *δ* 7.97–7.94 (m, 2H), 7.07 (td, *J* = 7.8, 1.7 Hz, 1H), 6.96 (dd, *J* = 7.6, 1.6 Hz, 1H), 6.91–6.89 (m, 2H), 6.71 (d, *J* = 8.2 Hz, 1H), 6.61–6.58 (m, 1H), 4.49–4.46 (m, 2H), 4.01 (t, *J* = 6.6 Hz, 2H), 3.66 (t, *J* = 6.2 Hz, 2H), 3.42–3.40 (m, 2H), 2.77 (t, *J* = 6.4 Hz, 2H), 1.99–1.94 (m, 2H), 1.80 (dq, *J* = 8.5, 6.6 Hz, 2H), 1.47 (p, *J* = 7.6 Hz, 2H), 1.38–1.33 (m, 5H), 0.95–0.89 (m, 4H) ppm. ^13^C NMR (126 MHz, CDCl_3_): *δ* 166.6, 163.2, 145.0, 131.8, 129.5, 127.3, 122.5, 122.3, 116.1, 114.2, 110.8, 68.4, 61.7, 50.4, 50.2, 31.7, 29.2, 28.3, 25.8, 22.7, 22.4, 14.2 ppm. HRMS [M+H]^+^: 382.23749 (calc. for [C_24_H_32_NO_3_]^+^: 382.23740), 98 % purity. **6l** was isolated as an oil in 18% yield (111 mg).

*(1-Methylpiperidin-2-yl)methyl 4-(hexyloxy)benzoate (**6m**)*: Prepared from the carboxylic acid **4** and 1-methyl-2-piperidinementhanol (**8m**) by general procedure **A**. The column chromatography used a mobile phase DCM/MeOH/NH_3_ (20/1/0.1). ^1^H NMR (500 MHz, CDCl_3_): *δ* 7.99–7.96 (m, 2H), 6.91–6.89 (m, 2H), 4.44–4.39 (m, 1H), 4.34–4.30 (m, 1H), 4.00 (t, *J* = 6.6 Hz, 2H), 2.92 (tt, *J* = 9.0, 3.6 Hz, 1H), 2.45–2.42 (m, 3H), 2.34–2.30 (m, 1H), 2.22–2.15 (m, 1H), 1.86–1.74 (m, 4H), 1.71–1.63 (m, 2H), 1.51–1.42 (m, 3H), 1.38–1.28 (m, 5H), 0.93–0.86 (m, 3H) ppm. ^13^C NMR (126 MHz, CDCl_3_): *δ* 166.2, 163.1, 131.7, 122.2, 114.1, 68.2, 66.6, 63.0, 57.1, 43.7, 31.6, 29.1, 25.7, 23.7, 22.6, 14.0 ppm. HRMS [M+H]^+^: 334.23730 (calc. for [C_20_H_32_NO_3_]^+^: 334.23767), 95 % purity. **6m** was isolated as an amorphous solid in 23% yield (107 mg).

*[(2S)-1-Methylpiperidin-2-yl]methyl 4-(hexyloxy)benzoate (**6n**):* Prepared from the carboxylic acid **4** and (*S*)-(1-methylpiperidin-2-yl)methanol (**8n**) by general procedure **A**. The column chromatography used a mobile phase DCM/MeOH/NH_3_ (20/1/0.1). ^1^H NMR (500 MHz, CDCl_3_): *δ* 7.99–7.96 (m, 2H), 6.91–6.88 (m, 2H), 4.41 (dt, *J* = 11.2, 4.3 Hz, 1H), 4.32 (dt, *J* = 12.1, 4.2 Hz, 1H), 4.00 (t, *J* = 6.6 Hz, 2H), 2.94–2.90 (m,1H), 2.46–2.43 (m,2H), 2.35–2.30 (m, 1H), 2.22–2.15 (m, 1H), 1.86–1.76 (m, 4H), 1.70–1.63 (m, 2H), 1.56–1.42 (m, 3H), 1.37–1.28 (m, 5H), 0.92–0.88 (m, 3H) ppm. ^13^C NMR (126 MHz, CDCl_3_): *δ* 166.4, 163.2, 131.8, 122.3, 114.3, 68.4, 66.8, 63.2, 57.2, 43.8, 31.7, 29.2, 25.8, 23.8, 22.7, 14.2 ppm. HRMS [M+H]^+^: 334.23730 (calc. for [C_20_H_32_NO_3_]^+^: 334.23767), 97% purity. **6n** was isolated as an amorphous solid in 28% yield (135 mg).

*[(3S)-1-Methylpiperidin-3-yl]methyl 4-(hexyloxy)benzoate (**6o**)*: Prepared from carboxylic acid **4** and (*S*)-1-methyl-3-(hydroxymethyl)piperidine (**8o**). The column chromatography used a mobile phase DCM/MeOH/NH_3_ (20/1/0.1). ^1^H NMR (500 MHz, CDCl_3_): *δ* 7.97–7.94 (m, 2H), 6.91–6.87 (m, 2H), 4.19 (dd, *J* = 11.0, 5.5 Hz, 1H), 4.11 (dd, *J* = 11.0, 7.3 Hz, 1H), 4.00 (t, *J* = 6.6 Hz, 2H), 3.03–3.00 (m, 1H), 2.89 (d, *J* = 11.3 Hz, 1H), 2.34 (s, 3H), 2.23–2.16 (m, 1H), 2.03–1.96 (m, 1H), 1.88 (t, *J* = 11.0 Hz, 1H), 1.80–1.71 (m, 4H), 1.49–1.42 (m, 2H), 1.37–1.30 (m, 4H), 0.94–0.84 (m, 4H) ppm. ^13^C NMR (126 MHz, CDCl_3_): *δ* 166.4, 163.2, 131.7, 122.4, 114.2, 68.4, 67.2, 59.1, 56.0, 46.4, 35.9, 31.7, 29.2, 26.7, 25.8, 24.6, 22.7, 14.1 ppm. HRMS [M+H]^+^: 334.23740 (calc. for [C_20_H_32_NO_3_]^+^: 334.23767), 98% purity. **6o** was isolated as an oil in 45% yield (228 mg).

*(1-Methylpiperidin-3-yl)methyl 4-(hexyloxy)benzoate (**6p**)*: Prepared from carboxylic acid **4** and 1-methyl-3-piperidinemethanol (**8p**). The column chromatography used a mobile phase DCM/MeOH/NH_3_ (20/1/0.1). ^1^H NMR (500 MHz, CDCl_3_): *δ* 7.99–7.95 (m, 2H), 6.90 (d, *J* = 8.5 Hz, 2H), 4.21–4.17 (m, 1H), 4.13–4.09 (m, 1H), 4.00 (t, *J* = 6.6 Hz, 2H), 3.03–2.99 (m, 1H), 2.92–2.86 (m, 1H), 2.34–2.30 (m, 3H), 2.22–2.15 (m, 1H), 2.03–1.95 (m, 1H), 1.90–1.7 (m, 5H), 1.51–1.42 (m, 2H), 1.39–1.29 (m, 4H), 0.94–0.83 (m, 4H) ppm. ^13^C NMR (126 MHz, CDCl_3_): *δ* 166.4, 163.2, 131.7, 122.4, 114.2, 68.4, 67.2, 59.1, 56.1, 46.5, 31.7, 29.2, 26.7, 25.8, 24.6, 22.7, 14.1 ppm. HRMS [M+H]^+^: 334.23715 (calc. for [C_20_H_32_NO_3_]^+^: 334.23767), 95% purity. **6p** was isolated as an oil in 47% yield (124 mg).

*[(3R)-1-Methylpiperidin-3-yl]methyl 4-(hexyloxy)benzoate (**6r**):* Prepared from carboxylic acid **4** and (*R*)-1-methyl-3-(hydroxymethyl)piperidine (**8r**). The column chromatography used a mobile phase DCM/MeOH/NH_3_ (20/1/0.1). ^1^H NMR (500 MHz, CDCl_3_): *δ* 7.97–7.94 (m, 2H), 6.91–6.88 (m, 2H), 4.18 (dd, *J* = 11.0, 5.6 Hz, 1H), 4.11 (dd, *J* = 11.0, 7.3 Hz, 1H), 3.99 (t, *J* = 6.6 Hz, 2H), 3.00–2.97 (m, 1H), 2.84 (dd, *J* = 9.4, 5.7 Hz, 1H), 2.31 (s, 3H), 2.19–2.13 (m, 1H), 1.97 (td, *J* = 11.2, 3.2 Hz, 1H), 1.82–1.67 (m, 5H), 1.48–1.42 (m, 2H), 1.36–1.31 (m, 4H), 0.92–0.88 (m, 4H) ppm. ^13^C NMR (126 MHz, CDCl_3_): *δ* 166.4, 163.1, 131.7, 122.5, 114.2, 68.3, 67.3, 59.2, 56.1, 46.6, 36.0, 31.7, 29.2, 26.7, 25.8, 24.7, 22.7, 14.1 ppm. HRMS [M+H]^+^: 334.23730 (calc. for [C_20_H_32_NO_3_]^+^: 334.23767), 99% purity. **6r** was isolated as an oil in 25% yield (83 mg).

*(1-Methylpiperidin-4-yl)methyl 4-(hexyloxy)benzoate (**6s**)*: Prepared from carboxylic acid 4 and 4-hydroxy-1-methylpiperidine (8s). The column chromatography used a mobile phase PE/EA (4/1). ^1^H NMR (500 MHz, CDCl_3_): *δ* 7.99–7.95 (m, 2H), 6.90–6.87 (m, 2H), 5.03–4.99 (m, 1H), 3.98 (t, *J* = 6.6 Hz, 2H), 2.68 (s, 2H), 2.40–2.28 (m, 5H), 2.03–1.98 (m, 2H), 1.88–1.81 (m, 2H), 1.77 (dq, *J* = 7.9, 6.6 Hz, 2H), 1.47–1.41 (m, 2H), 1.35–1.29 (m, 4H), 0.91–0.84 (m, 3H) ppm. ^13^C NMR (126 MHz, CDCl_3_): *δ* 165.8, 163.0, 131.6, 122.8, 114.1, 68.3, 63.7, 60.5, 52.9, 46.2, 31.6, 30.9, 29.2, 25.8, 22.7, 14.1 ppm. HRMS [M+H]^+^: 320.22240 (calc. for [C_19_H_30_NO_3_]^+^: 320.22202), 97% purity. **6s** was isolated as an amorphous solid in 42% yield (189 mg).

*(1-Methylpyrrolidin-2-yl)methyl 4-(hexyloxy)benzoate (**6t**)*: Prepared from carboxylic acid **4** and 1-methyl-2-pyrrolidinementhanol (**8t**). Column chromatography used a mobile phase DCM/MeOH/NH_3_ (20/1/0.1). ^1^H NMR (500 MHz, CDCl_3_): *δ* 7.99–7.96 (m, 2H), 6.91–6.88 (m, 2H), 4.34–4.31 (m, 2H), 4.00 (t, *J* = 6.6 Hz, 2H), 3.18–3.14 (m, 1H), 2.72–2.67 (m, 1H), 2.51 (s, 3H), 2.38–2.32 (m, 1H), 2.06–2.00 (m, 1H), 1.90–1.70 (m, 3H), 1.50–1.42 (m, 2H), 1.38–1.26 (m, 4H), 0.94–0.86 (m, 3H) ppm. ^13^C NMR (126 MHz, CDCl_3_): *δ* 166.4, 163.2, 131.8, 122.4, 114.2, 68.4, 67.0, 64.4, 57.8, 41.6, 31.7, 29.2, 28.6, 25.8, 23.0, 22.7, 14.2 ppm. HRMS [M+H]^+^: 320.22218 (calc. for [C_19_H_30_NO_3_]^+^: 320.22202), 96% purity. **6t** was isolated as an oil in 39% yield (166 mg).

*[(2S)-1-Methylpyrrolidin-2-yl]methyl 4-(hexyloxy)benzoate (**6w**):* Prepared from carboxylic acid **4** and (*S*)-1-methyl-2-pyrrolidinemethanol (**8w**). The column chromatography used a mobile phase DCM/MeOH/NH_3_ (20/1/0.1). ^1^H NMR (500 MHz, CDCl_3_): *δ* 7.99–7.96 (m, 2H), 6.91–6.88 (m, 2H), 4.33–4.32 (m, 2H), 4.00 (t, *J* = 6.6 Hz, 2H), 3.18–3.15 (m, 1H), 2.70 (s, 1H), 2.52 (s, 3H), 2.38–2.33 (m, 1H), 2.08–2.00 (m, 1H), 1.95–1.85 (m, 1H), 1.82–1.70 (m, 3H), 1.48–1.42 (m, 2H), 1.37–1.30 (m, 4H), 0.93–0.87 (m, 3H) ppm. ^13^C NMR (126 MHz, CDCl_3_): *δ* 166.4, 163.2, 131.8, 122.4, 114.2, 68.4, 67.0, 64.4, 57.8, 41.7, 31.7, 29.2, 28.6, 25.8, 22.7, 14.2 ppm. HRMS [M+H]^+^: 320.22215 (calc. for [C_19_H_30_NO_3_]^+^: 320.22202), 98% purity. **6w** was isolated as an oil in 43% yield (195 mg).

*[(2R)-1-Methylpyrrolidin-2-yl]methyl 4-(hexyloxy)benzoate (**6x**)*: Prepared from carboxylic acid **4** and *N*-methyl-D-prolinol (**8x**). The column chromatography used a mobile phase DCM/MeOH/NH_3_ (20/1/0.1). ^1^H NMR (500 MHz, CDCl_3_): *δ* 7.99–7.96 (m, 2H), 6.91–6.88 (m, 2H), 4.27 (t, *J* = 5.5 Hz, 2H), 3.93 (t, *J* = 6.6 Hz, 2H), 3.19–3.14 (m, 1H), 2.73–2.67 (m, 1H), 2.52–2.51 (m, 3H), 2.39–2.34 (m, 1H), 2.08–2.00 (m, 1H), 1.96–1.85 (m, 1H), 1.82–1.66 (m, 3H), 1.48–1.42 (m, 2H), 1.37–1.31 (m, 4H), 0.92–0.88 (m, 3H) ppm. ^13^C NMR (126 MHz, CDCl_3_): *δ* 166.4, 163.2, 131.8, 122.4, 114.2, 68.4, 66.9, 64.4, 57.8, 41.6, 31.7, 29.2, 28.6, 25.8, 22.9, 22.7, 14.1 ppm. HRMS [M+H]^+^: 320.22217 (calc. for [C_19_H_30_NO_3_]^+^: 320.22202), 98% purity. **6x** was isolated as an oil in 47% yield (210 mg).

*(1-Methylpyrrolidin-3-yl)methyl 4-(hexyloxy)benzoate (**6y**)*: Prepared from carboxylic acid **4** and (1-methylpyrrolidin-3-yl)methanol (**8y**). The column chromatography used a mobile phase DCM/MeOH/NH_3_ (20/1/0.1). ^1^H NMR (500 MHz, CDCl_3_): *δ* 7.97–7.94 (m, 2H), 6.91–6.88 (m, 2H), 4.28–4.24 (m, 1H), 4.21–4.17 (m, 1H), 3.99 (t, *J* = 6.6 Hz, 2H), 2.88–2.83 (m, 1H), 2.74–2.68 (m, 2H), 2.63–2.58 (m, 1H), 2.47–2.38 (m, 4H), 2.12–2.06 (m, 1H), 1.81–1.76 (m, 2H), 1.67–1.62 (m, 1H), 1.48–1.42 (m, 2H), 1.37–1.29 (m, 4H), 0.93–0.85 (m, 3H) ppm. ^13^C NMR (126 MHz, CDCl_3_): *δ* 166.5, 163.2, 131.7, 122.4, 114.2, 68.4, 67.3, 59.3, 55.9, 42.1, 37.3, 31.7, 29.2, 28.0, 25.8, 22.7, 14.1 ppm. HRMS [M+H]^+^: 320.22246 (calc. for [C_19_H_30_NO_3_]^+^: 320.22202), 98% purity. **6y** was isolated as an oil in 31% yield (111 mg).

*[(3S)-1-Methylpyrrolidin-3-yl]methyl 4-(hexyloxy)benzoate (**6z**)*: Prepared from carboxylic acid **4** and (*S*)-(1-methylpyrrolidin-3-yl)methanol (**8z**). The column chromatography used a mobile phase DCM/MeOH/NH_3_ (20/1/0.1). ^1^H NMR (500 MHz, CDCl_3_): *δ* 7.97–7.94 (m, 2H), 6.91–6.88 (m, 2H), 4.29–4.25 (m, 1H), 4.22–4.18 (m, 1H), 4.00 (t, *J* = 6.6 Hz, 2H), 2.90–2.85 (m, 1H), 2.75–2.69 (m, 2H), 2.65–2.59 (m, 1H), 2.49–2.38 (m, 4H), 2.14–2.07 (m, 1H), 1.82–1.76 (m, 2H), 1.70–1.63 (m, 1H), 1.48–1.42 (m, 2H), 1.37–1.30 (m, 4H), 0.90–0.87 (m, 3H) ppm. ^13^C NMR (126 MHz, CDCl_3_): *δ* 166.4, 163.2, 131.7, 122.4, 114.2, 68.4, 67.2, 59.3, 55.9, 42.1, 37.3, 31.7, 29.2, 28.0, 25.8, 22.7, 14.2 ppm. HRMS [M+H]^+^: 320.22235 (calc. for [C_19_H_30_NO_3_]^+^: 320.22202), 99% purity. **6z** was isolated as an oil in 28% yield (102 mg).

*(2R)-1-{2-[4-(Hexyloxy)benzoyloxy]ethyl}-1,2-dimethylpiperidin-1-ium iodide (**7e**)*: Prepared from **6e** by general procedure **C**. ^1^H NMR (500 MHz, CDCl_3_): *δ* 7.98–7.90 (m, 2H), 6.93–6.88 (m, 2H), 4.86 (t, *J* = 4.9 Hz, 1H), 4.29–4.25 (m, 1H), 4.01–3.97 (m, 2H), 3.23 (s, 2H), 3.16 (q, *J* = 7.4 Hz, 4H), 2.02–1.91 (m, 2H), 1.90–1.82 (m, 1H), 1.82–1.74 (m, 2H), 1.49–1.41 (m, 8H), 1.39 (s, 1H), 1.35–1.30 (m, 4H), 1.27 (s, 1H), 0.91–0.87 (m, 3H) ppm. ^13^C NMR (126 MHz, CDCl_3_): *δ* 164.6, 162.8, 131.0, 119.5, 113.5, 67.4, 66.2, 62.1, 60.7, 56.5, 45.2, 30.6, 30.5, 28.7, 28.0, 27.4, 24.6, 21.6, 19.4, 15.1, 13.0, 7.7 ppm. HRMS [M+H]^+^: 362.26897 (calc. for [C_22_H_36_NO_3_]^+^: 362.26944), 98% purity. **7e** was isolated as a yellow oil in 37% yield (26 mg).

*(2S)-1-{2-[4-(Hexyloxy)benzoyloxy]ethyl}-1,2-dimethylpiperidin-1-ium iodide (**7f**)*: Prepared from **6f** by general procedure **C**. ^1^H NMR (500 MHz, CDCl_3_): *δ* 7.97–7.90 (m, 2H), 6.93–6.89 (m, 2H), 4.86 (t, *J* = 5.0 Hz, 1H), 4.84–4.81 (m, 1H), 4.34–4.29 (m, 1H), 4.07–3.93 (m, 4H), 3.57 (s, 1H), 3.24 (s, 2H), 2.01–1.95 (m, 2H), 1.94–1.90 (m, 2H), 1.89–1.83 (m, 2H), 1.83–1.76 (m, 3H), 1.74 (s, 1H), 1.50 (d, *J* = 6.6 Hz, 1H), 1.48–1.42 (m, 4H), 1.36–1.30 (m, 4H), 0.92–0.88 (m, 3H) ppm. ^13^C NMR (126 MHz, CDCl_3_): *δ* 165.6, 165.6, 163.9, 163.8, 132.0, 131.9, 120.5, 120.4, 114.6, 114.5, 68.4, 66.9, 63.0, 61.6, 57.5, 57.3, 31.5, 29.0, 25.6, 22.6, 16.0, 14.0 ppm. HRMS [M+H]^+^: 362.26897 (calc. for [C_22_H_36_NO_3_]^+^: 362.26944), 97% purity. **7f** was isolated as a white solid (melting point 152– 154 °C) in 67% yield (47 mg).

*2-{2-[4-(Hexyloxy)benzoyloxy]ethyl}-2-methyl-2-azabicyclo[2.2.2]octan-2-ium iodide (**7g**)*: Prepared from **6g** by general procedure **C**. ^1^H NMR (500 MHz, CDCl_3_): *δ* 7.97–7.92 (m, 2H), 6.94–6.89 (m, 2H), 4.88–4.77 (m, 2H), 4.41–4.34 (m, 1H), 4.25–4.18 (m, 1H), 4.00 (t, *J* = 6.5 Hz, 2H), 3.92–3.88 (m, 1H), 3.83–3.78 (m, 1H), 3.73–3.68 (m, 1H), 3.56 (s, 3H), 2.48–2.38 (m, 2H), 2.19–2.13 (m, 1H), 1.98–1.83 (m, 4H), 1.82–1.75 (m, 3H), 1.49–1.40 (m, 2H), 1.37–1.30 (m, 4H), 1.25 (s, 1H), 0.93–0.88 (m, 3H) ppm. ^13^C NMR (126 MHz, CDCl_3_): *δ* 165.6, 163.8, 132.0, 120.5, 114.6, 69.2, 68.4, 64.1, 61.6, 58.2, 52.1, 31.5, 29.0, 25.6, 24.7, 22.6, 21.8, 21.8, 21.6, 21.4, 14.0 ppm. HRMS [M+H]^+^: 374.26868 (calc. for [C_23_H_36_NO_3_]^+^: 374.26897), 99% purity. **7g** was isolated as a yellow solid (melting point 177–178 °C) in 71% yield (35 mg).

*1-{2-[4-(Hexyloxy)benzoyloxy]ethyl}-1-methyl-2,5-dihydro-1H-pyrrol-1-ium iodide (**7h**)*: Prepared from **6h** by general procedure **C**. ^1^H NMR (500 MHz, CDCl_3_): *δ* 7.99–7.93 (m, 2H), 6.93–6.89 (m, 2H), 6.02 (s, 2H), 4.86 (s, 1H), 4.83–4.79 (m, 2H), 4.62 (d, *J* = 2.6 Hz, 1H), 4.59 (d, *J* = 3.1 Hz, 1H), 4.54–4.50 (m, 2H), 3.98 (t, *J* = 6.5 Hz, 2H), 3.62 (s, 3H), 1.81–1.73 (m, 2H), 1.48–1.40 (m, 2H), 1.36–1.30 (m, 4H), 1.24 (s, 1H), 0.91–0.87 (m, 3H) ppm. ^13^C NMR (126 MHz, CDCl_3_): *δ* 165.6, 163.8, 132.0, 124.7, 120.5, 114.6, 72.3, 68.4, 63.8, 58.9, 52.7, 31.9, 31.5, 29.7, 29.0, 25.6, 22.7, 22.6, 14.1, 14.0 ppm. HRMS [M+H]^+^: 332.22189 (calc. for [C_20_H_30_NO_3_]^+^: 332.22202), 96% purity. **7h** was isolated as a purple solid (melting point 88–90 °C) in 51% yield (56 mg).

*{2-[4-(Hexyloxy)benzoyloxy]ethyl}-1-methylpyrrolidin-1-ium iodide (**7j**)*: Prepared from **6j** by general procedure **C**. ^1^H NMR (500 MHz, CDCl_3_): *δ* 7.92–7.88 (m, 2H), 6.88–6.84 (m, 2H), 4.78–4.72 (m, 2H), 4.24–4.20 (m, 2H), 3.96–3.89 (m, 3H), 3.35 (s, 3H), 2.38–2.21 (m, 4H), 1.76–1.68 (m, 2H), 1.43–1.35 (m, 2H), 1.30–1.24 (m, 4H), 1.19 (s, 3H), 0.86–0.82 (m, 3H) ppm. ^13^C NMR (126 MHz, CDCl_3_): *δ* 165.5, 163.9, 132.0, 120.4, 114.6, 68.5, 66.0, 63.0, 58.5, 49.5, 31.9, 31.5, 29.7, 29.4, 29.0, 25.6, 22.7, 22.6, 21.7, 14.0 ppm. HRMS [M+H]^+^: 334.23450 (calc. for [C_20_H_32_NO_3_]^+^: 334.23767), 98% purity. **7j** was isolated as a yellow solid (melting point 88– 90 °C) in 63% yield (35 mg).

*{2-[4-(Hexyloxy)benzoyloxy]ethyl}-2-methyl-2,3-dihydro-1H-isoindol-2-ium iodide (**7k**)*: Prepared from **6k** by general procedure **C**. ^1^H NMR (500 MHz, CDCl_3_): *δ* 7.99–7.95 (m, 2H), 7.40–7.35 (m, 4H), 6.93–6.90 (m, 2H),

5.33 (d, *J* = 14.2 Hz, 2H), 5.13 (d, *J* = 14.2 Hz, 2H), 4.83–4.81 (m, 2H), 4.68–4.66 (m, 2H), 3.99 (t, *J* = 6.6 Hz, 2H), 3.64 (s, 3H), 1.78 (dq, *J* = 8.3, 6.6 Hz, 2H), 1.45 (dtd, *J* = 9.4, 7.2, 5.2 Hz, 2H), 1.33 (dq, *J* = 7.2, 3.4 Hz, 4H), 0.92–0.89 (m, 3H) ppm. ^13^C NMR (126 MHz, CDCl_3_): *δ* 165.6, 163.9, 132.2, 132.0, 129.9, 123.8, 120.6, 114.7, 70.8, 68.5, 63.5, 58.9, 51.5, 31.7, 29.1, 25.8, 22.7, 14.1 ppm. HRMS [M+H]^+^: 382.23737 (calc. for [C_24_H_32_NO_3_]^+^: 382.23767), 96% purity. **7k** was isolated as an amorphous solid in 25% yield (31 mg).

*2-{[4-(Hexyloxy)benzoyloxy]methyl}-1,1-dimethylpiperidin-1-ium iodide (**7m**)*: Prepared from **6m** by general procedure **C**. HRMS [M+H]^+^: 348.25341 (calc. for [C_21_H_34_NO_3_]^+^: 348.25332), 97% purity. **7m** was isolated as a slightly yellow solid (decomposition from 158 °C) in 17% yield (25 mg).

*(2R)-2-{[4-(Hexyloxy)benzoyloxy]methyl}-1,1-dimethylpiperidin-1-ium iodide (**7n**)*: Prepared from **6n** by general procedure **C**. HRMS [M+H]^+^: 348.25347 (calc. for [C_21_H_34_NO_3_]^+^: 348.25332), 99% purity. **7m** was isolated as a white solid (melting point 219–221 °C) in 15% yield (18 mg).

*(3S)-3-{[4-(hexyloxy)benzoyloxy]methyl}-1,1-dimethylpiperidin-1-ium iodide (**7o**)*: Prepared from **6o** by general procedure **C**. ^1^H NMR (500 MHz, CD_3_OD): *δ* 8.03–8.00 (m, 2H), 7.00–6.97 (m, 2H), 4.30 (dd, *J* = 11.3, 4.8 Hz, 1H), 4.20 (dd, *J* = 11.3, 6.6 Hz, 1H), 4.05 (t, *J* = 6.4 Hz, 2H), 3.67 (ddt, *J* = 12.5, 3.9, 2.0 Hz, 1H), 3.54 (ddq, *J* = 12.4, 3.8, 1.8 Hz, 1H), 3.38 (td, *J* = 13.1, 3.5 Hz, 1H), 3.27–3.25 (m, 4H), 3.20 (s, 3H), 2.62–2.57 (m, 1H), 2.13–2.07 (m, 1H), 2.02–1.95 (m, 2H), 1.82–1.77 (m, 2H), 1.52–1.44 (m, 3H), 1.41–1.32 (m, 4H), 0.95–0.91 (m, 3H) ppm. ^13^C NMR (126 MHz, CD_3_OD): *δ* 167.5, 164.9, 132.9, 122.8, 115.4, 69.4, 66.6, 65.6, 63.6, 57.5, 32.7, 32.7, 30.2, 26.8, 25.3, 23.7, 20.8, 14.4 ppm. HRMS [M+H]^+^: 348.25374 (calc. for [C_21_H_34_NO_3_]^+^: 348.25332), 100% purity. **7k** was isolated as a white solid (melting point 172–174 °C) in 21% yield (34 mg).

*3-{[4-(hexyloxy)benzoyloxy]methyl}-1,1-dimethylpiperidin-1-ium iodide (**7p**)*: Prepared from **6p** by general procedure **C**. ^1^H NMR (500 MHz, CD_3_OD): *δ* 8.03–7.99 (m, 2H), 7.00–6.97 (m, 2H), 4.30 (dd, *J* = 11.3, 4.8 Hz, 1H), 4.20 (dd, *J* = 11.3, 6.7 Hz, 1H), 4.05 (t, *J* = 6.4 Hz, 2H), 3.67 (ddt, *J* = 12.5, 3.9, 2.0 Hz, 1H), 3.56–3.52 (m, 1H), 3.39 (td, *J* = 13.1, 3.4 Hz, 1H), 3.28–3.24 (m, 4H), 3.21 (s, 3H), 2.63–2.57 (m, 1H), 2.13–2.06 (m, 1H), 2.02–1.95 (m, 2H), 1.82–1.76 (m, 2H), 1.52–1.47 (m, 3H), 1.40–1.33 (m, 4H), 0.95–0.91 (m, 3H) ppm. ^13^C NMR (126 MHz, CD_3_OD): *δ* 167.5, 164.9, 132.9, 122.8, 115.4, 69.4, 66.6, 65.6, 63.6, 57.5, 32.7, 32.6, 30.2, 26.8, 25.3, 23.7, 20.8, 14.4 ppm. HRMS [M+H]^+^: 348.25345 (calc. for [C_21_H_34_NO_3_]^+^: 348.25332), 100% purity. **7p** was isolated as a slightly yellow solid (melting point 141–143 °C) in 28% yield (41 mg).

*(3R)-3-{[4-(hexyloxy)benzoyloxy]methyl}-1,1-dimethylpiperidin-1-ium iodide (**7r**)*: Prepared from **6r** by general procedure **C**. ^1^H NMR (500 MHz, CD_3_OD): *δ* 8.03–8.00 (m, 2H), 7.00–6.97 (m, 2H), 4.29 (dd, *J* = 11.3, 4.8 Hz, 1H), 4.20 (dd, *J* = 11.3, 6.6 Hz, 1H), 4.06–4.03 (m, 2H), 3.68 (ddt, *J* = 12.4, 4.1, 2.1 Hz, 1H), 3.54 (ddt, *J* = 12.5, 4.1, 2.1 Hz, 1H), 3.40 (td, *J* = 13.0, 3.6 Hz, 1H), 3.29–3.27 (m, 4H), 3.21 (s, 3H), 2.64–2.57 (m, 1H), 2.16–2.06 (m, 1H), 2.01–1.95 (m, 2H), 1.82–1.76 (m, 2H), 1.52–1.42 (m, 3H), 1.40–1.33 (m, 4H), 0.94–0.91 (m, 3H) ppm. ^13^C NMR (126 MHz, CD_3_OD): *δ* 167.5, 164.9, 132.9, 122.8, 115.4, 69.4, 66.6, 65.6, 63.6, 57.5, 32.7, 32.6, 30.2, 26.8, 25.3, 23.7, 20.8, 14.4 ppm. HRMS [M+H]^+^: 348.25345 (calc. for [C_21_H_34_NO_3_]^+^: 348.25332), 100% purity. **7r** was isolated as a white solid (melting point 179–181 °C) in 32% yield (40 mg).

*4-[4-(hexyloxy)benzoyloxy]-1,1-dimethylpiperidin-1-ium iodide (**7s**)*: Prepared from **6s** by general procedure **C**. ^1^H NMR (500 MHz, CD_3_OD): *δ* 8.06–8.02 (m, 2H), 7.01–6.98 (m, 2H), 5.25 (tt, *J* = 6.1, 3.5 Hz, 1H), 4.05 (t, *J* = 6.4 Hz, 2H), 3.66 (ddd, *J* = 13.4, 9.9, 3.7 Hz, 2H), 3.56 (dt, *J* = 13.5, 5.2 Hz, 2H), 3.31–3.30 (m, 3H), 3.25 (s, 3H), 2.42–2.36 (m, 2H), 2.23–2.17 (m, 2H), 1.79 (dq, *J* = 8.5, 6.5 Hz, 2H), 1.52–1.45 (m, 2H), 1.40–1.33 (m, 4H), 0.95–0.90 (m, 3H) ppm. ^13^C NMR (126 MHz, CD_3_OD): *δ* 166.7, 165.0, 132.9, 122.9, 115.4, 69.4, 66.0, 60.3, 32.7, 30.2, 26.8, 26.2, 23.7, 14.4 ppm. HRMS [M+H]^+^: 334.23764 (calc. for [C_20_H_32_NO_3_]^+^: 334.23767), 100% purity. **7s** was isolated as a white solid (melting point 168–170 °C) in 25% yield (35 mg).

*2-{[4-(hexyloxy)benzoyloxy]methyl}-1,1-dimethylpyrrolidin-1-ium iodide (**7t**)*: Prepared from **6t** by general procedure **C**. ^1^H NMR (500 MHz, CD_3_OD): *δ* 8.02–7.99 (m, 2H), 7.03–7.00 (m, 2H), 4.75–4.65 (m, 2H), 4.19 (qd, *J* = 7.8, 4.2 Hz, 1H), 4.06 (t, *J* = 6.5 Hz, 2H), 3.81–3.69 (m, 2H), 3.36 (s, 3H), 3.16 (s, 3H), 2.48 (dq, *J* = 15.6, 7.8 Hz, 1H), 2.28–2.22 (m, 2H), 2.19–2.11 (m, 1H), 1.82–1.7 (m, 2H), 1.52–1.45 (m, 2H), 1.40–1.33 (m, 4H), 0.96–0.87 (m, 3H) ppm. ^13^C NMR (126 MHz, CD_3_OD): *δ* 166.7, 165.2, 132.9, 122.2, 115.6, 75.2, 69.5, 69.1, 61.2, 53.5, 46.5, 32.7, 30.2, 26.8, 25.5, 23.7, 20.5, 14.4 ppm. HRMS [M+H]^+^: 334.23749 (calc. for [C_20_H_32_NO_3_]^+^: 334.23767), 100% purity. **7t** was isolated as an oil in 27% yield (36 mg).

*(2S)-2-{[4-(hexyloxy)benzoyloxy]methyl}-1,1-dimethylpyrrolidin-1-ium iodide (**7w**)*: Prepared from **6w** by general procedure **C**. ^1^H NMR (500 MHz, CD_3_OD): *δ* 8.02–7.99 (m, 2H), 7.02–7.00 (m, 2H), 4.75–4.66 (m, 2H), 4.24–4.17 (m, 1H), 4.06 (t, *J* = 6.4 Hz, 2H), 3.82–3.70 (m, 2H), 3.37 (s, 3H), 3.17 (s, 3H), 2.51–2.44 (m, 1H), 2.29–2.21 (m, 2H), 2.17–2.11 (m, 1H), 1.82–1.76 (m, 2H), 1.52–1.45 (m, 2H), 1.40–1.33 (m, 4H), 0.94–0.91 (m, 3H) ppm. ^13^C NMR (126 MHz, CD_3_OD): *δ* 166.7, 165.1, 132.9, 122.1, 115.6, 75.2, 69.5, 69.1, 61.2, 53.5, 46.6, 32.7, 30.2, 26.8, 25.5, 23.6, 20.5, 14.4 ppm. HRMS [M+H]^+^: 334.23761 (calc. for [C_20_H_32_NO_3_]^+^: 334.23767), 100% purity. **7w** was isolated as a slightly yellow solid (melting point 83–85 °C) in 17% yield (25 mg).

*(2R)-2-{[4-(hexyloxy)benzoyloxy]methyl}-1,1-dimethylpyrrolidin-1-ium iodide (**7x**)*: Prepared from **6x** by general procedure **C**. ^1^H NMR (500 MHz, CD_3_OD): *δ* 8.01–7.99 (m, 2H), 7.03–7.01 (m, 2H), 4.75–4.65 (m, 2H), 4.23–4.15 (m, 1H), 4.06 (t, *J* = 6.4 Hz, 2H), 3.81–3.69 (m, 2H), 3.36 (s, 3H), 3.16 (s, 3H), 2.48 (dq, *J* = 15.2, 7.9 Hz, 1H), 2.28–2.18 (m, 2H), 2.17–2.10 (m, 1H), 1.84–1.77 (m, 2H), 1.52–1.45 (m, 2H), 1.39–1.33 (m, 4H), 0.95–0.90 (m, 3H) ppm. ^13^C NMR (126 MHz, CD_3_OD): *δ* 166.7, 165.2, 132.9, 122.2, 115.6, 75.2, 69.5, 69.1, 61.2, 53.5, 46.6, 32.7, 30.2, 26.8, 25.5, 23.7, 20.5, 14.4 ppm. HRMS [M+H]^+^: 334.23767 (calc. for [C_20_H_32_NO_3_]^+^: 334.23767), 99% purity. **7x** was isolated as a slightly yellow solid (decomposition from 85 °C) in 26% yield (33 mg).

*3-{[4-(hexyloxy)benzoyloxy]methyl}-1,1-dimethylpyrrolidin-1-ium iodide (**7y**)*: Prepared from **6y** by general procedure **C**. ^1^H NMR (500 MHz, CD_3_OD): *δ* 8.03–8.00 (m, 2H), 7.03–7.01 (m, 2H), 4.47 (dd, *J* = 11.1, 5.8 Hz, 1H), 4.40 (dd, *J* = 11.1, 7.3 Hz, 1H), 4.08 (t, *J* = 6.4 Hz, 2H), 3.95 (qd, *J* = 7.3, 3.1 Hz, 1H), 3.78–3.69 (m, 2H), 3.53 (ddt, *J* = 11.7, 5.7, 3.0 Hz, 1H), 3.34 (s, 3H), 3.27–3.23 (m, 4H), 2.58–2.48 (m, 1H), 2.23–2.17 (m, 1H), 1.89–1.80 (m, 2H), 1.55–1.48 (m, 2H), 1.43–1.33 (m, 4H), 0.97–0.95 (m, 3H) ppm. ^13^C NMR (126 MHz, CD_3_OD): *δ* 167.5, 164.9, 132.8, 122.8, 115.4, 69.4, 69.3, 66.8, 66.0, 54.0, 52.9, 37.3, 32.7, 30.2, 26.8, 26.7, 23.7, 14.4 ppm. HRMS [M+H]^+^: 334.23758 (calc. for [C_20_H_32_NO_3_]^+^: 334.23767), 99% purity. **7y** was isolated as an amorphous solid in 37% yield (28 mg).

*(3S)-3-{[4-(hexyloxy)benzoyloxy]methyl}-1,1-dimethylpyrrolidin-1-ium iodide (**7z**)*: Prepared from **6z** by general procedure **C**. ^1^H NMR (500 MHz, CD_3_OD): *δ* 8.03–8.00 (m, 2H), 7.04–7.01 (m, 2H), 4.47 (dd, *J* = 11.1, 5.8 Hz, 1H), 4.40 (dd, *J* = 11.1, 7.3 Hz, 1H), 4.08 (t, *J* = 6.4 Hz, 2H), 3.94 (dd, *J* = 12.1, 8.3 Hz, 1H), 3.78–3.68 (m, 2H), 3.53 (dd, *J* = 12.1, 9.6 Hz, 1H), 3.34 (s, 3H), 3.26 (s, 4H), 2.57–2.51 (m, 1H), 2.22–2.17 (m, 1H), 1.85–1.80 (m, 2H), 1.55–1.48 (m, 2H), 1.43–1.37 (m, 4H), 0.98–0.94 (m, 3H) ppm. ^13^C NMR (126 MHz, CD_3_OD): *δ* 167.5, 164.9, 132.8, 122.8, 115.4, 69.4, 69.3, 66.8, 66.0, 53.9, 52.9, 37.3, 32.7, 30.2, 26.8, 26.7, 23.7, 14.4 ppm. HRMS [M+H]^+^: 334.23767 (calc. for [C_20_H_32_NO_3_]^+^: 334.23767), 99% purity. **7z** was isolated as a white solid (decomposition from 122 °C) in 42% yield (35 mg).

*Quinuclidin-3-yl 4-hexyloxybenzoate:* Prepared from 3-quinuclidinol and *p*-hexyloxybenzoyl chloride by general procedure **D** in 59% yield (1.5 g). The crude product was used without purification in the following reactions.

*Quinuclidin-3-yl 4-hexyloxybenzoate hydrochloride (**12a**):* Prepared from quinuclidin-3-yl 4-hexyloxybenzoate by general procedure **E**. ^1^H NMR (500 MHz, CD_3_OD): *δ* 8.03–7.99 (m, 2H), 7.02–6.99 (m, 2H), 5.30 (dddd, *J* = 8.5, 4.3, 2.8, 1.3 Hz, 1H), 4.06 (t, *J* = 6.5 Hz, 2H), 3.84 (ddd, *J* = 14.1, 8.6, 2.1 Hz, 1H), 3.48–3.32 (m, 5H), 2.51 (dtd, *J* = 6.6, 4.0, 2.5 Hz, 1H), 2.34 (dddd, *J* = 15.2, 9.1, 4.8, 2.6 Hz, 1H), 2.17–2.10 (m, 1H), 2.07–1.94 (m, 2H), 1.80 (ddt, *J* = 9.2, 7.9, 6.4 Hz, 2H), 1.52–1.46 (m, 2H), 1.41–1.29 (m, 4H), 0.96–0.89 (m, 3H) ppm. ^13^C NMR (126 MHz, CD_3_OD): *δ* 167.0, 165.1, 132.9, 122.5, 115.4, 69.4, 68.5, 54.6, 47.8, 47.0, 32.7, 30.2, 26.8, 25.3, 23.7, 21.2, 18.1, 14.3 ppm. HRMS [M+H]^+^: 332.22125 (calc. for [C_20_H_20_NO_3_]^+^: 322.22202), 97% purity. **12a** was isolated as a white solid (melting point 187–188 °C) in 40% yield (401 mg).

*1-methylquinuclidin-3-yl 4-hexyloxybenzoate iodide (**12b**):* Prepared from quinuclidin-3-yl 4-hexyloxybenzoate by general procedure **C**. ^1^H NMR (CD_3_OD): *δ* 7.93–7.90 (m, 2H), 6.92–6.89 (m, 2H), 5.36–5.31 (m, 1H), 4.07–4.05 (m, 2H), 4.01–3.96 (m, 1H), 3.63–3.50 (m, 3H), 3.47–3.40 (m, 1H), 3.05 (s, 3H), 2.58–2.54 (m, 1H), 2.45–2.38 (m, 1H), 2.23–2.16 (m, 1H), 2.14–2.03 (m, 2H), 1.83–1.77 (m, 2H), 1.52–1.46 (m, 2H), 1.41–1.34 (m, 5H), 0.95–0.82 (m, 3H) ppm. ^13^C NMR (126 MHz, CD_3_OD): *δ* 166.9, 165.2, 132.9, 122.4, 115.5, 69.4, 69.0, 64.1, 58.0, 57.2, 52.2, 32.7, 30.2, 26.8, 25.2, 23.7, 22.3, 19.6, 14.3 ppm. HRMS [M+H]^+^: 346.23672 (calc. for [C_21_H_32_NO_3_]^+^: 346.23767), 92% purity. **12b** was isolated as a white solid (melting point 143–145 °C) in 78% yield (504 mg).

*Quinuclidin-3-yl 4-butoxybenzoate:* Prepared from 3-quinuclidinol and *p*-butoxybenzoyl chloride by general procedure **D** in 32% yield (463 mg). The crude product was used without purification in the following reactions.

*Quinuclidin-3-yl 4-butoxybenzoate hydrochloride (**13a**):* Prepared from quinuclidin-3-yl 4-butoxybenzoate by general procedure **E**. ^1^H NMR (500 MHz, CD_3_OD): *δ* 8.03–7.99 (m, 2H), 7.02–6.99 (m, 2H), 5.30 (dddd, *J* = 8.4, 4.2, 2.9, 1.3 Hz, 1H), 4.07 (t, *J* = 6.4 Hz, 2H), 3.87–3.82 (m, 1H), 3.48–3.32 (m, 5H), 2.51 (dtd, *J* = 6.6, 4.0, 2.5 Hz, 1H), 2.33 (dddd, *J* = 15.6, 13.9, 7.3, 4.1 Hz, 1H), 2.17–2.09 (m, 1H), 2.08–1.97 (m, 2H), 1.81–1.76 (m, 2H),

1.56–1.48 (m, 2H), 0.9 (t, *J* = 7.4 Hz, 3H) ppm. ^13^C NMR (126 MHz, CD_3_OD): *δ* 167.0, 165.1, 132.9, 122.6, 115.4, 69.1, 68.5, 54.6, 47.8, 47.0, 32.3, 25.3, 21.2, 20.2, 18.1, 14.1 ppm. HRMS [M+H]^+^: 304.19031 (calc. for [C_20_H_20_NO_3_]^+^: 304.19072), 99% purity. **13a** was isolated as a white solid (melting point 180–181 °C) in 87% yield (220 mg).

*1-Methylquinuclidin-3-yl 4-butoxybenzoate iodide (**13b**):* Prepared from quinuclidin-3-yl 4-butoxybenzoate by general procedure **C**. ^1^H NMR (500 MHz, CD_3_OD): *δ* 8.04–8.01 (m, 2H), 7.02–6.98 (m, 2H), 5.35–5.32 (m, 1H), 4.07 (t, *J* = 6.4 Hz, 2H), 4.03–3.97 (m, 1H), 3.65–3.53 (m, 4H), 3.49–3.41 (m, 1H), 3.08–3.05 (m, 3H), 2.58–2.54 (m, 1H), 2.43–2.37 (m, 1H), 2.22–2.17 (m, 1H), 2.14–2.04 (m, 2H), 1.79 (dq, *J* = 8.3, 6.5 Hz, 2H), 1.52 (h, *J* = 7.4 Hz, 2H), 1.00 (t, *J* = 7.4 Hz, 3H) ppm. ^13^C NMR (126 MHz, CD_3_OD): *δ* 166.9, 165.2, 132.9, 122.5, 115.4, 69.1, 69.0, 64.1, 58.0, 57.3, 52.3, 32.3, 25.2, 22.3, 20.2, 19.6, 14.1 ppm. HRMS [M+H]^+^: 318.20587 (calc. for [C_19_H_28_NO_3_]^+^: 318.20637), 95% purity. **13b** was isolated as a white solid (melting point 135–136 °C) in 86% yield (350 mg).

*Quinuclidin-3-yl 4-butoxybenzoate:* Prepared from 3-quinuclidinol and *p*-methoxybenzoyl chloride by general procedure **D** in 25% yield (0.770 g). The crude product was used without purification in the following reaction.

*Quinuclidin-3-yl 4-methoxybenzoate hydrochloride (**14a**):* Prepared from quinuclidin-3-yl 4-butoxybenzoate by general procedure **E**. ^1^H NMR (500 MHz, CD_3_OD): *δ* 8.04–8.01 (m, 2H), 7.04–7.01 (m, 2H), 5.32–5.29 (m, 1H), 3.87 (s, 3H), 3.87–3.82 (m, 1H), 3.49–3.41 (m, 3H), 3.40–3.33 (m, 2H), 2.51–2.49 (m, 1H), 2.37–2.30 (m, 1H), 2.18–2.09 (m, 1H), 2.07–1.94 (m, 2H) ppm. ^13^C NMR (126 MHz, CD_3_OD): *δ* 166.9, 165.6, 132.9, 122.8, 115.0, 68.6, 56.1, 54.6, 47.8, 47.0, 25.3, 21.2, 18.1 ppm. HRMS [M+H]^+^: 262.14304 (calc. for [C_20_H_20_NO_3_]^+^: 262.14377), 100% purity. **14a** was isolated as a white solid (melting point 208–210 °C) in 91% yield (310 mg).

*1-methylquinuclidin-3-yl 4-methoxybenzoate iodide (**14b**):* Prepared from quinuclidin-3-yl 4-butoxybenzoate by general procedure **C**. ^1^H NMR (500 MHz, CD_3_OD): *δ* 8.06–8.02 (m, 2H), 7.08–6.95 (m, 2H), 5.35–5.32 (m, 1H), 4.01 (ddd, *J* = 13.8, 8.3, 2.5 Hz, 1H), 3.88 (s, 3H), 3.59–3.54 (m, 4H), 3.46 (dddt, *J* = 15.3, 6.7, 5.0, 2.5 Hz, 1H), 3.05 (s, 3H), 2.56 (qt, *J* = 4.3, 2.5 Hz, 1H), 2.41–2.39 (m, 1H), 2.20 (ddt, *J* = 14.4, 10.0, 4.3 Hz, 1H), 2.14–2.06 (m, 2H) ppm. ^13^C NMR (126 MHz, CD_3_OD): *δ* 166.8, 165.6, 133.0, 122.7, 115.0, 69.1, 64.1, 58.0, 57.3, 56.1, 52.3, 25.2, 22.3, 19.6 ppm. HRMS [M+H]^+^: 276.15860 (calc. for [C_16_H_22_NO_3_]^+^: 276.15942), 96% purity. **14b** was isolated as a white solid (melting point 205–207 °C) in 89% yield (410 mg).

### 5.2. Molecular modelling

#### 5.2.1. Preparation of receptor structure

The structures of the muscarinic acetylcholine receptors were downloaded from the RCSB Protein Data Bank (https://www.rcsb.org/). Non-protein and nanobody molecules were deleted, and the resulting receptor protein was processed in Maestro using the Protein Preparation Wizard according to Sastry et al. guidelines [33].

#### 5.2.2. Ligand docking

Structures of ligands were sketched in ChemAxon MarvinSketch and imported into YASARA, where they were parametrised and energy minimised using the NOVA forcefield [34]. Antagonists KH-5 and compounds **6a-z**, **7a-z**, **13a**, **13b**, and **14a** and **14b** were docked to receptors in inactive conformations (5CXV of M_1_, 3UON of M_2_, 4DAJ of M_3_, 5DSG of M_4_ and 6OL9 of M_5_) and xanomeline was docked to receptors in active conformations (6OIJ for M_1_, 4MQT for M_2_ and 8FX5 of M_4_). The orthosteric binding site was defined as a 5 Å extended cuboid around a co-crystallised orthosteric ligand. The ligands were docked to the orthosteric binding site using the AutoDock local search procedure for 888 poses [35]. All poses were energy minimised and rescored using AutoDock VINA’s local search, confined closely to the original ligand pose. The wash-resistant binding site was defined as a 3 Å extended cuboid around low-scoring poses penetrating the hexyloxy chain among transmembrane helices towards the membrane. The ligands were redocked to the wash-resistant site and rescored. The pose with the highest rescore value was selected for further work.

#### 5.2.3. Ligand descriptors

Eight molecular descriptors of energy-minimised (NOVA forcefield [34]) compounds were determined: 1. The distance between ketoxy oxygen and nitrogen. 2. The improper angle of ketoxy oxygen – ketoxy carbon – ester oxygen – nitrogen. 3. The volume of the compound molecule. 4. The Van der Vals volume of the basic centre. 5. The geometric radius of the basic centre. 6. The solvent-accessible surface area according to the NOVA force field. 7. The electrostatic potential calculated by the Particle Mesh Ewald (PME) method in YASARA. 8. The in-vacuo net charge of the compound molecule according to the NOVA force field.

#### 5.2.4. Simulation of molecular dynamics

To evaluate ligand binding to the receptor and quantify its interactions with the receptor, conventional molecular dynamics (cMD) was simulated using Desmond ver. 7.6 [36]. The simulated system consisted of the receptor-ligand complex in 1-palmitoyl-2-oleoyl-sn-glycero-3-phosphocholine (POPC) membrane set to an OPM-oriented receptor in water and 0.15M NaCl. The system was first relaxed by the standard Desmond protocol for membrane proteins, and then 120 ns γNPT (Noose-Hover chain thermostat at 300 K, Martyna-Tobias-Klein barostat at 1.01325 bar, isotropic coupling, Coulombic cutoff at 0.9 nm) molecular dynamics without restraints was simulated. MD was run three times with random initial velocities (3 independent replicas). The quality of the molecular dynamics simulation was assessed using Maestro’s simulation quality analysis tools and analysed using the simulation event analysis tool. Ligand-receptor interactions were identified using the simulation interaction diagram tool.

### 5.3. Cell culturing

CHO cells stably transfected with the genes of individual subtypes of human muscarinic receptors (M_1_-M_5_) were purchased from the Missouri S&T cDNA resource centre (Rolla, MO, USA) (https://www.cdna.org/home.php?cat=177). Cells were grown to confluence in 75-cm^2^ flasks in Dulbecco’s modified Eagle’s medium DMEM (Thermo Fisher Scientific; LOT: 2556745) supplemented with 10% fetal bovine serum and geneticin at the concentration of 50 µg/ml at 37 °C in a humidified incubator containing 5% –CO2. The medium was supplemented with 5 mM butyrate for the last 24 hours of culture to increase receptor expression. Cells were washed with phosphate-buffered saline (PBS), manually scraped from dishes and centrifuged at 1000 × g for 5 min. Cell pellets were kept at −80°C.

### 5.4. Membrane preparation

Membranes were prepared as described previously by Randakova et al. [11]. Cells harvested from twenty 100 mm Petri dishes (CHO) or 30 ml of cell suspension (Sf9) were suspended in 20 ml of ice-cold incubation medium (100 mM NaCl, 20 mM Na-HEPES, 10 mM MgCl_2_, pH 7.4) supplemented with 10 mM EDTA and homogenized on ice by two 30-s strokes using a Polytron homogenizer (Ultra-Turrax; Janke & Kunkel GmbH & Co. KG, IKA-Labortechnik, Staufen, Germany) with a 30 s pause between strokes. Cell homogenates were centrifuged for 5 min at 1,000×g to remove whole cells and cell nuclei. The resulting supernatants were centrifuged for 30 min at 30,000×g. Pellets were suspended in a fresh incubation medium, incubated on ice for 30 min, and centrifuged again. The resulting membrane pellets were kept at –80 °C until assayed within 2 months.

### 5.5. Radioligand Binding

All radioligand binding experiments were optimised and carried out according to general guidelines [37]. Briefly, membranes were incubated in 96-well plates at 30 °C in the incubation medium described above. Incubation volume was 400 μl or 800 μl for competition and saturation experiments with [^3^H]NMS, respectively. Approximately 30 μg of membrane proteins per sample were used. N-methylscopolamine binding was measured directly in saturation experiments using six concentrations (30 pM to 1000 pM) of [^3^H]NMS during incubation for 5 hours. For calculations of the equilibrium dissociation constant (K_D_), concentrations of free [^3^H]NMS were calculated by subtraction of bound radioactivity from total radioactivity in the sample and fitting Equation 1 (Data analysis section). The binding of tested ligands was determined in competition experiments with 1 nM [^3^H]NMS, and the inhibition constant K_I_ was calculated according to Equation 3. Incubation time was 15 hours. Non-specific binding was determined in the presence of 1 μM atropine. Incubations were terminated by filtration through Unifilter GF/C glass fibre filtration plates (Perkin Elmer) using a Brandel cell harvester (Brandel, Gaithersburg, MD, USA). Filtration plates were dried in a microwave oven, and then 40 μl of Rotiszint scintillation cocktail was added to each well. The filtration plates were counted in the Wallac Microbeta^2^ scintillation counter.

### 5.6. Inhibition of functional response to carbachol

The inhibitory effects of the tested compounds on the functional response to carbachol were quantified by measuring changes in the accumulation of inositol phosphates. M_2_ and M_4_ receptors that preferentially inhibit cAMP synthesis via G_i/o_ G-proteins were coupled to the phospholipase C-calcium pathway. For this purpose, new CHO cell lines stably expressing promiscuous G_15_ G-protein [38], besides M_2_ or M_4_ receptor, were generated by transfection with pCMV/hygro vector and hygromycin selection. Cells were seeded in 96-well plates, 20 thousand cells per well in 100 µl of DMEM. The next day, DMEM was removed, and 50 µl of DMEM supplemented with 30 nM [^3^H]myo-inositol was added for 12 hours. Then cells were washed with Krebs-HEPES buffer (KHB; 138 mM NaCl; 4 mM KCl; 1.3 mM CaCl_2_; 1 mM MgCl_2_; 1.2 mM NaH_2_PO_4_; 10 mM glucose; 20 mM Na-HEPES; pH = 7.4) supplemented with 10 mM LiCl. Cells were preincubated at 37°C for 30 minutes with the tested compound at the desired concentration, and then carbachol was added for an additional 30 minutes.

Then KHB was removed, and accumulation of inositol phosphates was stopped by the addition of 50 µl of 20 % trichloracetic acid (TCA). Plates were put to 4°C for 1 hour, then 40 µl of TCA extract was transferred to another 96-well plate, mixed with 200 µl of Rotiszint scintillation cocktail and counted in Wallac Microbeta^2^. The rest of the TCA extract was discarded, individual wells were washed with 50 µl of 20 % TCA, 50 µl of 1 M NaOH was added to each well and plates were shaken at room temperature for 15 min. Then 40 µl of NaOH lysate was transferred to another 96-well plate, mixed with 200 µl of Rotiszint scintillation cocktail and counted in Wallac Microbeta^2^. Levels of inositol phosphates were calculated as a fraction of soluble (TCA extract) to total (TCA extract plus NaOH lysate) radioactivity. When the reversibility of inhibition by tested compounds was determined, 30 min preincubation with tested compounds was carried out in the absence of LiCl, then cells were washed 3 times with 100 µl KHB, incubated either 1, 3 or 5 hours in 250 µl of fresh KHB at 37 °C, again washed 3-times with 100 µl of KHB, and finally incubated in the presence of 10 mM LiCl with carbachol for 30 minutes.

Alternatively, functional antagonism of carbachol-stimulated [^35^S]GTPγS binding to membranes prepared from CHO cells expressing individual subtypes of muscarinic receptors was measured. Membranes were preincubated with tested compounds, carbachol and GDP at 30 °C for 15 hours. Then [^35^S]GTPγS was added and samples were incubated for an additional 20 (M_2_, M_4_) or 30 minutes (M_1_, M_3_, M_5_). Final concentration of [^35^S]GTPγS was 500 pM (M_2_, M_4_) or 200 pM (M_1_, M_3_, M_5_). The final concentration of GDP was 20 μM (M_2_, M_4_) or 5 μM (M_1_, M_3_, M_5_). The incubation volume was 200 μl, and the incubation temperature was 30 °C. Incubations were terminated by filtration through Whatman GF/C glass fibre filters (Whatman) using a Brandel cell harvester (Brandel, Gaithersburg, MD, USA). Filters were dried in a microwave oven, and then solid scintillator Meltilex A was melted on the filters (105 °C, 70 s) using a hot plate. The filters were cooled and counted in the Wallac Microbeta scintillation counter.

### 5.7. Data analysis

Data from biological evaluation experiments were processed in Libre Office, analysed and then plotted using the program Grace (http://plasma-gate.weizmann.ac.il/Grace/). Curve fitting was done using Python and the Scipy library. Statistical analysis was performed using the statistical package R (http://www.r-project.org). For non-linear regression analysis, the following equations were used:

#### 5.7.1. [3H]NMS saturation binding

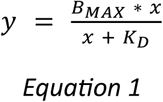

where y is specific binding at free concentration x, B_MAX_ is the maximum binding capacity, and K_D_ is the equilibrium dissociation constant.

#### 5.7.2. Competition binding

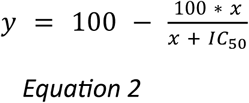

where y is specific radioligand binding at concentration x of competitor expressed as a percentage of binding in the absence of a competitor, and IC_50_ is the concentration causing 50 % inhibition of radioligand binding. The inhibition constant K_I_ was calculated as:

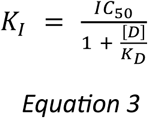

where [D] is a concentration of [^3^H]NMS used [39].

#### 5.7.3. Concentration response

After normalisation to basal, Equation 4 was fitted to data from measurements of functional response.

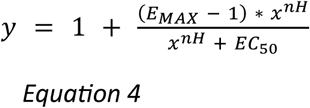

Where y is the response normalised to basal (in the absence of carbachol) at ligand concentration x, E_MAX_ is the maximal effect, EC_50_ is the concentration causing a half-maximal effect, and nH is the Hill coefficient.

The equilibrium dissociation constant of antagonism of functional response K_B_ was calculated from the shift in EC_50_ of functional response to carbachol by the antagonist according to the Schild equation (Equation 5) for a competitive antagonist [40].

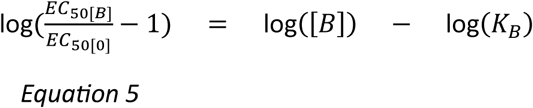

Where EC_50[0]_ is carbachol EC_50_ in the absence of the tested compound, and EC_50[B]_ is carbachol EC_50_ in the presence of the compound at concentration [B].

## Acknowledgements

The work was supported by the Czech Academy of Sciences institutional support [RVO:67985823], MHCZ-DRO (UHHK 00179906), and the Grant Agency of the Czech Republic grant [23-04670S]. Access to computing and storage facilities owned by parties and projects contributing to the Czech National Grid Infrastructure MetaCentrum provided under the programme “Projects of Large Research, Development, and Innovations Infrastructures” (CESNET LM2015042), is appreciated.

## Conflict of interest statement

The authors declare no conflicts of interest.

## Author contributions

All authors contributed to, read and approved the manuscript and are listed in alphabetical order (excluding the first two authors and corresponding authors). AJR and EM contributed equally. AJR designed, supervised and conducted cell culturing, ligand binding and functional experiments and data analysis. EM and BS performed chemical synthesis and analysed the NMR spectra. JB, NC, ED and DN performed cell culturing, ligand binding and functional experiments and raw data analysis. LP carried out LC-MS and HRMS analysis for the newly developed compounds. JJ performed molecular modelling and final data analysis. JK and JJ conceived the project and its experimental design, wrote and finalised the manuscript and secured project financing.

## Notes

### Competing Interest Statement

The authors have declared no competing interest.

### Summary of Updates

Minor changes to the manuscript according to the review in Biomedicine & Pharmacotherapy.

