## Supplementary Information for "Balancing Rigidity and Flexibility: Optimised 4-(Hexyloxy)benzoate Antagonists with Enhanced Affinity and Tuneable Duration at Muscarinic Receptors"

Supplementary Information: **Antagonistic properties of 4-(hexyloxy)benzoate derivatives and their N-methyl ammonium salts at muscarinic acetylcholine receptors**

Eva Dolejší[1a] (ORCID: 0000-0002-5845-182X), Eva Mezeiova[2a] (0000-0002-9986-5017), Jana Bláhová[1b], Nikolai Chetverikov[1] (0000-0003-4462-0319), Alena Janoušková-Randáková[1] (ORCID: 0000-0002-8403-6494), Dominik Nelic[1] (ORCID: 0000-0003-1926-3225), Lukas Prchal[2] (ORCID: 0000-0001-9698-4478), Barbora Svobodova[2] (0000-0001-8099-089X), John Boulos[3] (ORCID: 0000-0001-8561-6418), Jan Korábečný[2]* (ORCID: 0000-0001-6977-7596), Jan Jakubík[1]* (ORCID: 0000-0002-1737-1487)

[1], Department of Neurochemistry, Institute of Physiology, Czech Academy of Sciences, Prague, Czech Republic

[2], Biomedical Research Centre, University Hospital Hradec Kralove, Sokolska 581, 500 05 Hradec Kralove, Czech Republic

[3], Department of Physical Sciences, Barry University, Miami Shores, FL, USA

[a], contributed equally

[b], Current address: Department of Chemistry and Pharmacy, FAU Erlangen-Nürnberg, Germany

### 1. Synthesis

#### 1.1. General information

All solvents and chemical reagents were used in the highest available purity without further purification, and they were purchased from Sigma-Aldrich (Prague, Czech Republic). The reactions were monitored by thin-layer chromatography (TLC) on silica gel plates (60 F254, Merck, Prague, Czech Republic), and the spots were visualised by ultraviolet light (254 nm). Purification of crude products was carried out using columns of silica gel (silica gel 100, 0.063-0.200 mm, 70-230 mesh ASTM, Fluka, Prague, Czech Republic). NMR spectra were recorded in deuterated chloroform ($CDCl_3$) on a Varian S500 spectrometer. Chemical shifts (*δ*) are reported in parts per million (ppm), and spin multiplicates are given as first-order multiplets [singlet (s), triplet (t)] or more complex patterns involving different coupling constants [multiplet (m), doublet of doublets (dd), dt, doublet of quartets (dq), td, tt]. Coupling constants (*J*) are reported in Hz. The synthesised compounds were analysed by the LC-MS system consisting of UHPLC Dionex Ultimate 3000 coupled with Q Exactive™ Plus mass spectrometer to obtain high-resolution mass spectra (Thermo Fisher Scientific, Bremen, Germany).

#### 1.2. Syntheses of intermediates 3, 4 and 8k,l

##### 1.2.1. Preparation of aldehyde 3 and acid 4

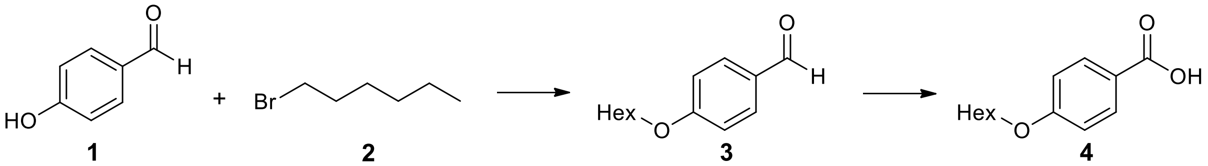

**1** **2** **3** **4**

**3** was prepared by s synthetic procedure taken from ref. [1]. NMR spectra were in good agreement with previously published data: $^1$H NMR (500 MHz, $CDCl_3$): *δ* 9.87 (s, 1H), 7.83–7.80 (m, 2H), 6.99–6.97 (m, 2H), 4.03 (t, *J* = 6.6 Hz, 2H), 1.83–1.78 (m, 2H), 1.50–1.43 (m, 2H), 1.37–1.33 (m, 4H), 0.92–0.89 (m, 3H). $^{13}$C NMR (126 MHz, $CDCl_3$): *δ* 190.9, 164.4, 132.1, 129.9, 114.9, 68.6, 31.6, 29.2, 25.8, 22.7, 14.1.

To a solution of aldehyde **3** (1.0 eq) in a mixture of solvents ($THF/H_2O/^tBuOH$ = 4/4/1, 0.1 M) were gradually added $NaH_2PO_4.H_2O$ (8.0 eq), 2-methyl-2-butene (10.0 eq), and $NaClO_2$ (4.0 eq). The resulting mixture was stirred for 1.5 hours at RT. The mixture was diluted with EA (10 × volume of the mixture of solvents put in the reaction) and 3 times washed with a 1M solution of HCl 1(⅓ × volume of EA). The organic layer was dried over $Na_2SO_4$, and the solvents were evaporated under reduced pressure. The crude product was used in subsequent reactions without any purification.

##### 1.2.2. Preparation of 2-(2,3-dihydro-1H-isoindol-2-yl)ethan-1-ol (8k)

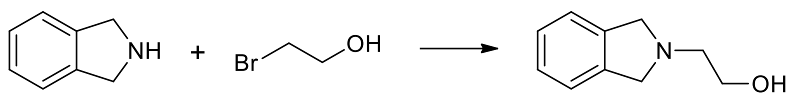

isoindoline 2-bromoethanol **8k**

**8k** was prepared by a modified procedure taken from ref.[2]. To a solution of isoindoline (1.0 eq) and 2-bromoethanol (1.0 eq) in dry toluene (0.25 M) and $Et_3N$ (2.0 eq) was added. The mixture was stirred overnight at reflux and then cooled to RT. The solvent was evaporated under reduced pressure, and the residue was dissolved in DCM (1.5 × volume of toluene put in the reaction) and washed with water (1 × volume of DCM). The water layer was again washed with DCM (1 x volume of water). Combined organic layers were dried over anhydrous $Na_2SO_4$, and the solvent was evaporated under reduced pressure. The crude product was purified by column chromatography (DCM/MeOH/$NH_3$ = 20:1:0.1. $^1$H NMR (500 MHz, $CDCl_3$): $\delta$ 7.21 (s, 3H), 3.99 (s, 4H), 3.72–3.70 (m, 2H), 2.93 (td, $J$ = 5.4, 0.9 Hz, 2H). $^{13}$C NMR (126 MHz, $CDCl_3$): $\delta$ 139.9, 127.0, 122.4, 60.0, 59.0, 57.6. These spectra are in good agreement with previously published data.[3]

##### 1.2.3. Preparation of 2-(1,2,3,4-tetrahydroquinolin-1-yl)ethan-1-ol (8l)

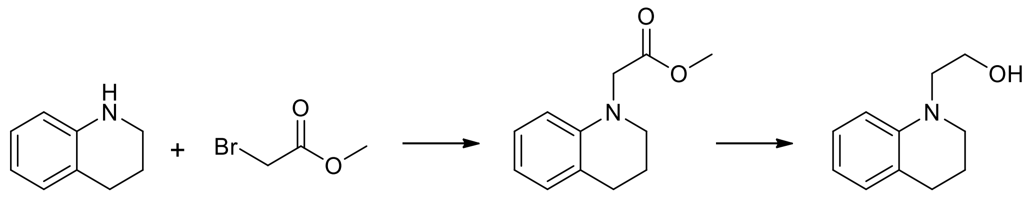

1,2,3,4-tetrahydro-quinoline methyl bromo-acetate **9** **8l**

**8l** was prepared by a modified two-step procedure taken from ref. [4]. To a solution of 1,2,3,4-tetrahydroquinoline (1.0 eq) in dry $CH_3CN$ and KI (2.4 eq), $K_2CO_3$ (2.0 eq), and methyl 2-bromoacetate (1.2 eq) were added. The resulting mixture was stirred for 2 days at reflux and then cooled to RT. The solid part was filtered off, and the solvent was evaporated under reduced pressure. The rest was diluted in EA (10 × volume of $CH_3CN$ put in the reaction) and washed with water (½ volume of EA) and brine (½ volume of EA). The organic layer was dried over anhydrous $Na_2SO_4$, and the solvent was evaporated. The crude product **9** was purified by column chromatography (PE/EA = 10:1).

To a cooled solution of **9** (1.0 eq) in dry THF (1M) was added a 2M solution of LAH (1.25 eq). The mixture was stirred overnight at RT. The mixture was then slowly diluted with a saturated aqueous solution of ammonium chloride (4 × volume of THF put in the reaction) and 3 times washed with DCM (1 × volume of ammonium chloride solution). Combined organic layers were dried over anhydrous $Na_2SO_4$, and the solvent was evaporated under reduced pressure. The crude product **8l** was purified by column chromatography (PE/EA = 7:3).

*Methyl 2-(1,2,3,4-tetrahydroquinolin-1-yl)acetate (**9**):* $^{1}$H NMR (500 MHz, $CDCl_3$): $\delta$ 7.05–7.01 (m, 1H), 6.98 (dt, $J$ = 7.3, 1.4 Hz, 1H), 6.63 (tt, $J$ = 7.3, 1.2 Hz, 1H), 6.41 (dt, $J$ = 8.2, 1.2 Hz, 1H), 4.02 (s, 2H), 3.73 (s, 3H), 3.42–3.39 (m, 2H), 2.80 (t, $J$ = 6.4 Hz, 2H), 2.03–1.98 (m, 2H). $^{13}$C NMR (126 MHz, $CDCl_3$): $\delta$ 171.7, 144.8, 129.3, 127.2, 122.9, 117.0, 110.4, 53.2, 52.0, 50.7, 28.0, 22.4. NMR spectra were in good agreement with previously published data.[4]

*2-(1,2,3,4-tetrahydroquinolin-1-yl)ethan-1-ol (**8l**)*: $^{1}$H NMR (500 MHz, $CDCl_3$): $\delta$ 7.07–7.04 (m, 1H), 6.97 (dq, $J$ = 7.3, 1.2 Hz, 1H), 6.69 (dd, $J$ = 8.3, 1.0 Hz, 1H), 6.62 (tt, $J$ = 7.3, 1.2 Hz, 1H), 3.82 (t, $J$ = 5.8 Hz, 2H), 3.45 (t, $J$ = 5.8 Hz, 2H), 3.34–3.32 (m, 2H), 2.78 (t, $J$ = 6.4 Hz, 2H), 2.00–1.95 (m, 2H), 1.86–1.82 (m, 1H). $^{13}$C NMR (126 MHz, $CDCl_3$): $\delta$ 146.0, 129.5, 127.2, 123.0, 116.6, 111.5, 60.0, 54.3, 50.5, 28.2, 22.4. NMR spectra were in good agreement with previously published data.[4]

#### 2. [1]H and [13]C NMR spectra

[1]H and [13]C NMR spectra of 4-(hexyloxy)benzaldehyde (**3**)

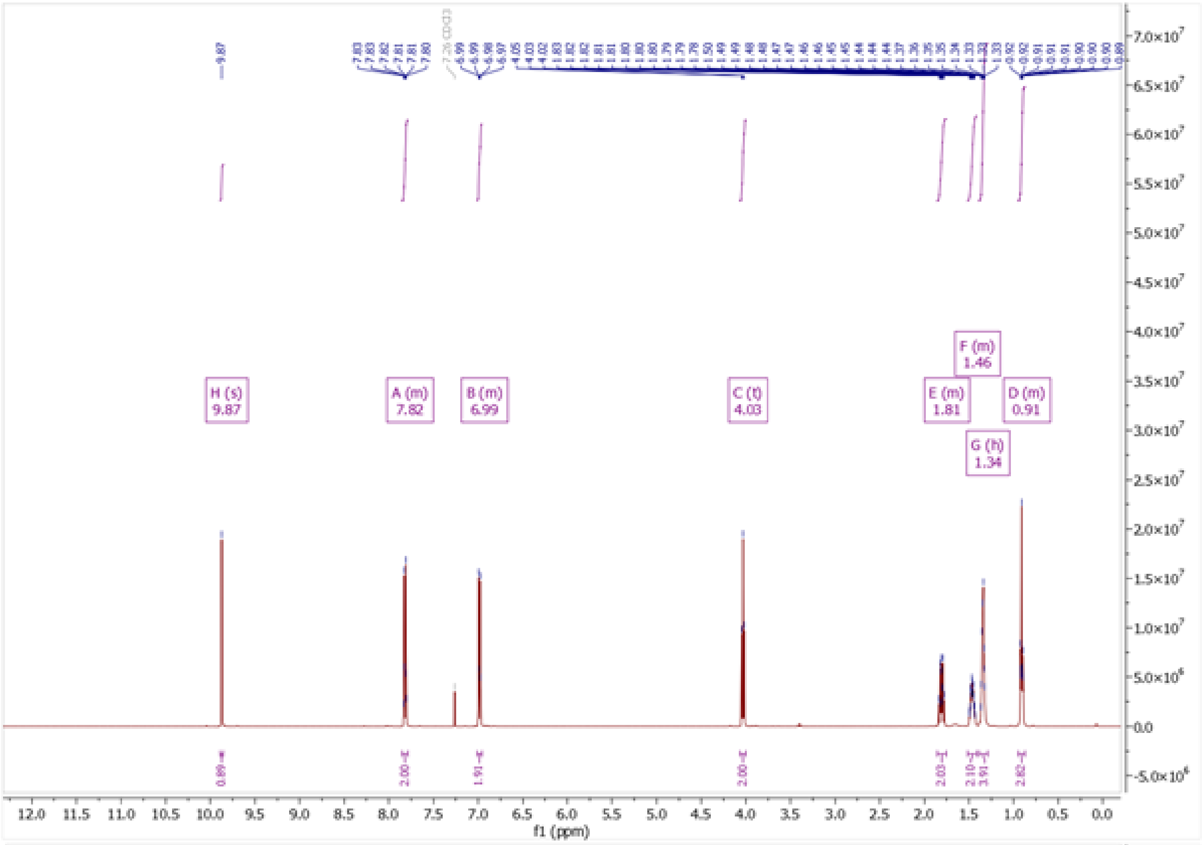

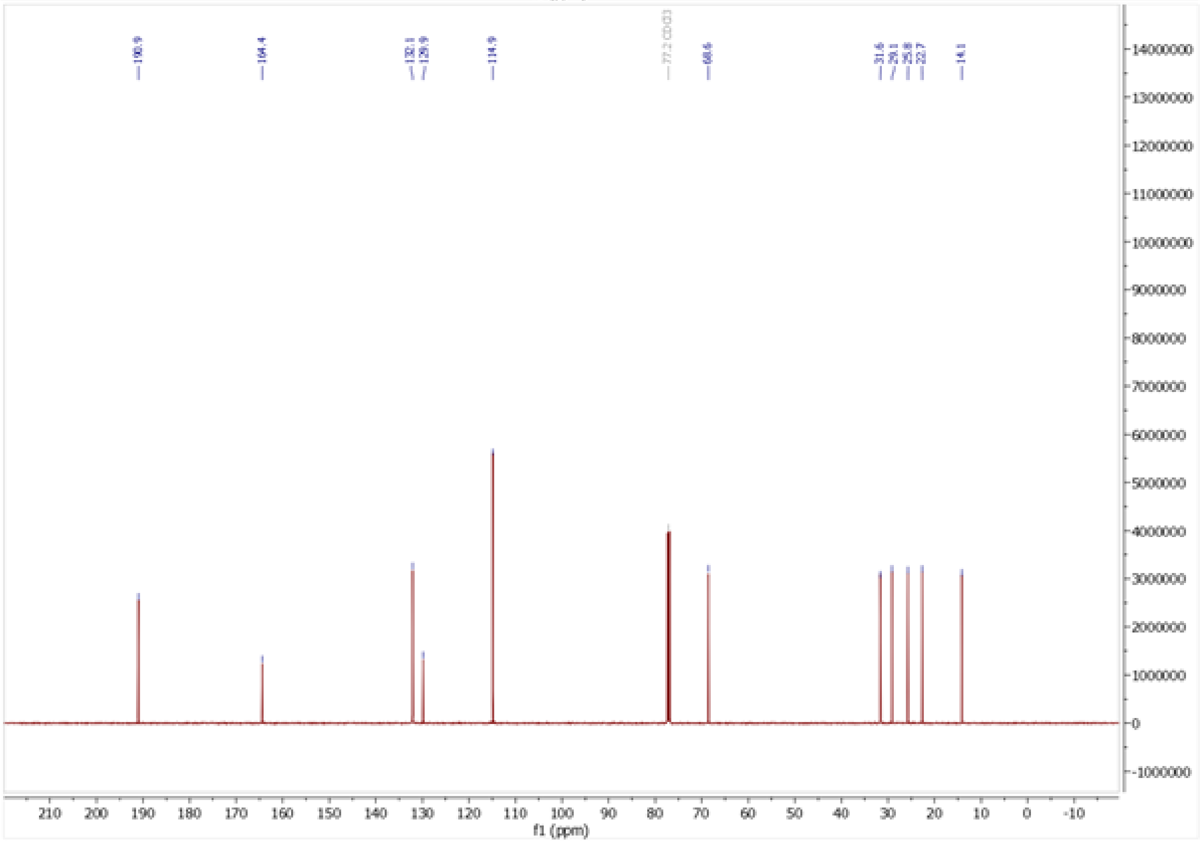

[1]H and [13]C NMR spectra of 2-(2,3-dihydro-1H-isoindol-2-yl)ethan-1-ol (**8k**)

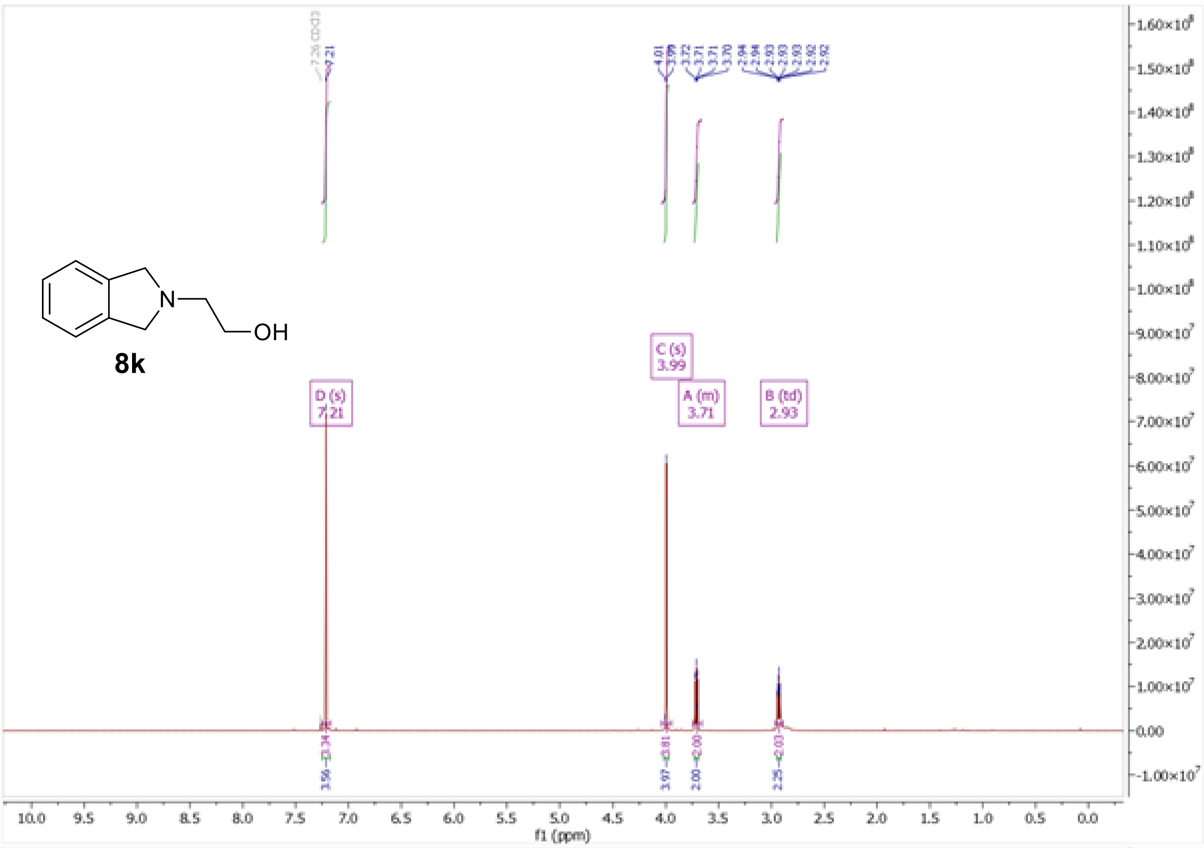

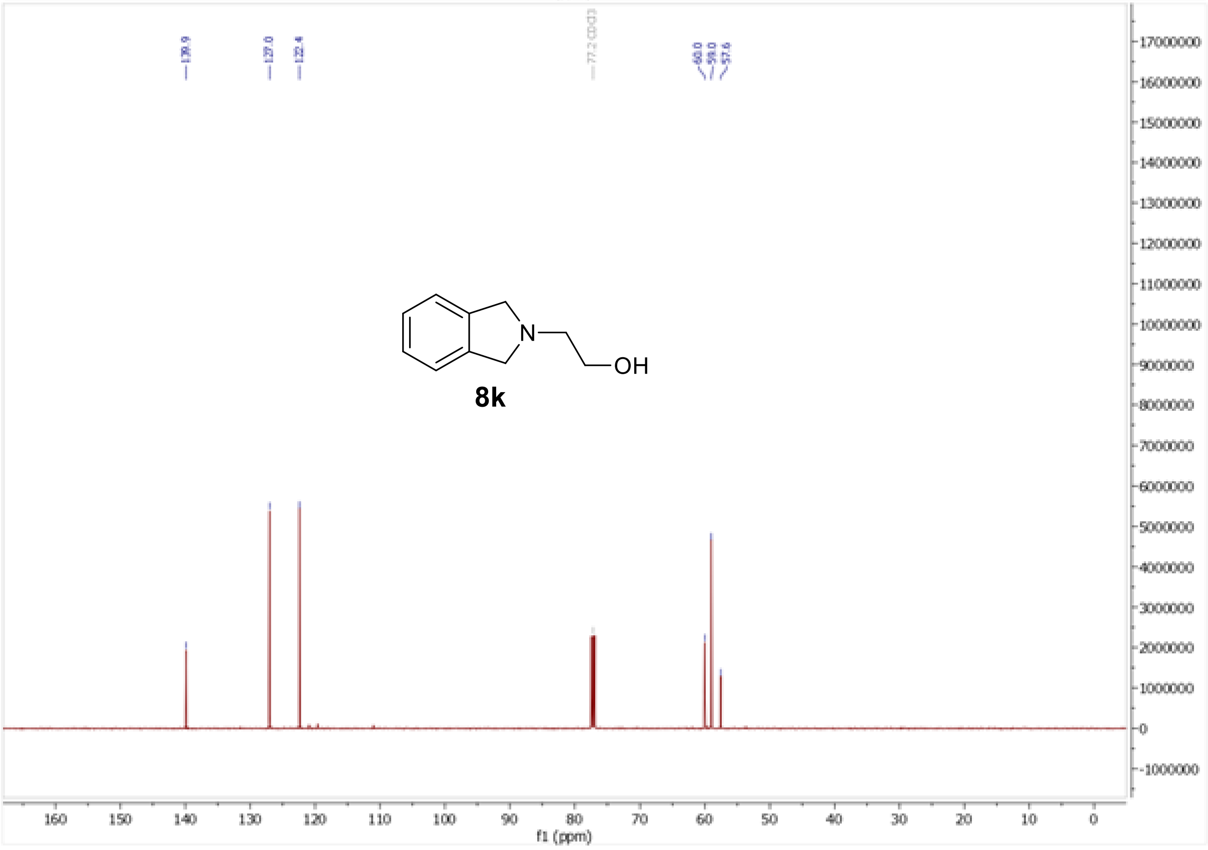

[1]H and [13]C NMR spectra of methyl 2-(1,2,3,4-tetrahydroquinolin-1-yl)acetate (**9**)

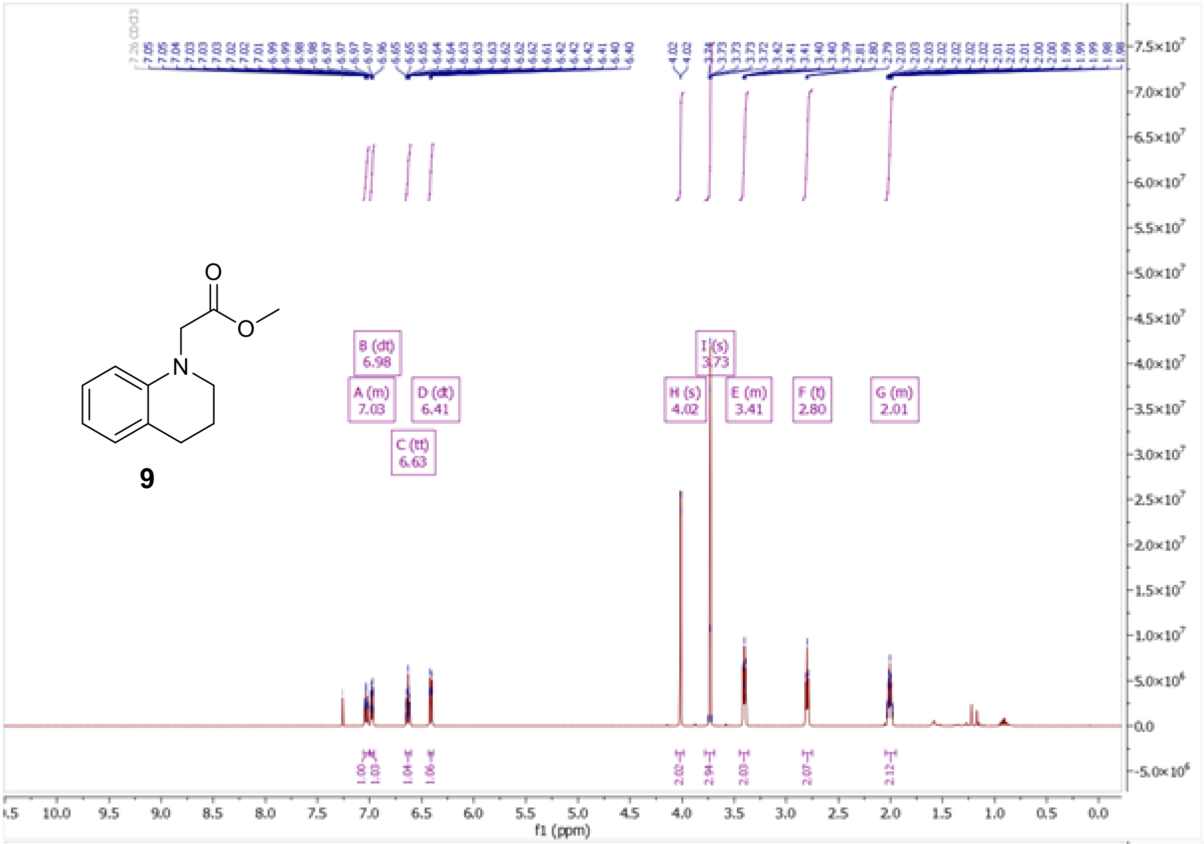

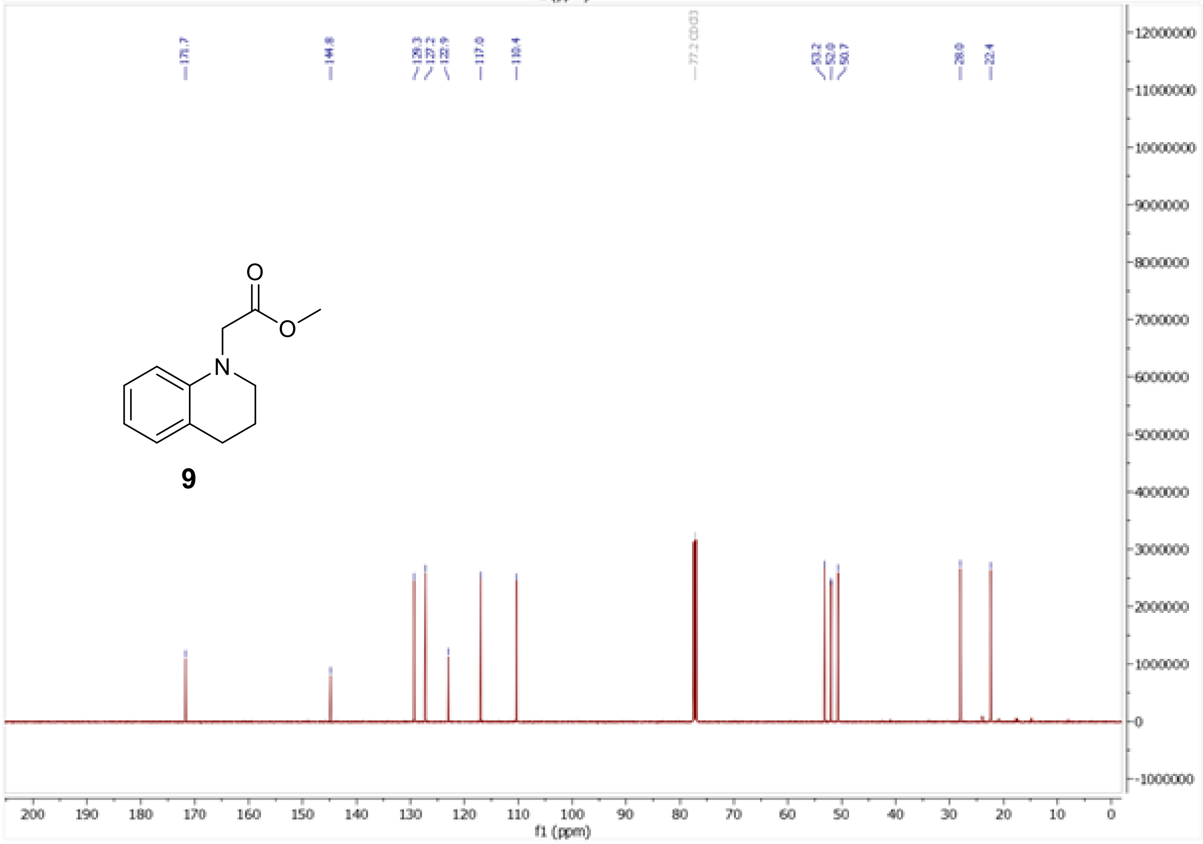

1H and 13C NMR spectra of 2-(1,2,3,4-tetrahydroquinolin-1-yl)ethan-1-ol (**8l**)

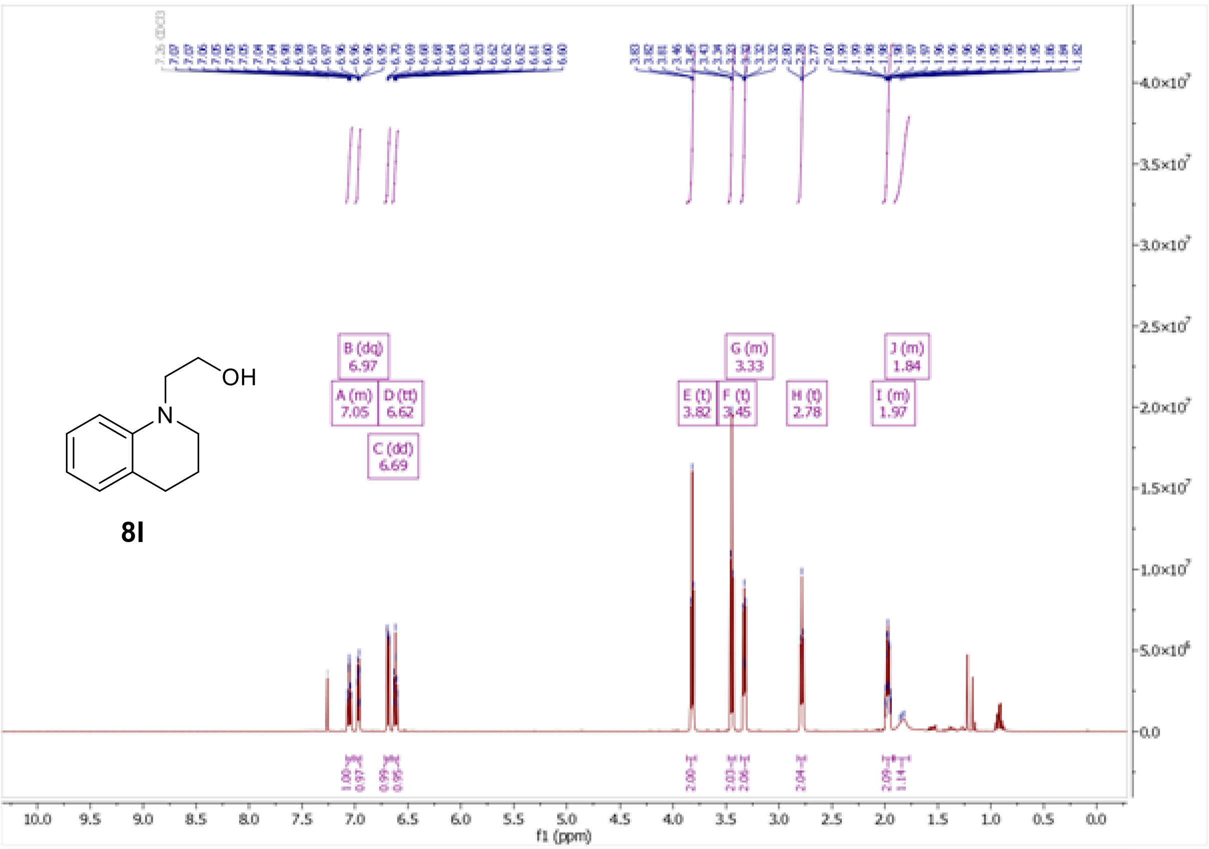

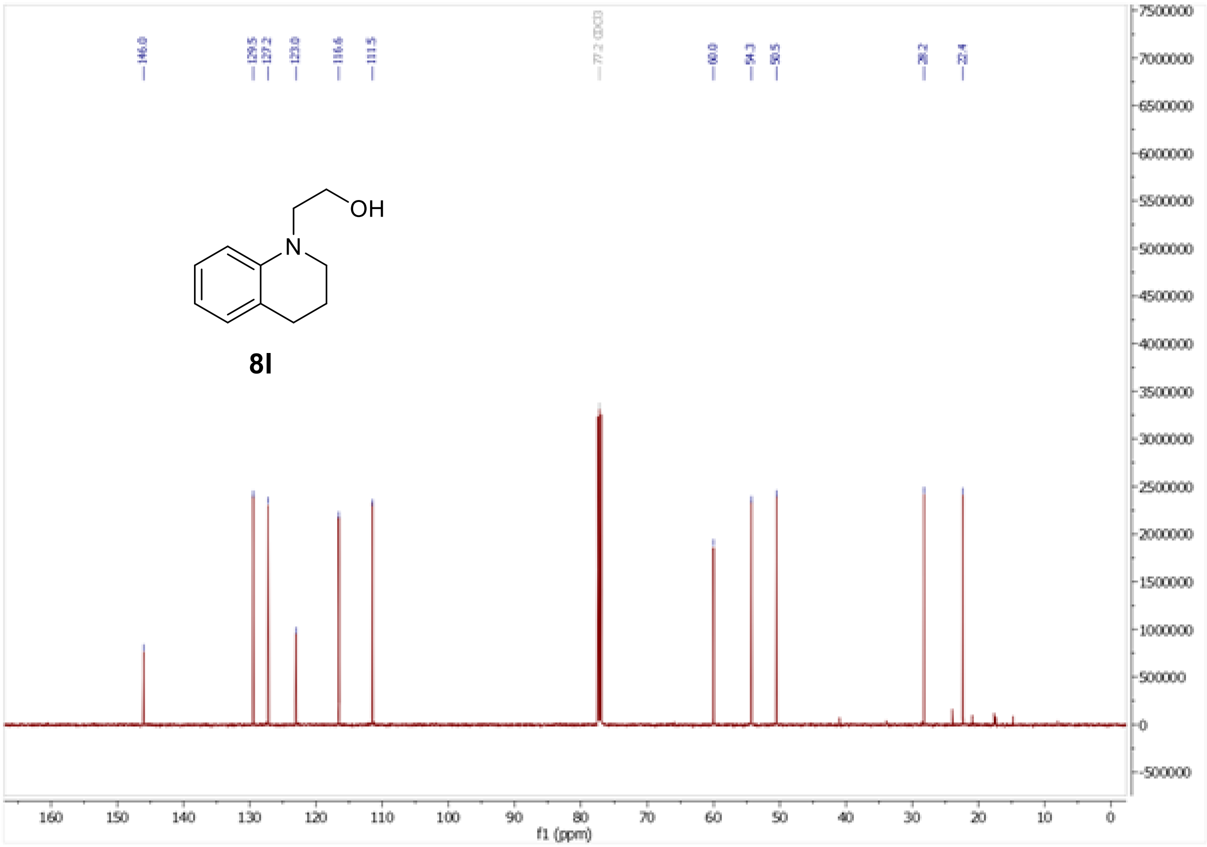

[1]H and [13]C NMR of 2-(1,3-dioxo-2,3-dihydro-1H-isoindol-2-yl)ethyl 4-(pentyloxy)benzoate (**6a**)

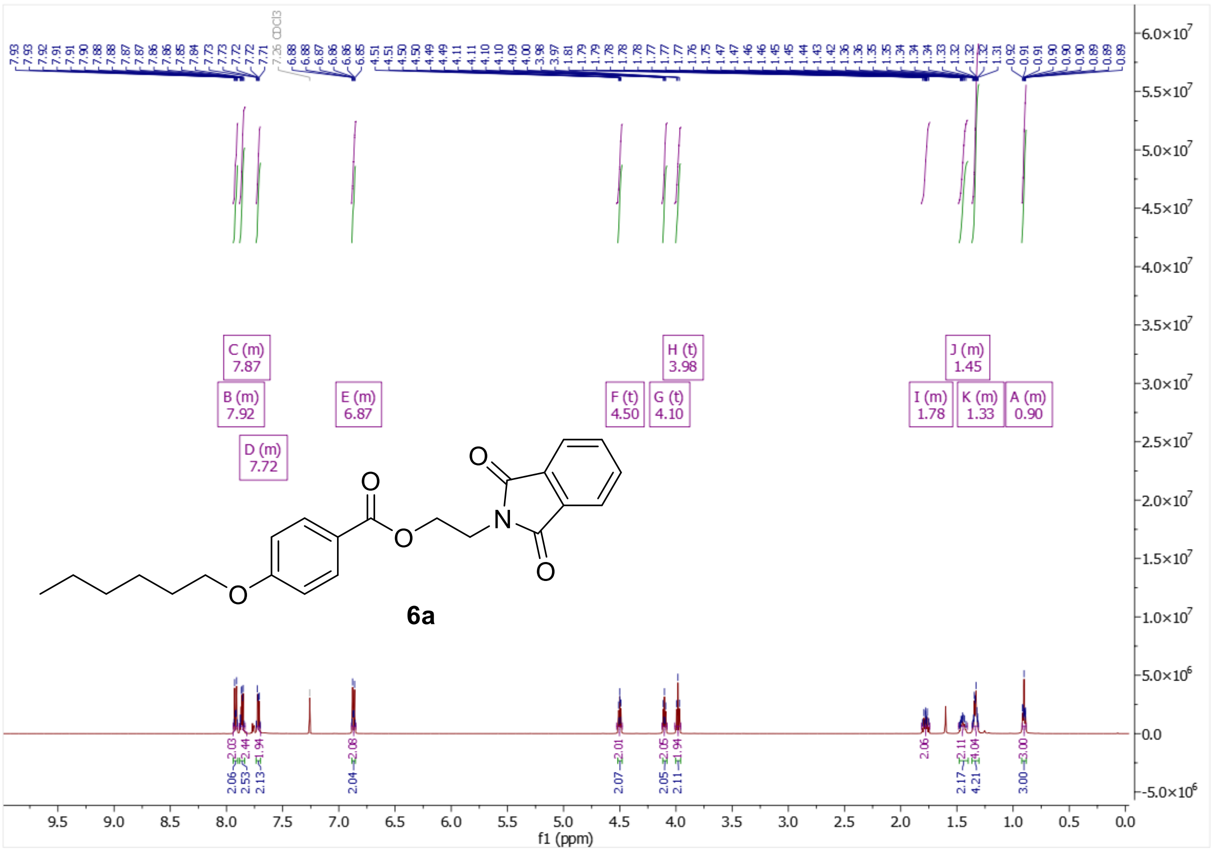

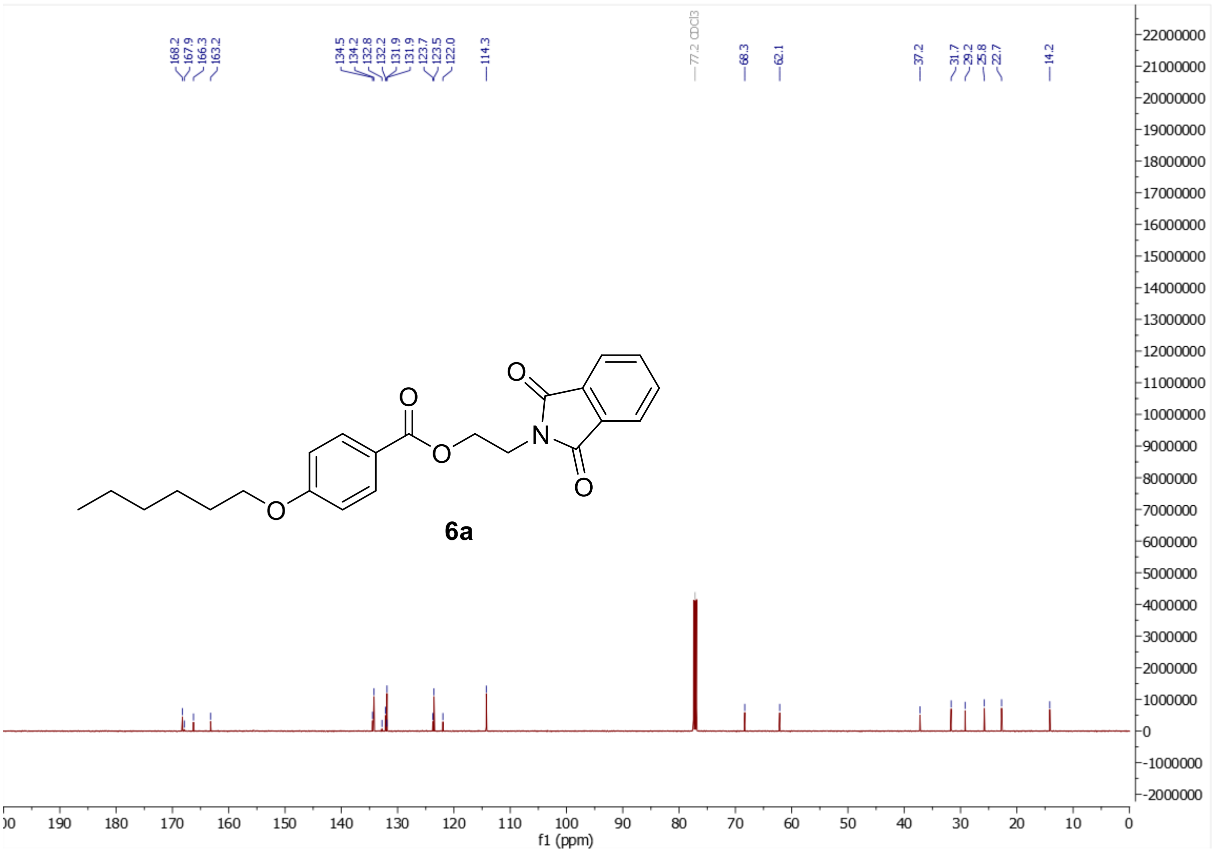

$^1$H and $^{13}$C NMR of 1-{2-[4-(hexyloxy)benzoyloxy]ethyl}-1-azabicyclo[2.2.2]octan-1-ium (**6b**)

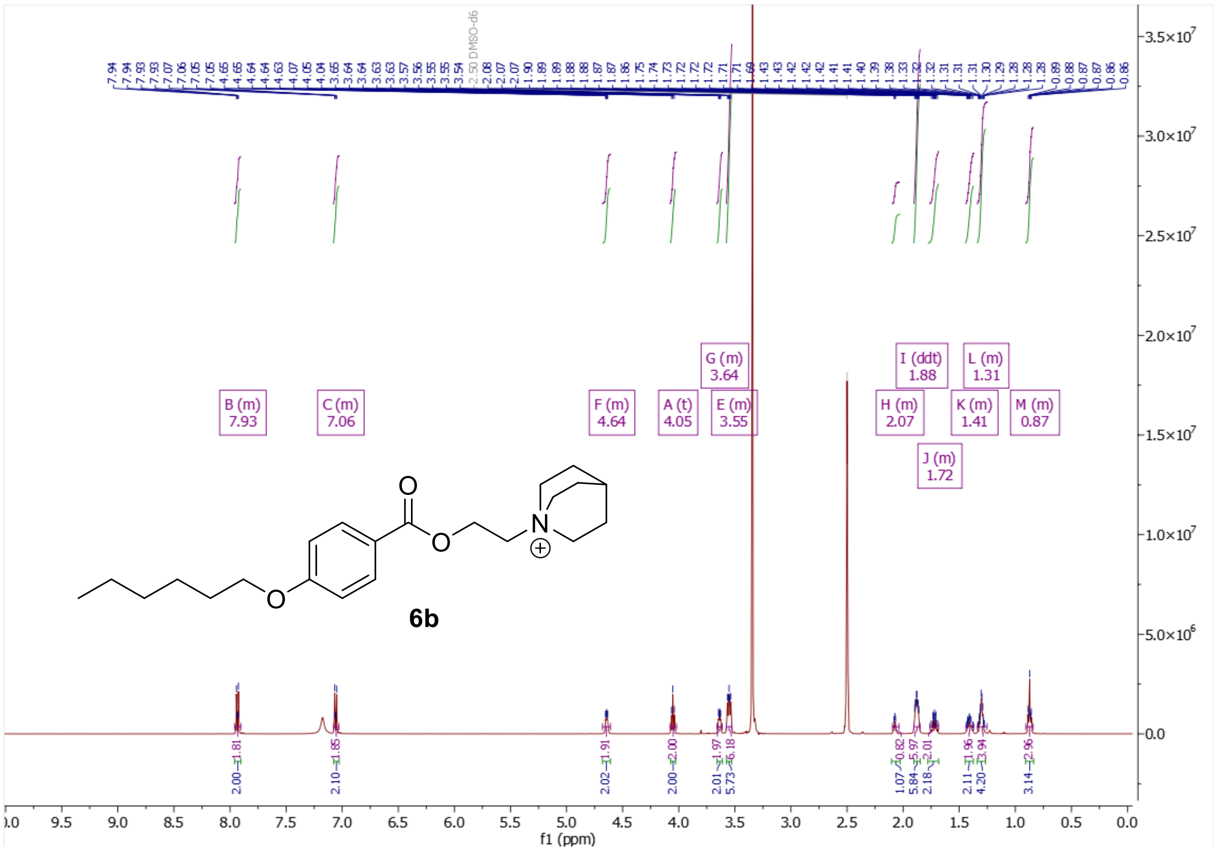

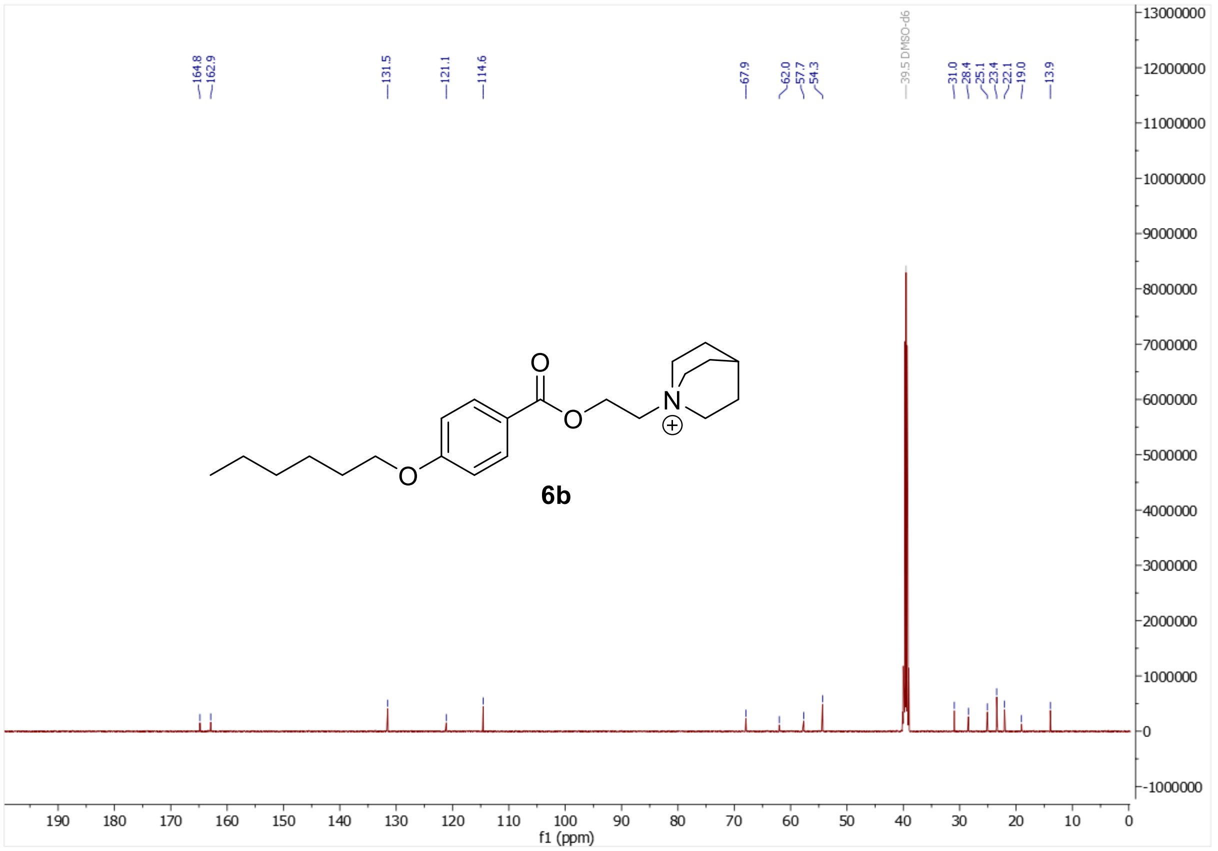

$^1H$ and $^{13}C$ NMR of 2-(2,5-dioxopyrrolidin-1-yl)ethyl 4-(hexyloxy)benzoate (**6c**)

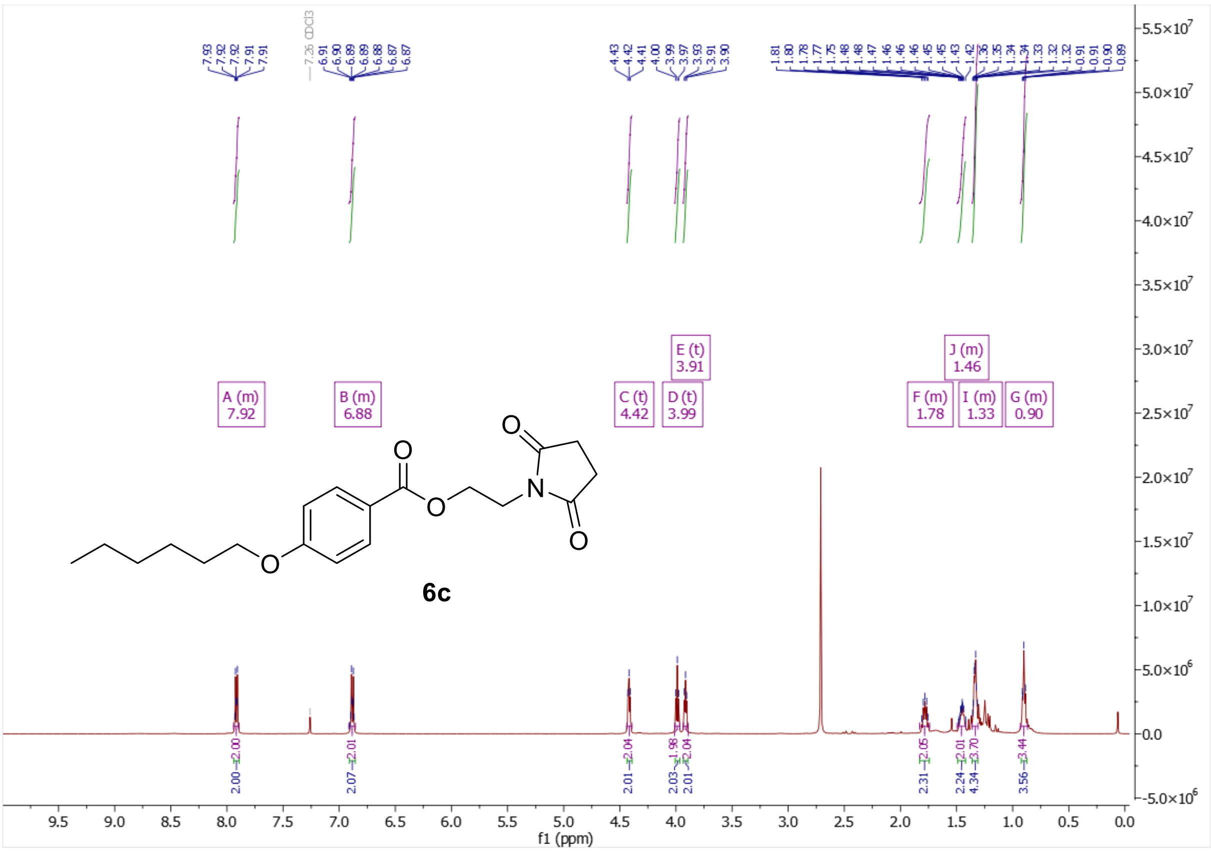

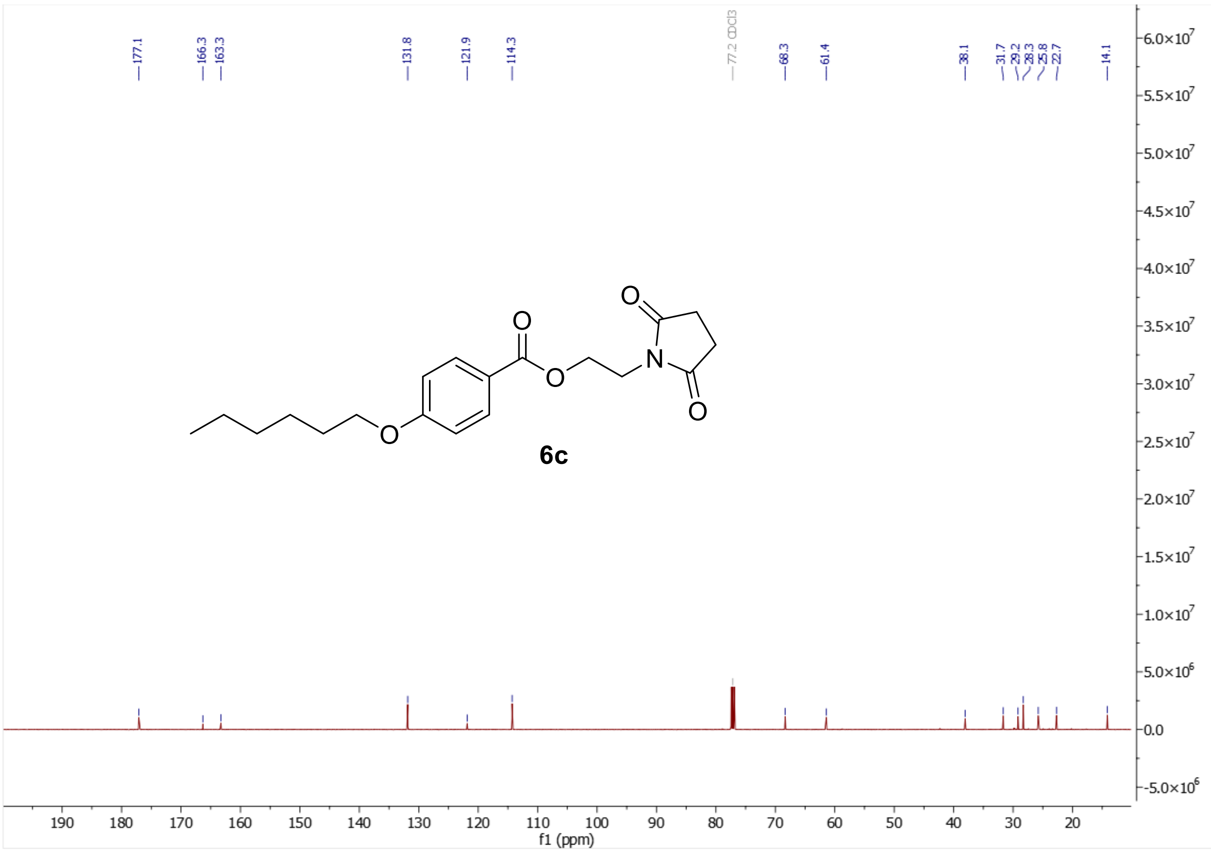

$^{1}H$ and $^{13}C$ NMR of 2-{3,5-dioxo-10-oxa-4-azatricyclo[5.2.1.0$^{2,6}$]dec-8-en-4-yl}ethyl 4-(hexyloxy)benzoate (**6d**)

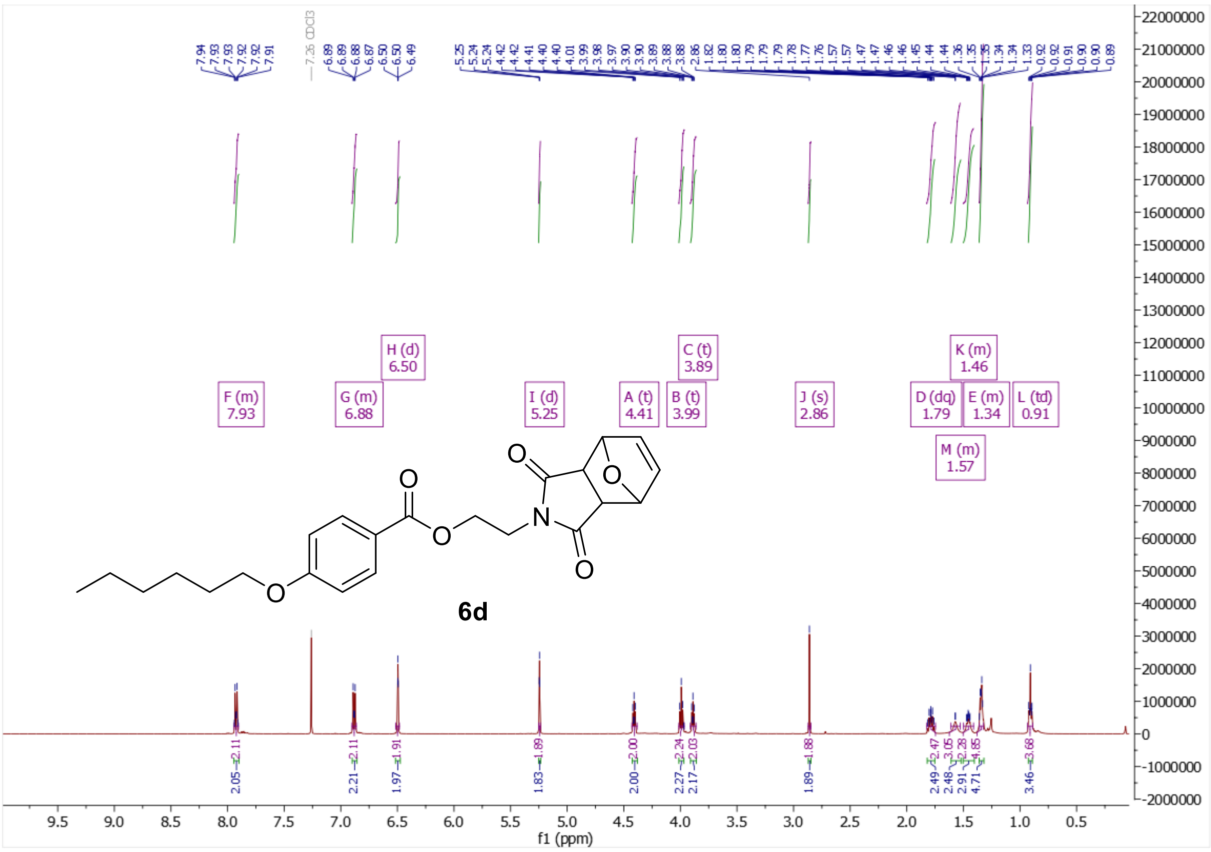

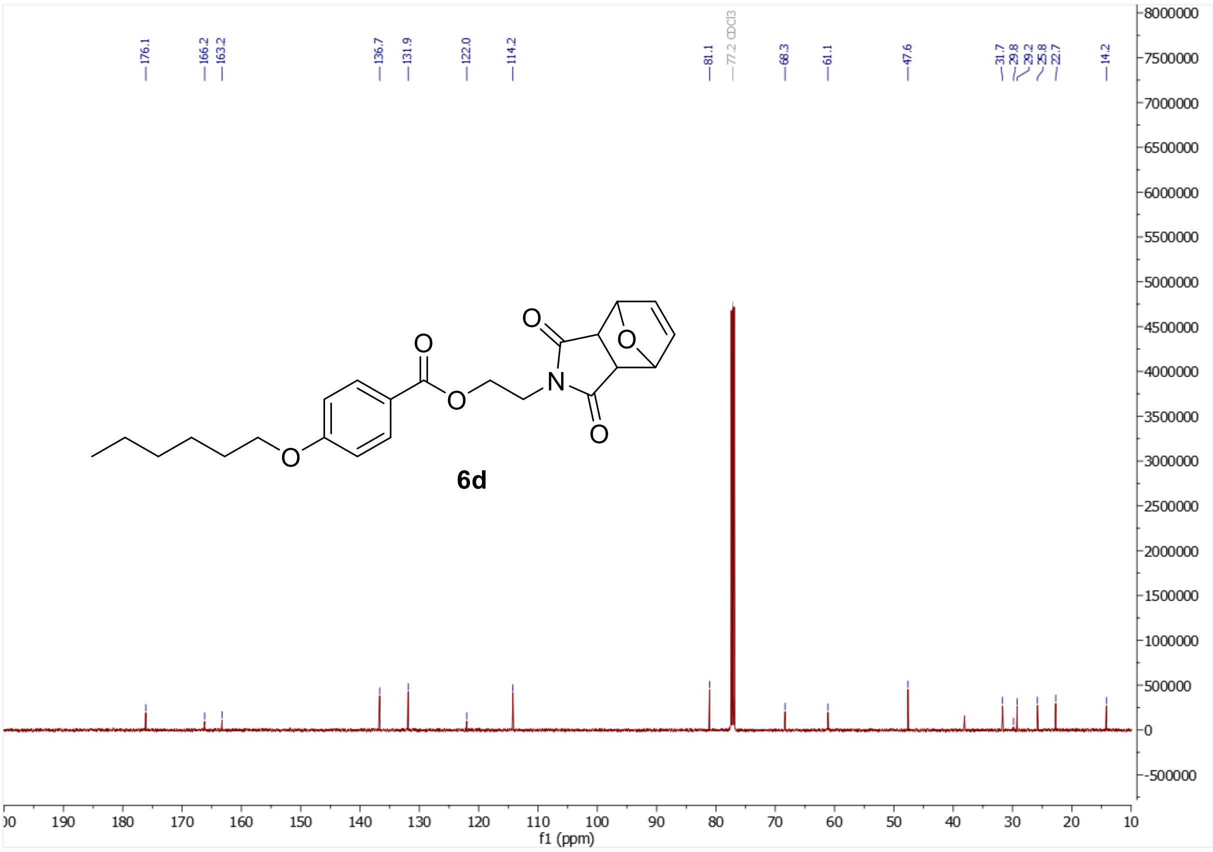

$^{1}H$ and $^{13}C$ NMR of 2-[(2*R*)-2-methylpiperidin-1-yl]ethyl 4-(hexyloxy)benzoate (**6e**)

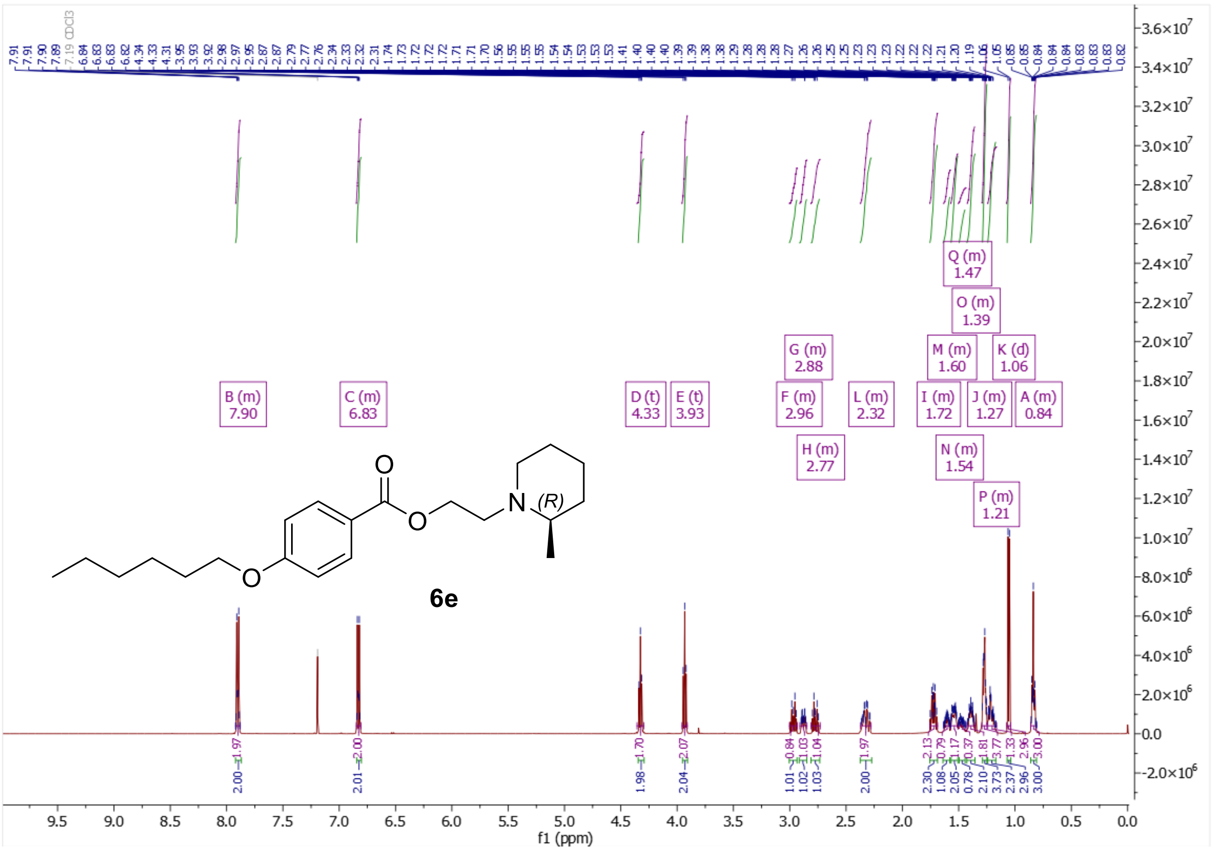

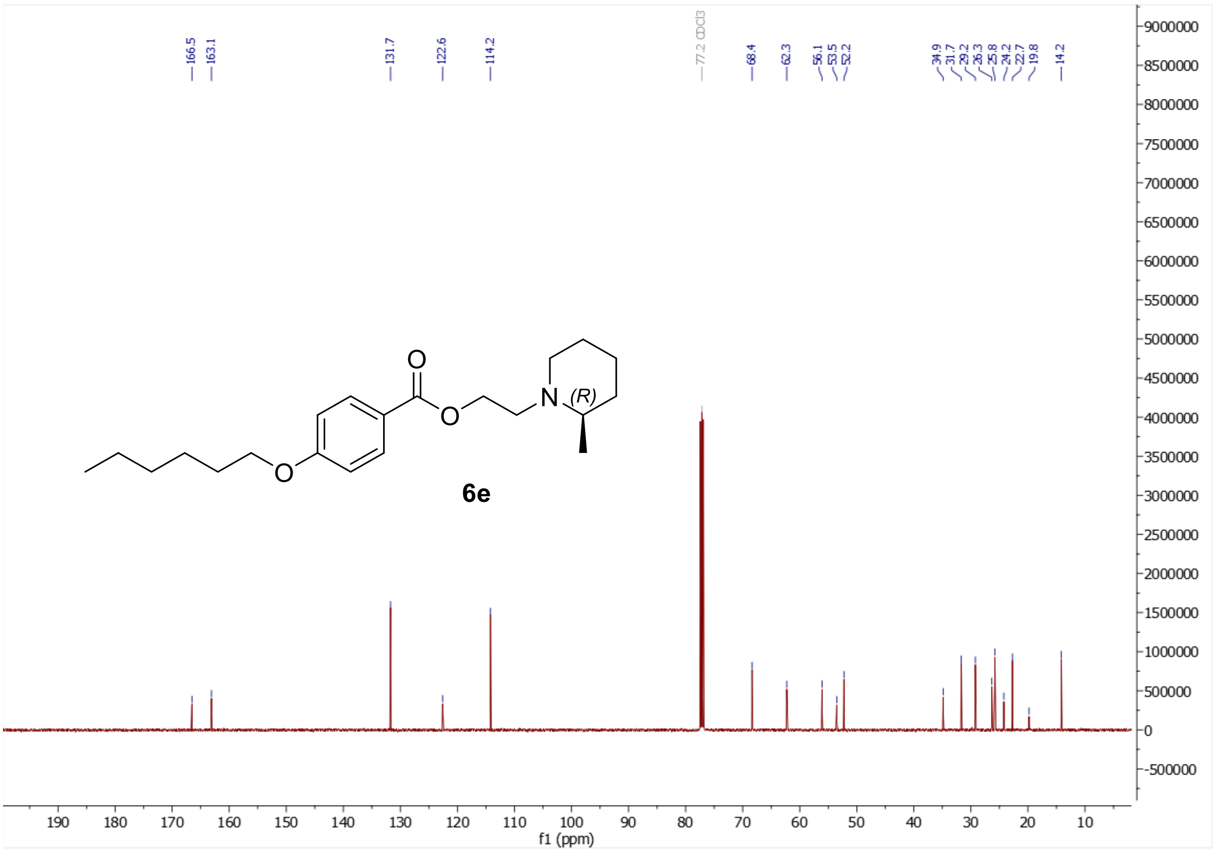

[1]H and [13]C NMR of 2-[(2*S*)-2-methylpiperidin-1-yl]ethyl 4-(hexyloxy)benzoate (**6f**)

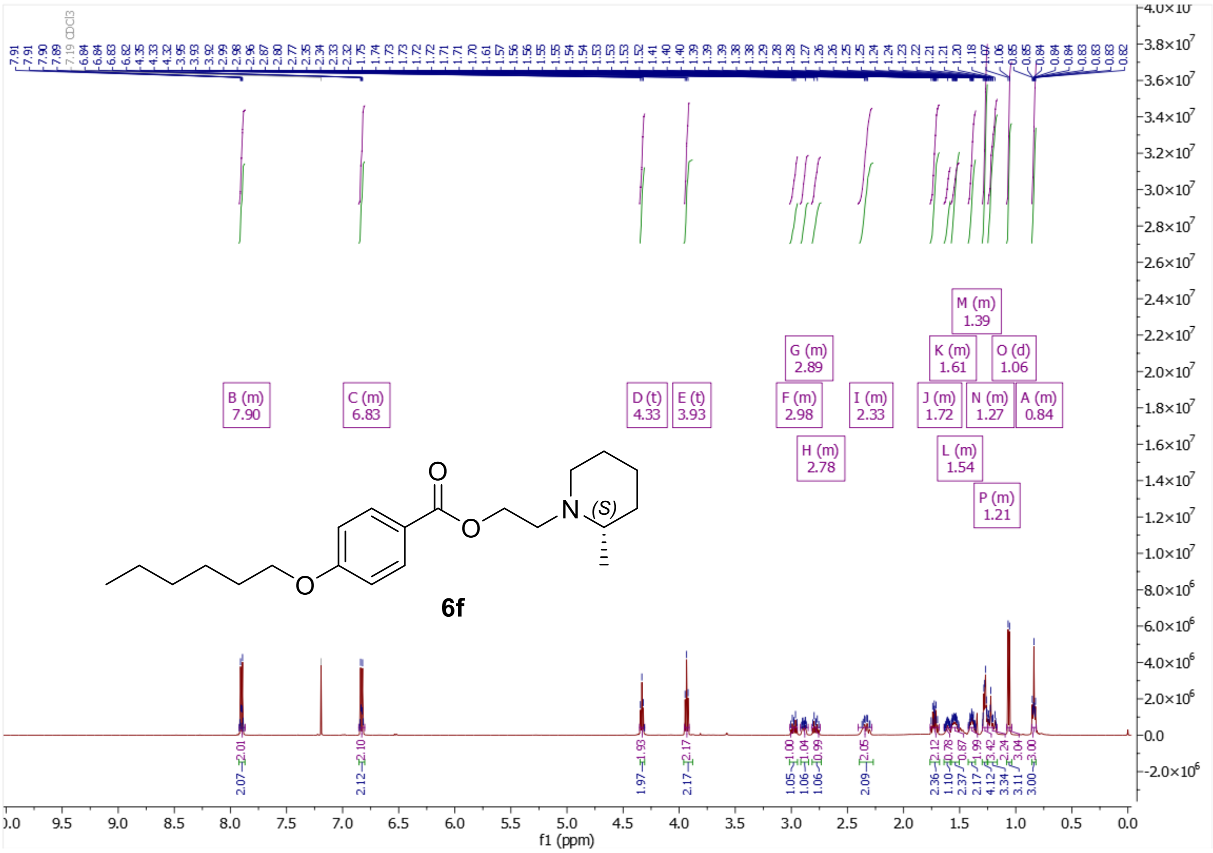

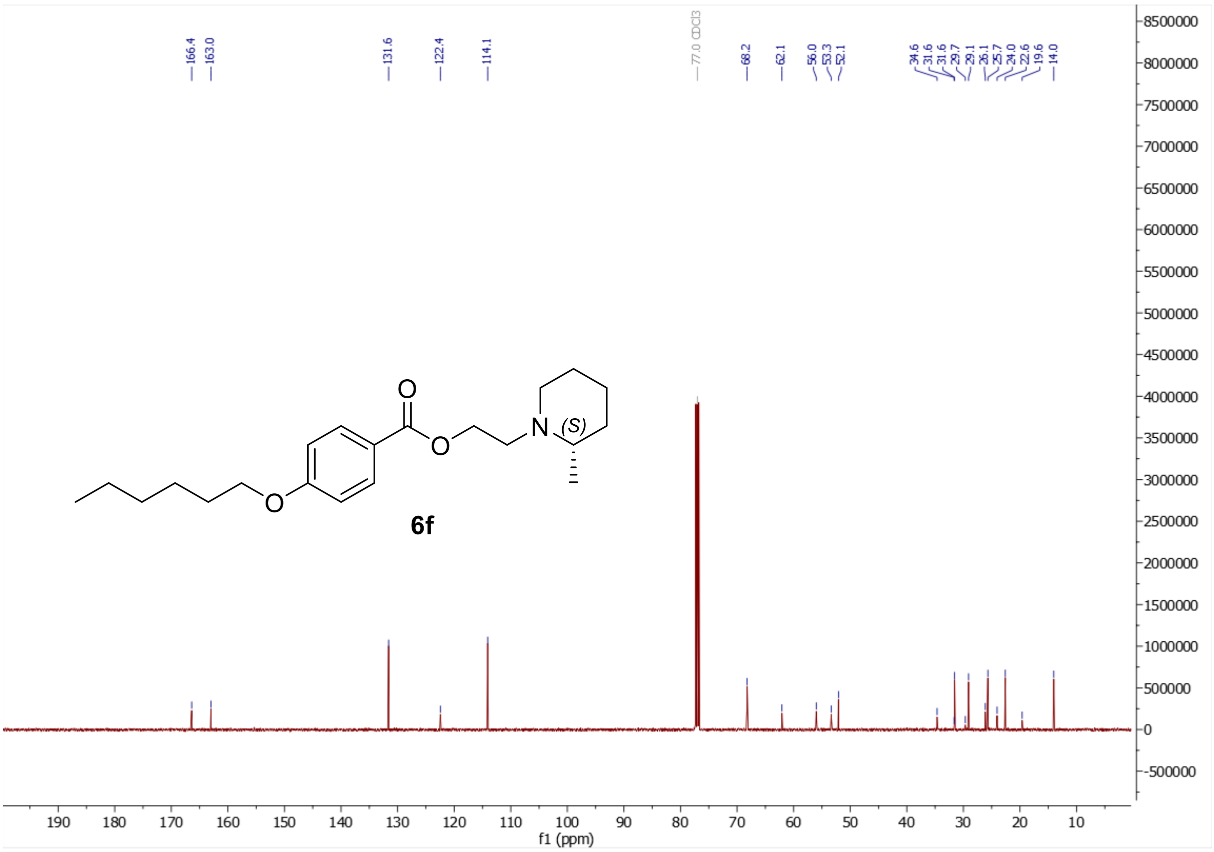

$^{1}H$ and $^{13}C$ NMR of 2-{2-azabicyclo[2.2.2]octan-2-yl}ethyl 4-(hexyloxy)benzoate (**6g**)

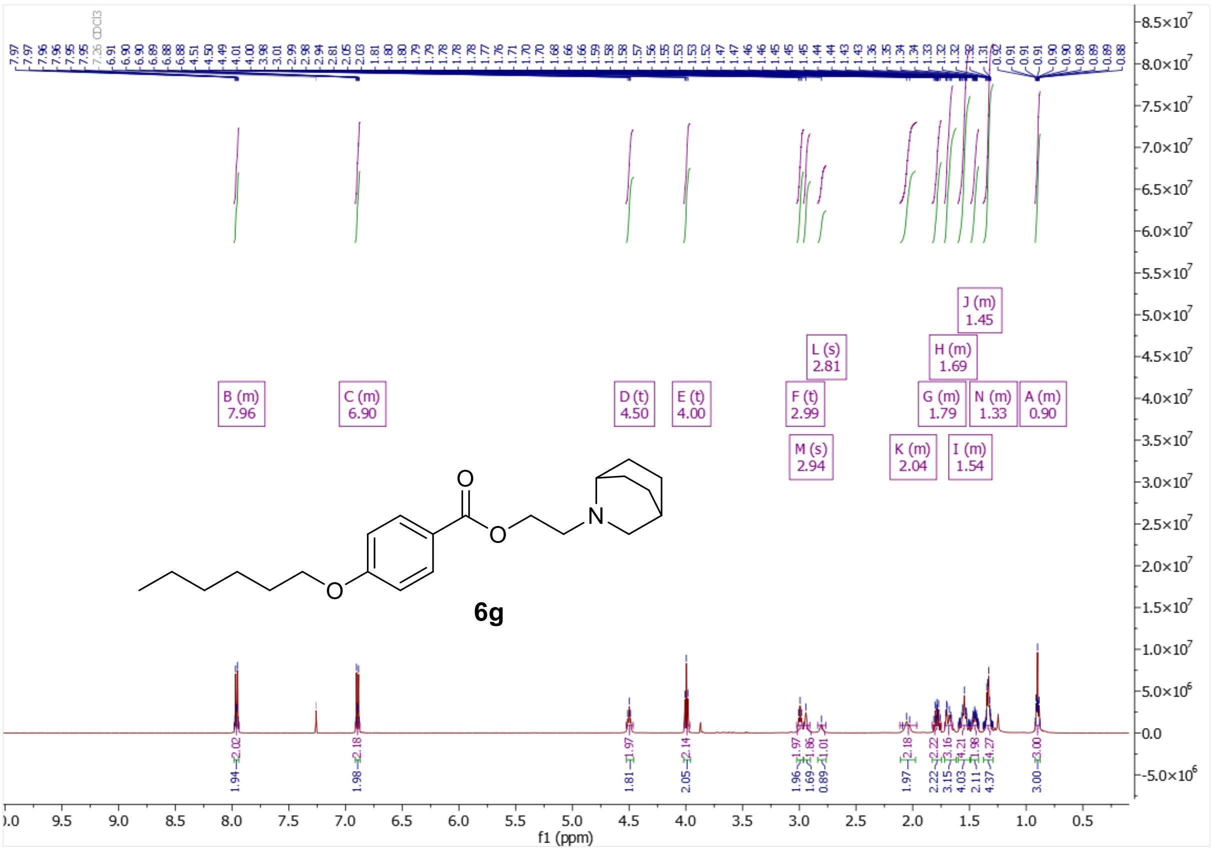

6g

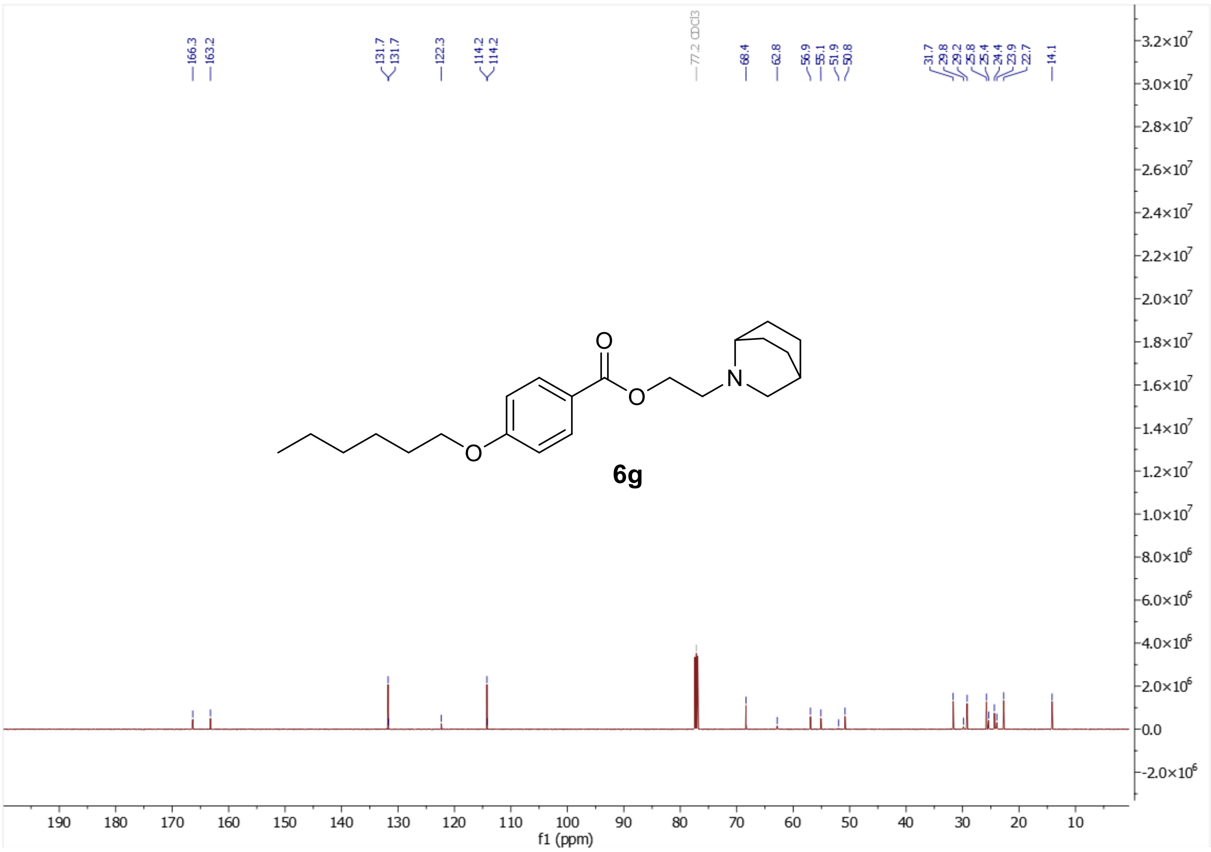

6g

$^{1}H$ and $^{13}C$ NMR of 2-(2,5-dihydro-1H-pyrrol-1-yl)ethyl 4-(hexyloxy)benzoate (**6h**)

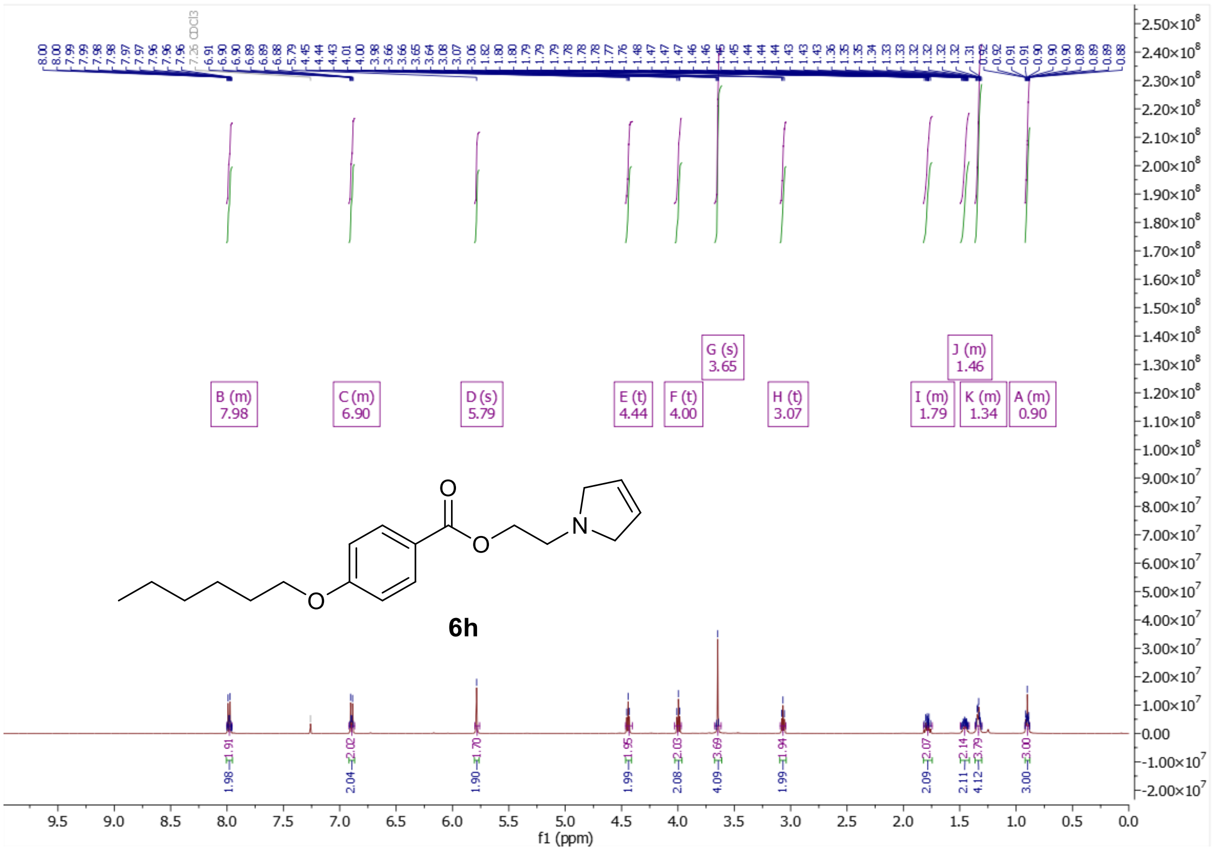

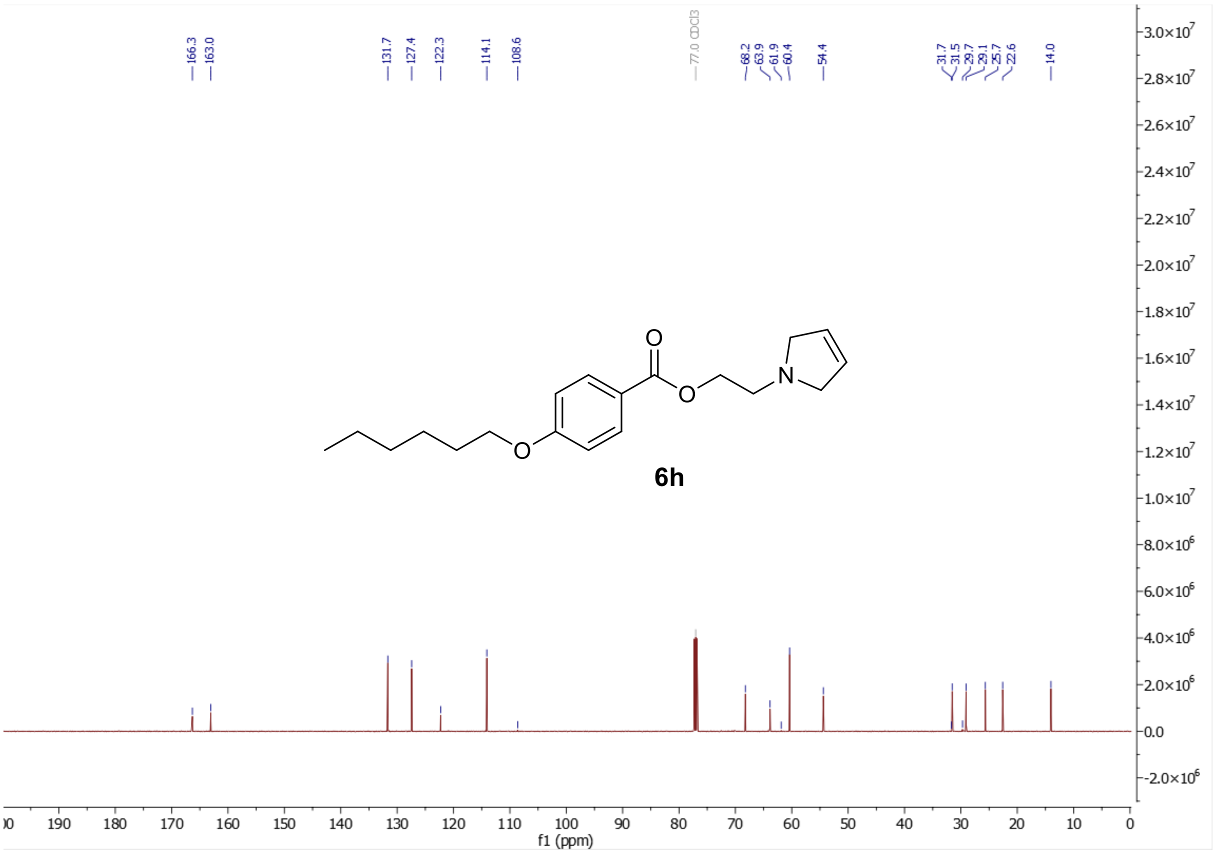

$^{1}H$ and $^{13}C$ NMR of 2-(pyrrolidin-1-yl)ethyl 4-(hexyloxy)benzoate (**6j**)

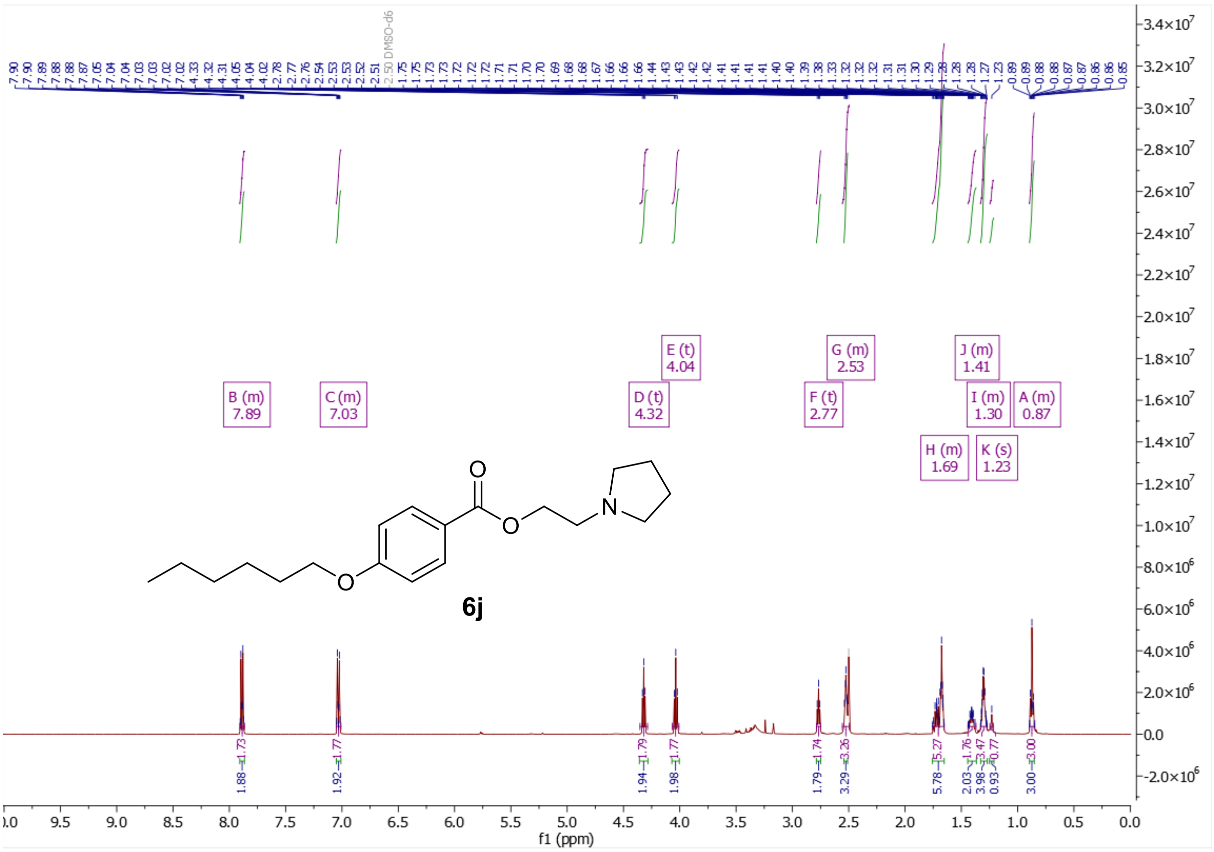

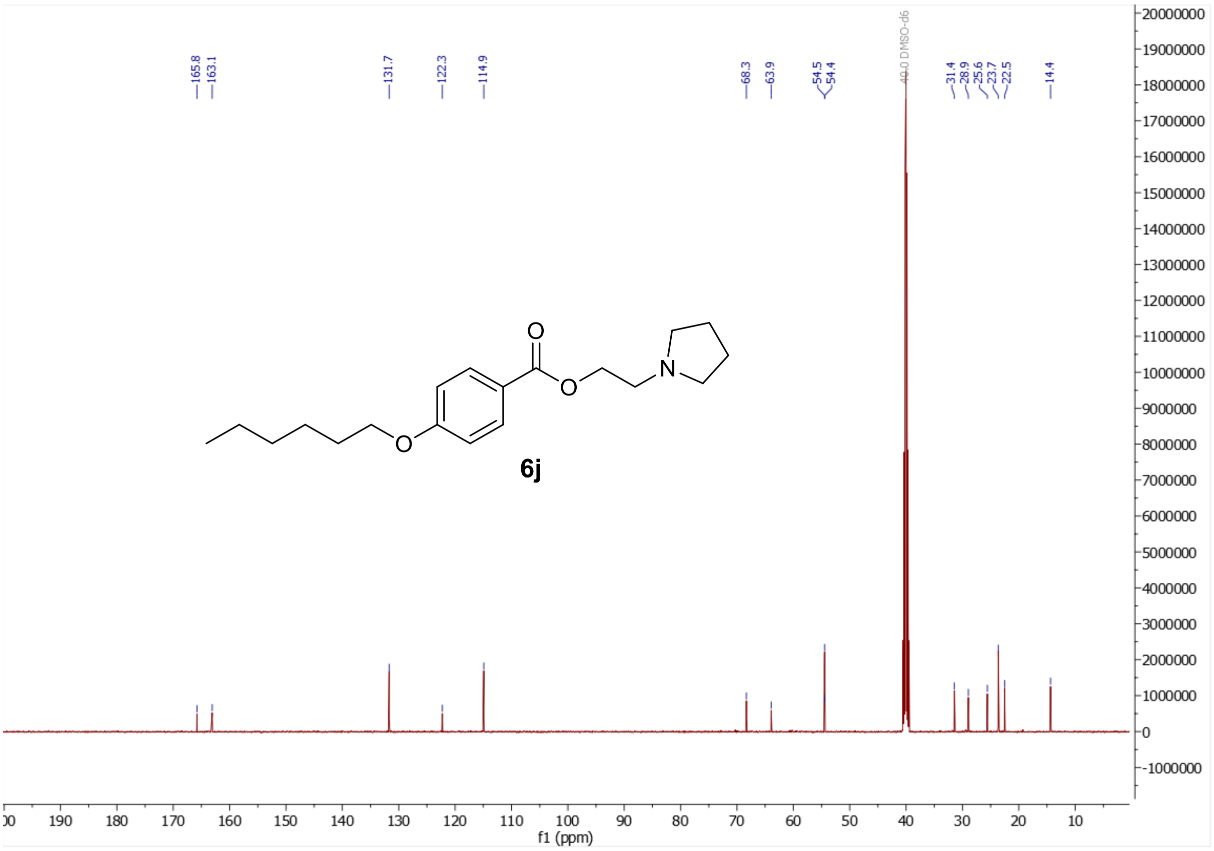

$^1$H and $^{13}$C NMR of 2-(2,3-dihydro-1H-isoindol-2-yl)ethyl 4-(hexyloxy)benzoate (**6k**)

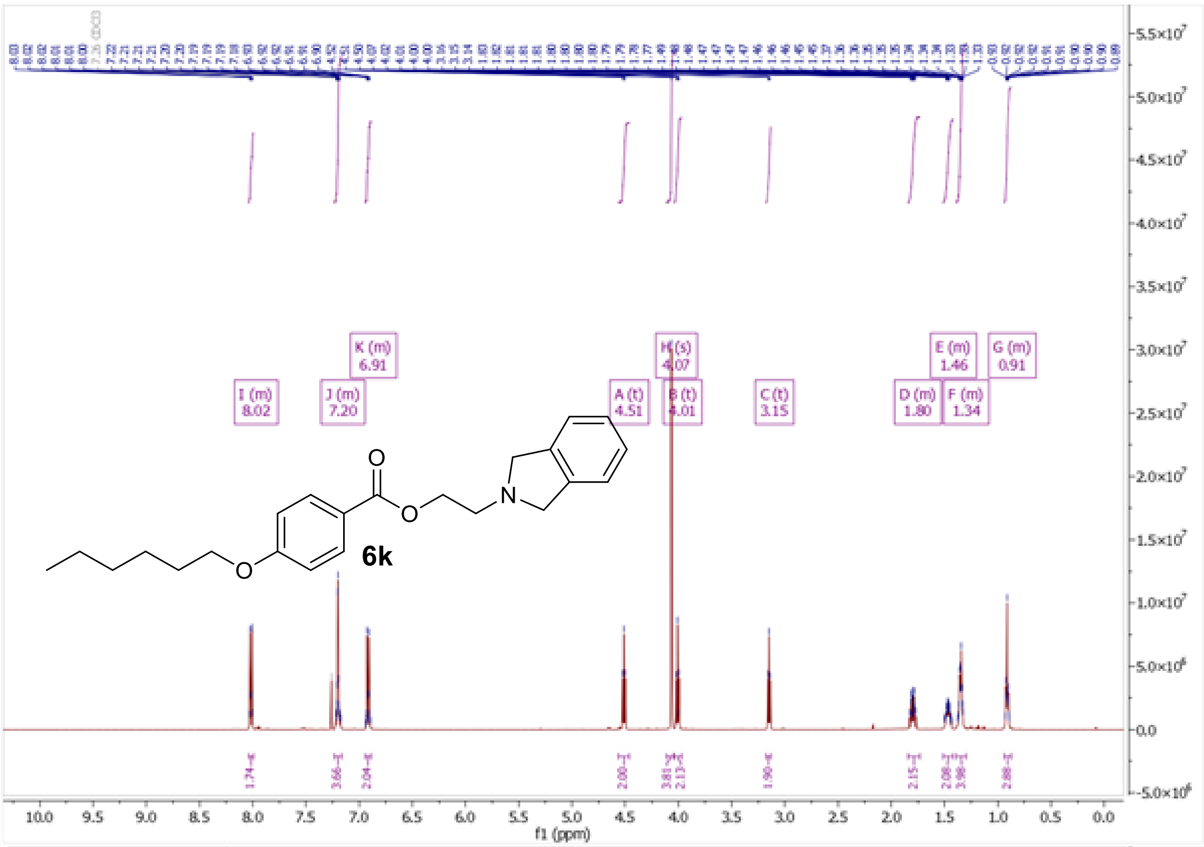

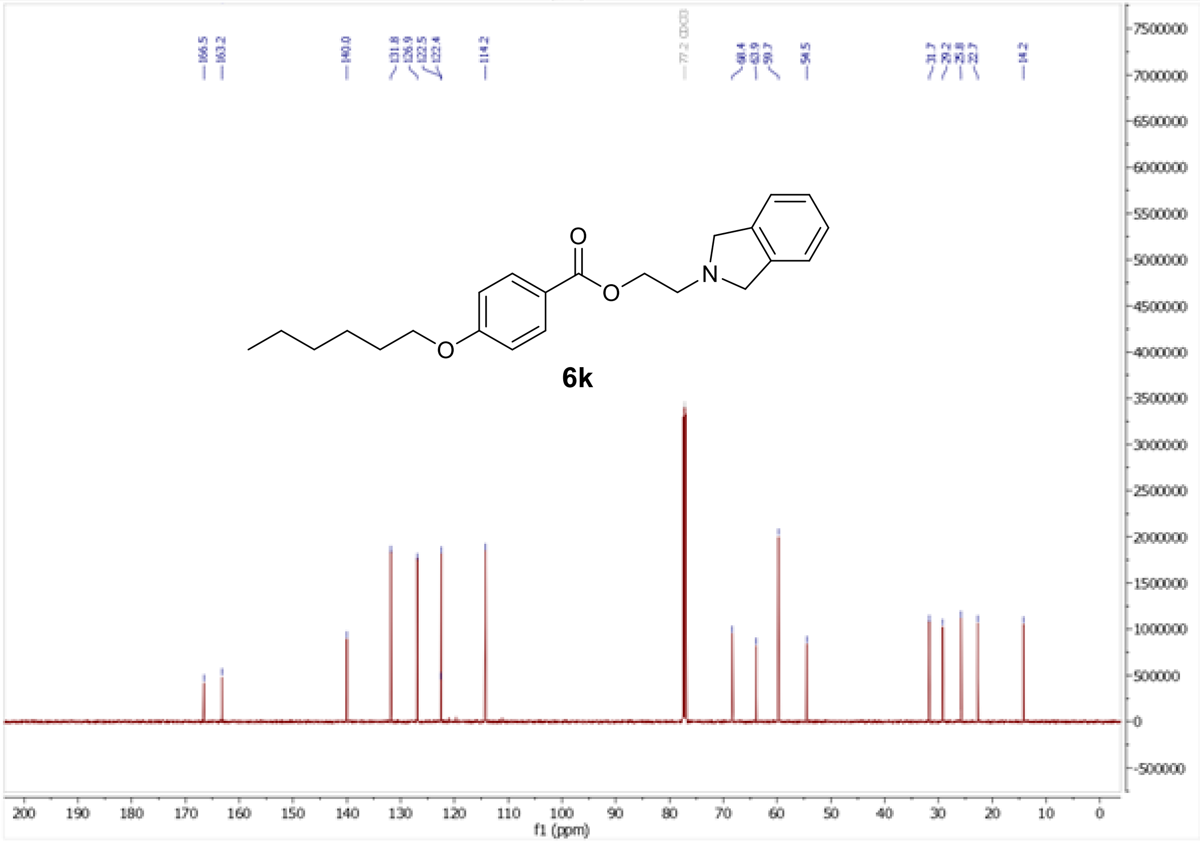

$^{1}H$ and $^{13}C$ NMR spectra of 2-(1,2,3,4-tetrahydroquinolin-1-yl)ethyl 4-(hexyloxy)benzoate (**6l**)

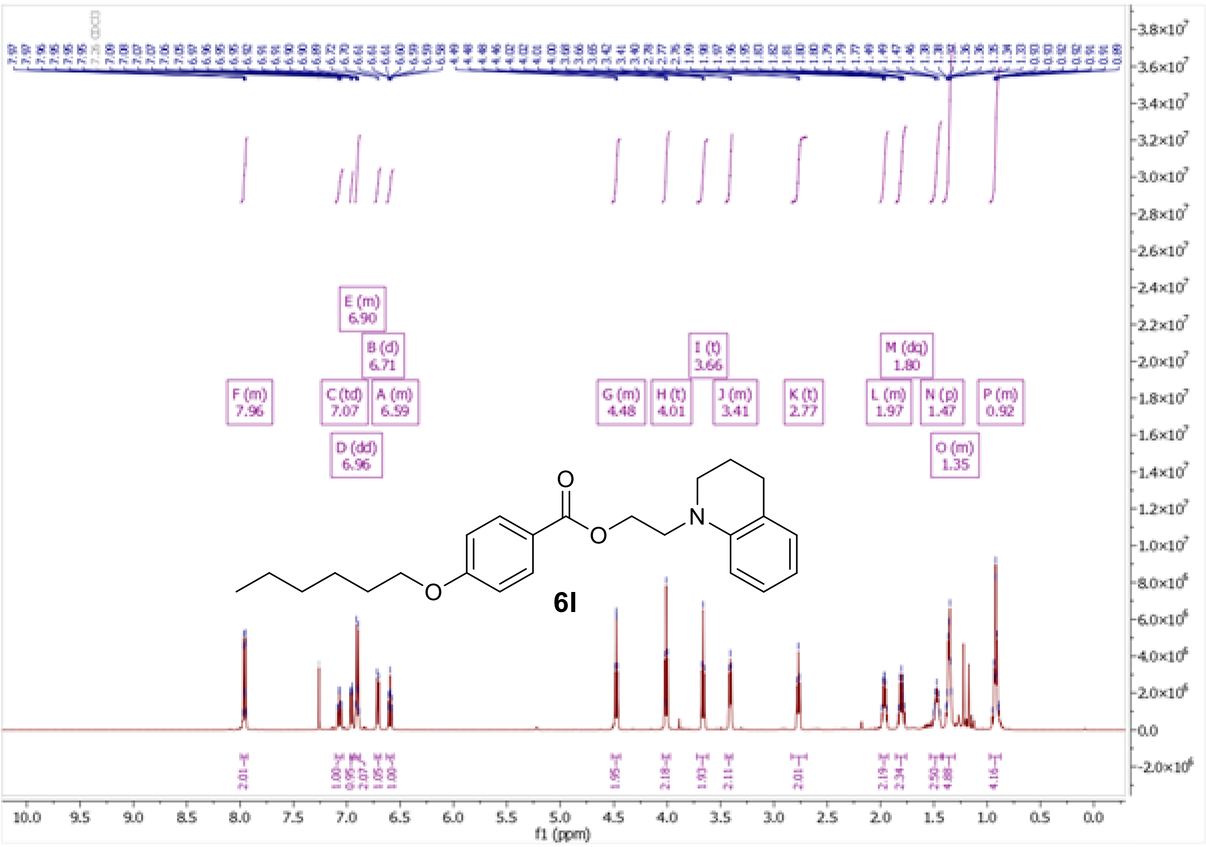

F (m)
7.96
E (m)
6.90
B (d)
6.71
C (td)
7.07
A (m)
6.59
D (dd)
6.96
G (m)
4.48
H (t)
4.01
I (t)
3.66
J (m)
3.41
K (t)
2.77
L (m)
1.97
M (dq)
1.80
N (p)
1.47
O (m)
1.35
P (m)
0.92
6l
f1 (ppm)

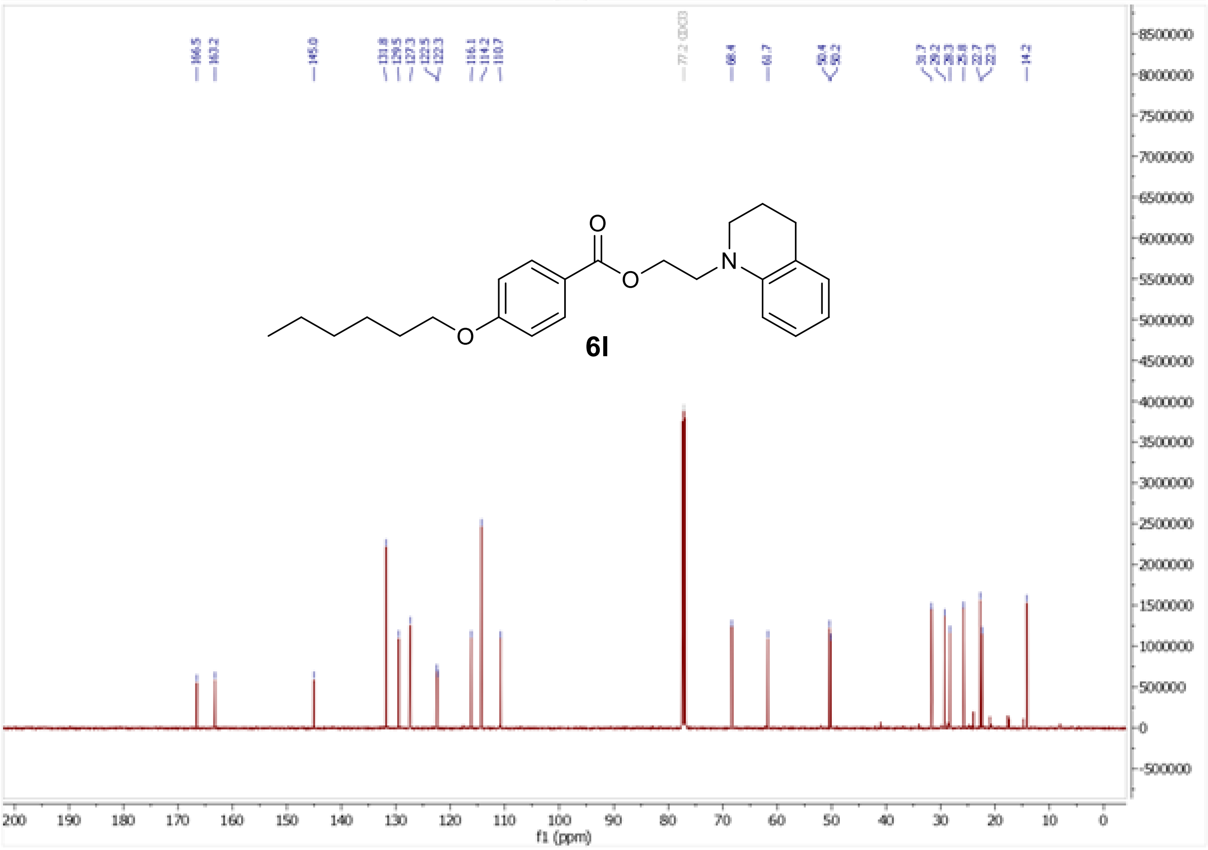

166.5
163.2
145.0
131.8
129.5
127.3
122.5
122.3
116.1
114.2
110.7
77.2 CDCl3
68.4
61.7
50.4
50.2
31.7
29.2
28.3
25.8
22.7
22.3
14.2
6l
f1 (ppm)

<sup></sup>

[1]H and [13]C NMR spectra of (1-methylpiperidin-2-yl)methyl 4-(hexyloxy)benzoate (**6m**)

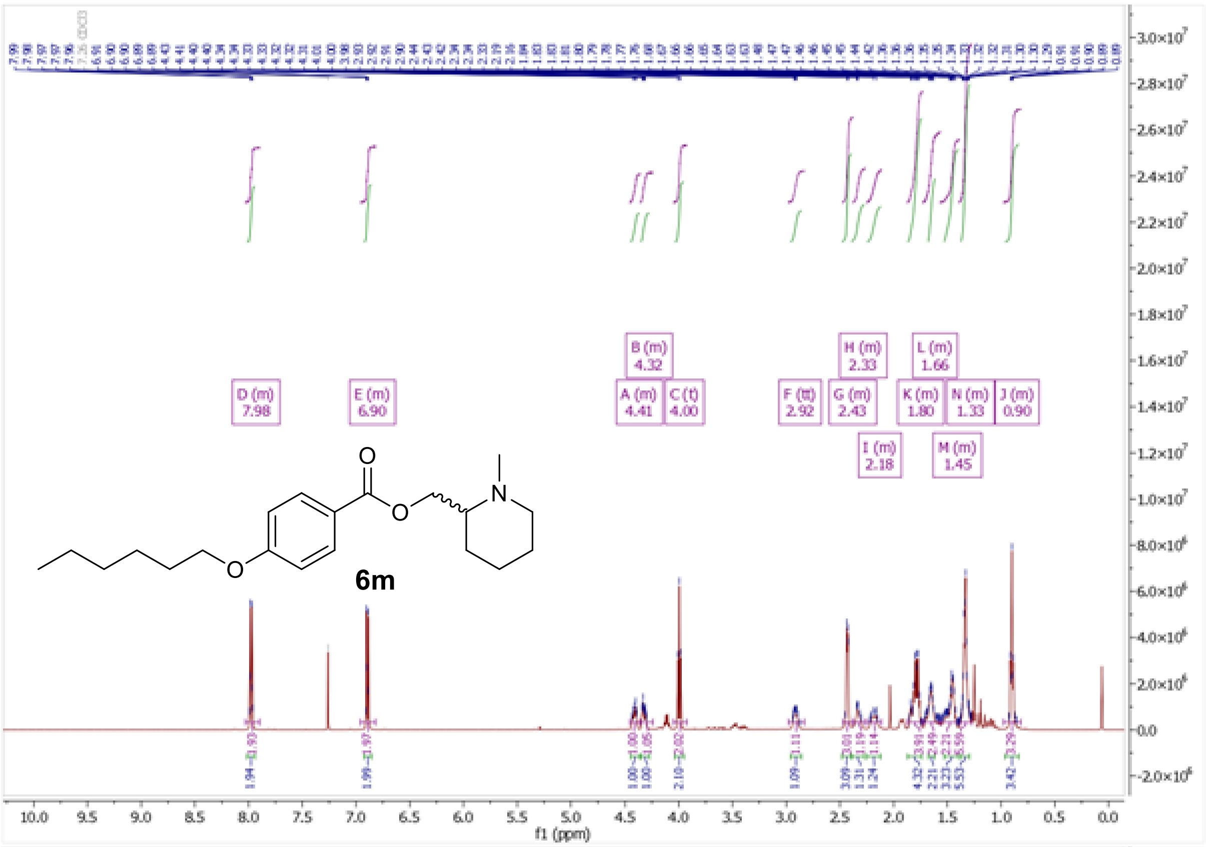

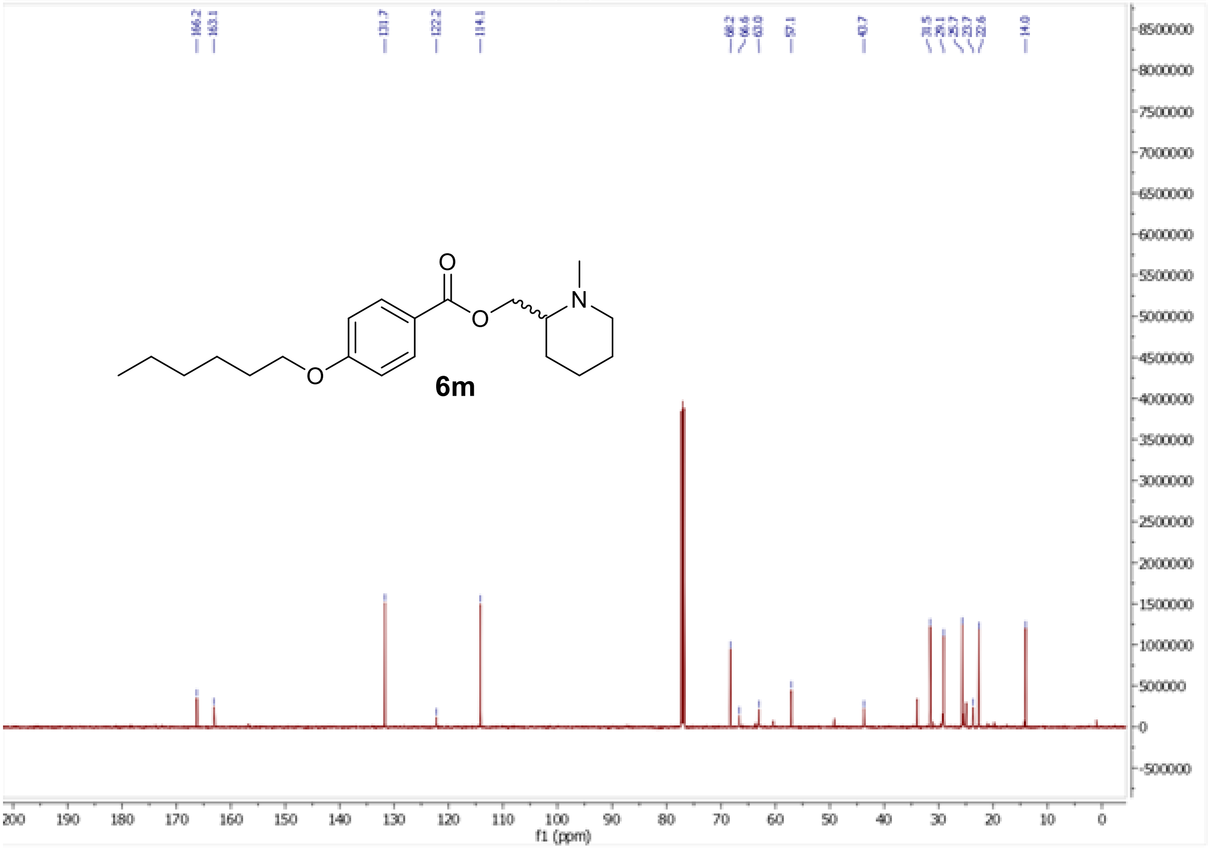

[1]H and [13]C NMR spectra of [(2*S*)-1-methylpiperidin-2-yl]methyl 4-(hexyloxy)benzoate (**6n**)

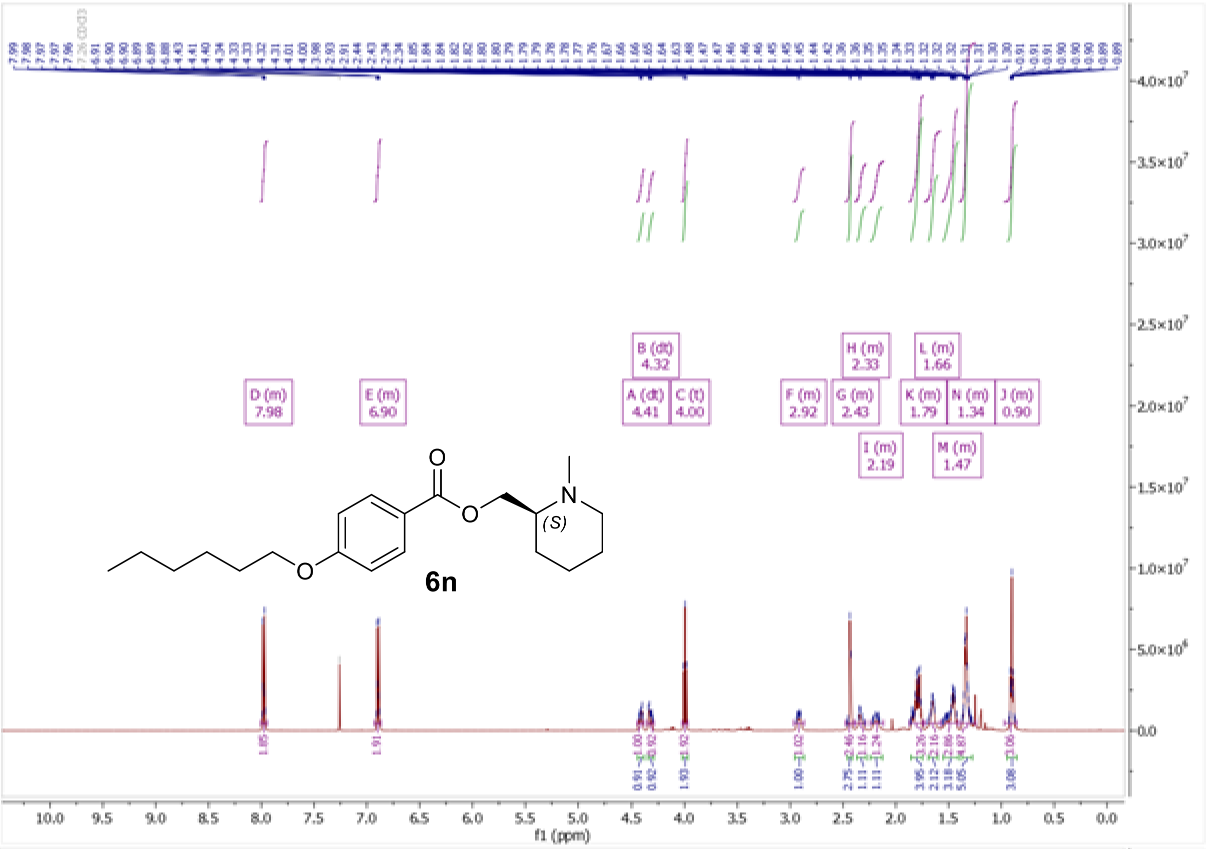

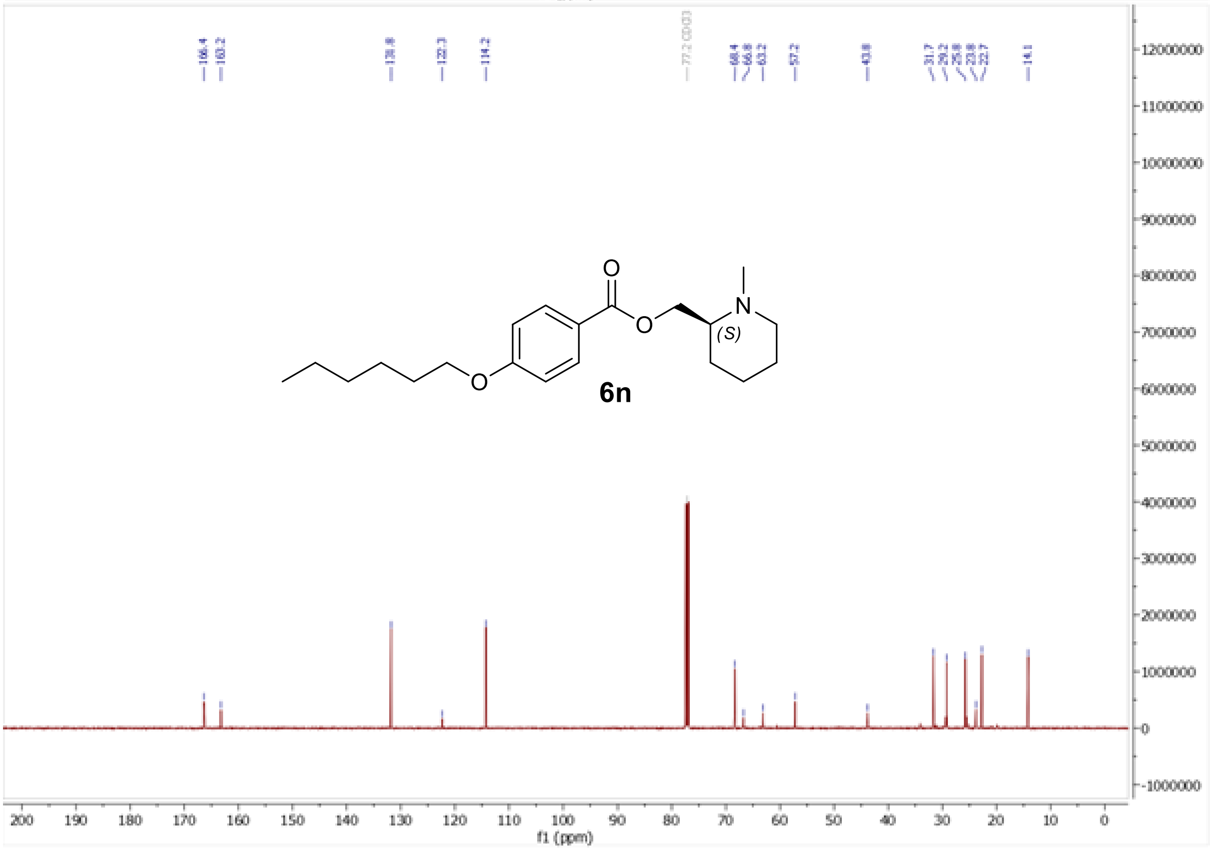

[1]H and [13]C NMR spectra of [(3*S*)-1-methylpiperidin-3-yl]methyl 4-(hexyloxy)benzoate (**6o**)

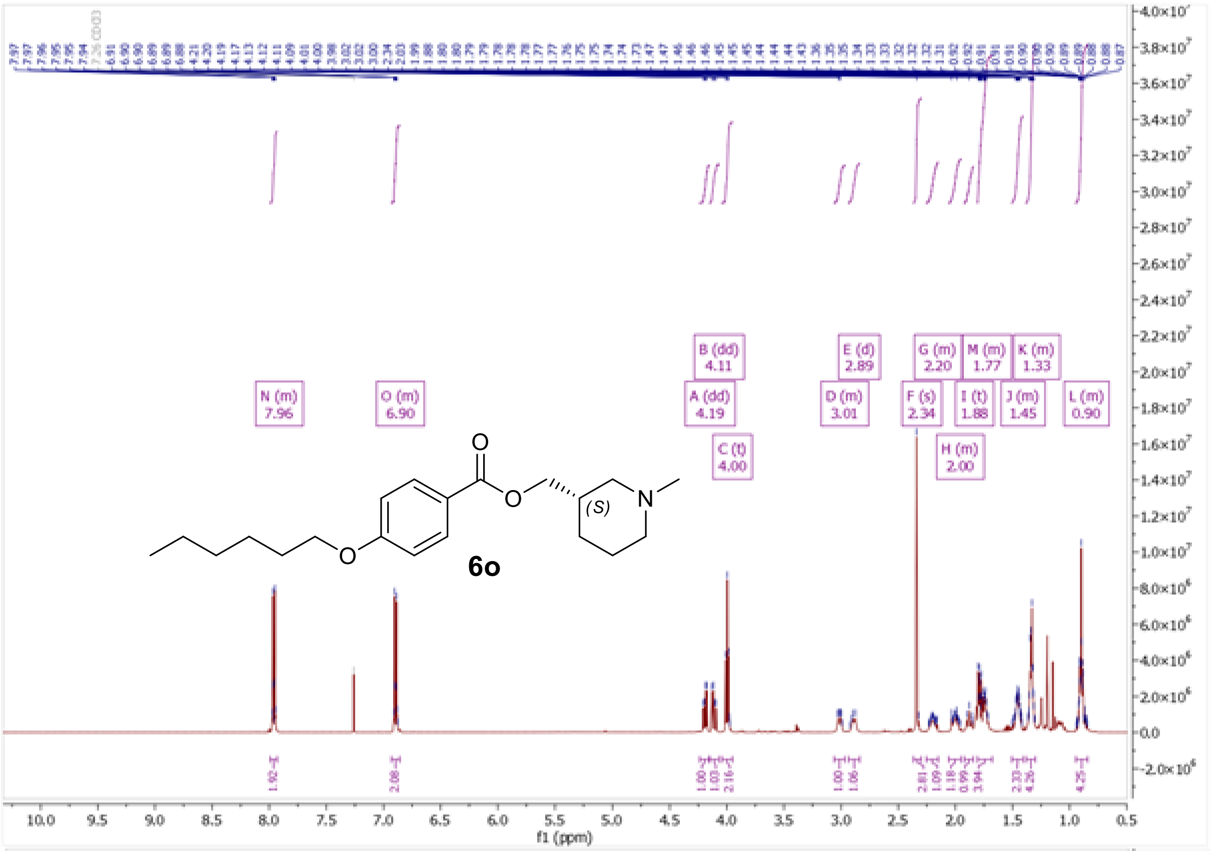

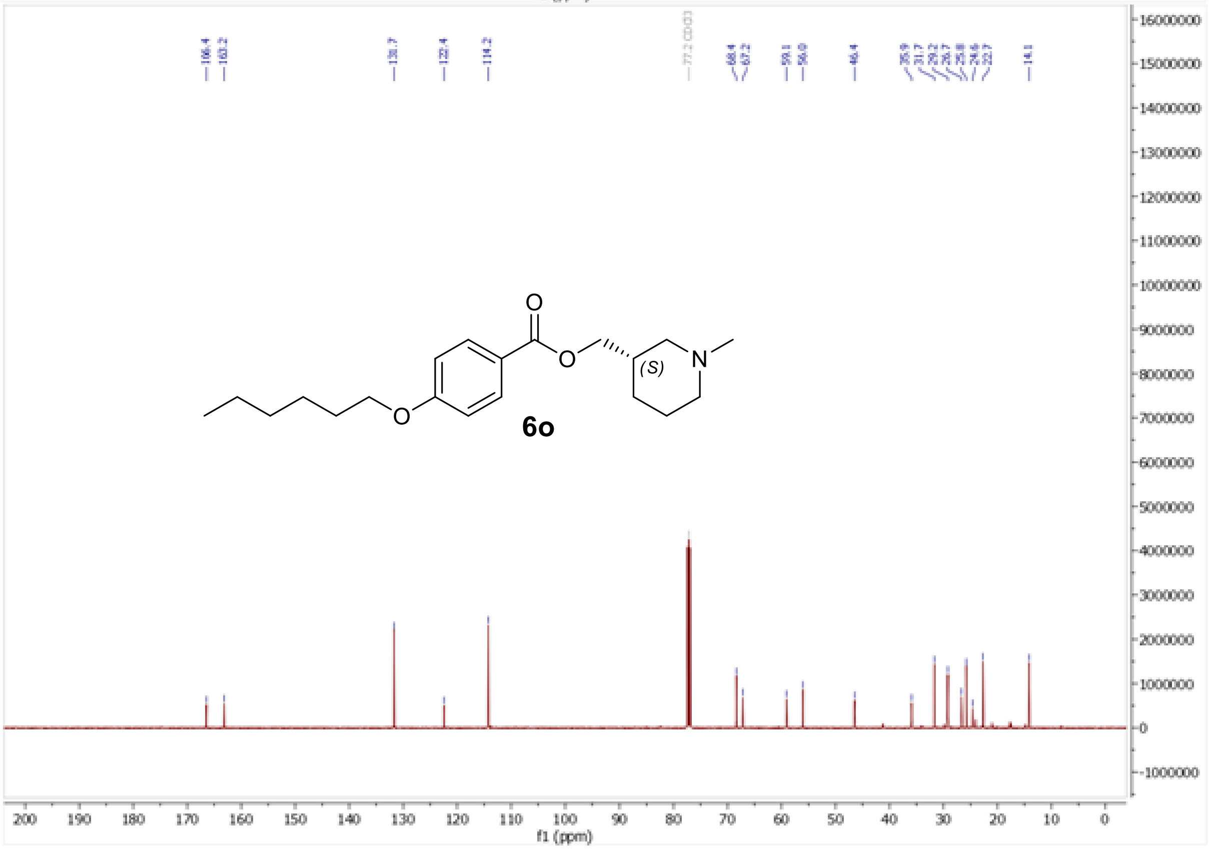

$^{1}H$ and $^{13}C$ NMR spectra of (1-methylpiperidin-3-yl)methyl 4-(hexyloxy)benzoate (**6p**)

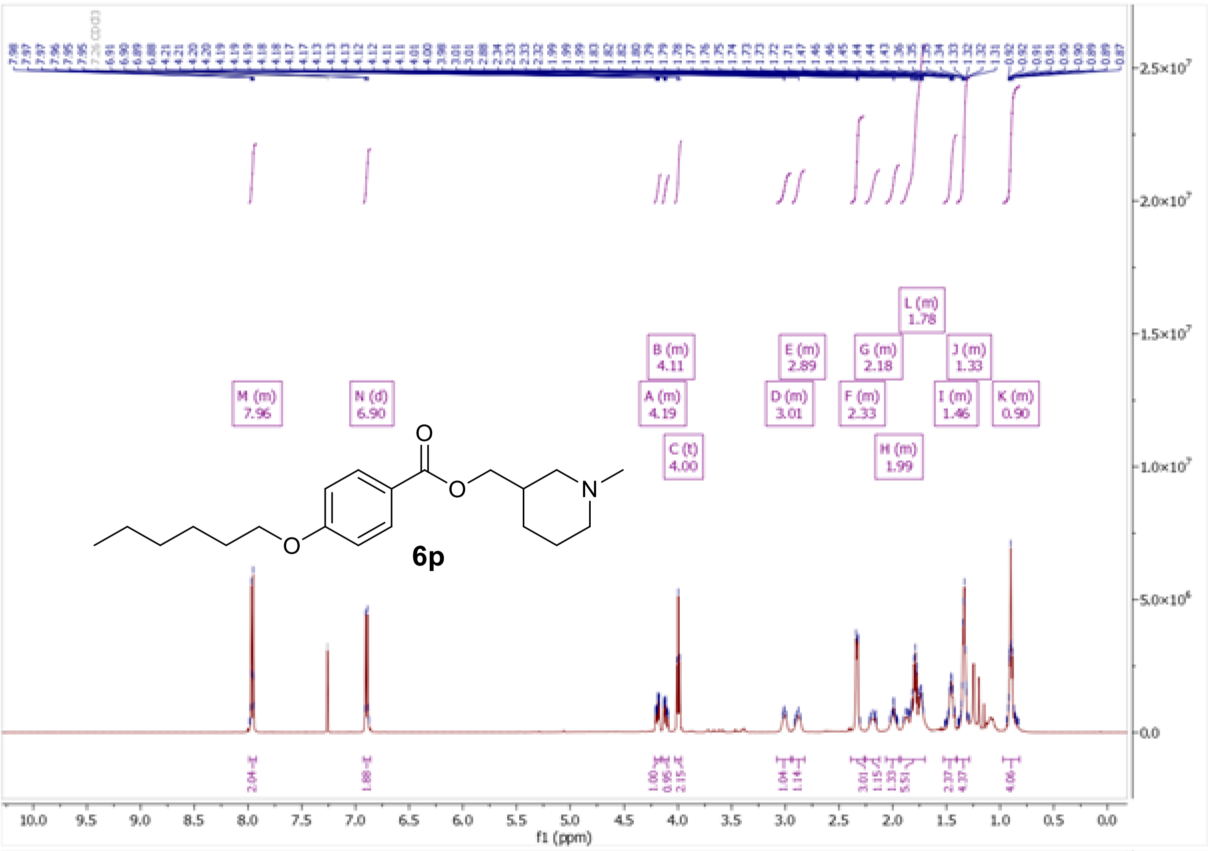
M (m)
7.96
N (d)
6.90
B (m)
4.11
A (m)
4.19
C (t)
4.00
E (m)
2.89
D (m)
3.01
G (m)
2.18
F (m)
2.33
H (m)
1.99
L (m)
1.78
J (m)
1.33
I (m)
1.46
K (m)
0.90
6p
f1 (ppm)

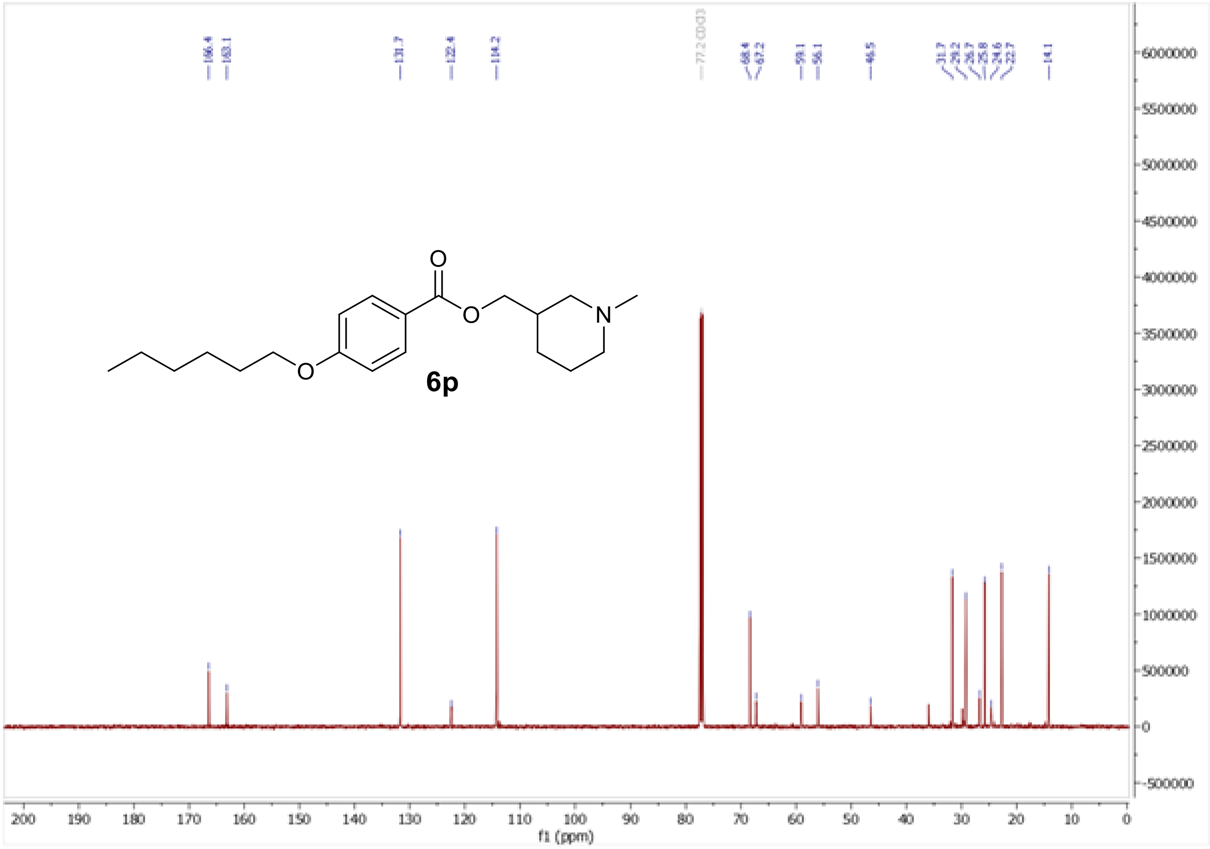
166.4
163.1
131.7
122.4
114.2
77.2 CDCl3
68.4
67.2
59.1
56.1
46.5
31.7
29.2
26.7
25.8
24.6
22.7
14.1
6p
f1 (ppm)

$^1H$ and $^{13}C$ NMR spectra of [(3*R*)-1-methylpiperidin-3-yl]methyl 4-(hexyloxy)benzoate (**6r**)

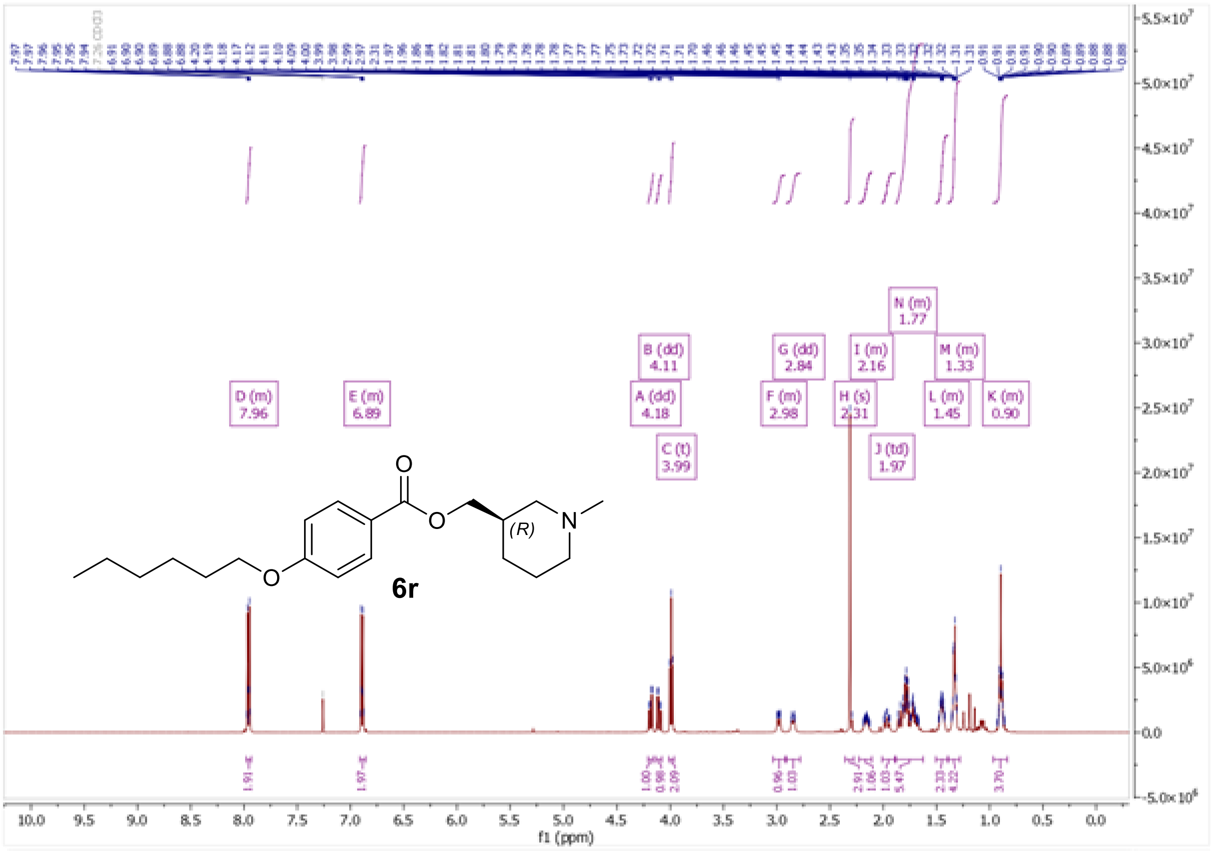

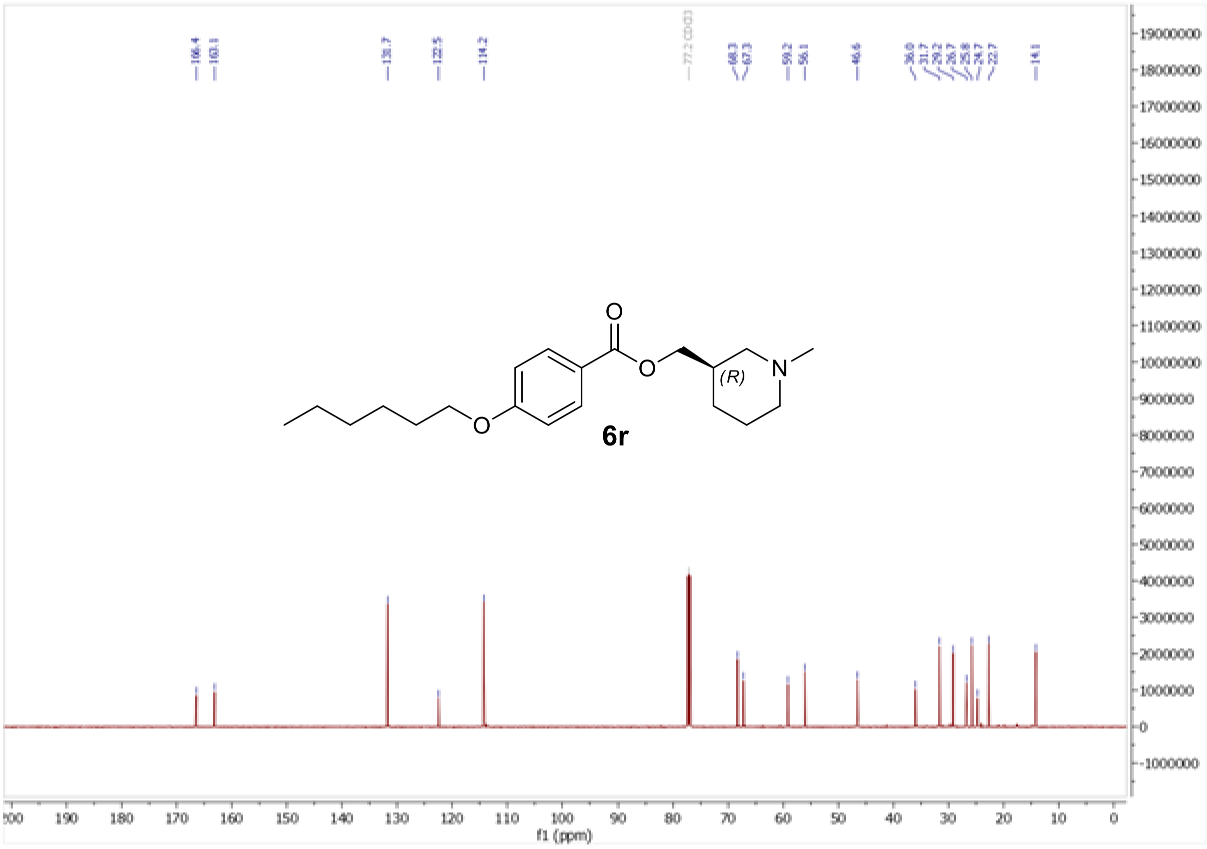

$^{1}H$ and $^{13}C$ NMR spectra of (1-methylpiperidin-4-yl)methyl 4-(hexyloxy)benzoate (**6s**)

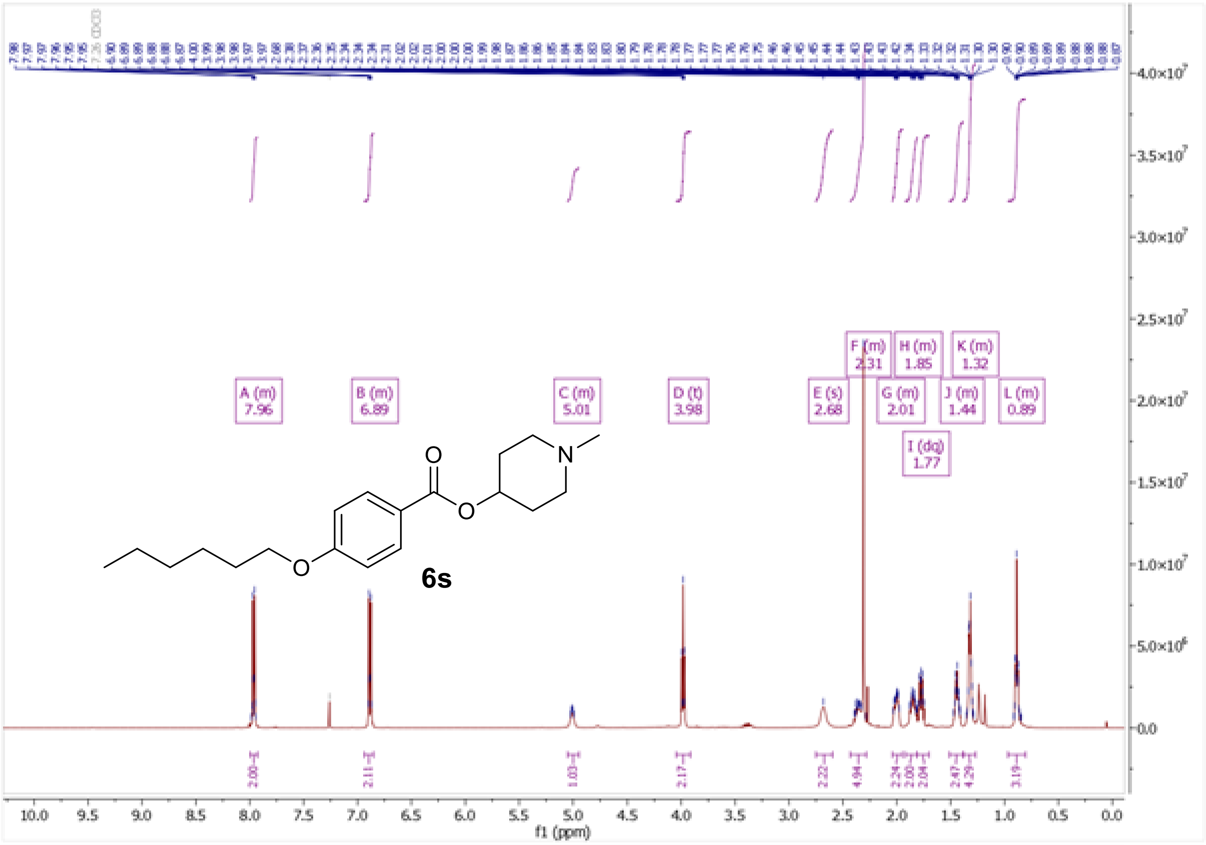

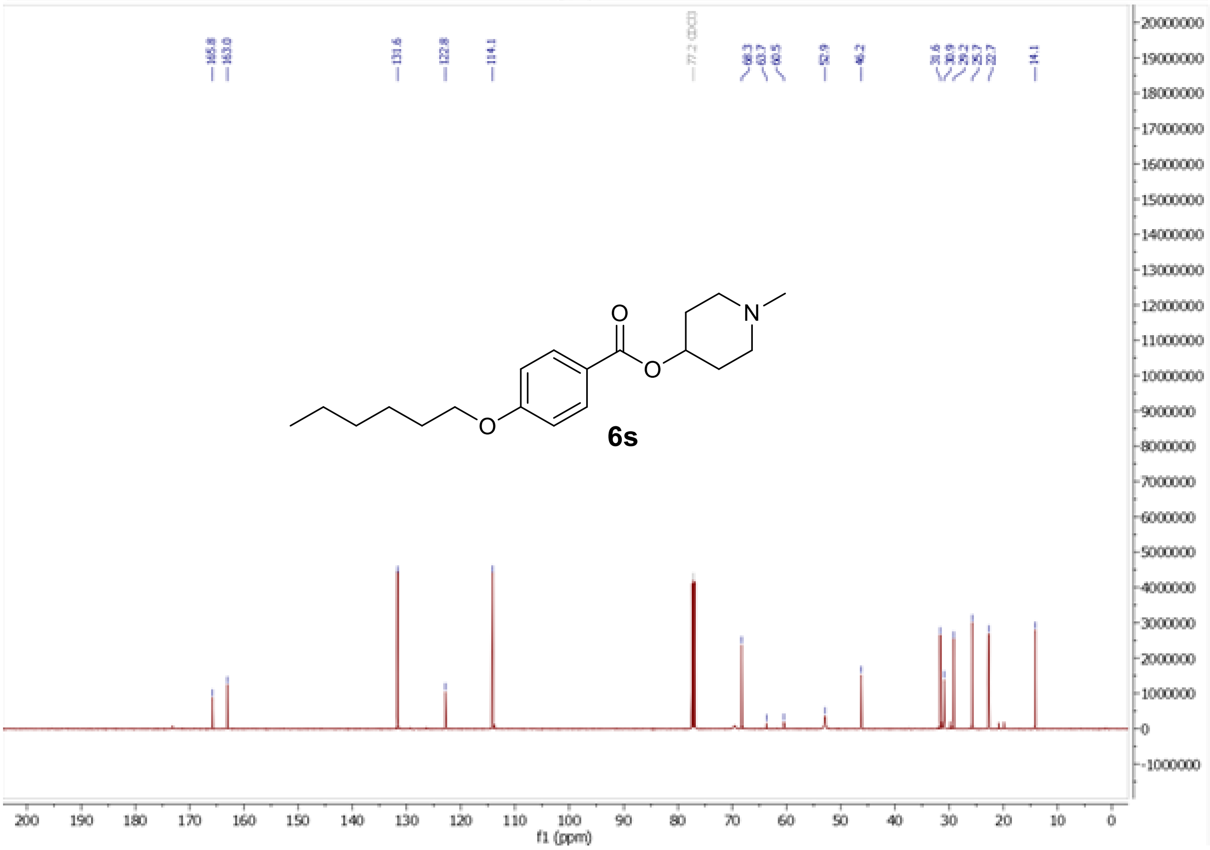

$^{1}H$ and $^{13}C$ NMR spectra of (1-methylpyrrolidin-2-yl)methyl 4-(hexyloxy)benzoate (**6t**)

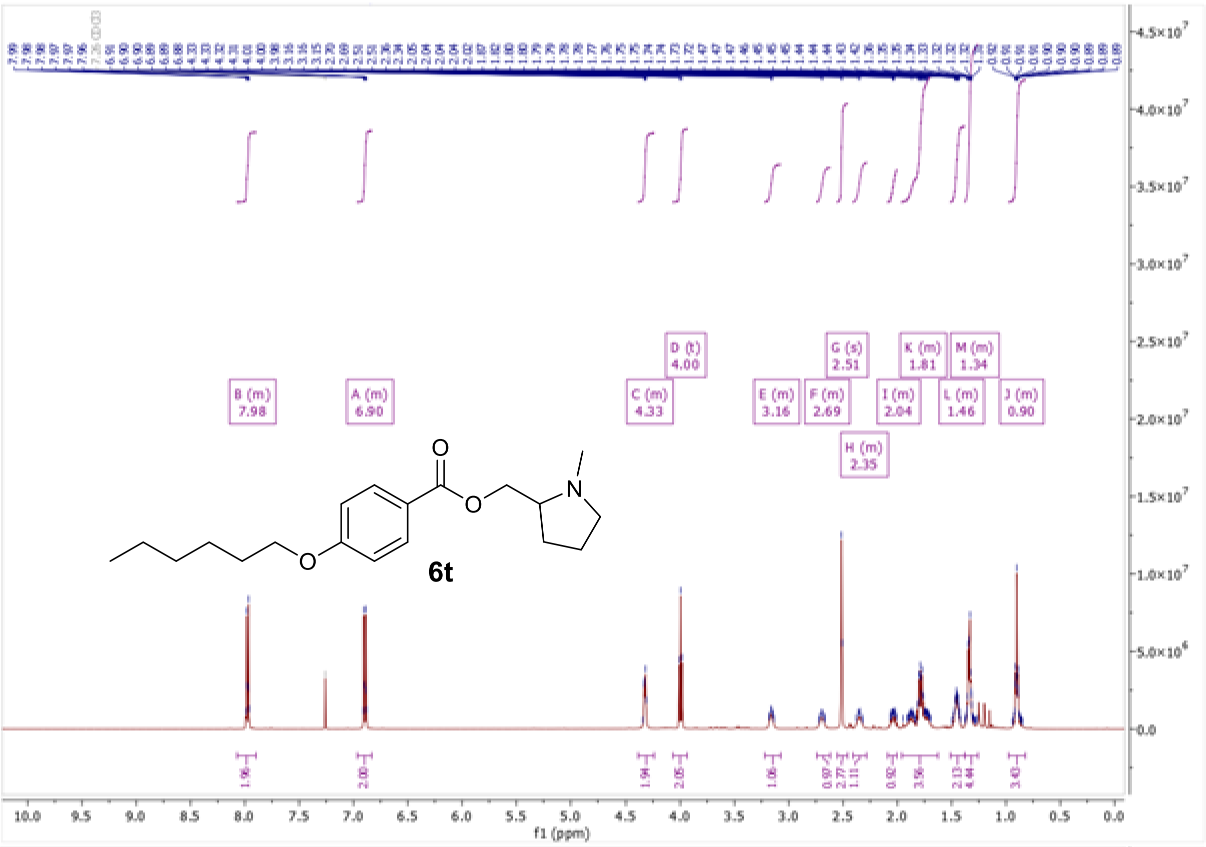

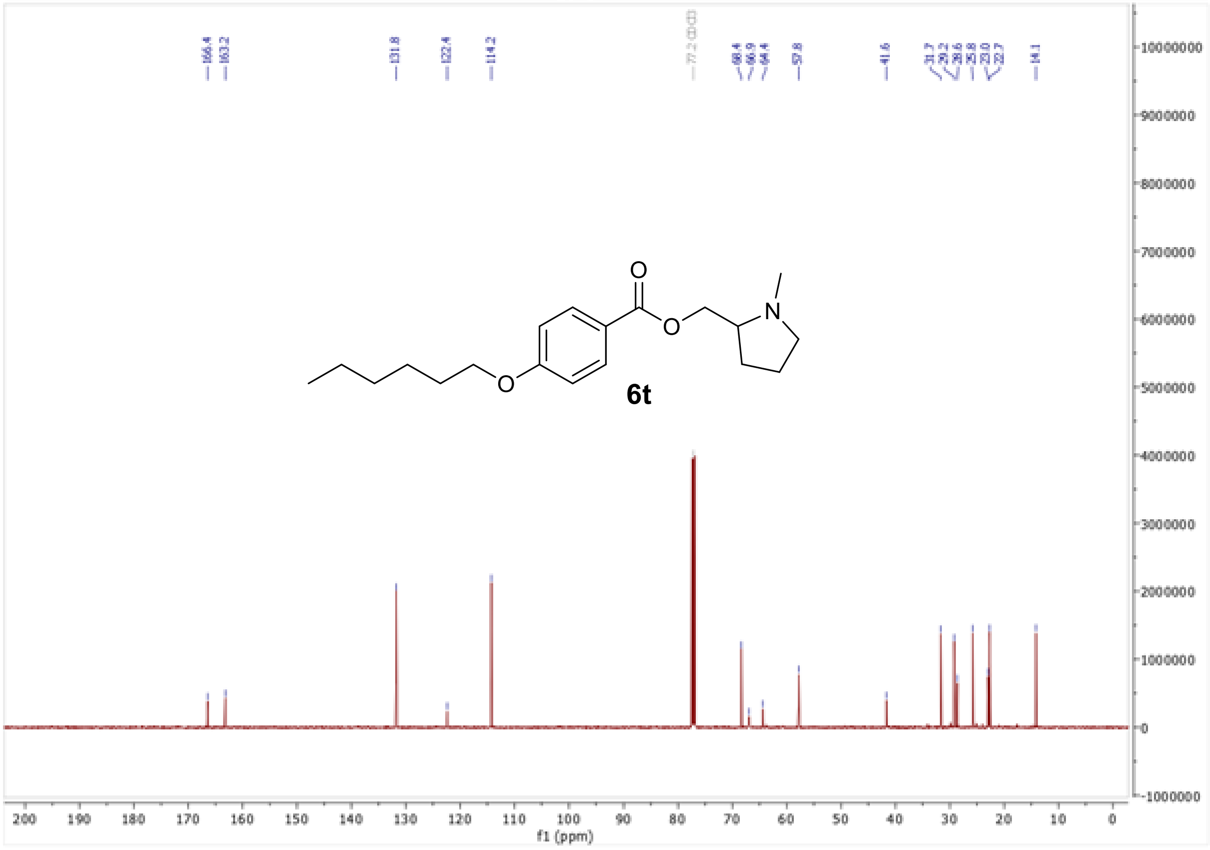

$^{1}H$ and $^{13}C$ NMR spectra of [(2*S*)-1-methylpyrrolidin-2-yl]methyl 4-(hexyloxy)benzoate (**6w**)

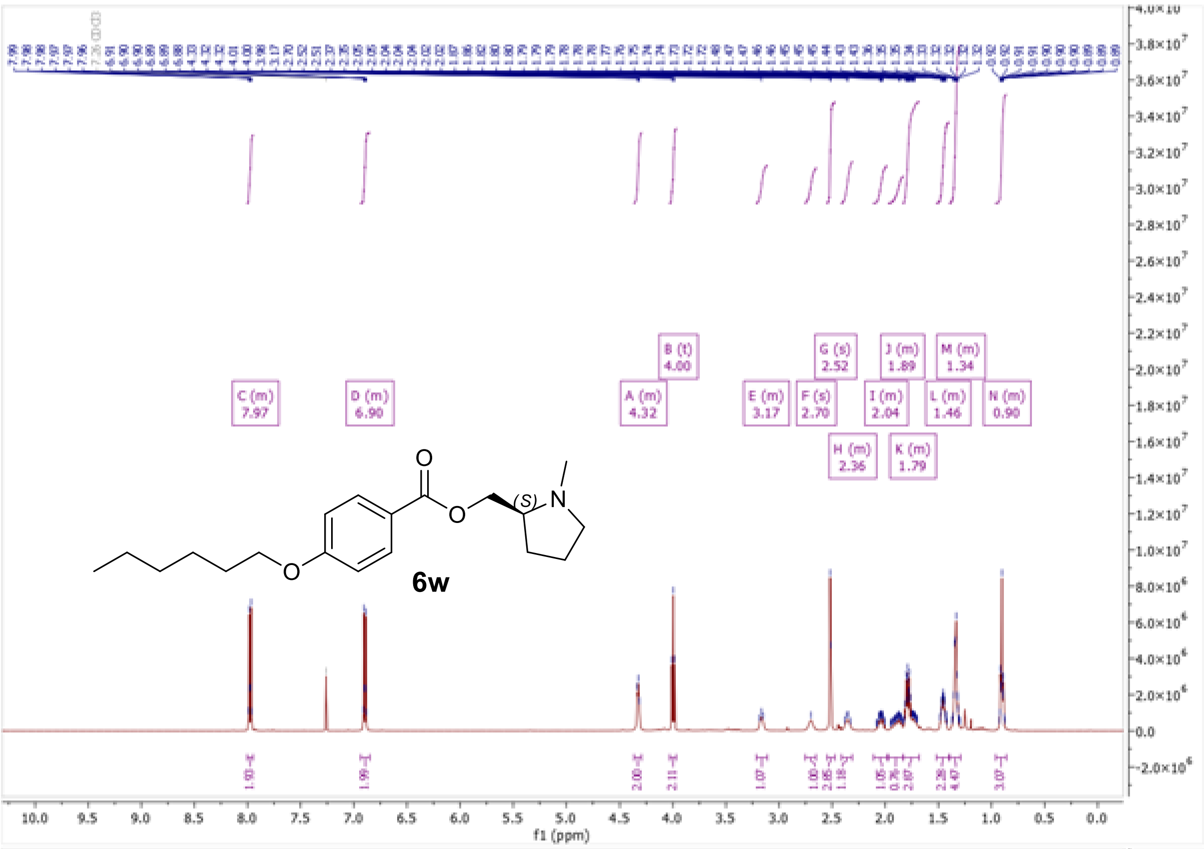

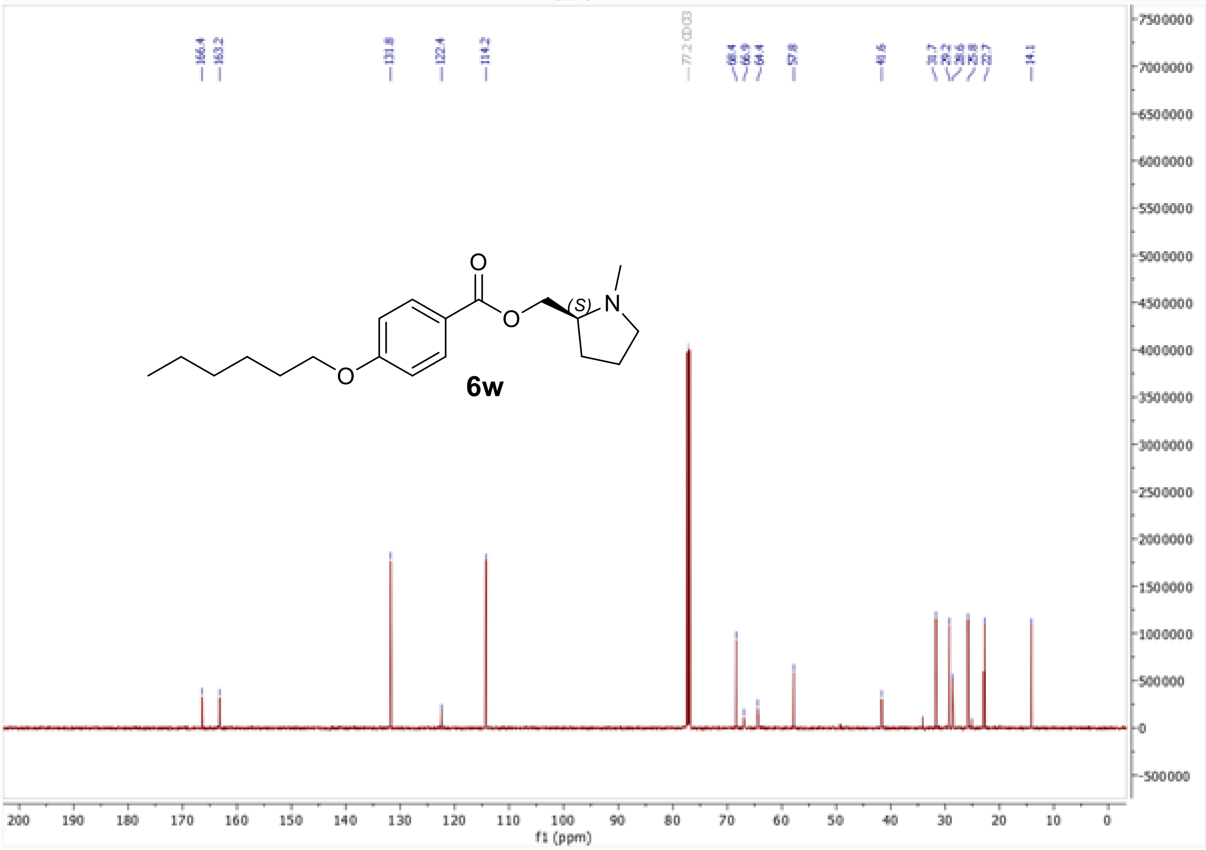

$^1H$ and $^{13}C$ NMR spectra of [(2*R*)-1-methylpyrrolidin-2-yl]methyl 4-(hexyloxy)benzoate (**6x**)

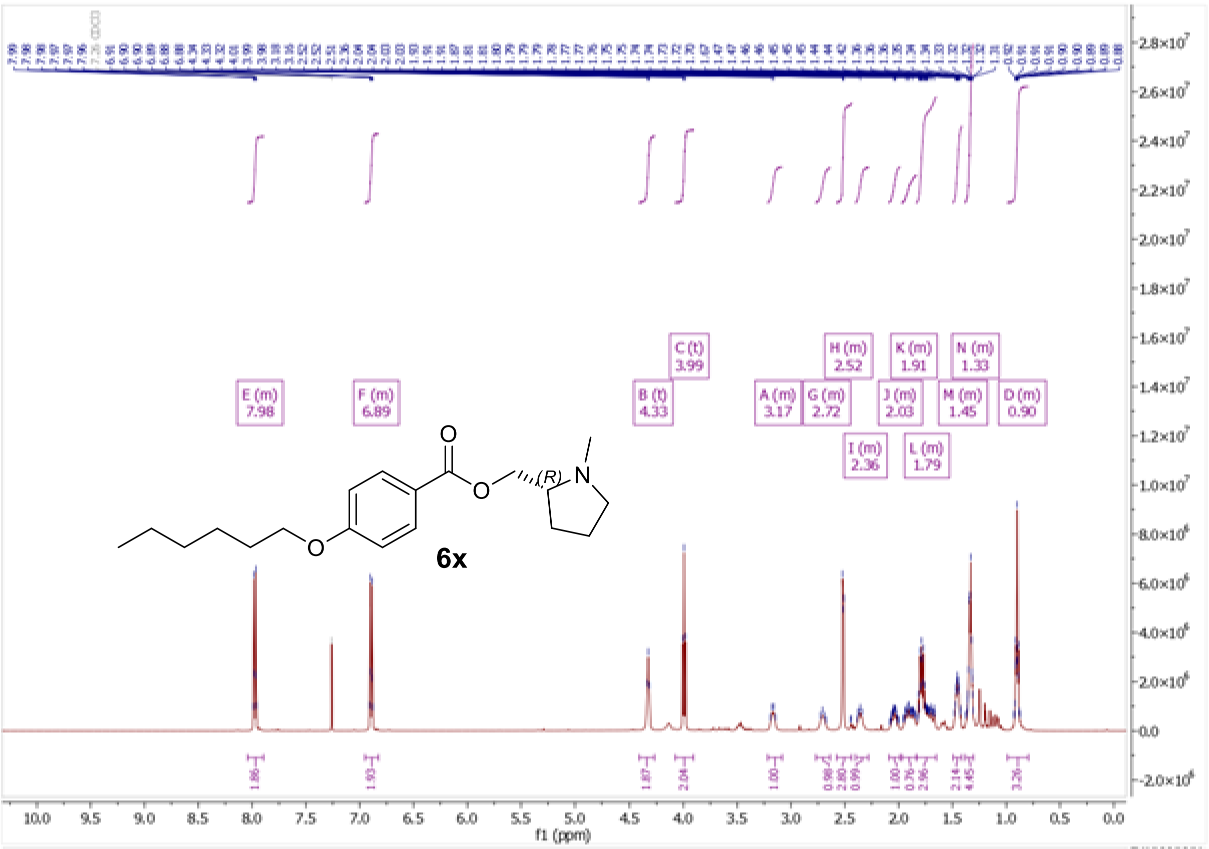
E (m)
7.98
F (m)
6.89
B (t)
4.33
C (t)
3.99
A (m)
3.17
G (m)
2.72
H (m)
2.52
I (m)
2.36
J (m)
2.03
K (m)
1.91
L (m)
1.79
M (m)
1.45
N (m)
1.33
D (m)
0.90
(R)
6x
f1 (ppm)

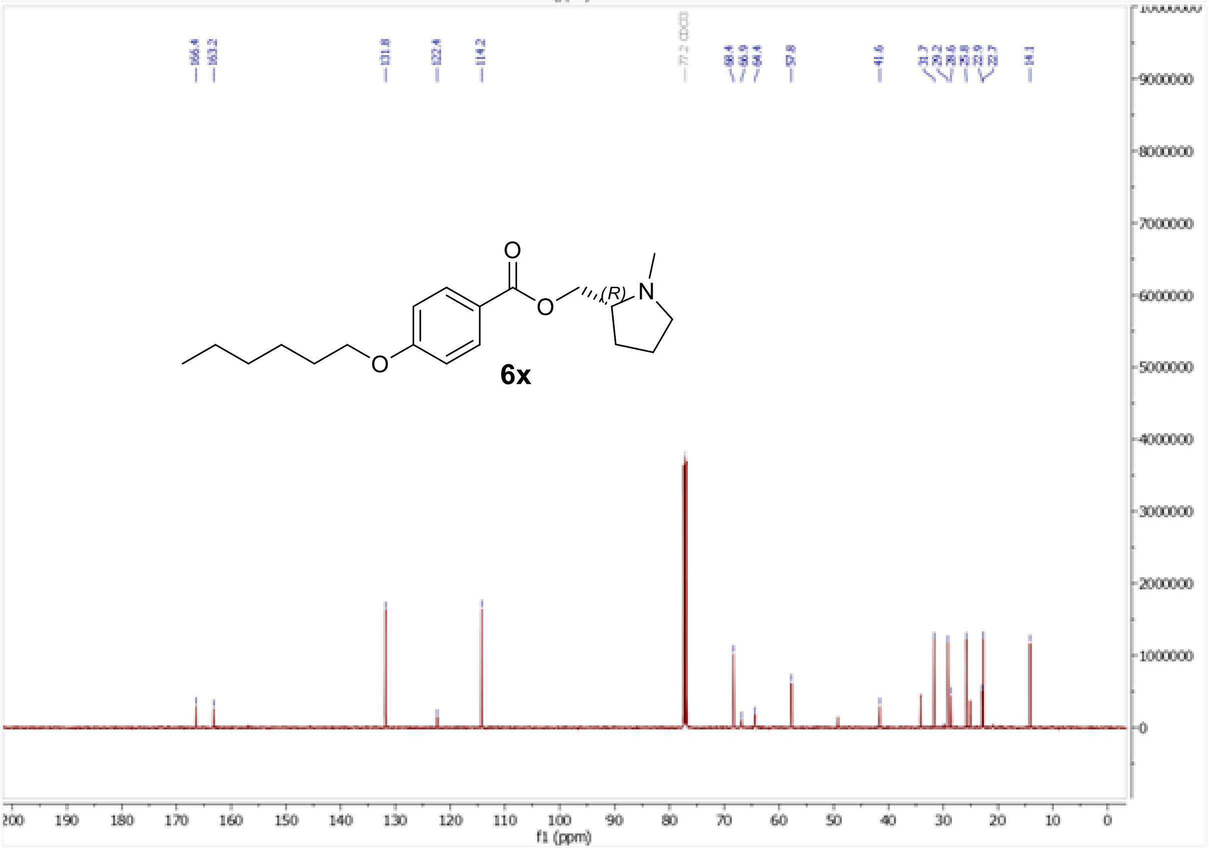
166.4
163.2
131.8
122.4
114.2
77.2 CDCl3
68.4
66.9
64.4
57.8
41.6
(R)
6x
f1 (ppm)

$^{1}H$ and $^{13}C$ NMR spectra of (1-methylpyrrolidin-3-yl)methyl 4-(hexyloxy)benzoate (**6y**)

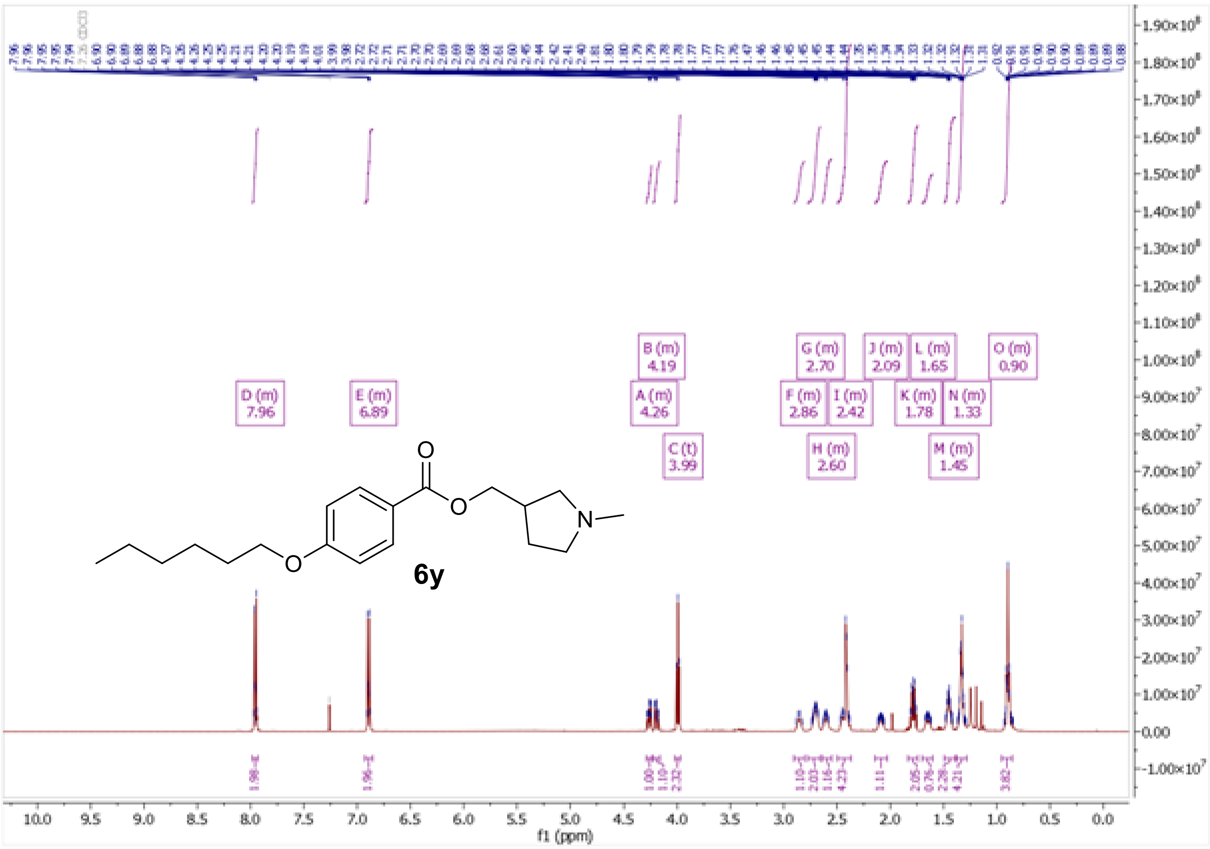

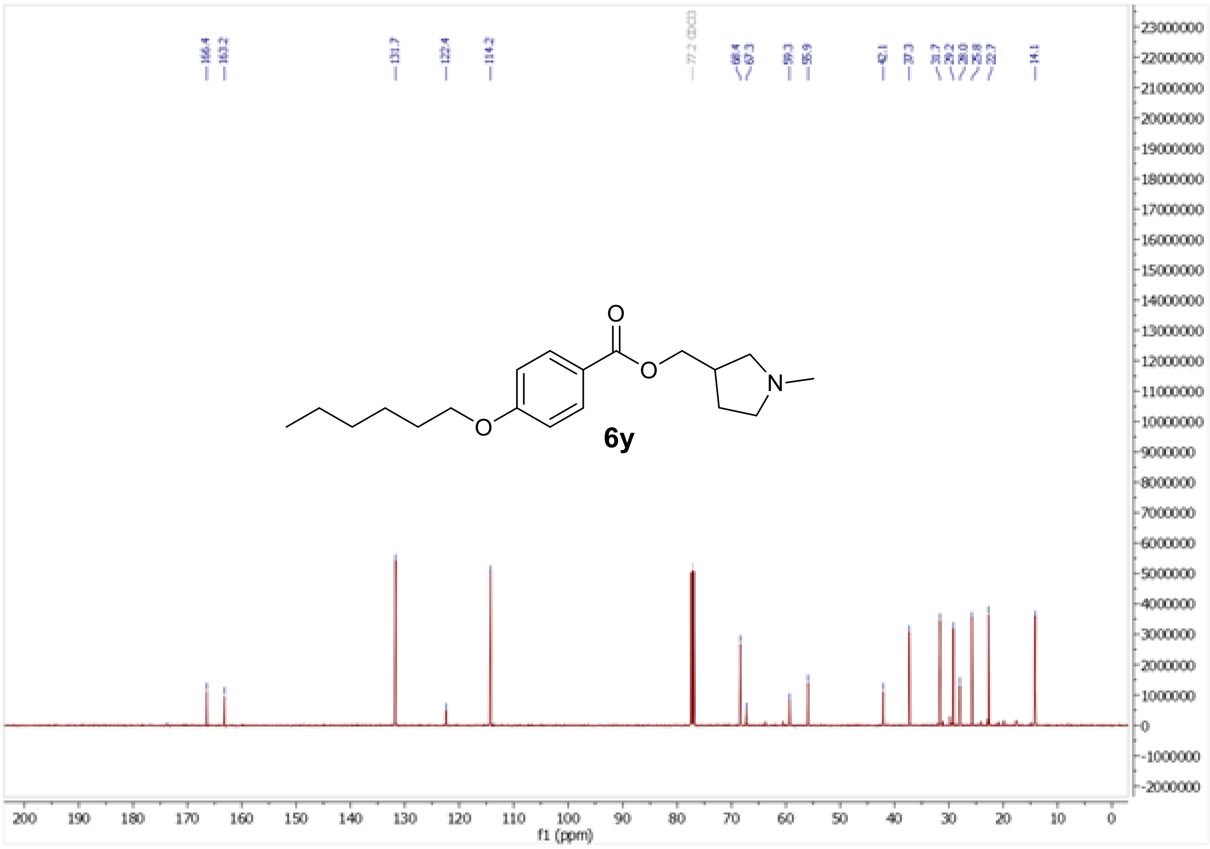

$^{1}H$ and $^{13}C$ NMR spectra of [(3*S*)-1-methylpyrrolidin-3-yl]methyl 4-(hexyloxy)benzoate (**6z**)

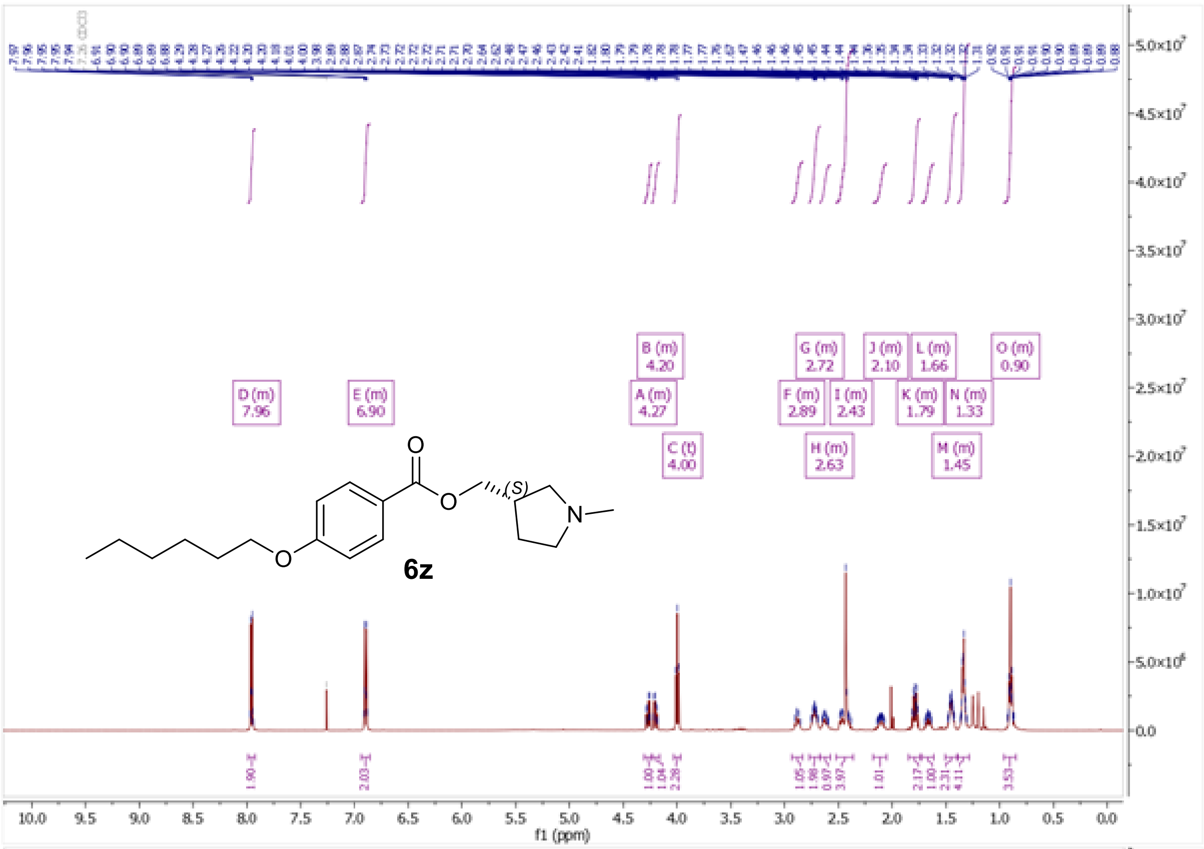

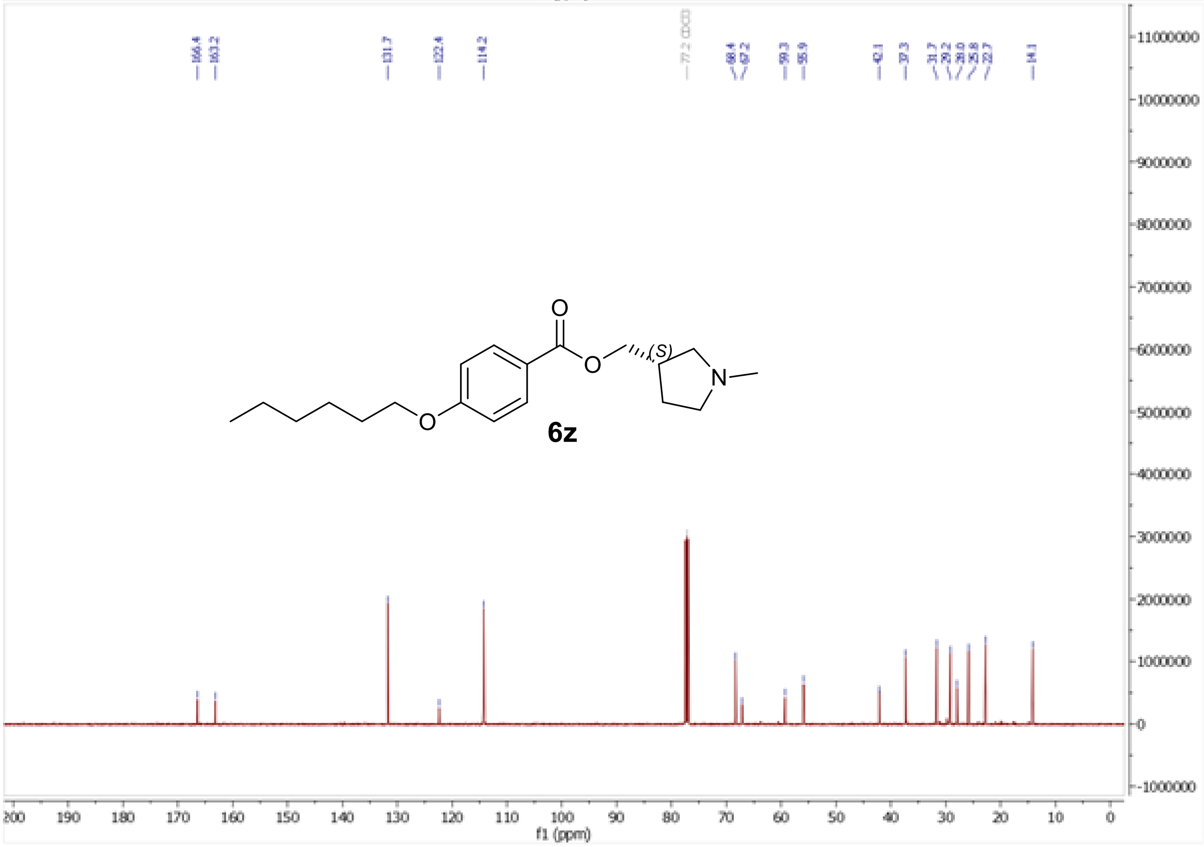

$^{1}H$ and $^{13}C$ NMR spectra of (2*R*)-1-{2-[4-(hexyloxy)benzoyloxy]ethyl}-1,2-dimethylpiperidin-1-ium iodide (**7e**)

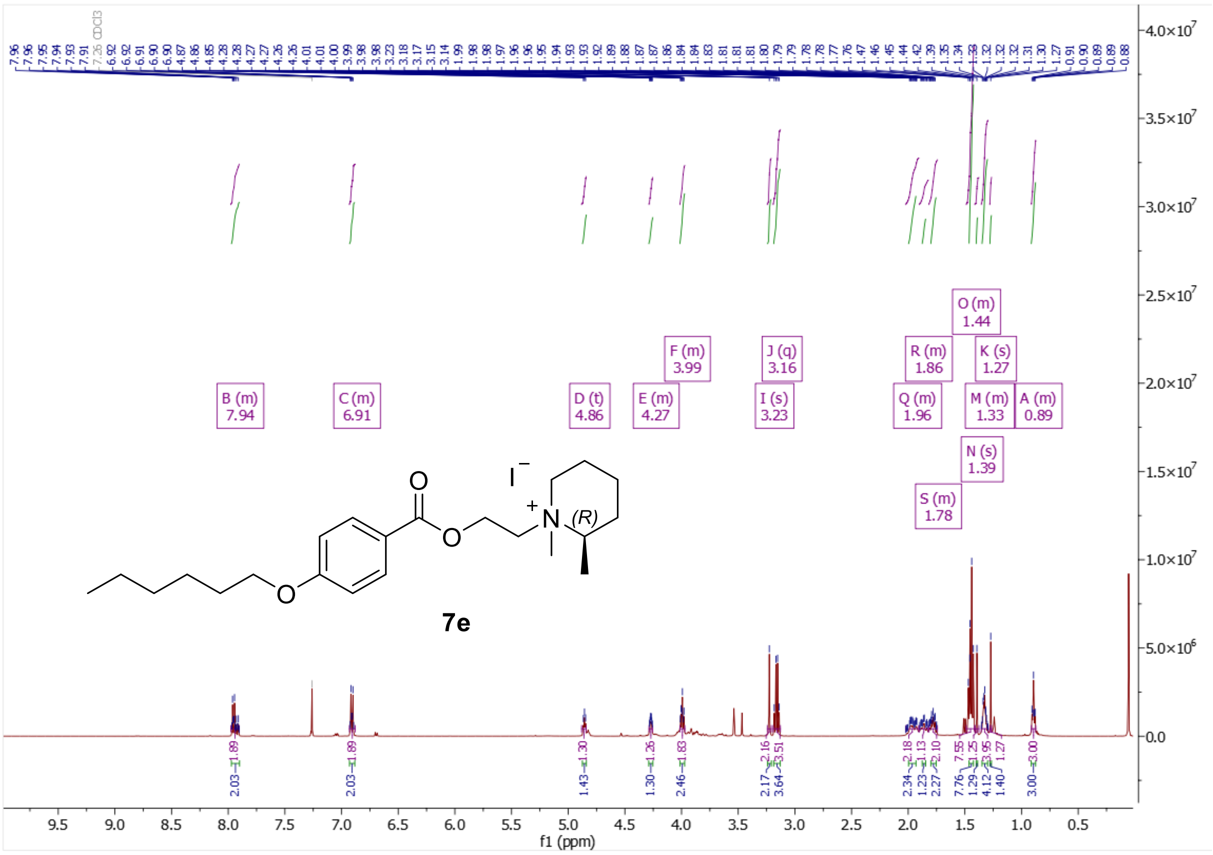

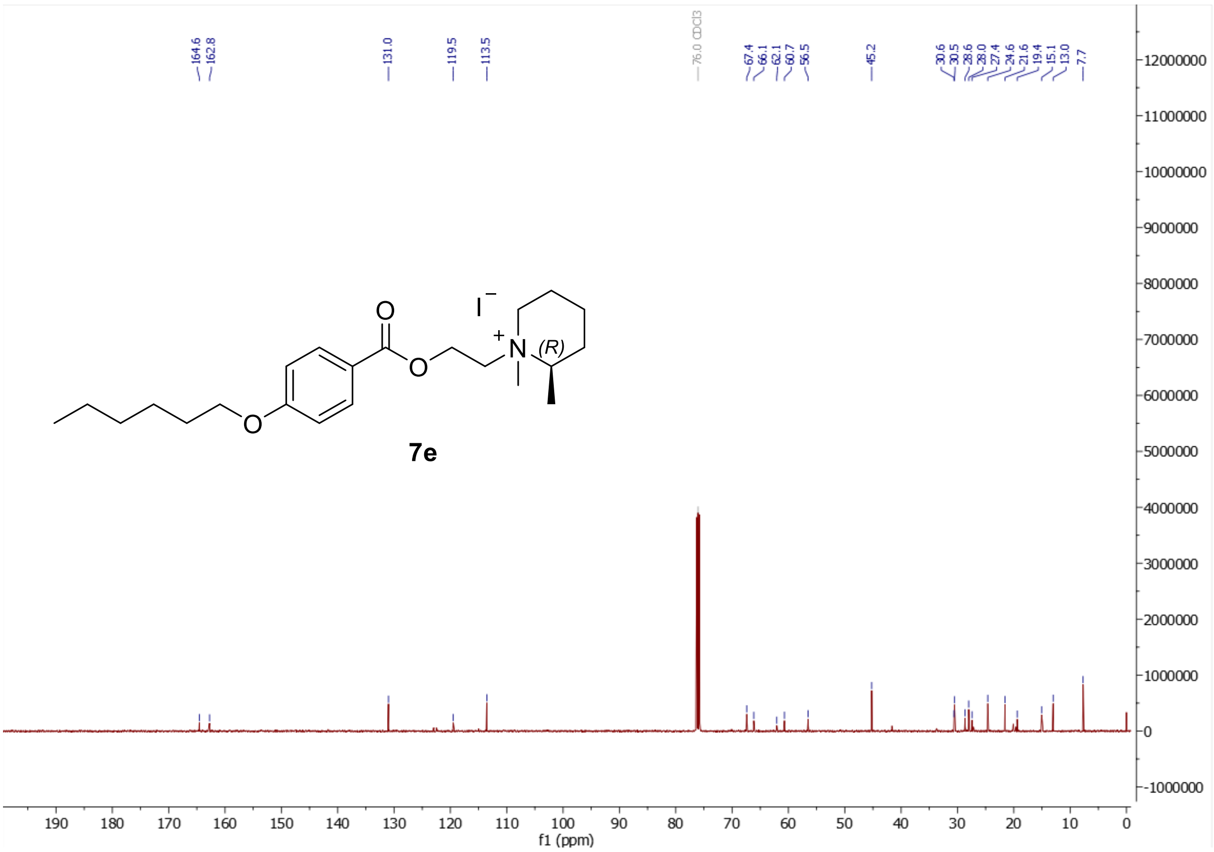

[1]H and [13]C NMR spectra of (2*S*)-1-{2-[4-(hexyloxy)benzoyloxy]ethyl}-1,2-dimethylpiperidin-1-ium iodide (**7f**)

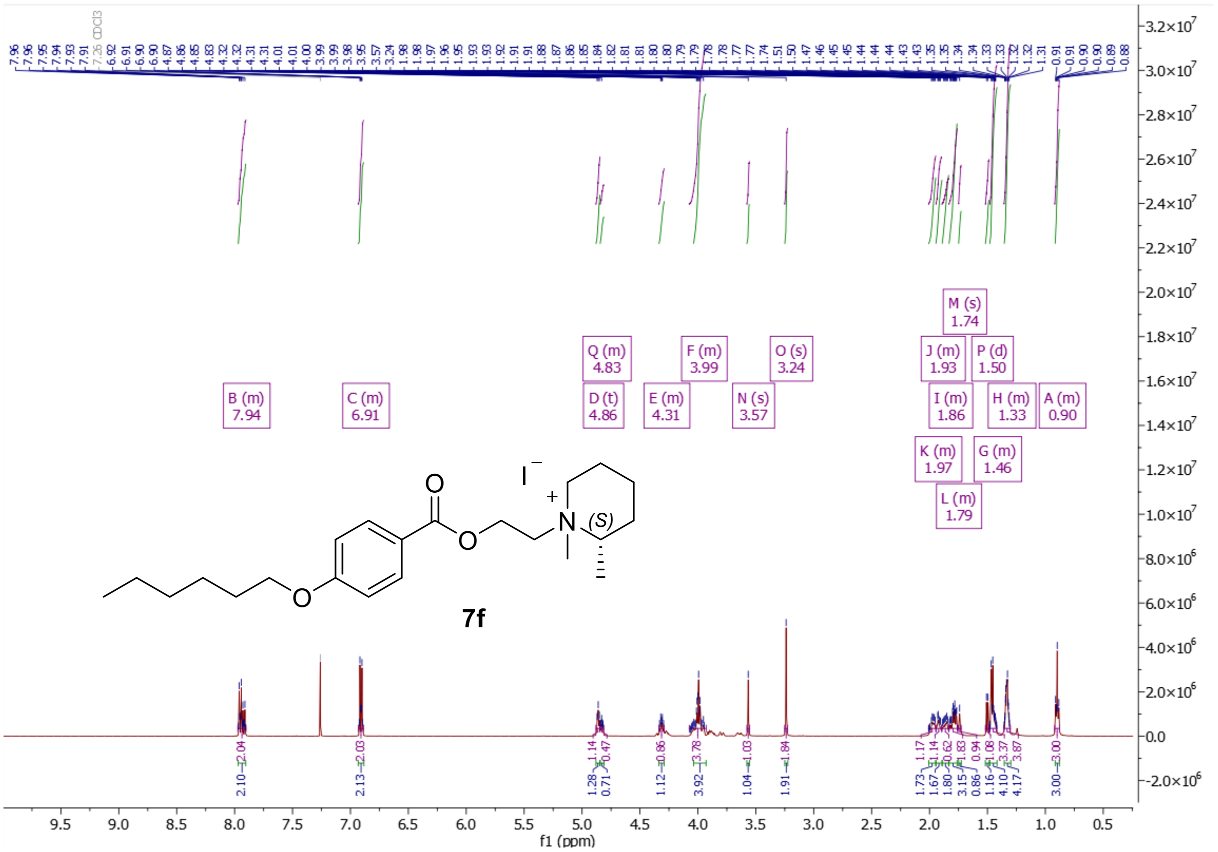

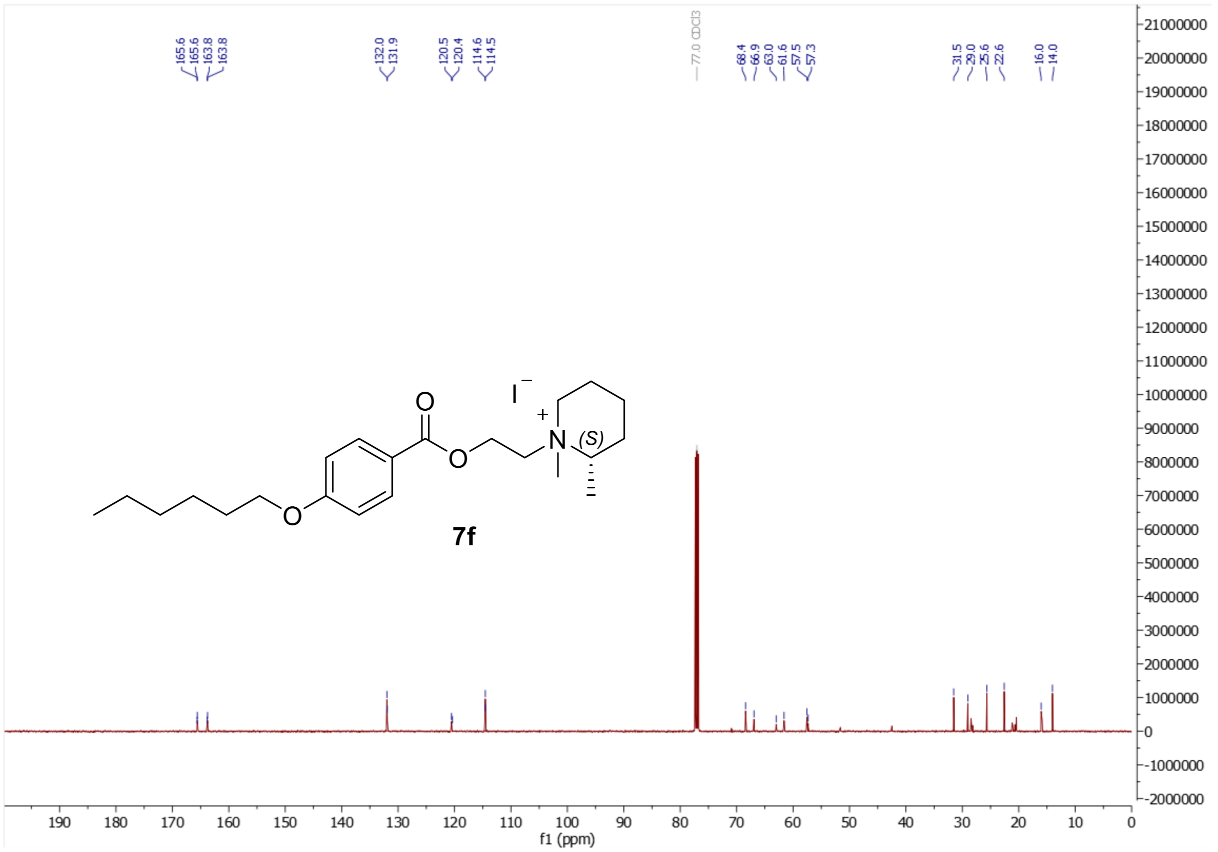

$^{1}H$ and $^{13}C$ NMR spectra of 2-{2-[4-(hexyloxy)benzoyloxy]ethyl}-2-methyl-2-azabicyclo[2.2.2]octan-2-ium iodide (**7g**)

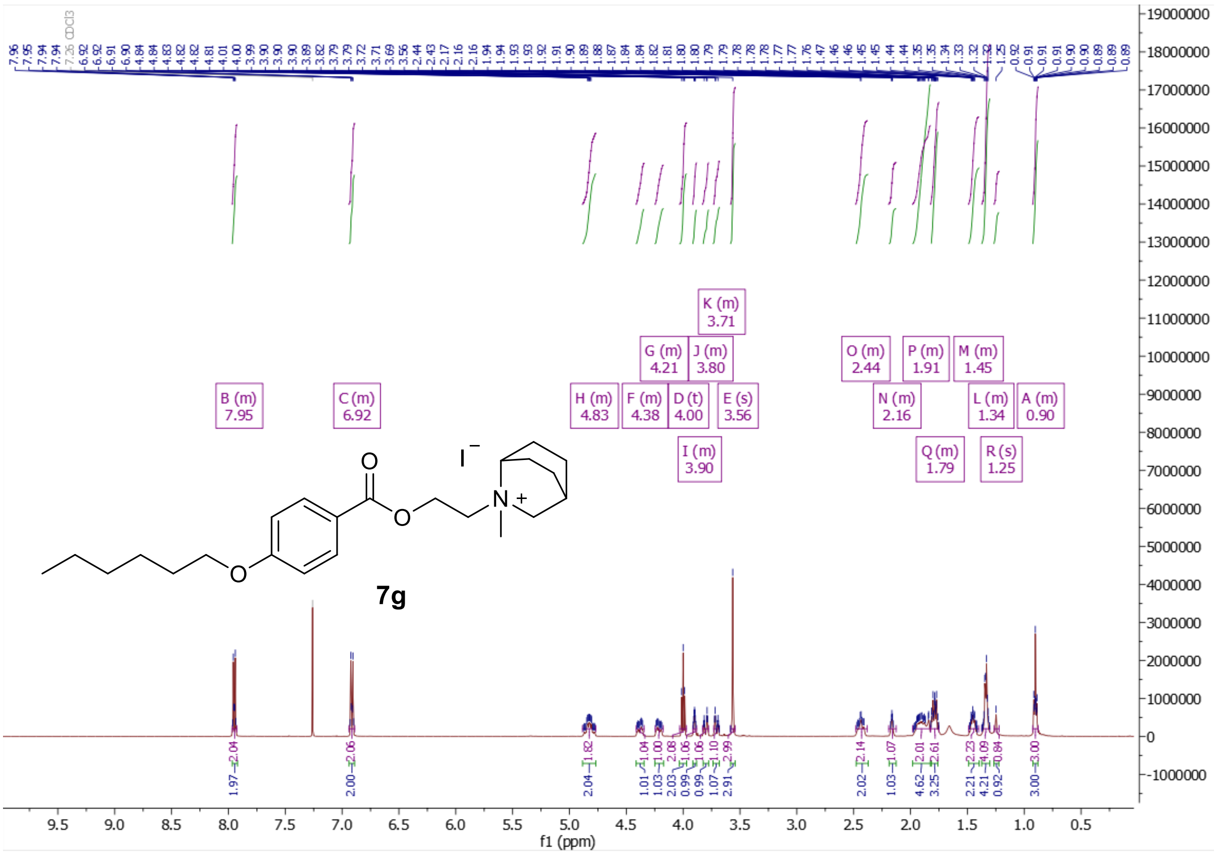

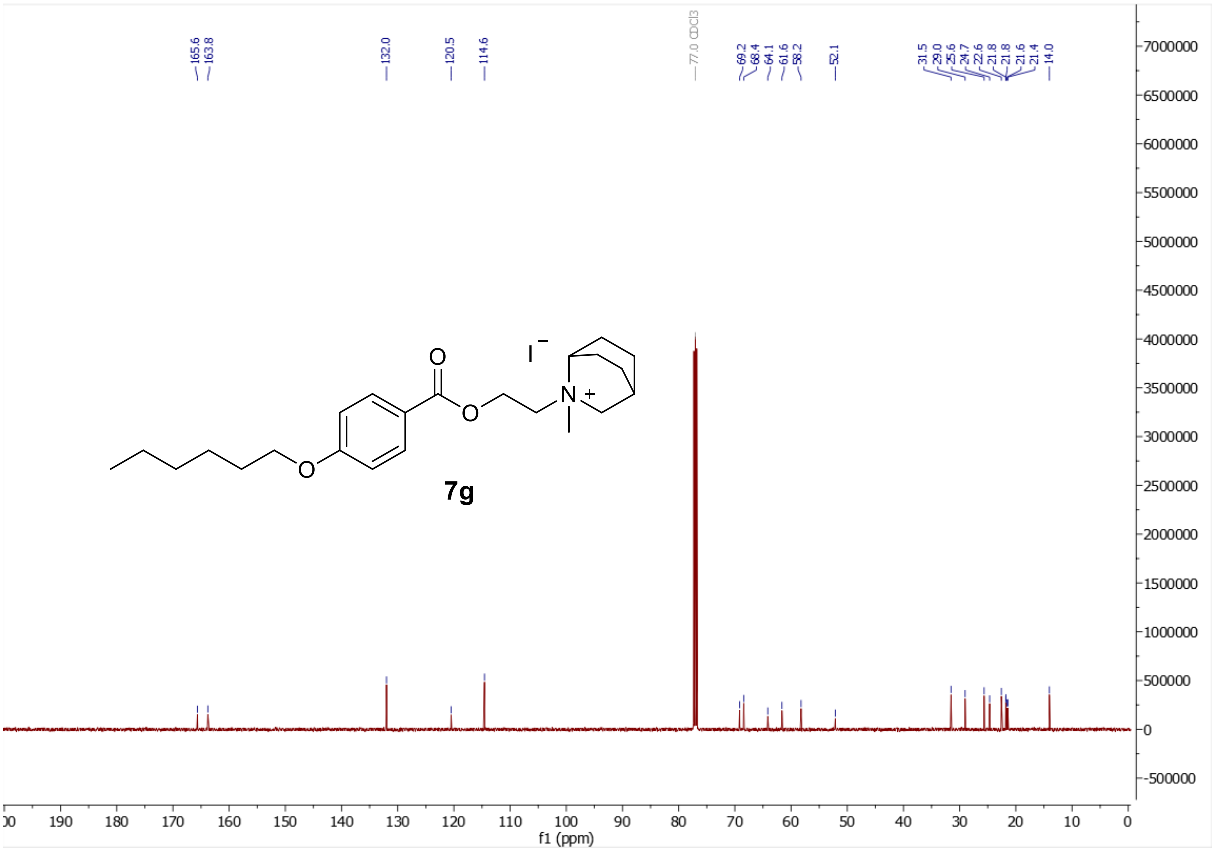

[1]H and [13]C NMR spectra of 1-{2-[4-(hexyloxy)benzoyloxy]ethyl}-1-methyl-2,5-dihydro-1H-pyrrol-1-ium iodide (**7h**)

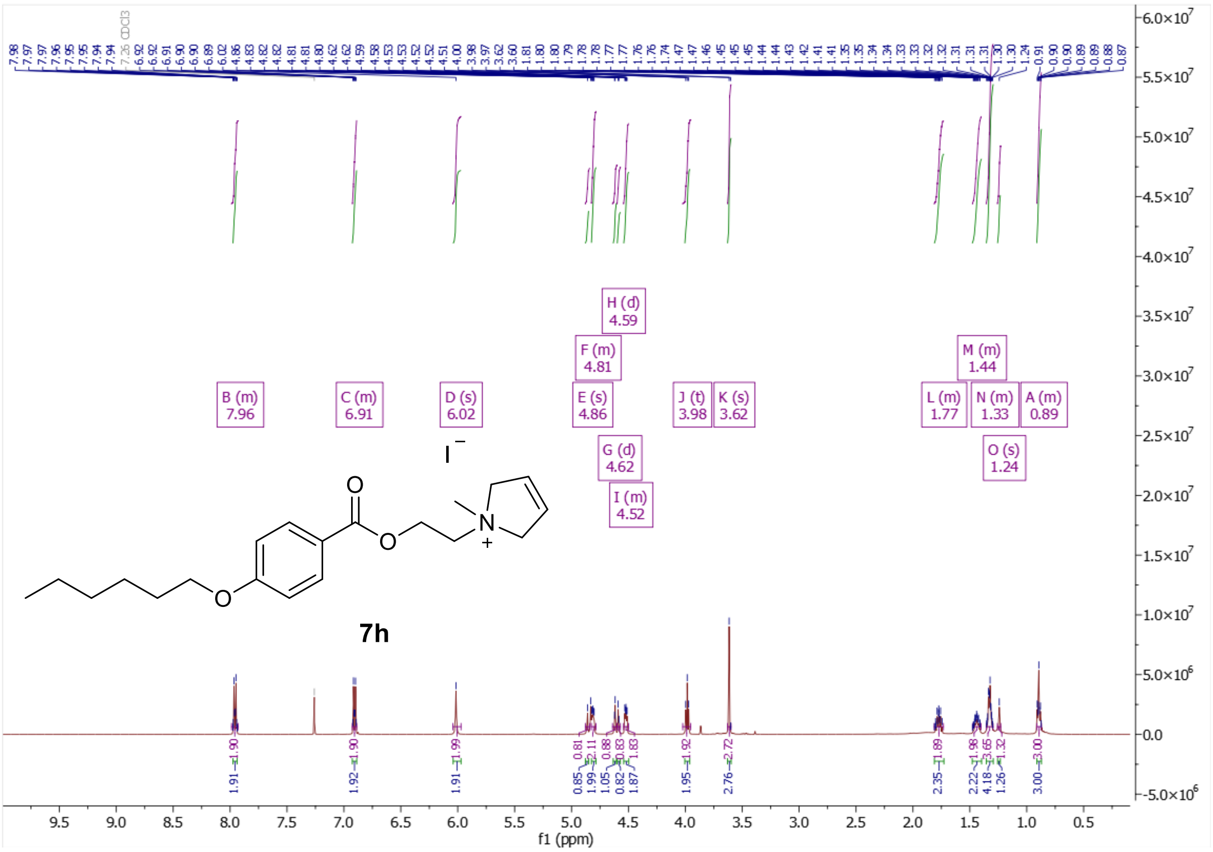

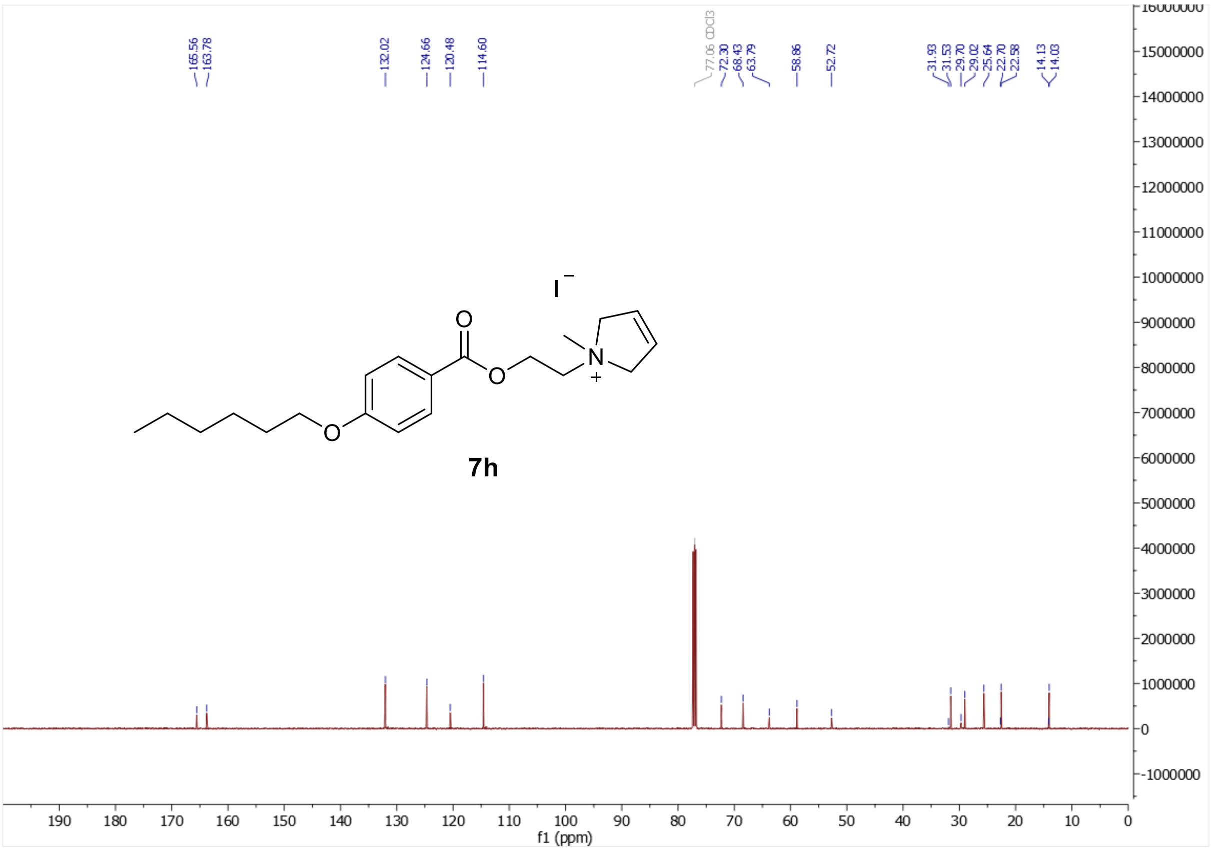

$^{1}$H and $^{13}$C NMR spectra of 1-{2-[4-(hexyloxy)benzoyloxy]ethyl}-1-methylpyrrolidin-1-ium iodide (**7j**)

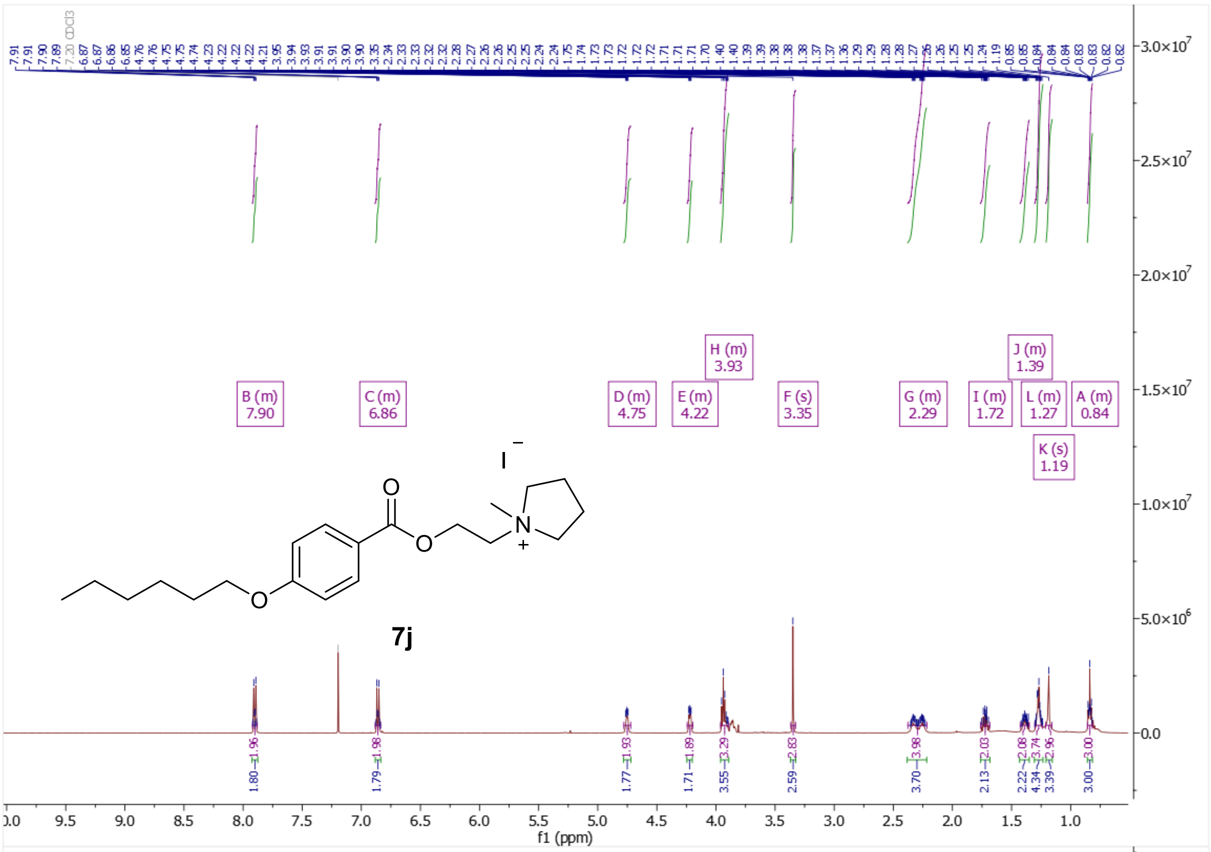

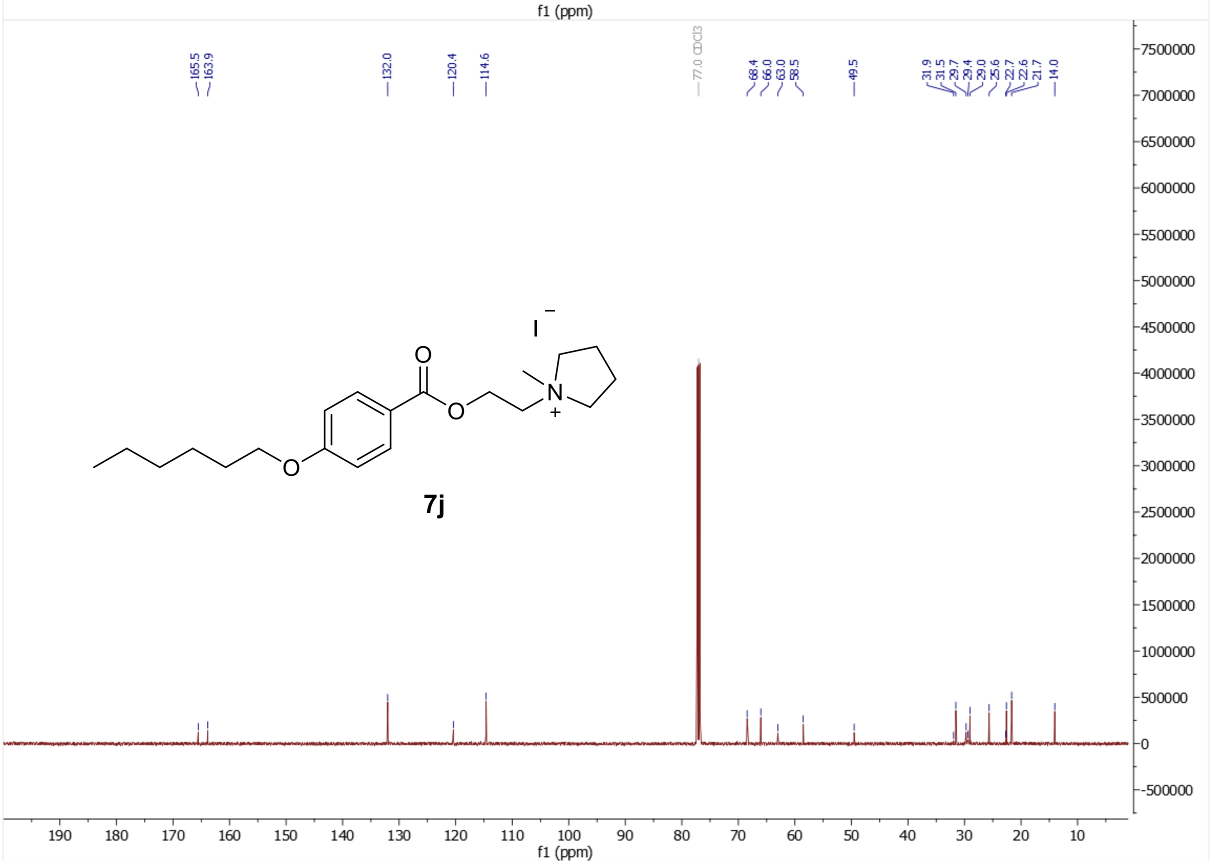

[1]H and [13]C NMR spectra of 2-{2-[4-(hexyloxy)benzoyloxy]ethyl}-2-methyl-2,3-dihydro-1H-isoindol-2-ium iodide (**7k**)

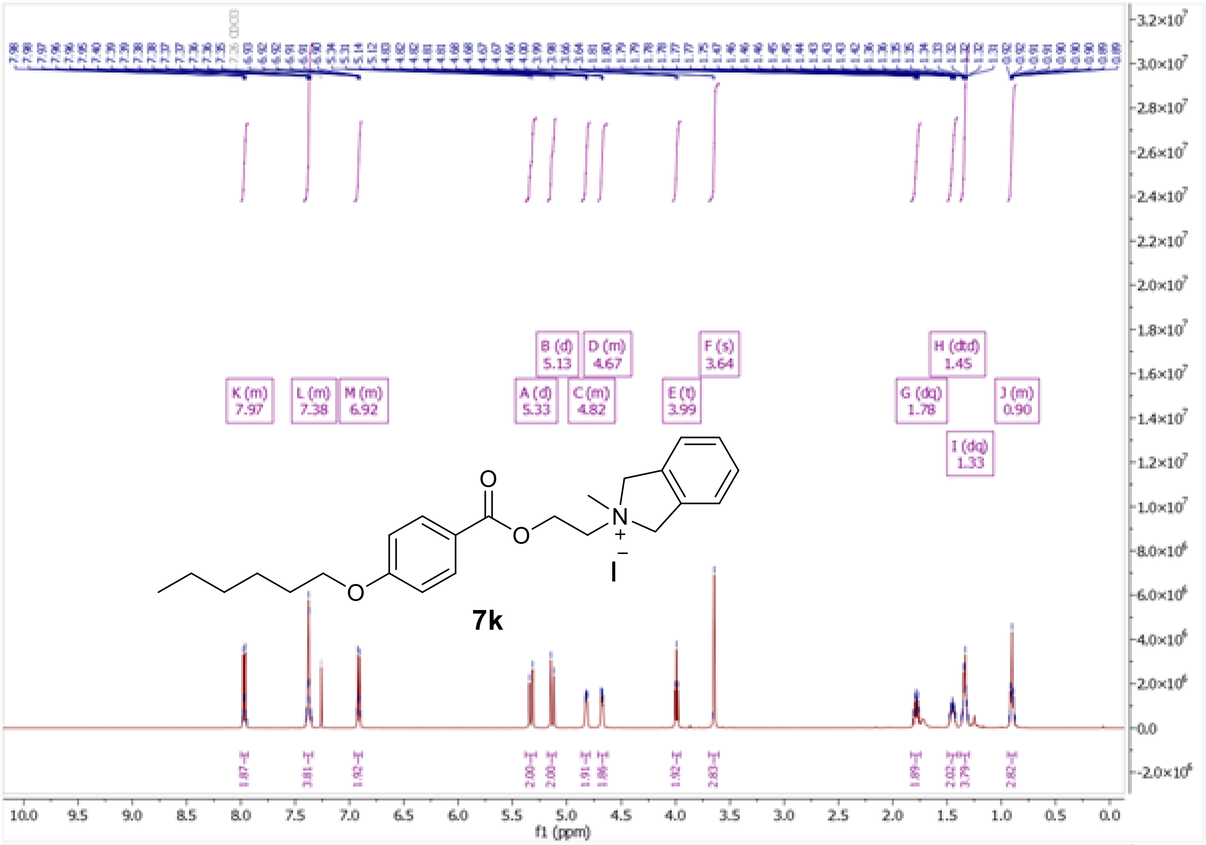

K (m)
7.97
L (m)
7.38
M (m)
6.92
A (d)
5.33
B (d)
5.13
C (m)
4.82
D (m)
4.67
E (t)
3.99
F (s)
3.64
G (dq)
1.78
H (dtd)
1.45
I (dq)
1.33
J (m)
0.90
7k
f1 (ppm)

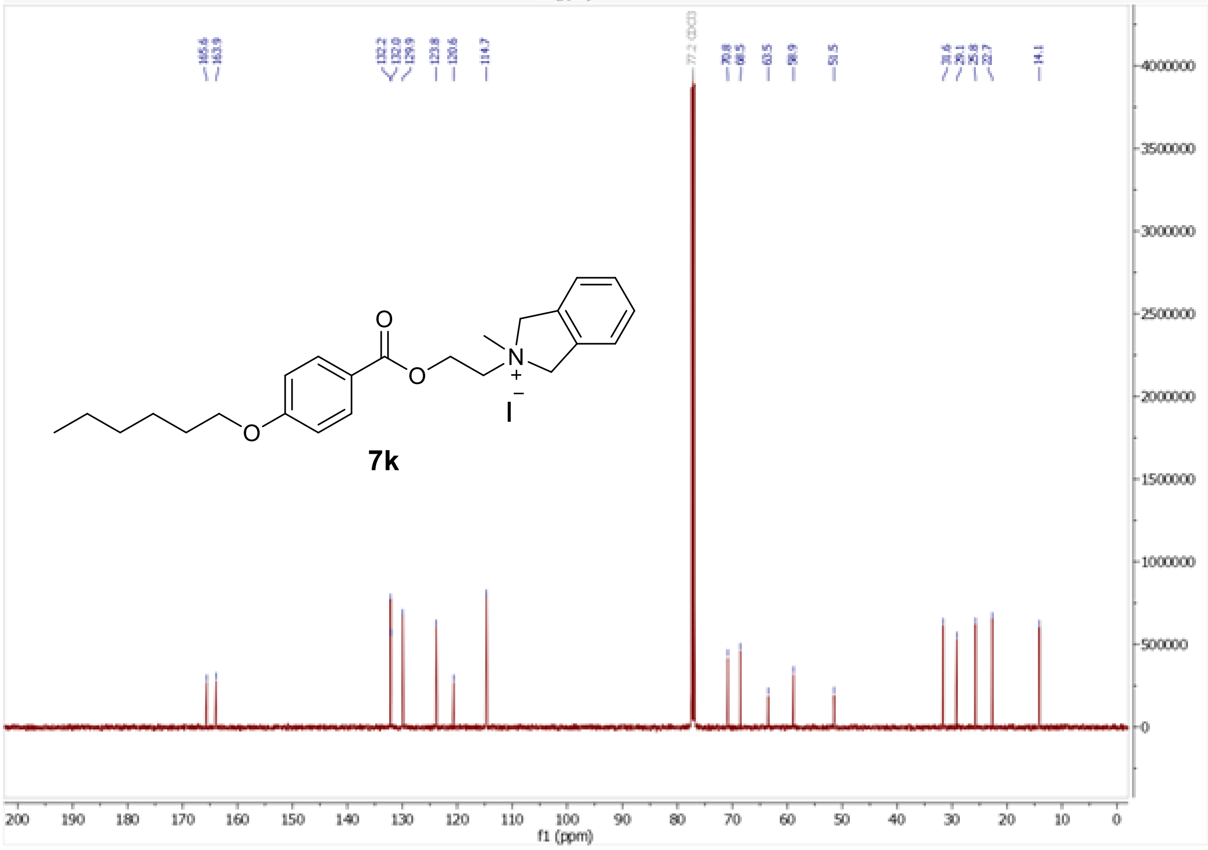

165.6
163.9
132.2
132.0
129.9
123.8
120.6
114.7
77.2 CDCl3
70.8
68.5
63.5
58.9
51.5
31.6
29.1
25.8
22.7
14.1
7k
f1 (ppm)

<sup></sup>[1]H and [13]C NMR spectra of (3*S*)-3-{[4-(hexyloxy)benzoyloxy]methyl}-1,1-dimethylpiperidin-1-ium iodide (**7o**)

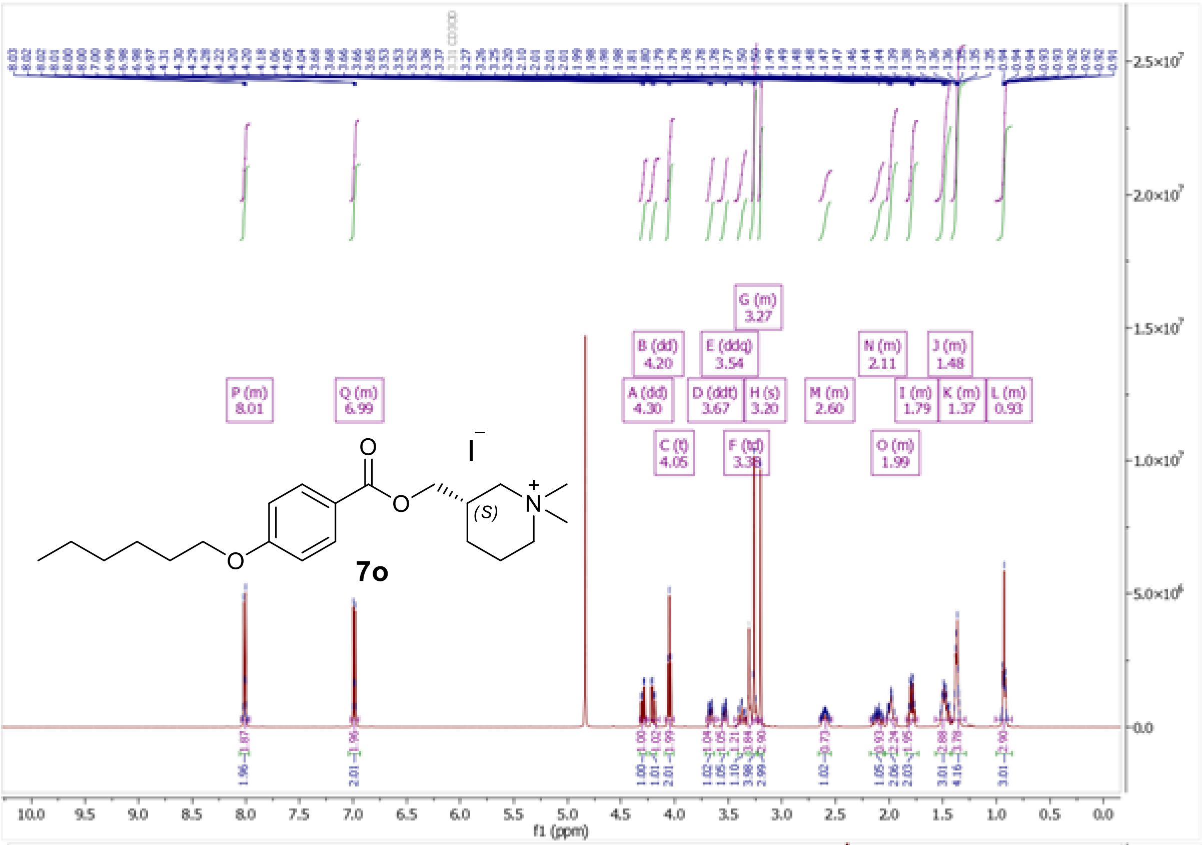

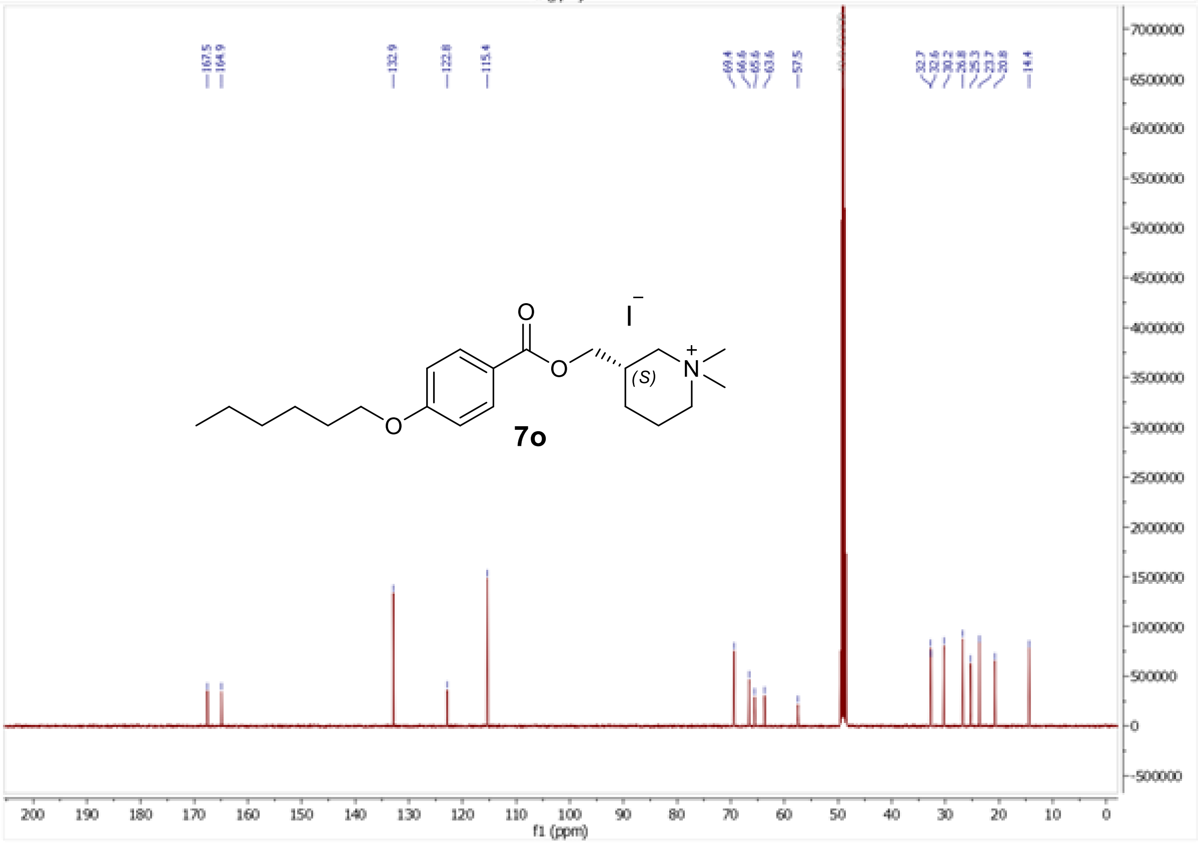

[1]H and [13]C NMR spectra of 3-{[4-(hexyloxy)benzoyloxy]methyl}-1,1-dimethylpiperidin-1-ium iodide (**7p**)

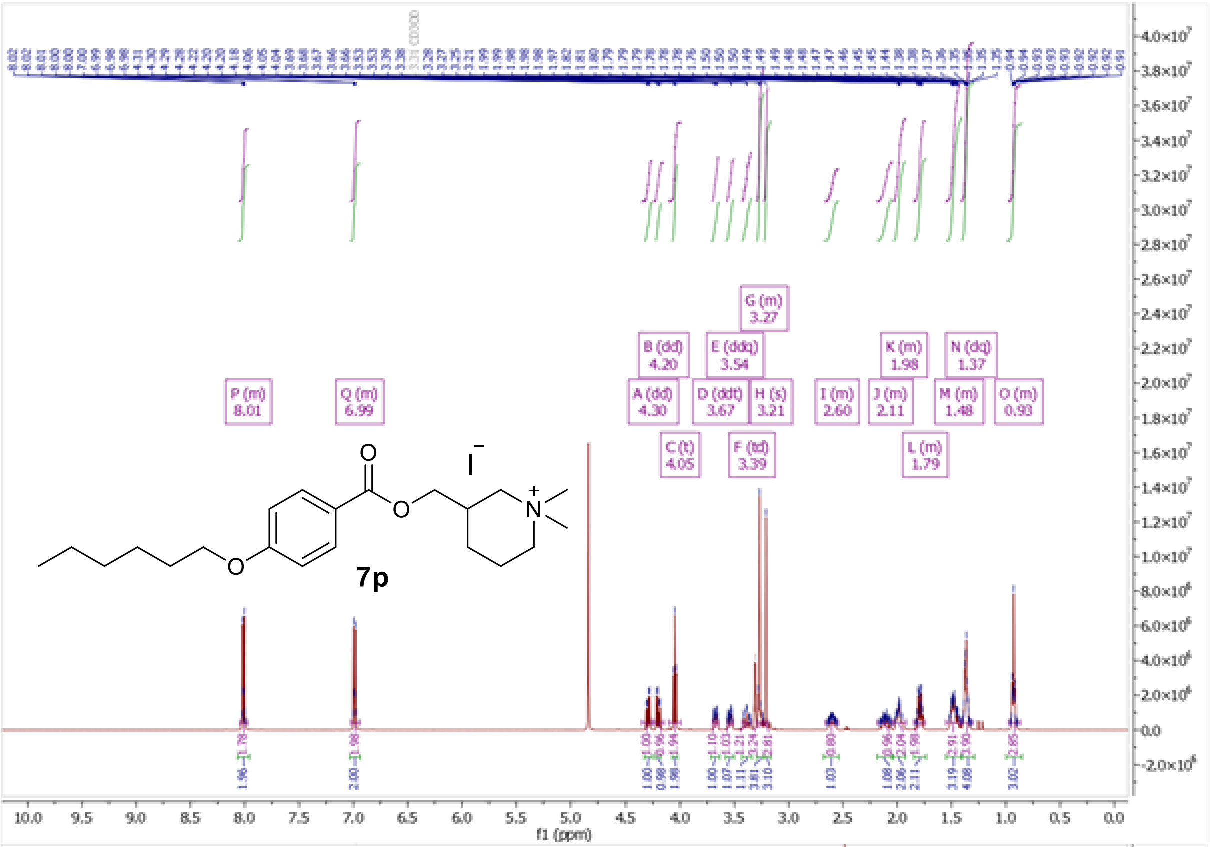

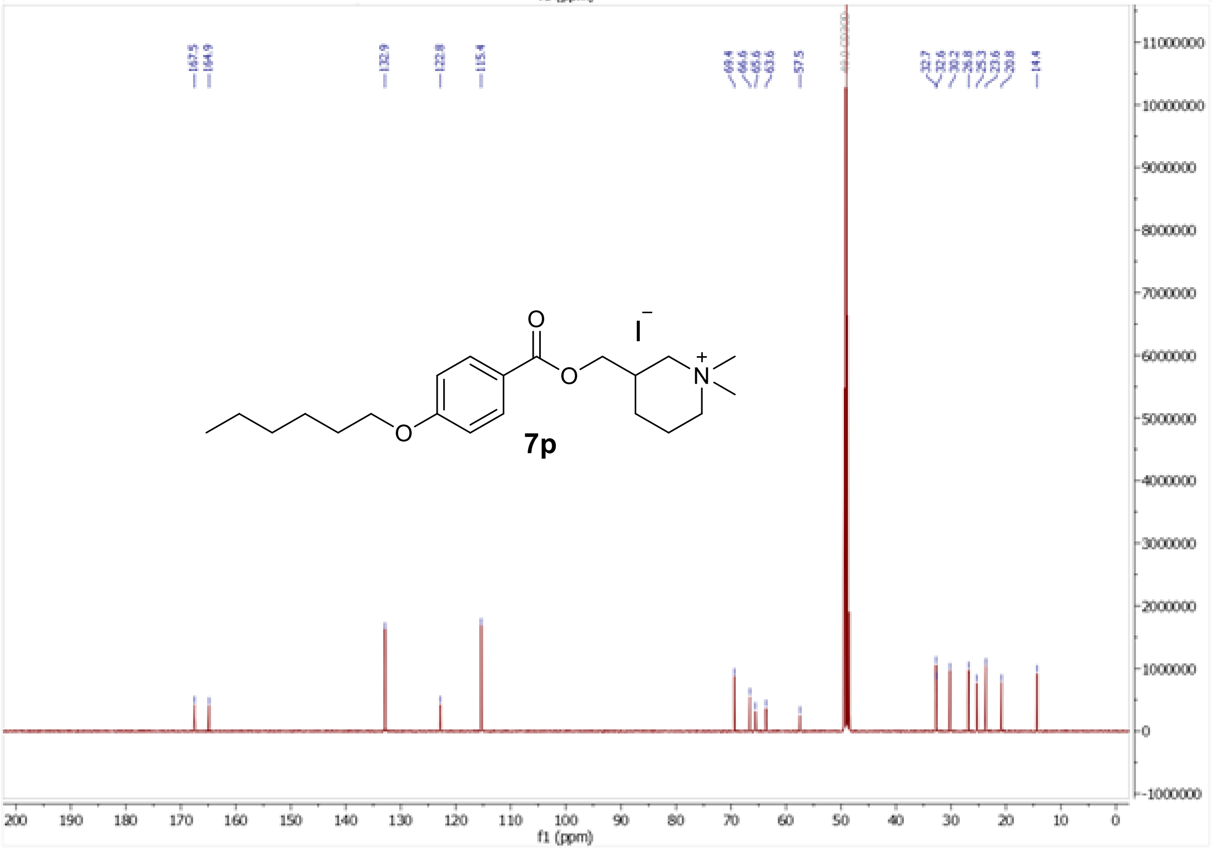

$^{1}H$ and $^{13}C$ NMR spectra of (3*R*)-3-{[4-(hexyloxy)benzoyloxy]methyl}-1,1-dimethylpiperidin-1-ium iodide (**7r**)

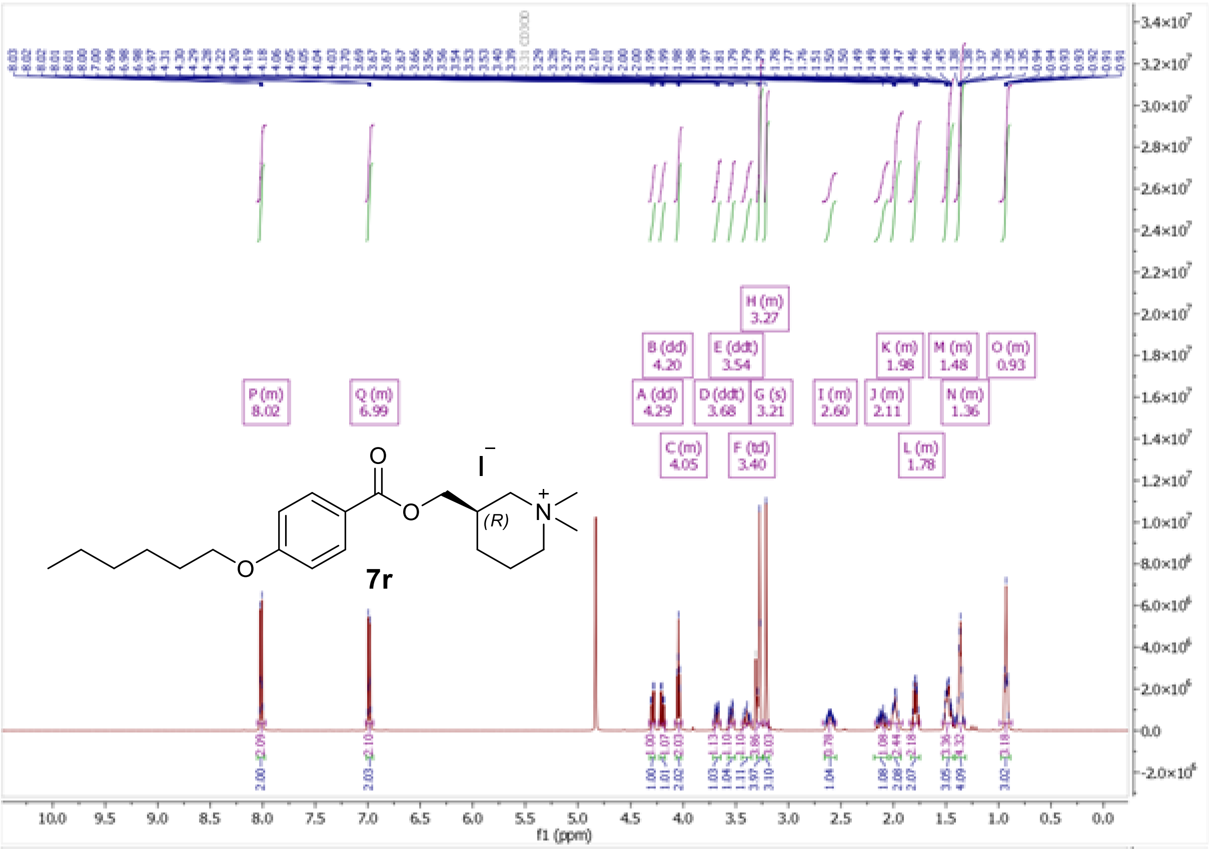

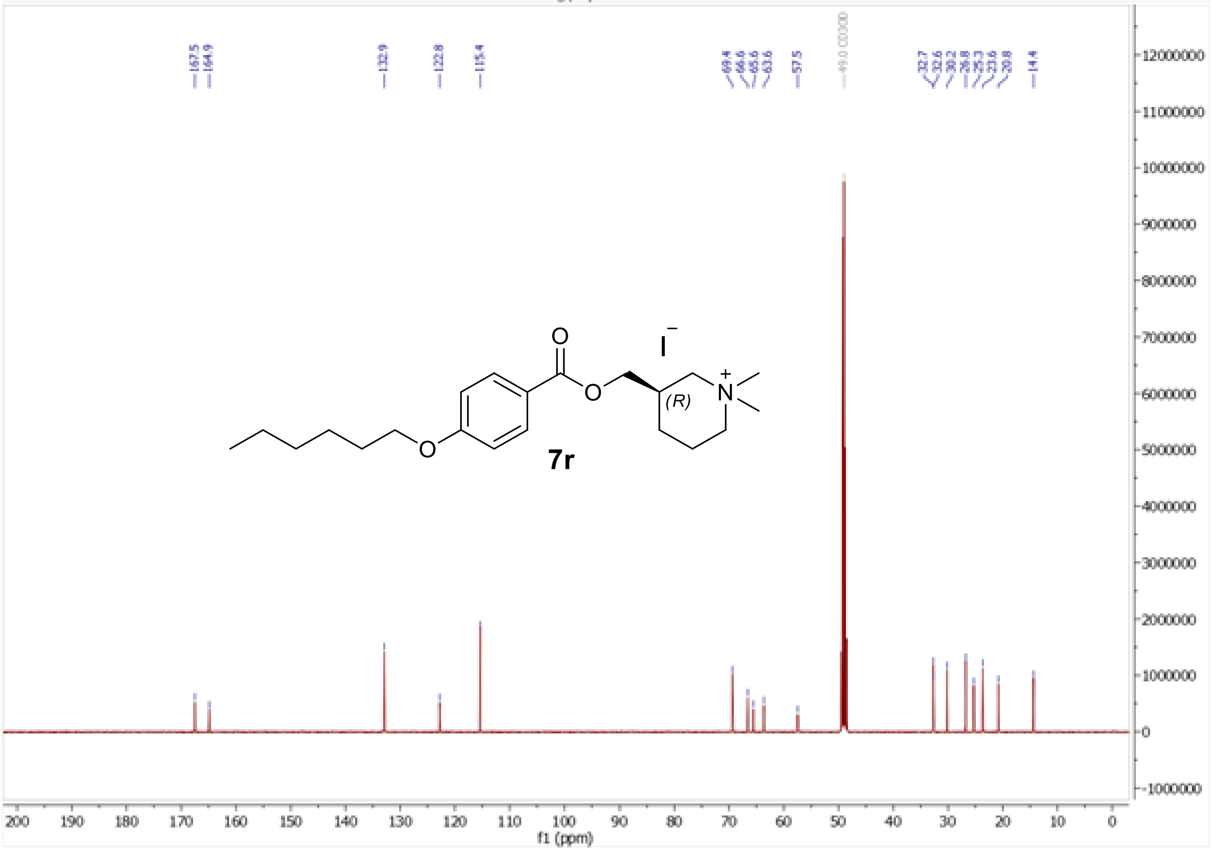

$^{1}H$ and $^{13}C$ NMR spectra of 4-[4-(hexyloxy)benzoyloxy]-1,1-dimethylpiperidin-1-ium iodide (**7s**)

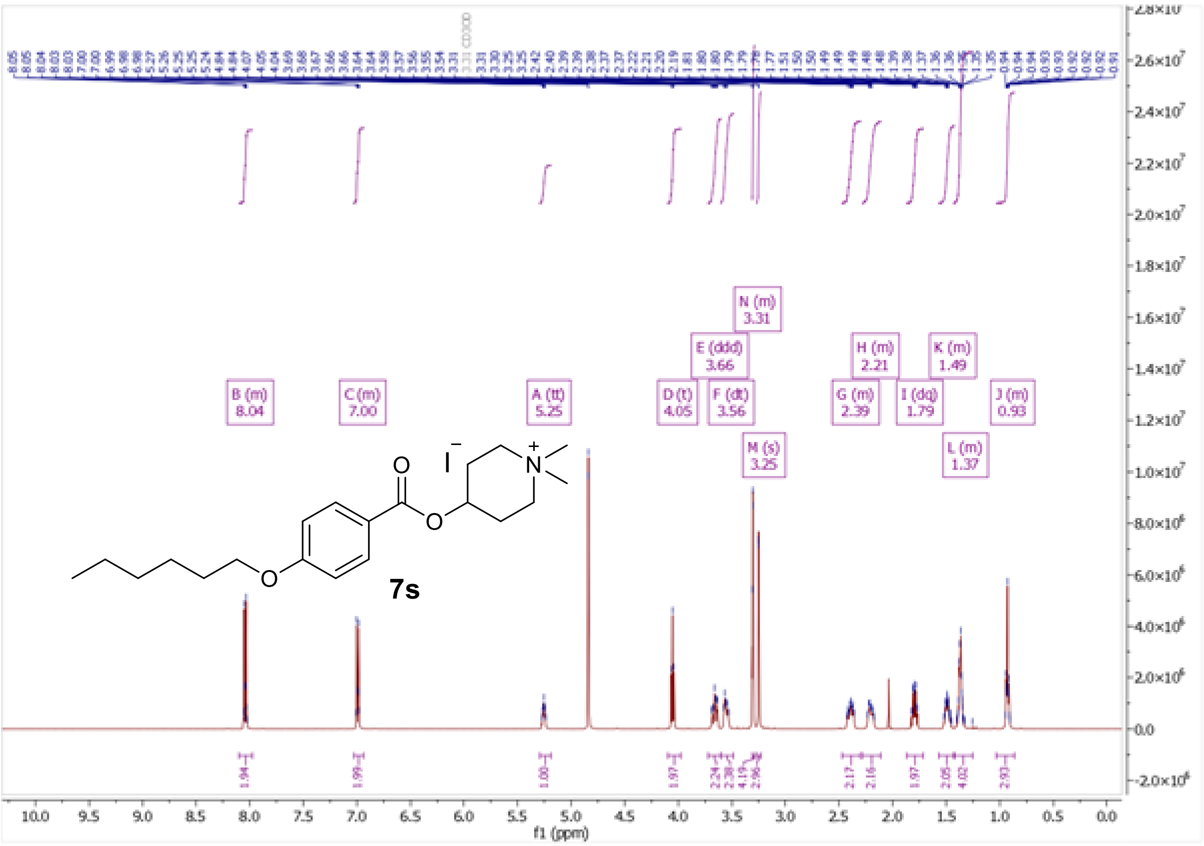
B (m)
8.04
C (m)
7.00
A (tt)
5.25
D (t)
4.05
E (ddd)
3.66
F (dt)
3.56
N (m)
3.31
M (s)
3.25
G (m)
2.39
H (m)
2.21
I (dq)
1.79
K (m)
1.49
L (m)
1.37
J (m)
0.93
7s
f1 (ppm)

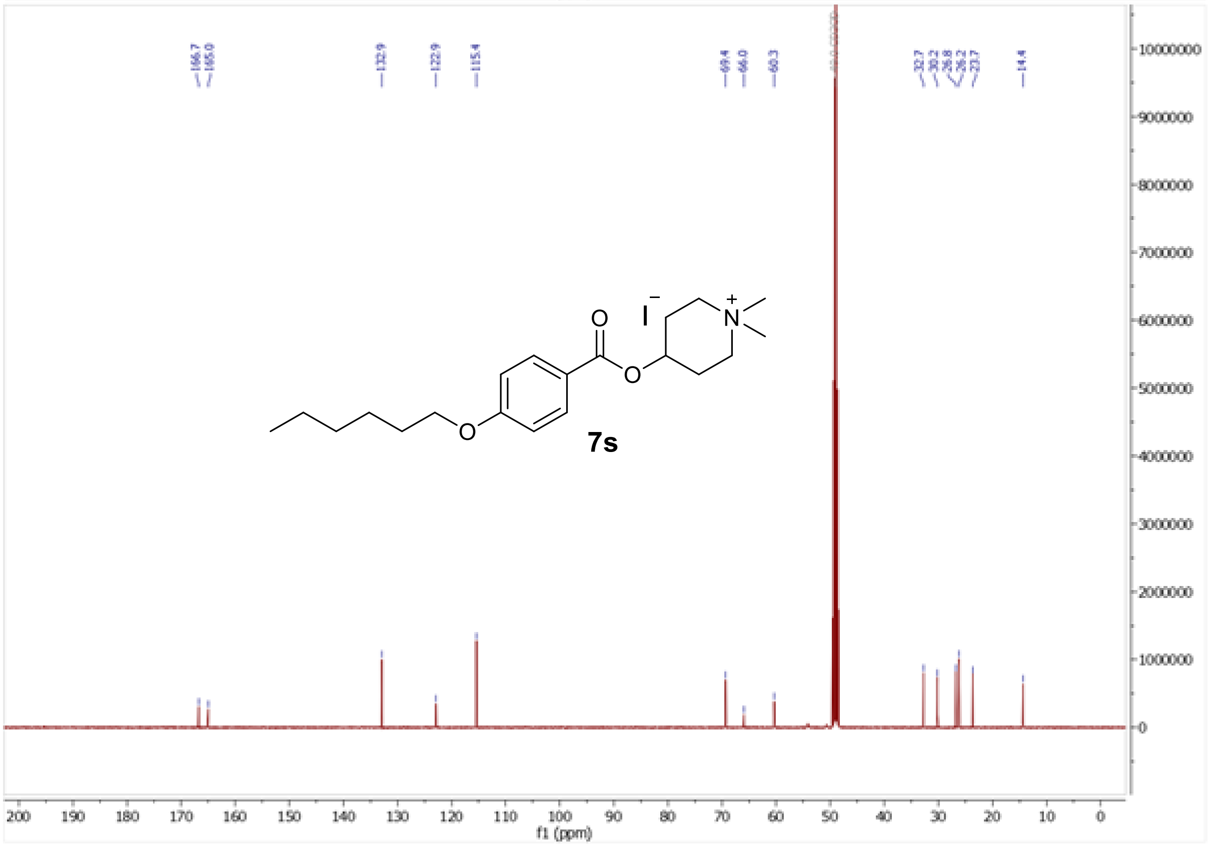
166.7
165.0
132.9
122.9
115.4
69.4
66.0
60.3
32.7
30.2
26.8
26.2
23.7
14.4
7s
f1 (ppm)

<sup></sup>[1]H and [13]C NMR spectra of 2-{[4-(hexyloxy)benzoyloxy]methyl}-1,1-dimethylpyrrolidin-1-ium iodide (**7t**)

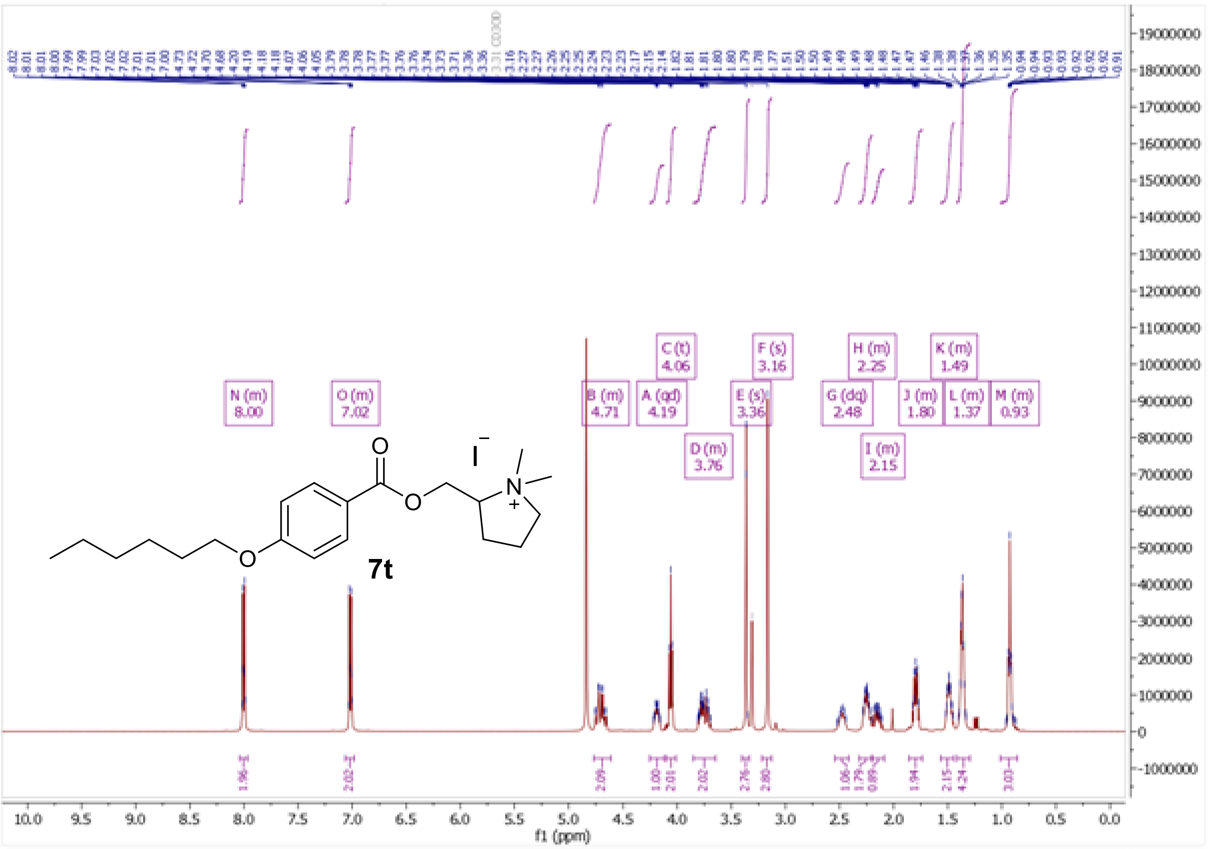

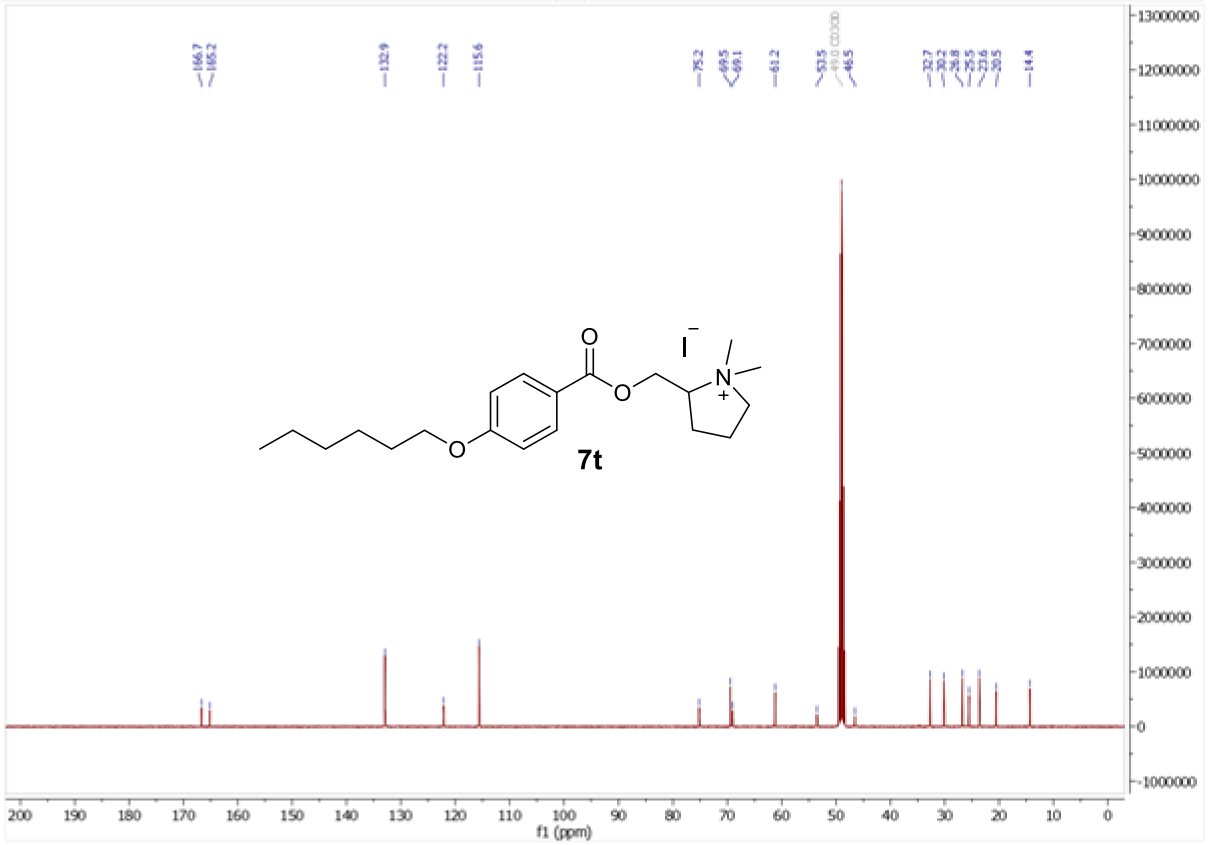

$^1$H and $^{13}$C NMR spectra of (2*S*)-2-{[4-(hexyloxy)benzoyloxy]methyl}-1,1-dimethylpyrrolidin-1-ium iodide (**7w**)

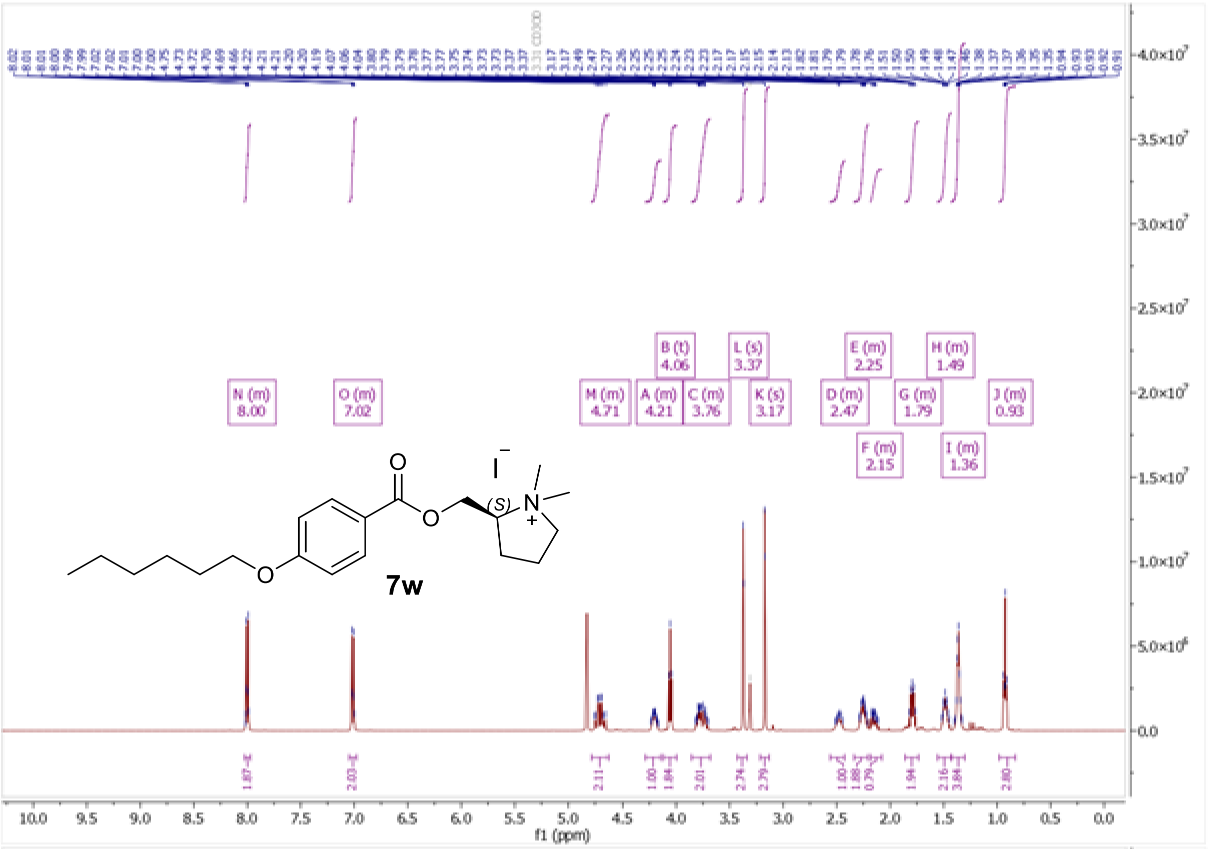

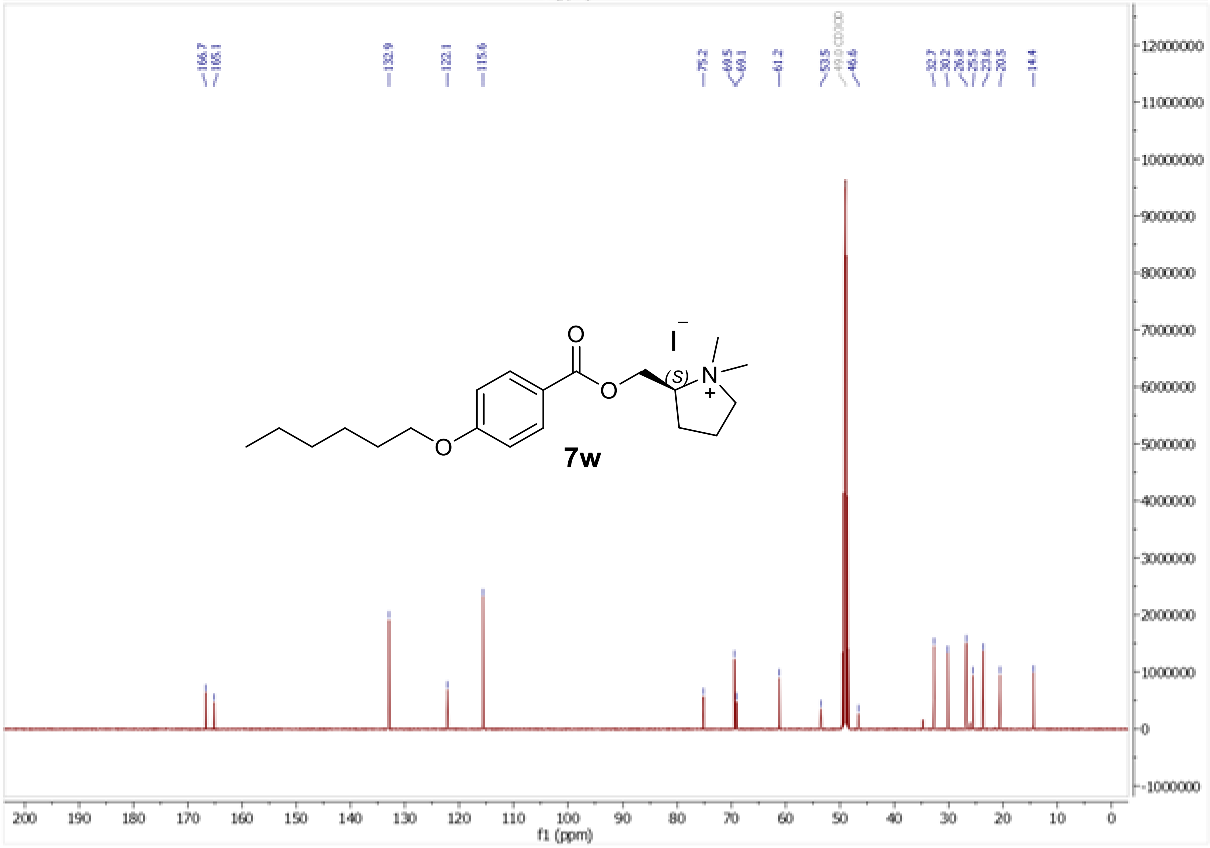

$^{1}H$ and $^{13}C$ NMR spectra of (2*R*)-2-{[4-(hexyloxy)benzoyloxy]methyl}-1,1-dimethylpyrrolidin-1-ium iodide (**7x**)

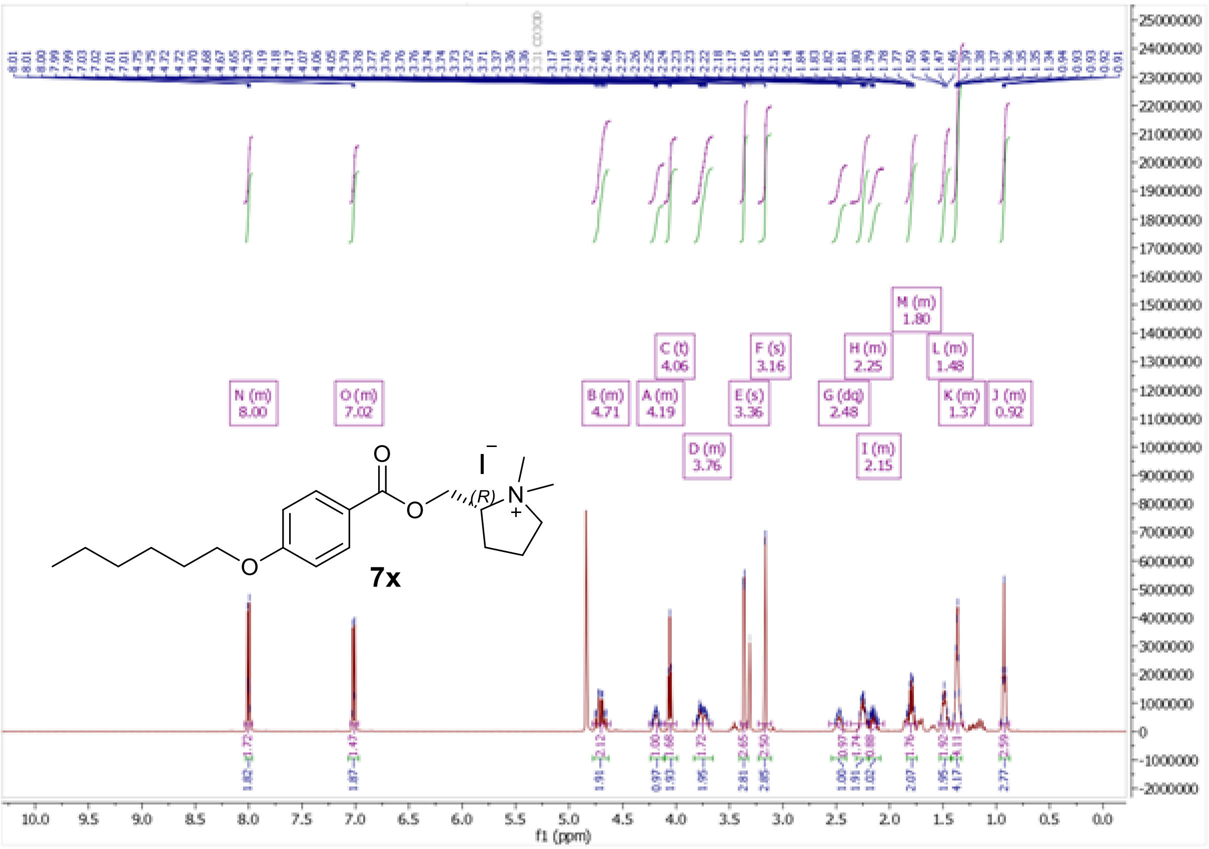

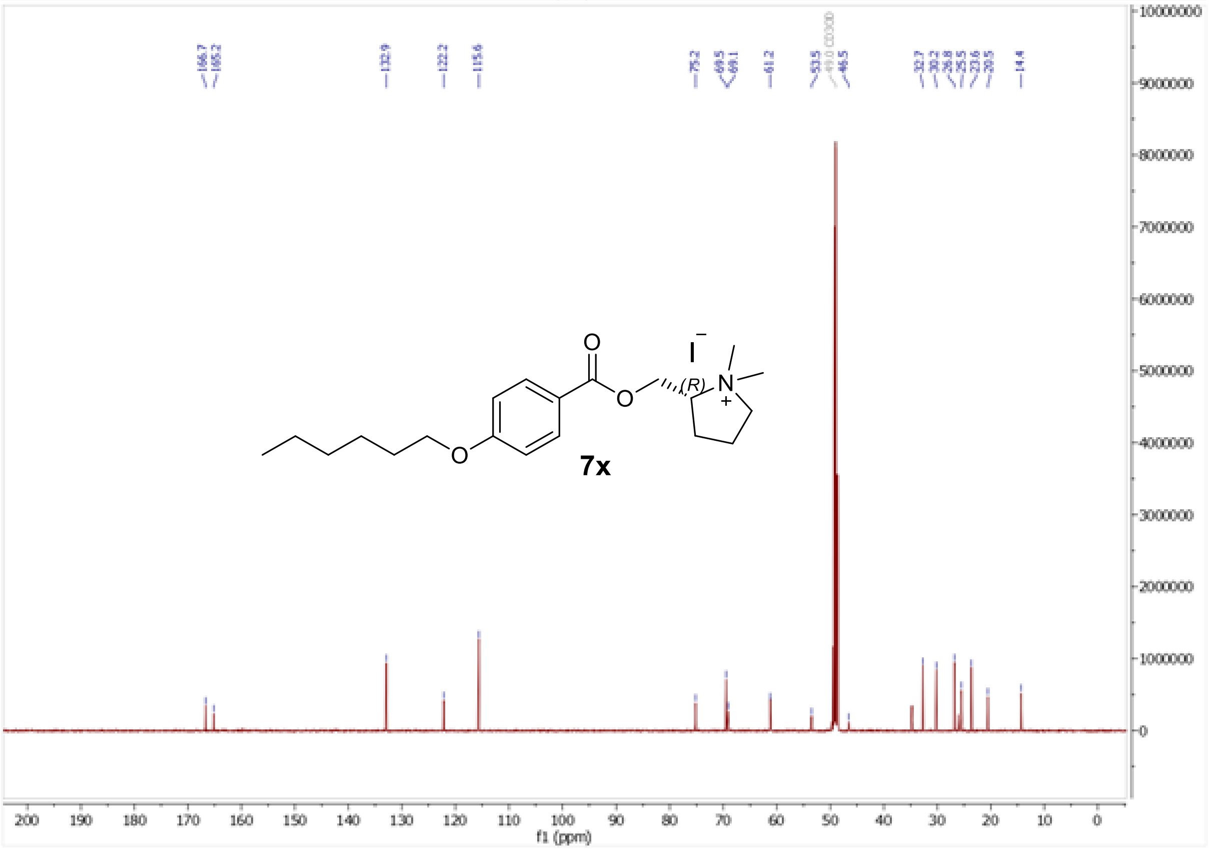

$^{1}H$ and $^{13}C$ NMR spectra of 3-{[4-(hexyloxy)benzoyloxy]methyl}-1,1-dimethylpyrrolidin-1-ium iodide (**7y**)

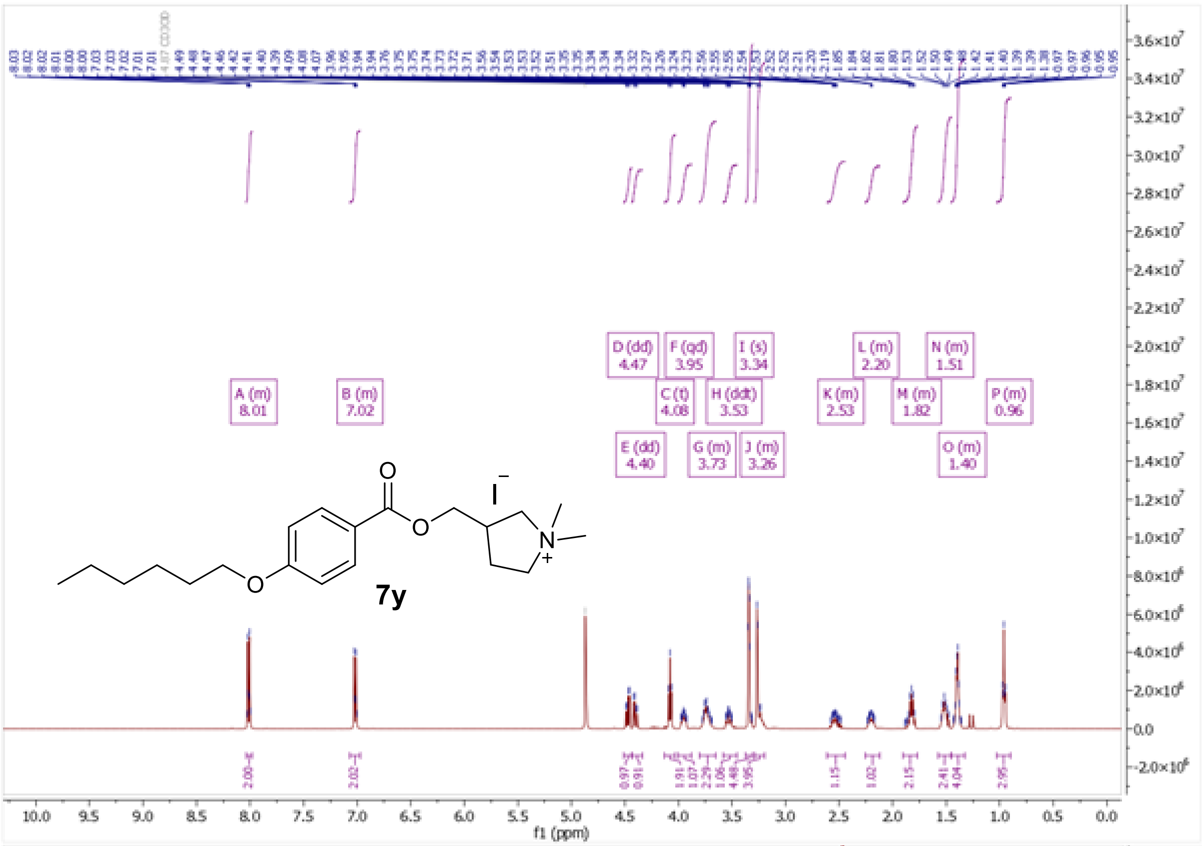
A (m) 8.01
B (m) 7.02
C (t) 4.08
D (dd) 4.47
E (dd) 4.40
F (qd) 3.95
G (m) 3.73
H (ddt) 3.53
I (s) 3.34
J (m) 3.26
K (m) 2.53
L (m) 2.20
M (m) 1.82
N (m) 1.51
O (m) 1.40
P (m) 0.96
7y
f1 (ppm)

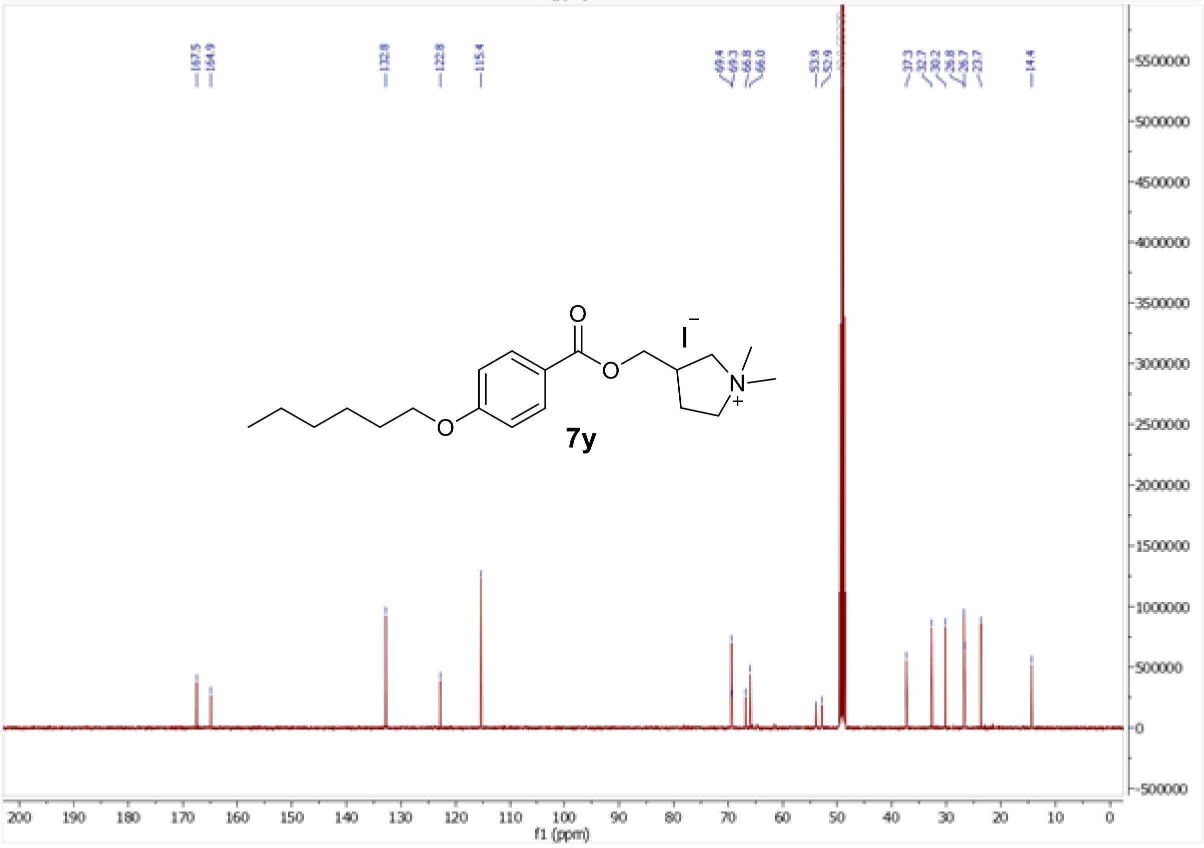
167.5
164.9
132.8
122.8
115.4
69.4
69.3
66.8
66.0
53.9
52.9
37.3
32.7
30.2
26.8
26.7
23.7
14.4
7y
f1 (ppm)

[1]H and [13]C NMR spectra of (3*S*)-3-{[4-(hexyloxy)benzoyloxy]methyl}-1,1-dimethylpyrrolidin-1-ium iodide (**7z**)

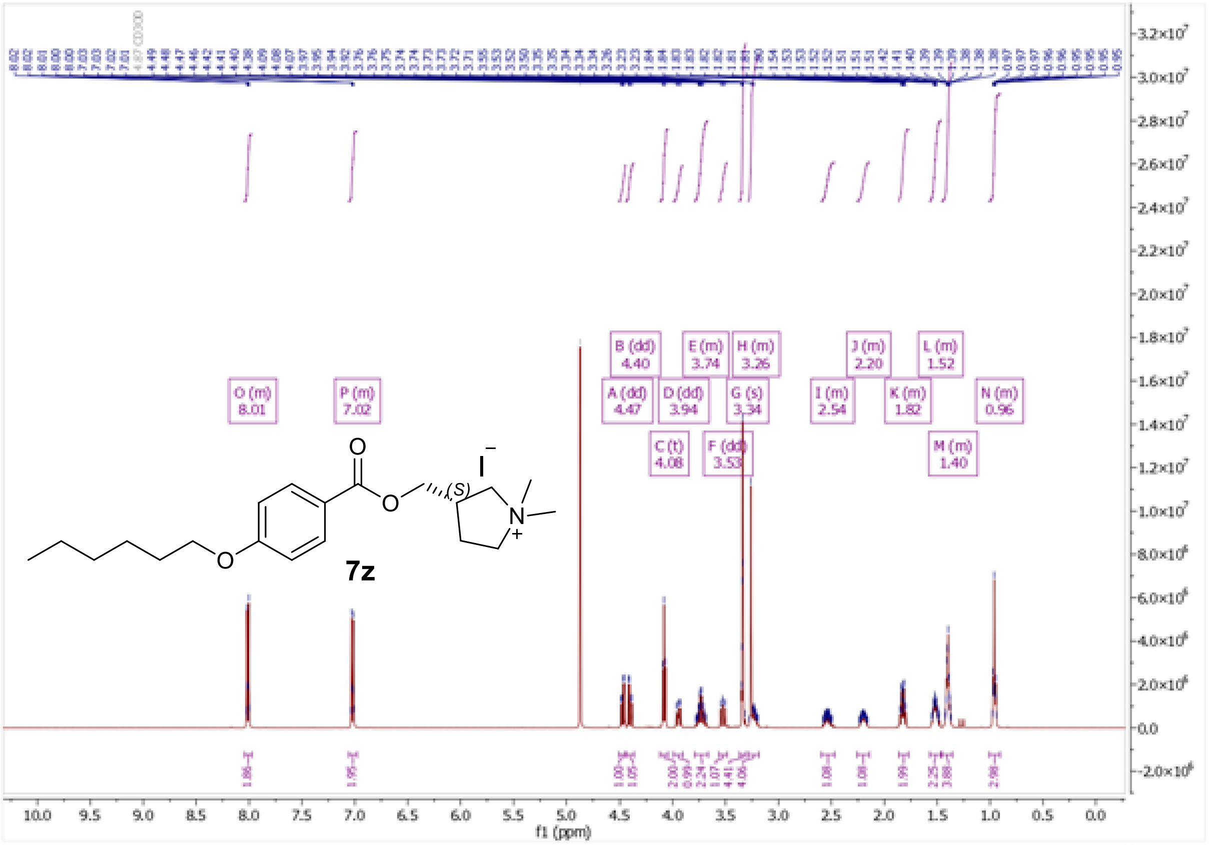

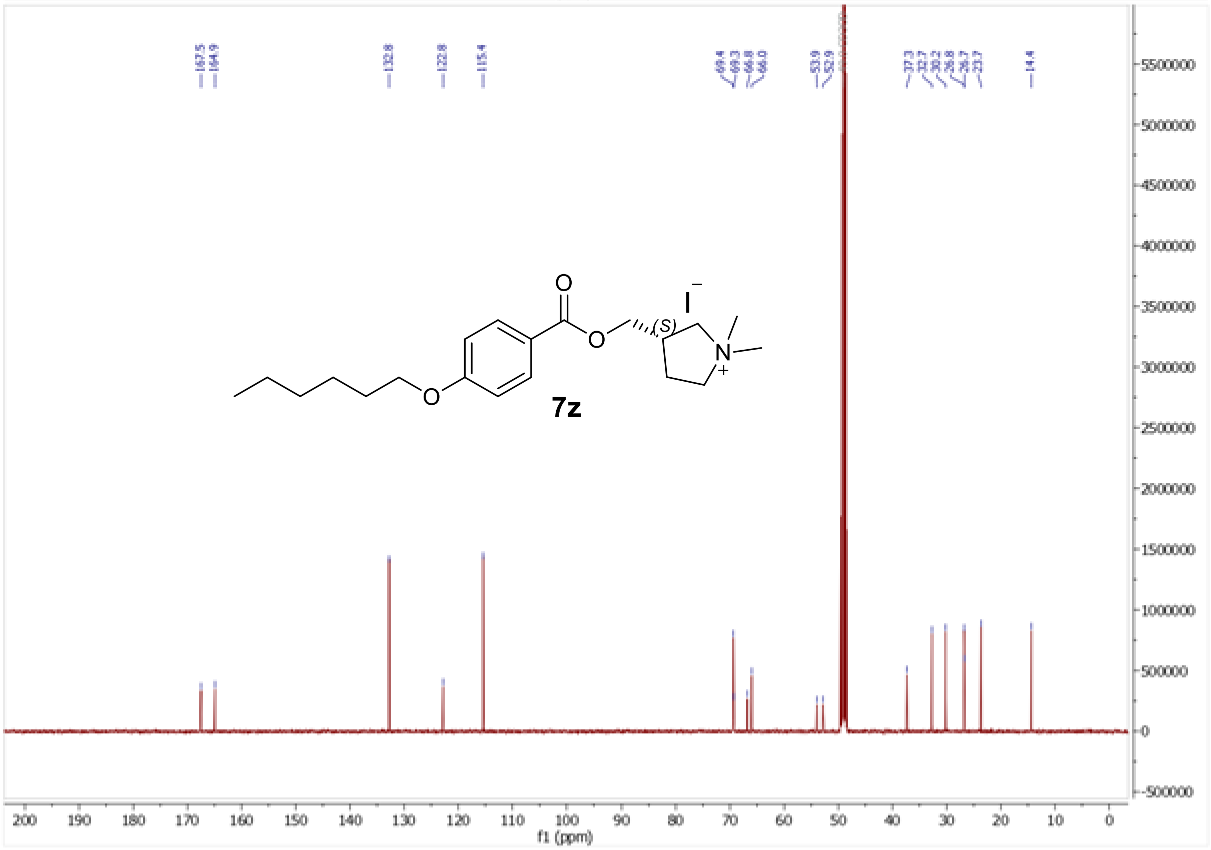

[1]H and [13]C NMR spectra of quinuclidin-3-yl 4-hexyloxybenzoate hydrochloride (**12a**)

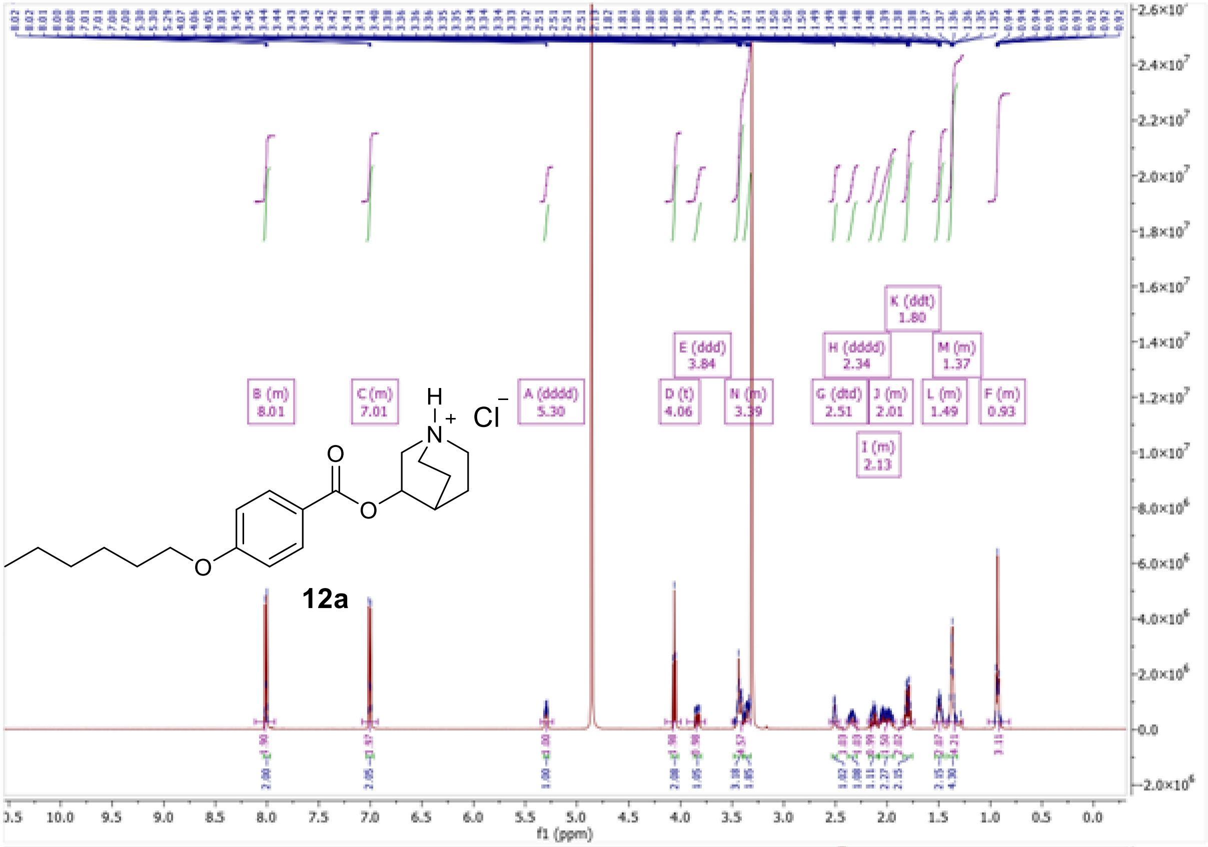

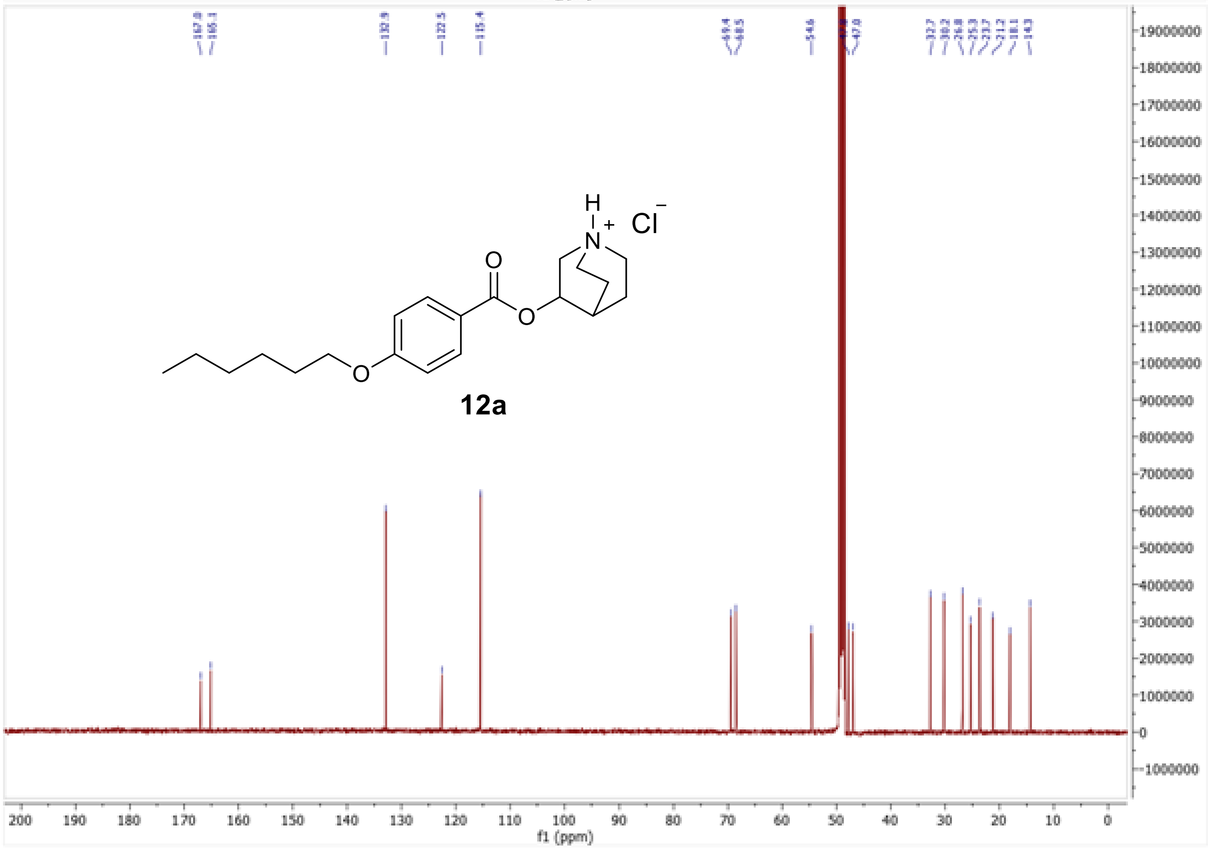

$^{1}H$ and $^{13}C$ NMR spectra of 1-methylquinuclidin-3-yl 4-hexyloxybenzoate iodide (**12b**)

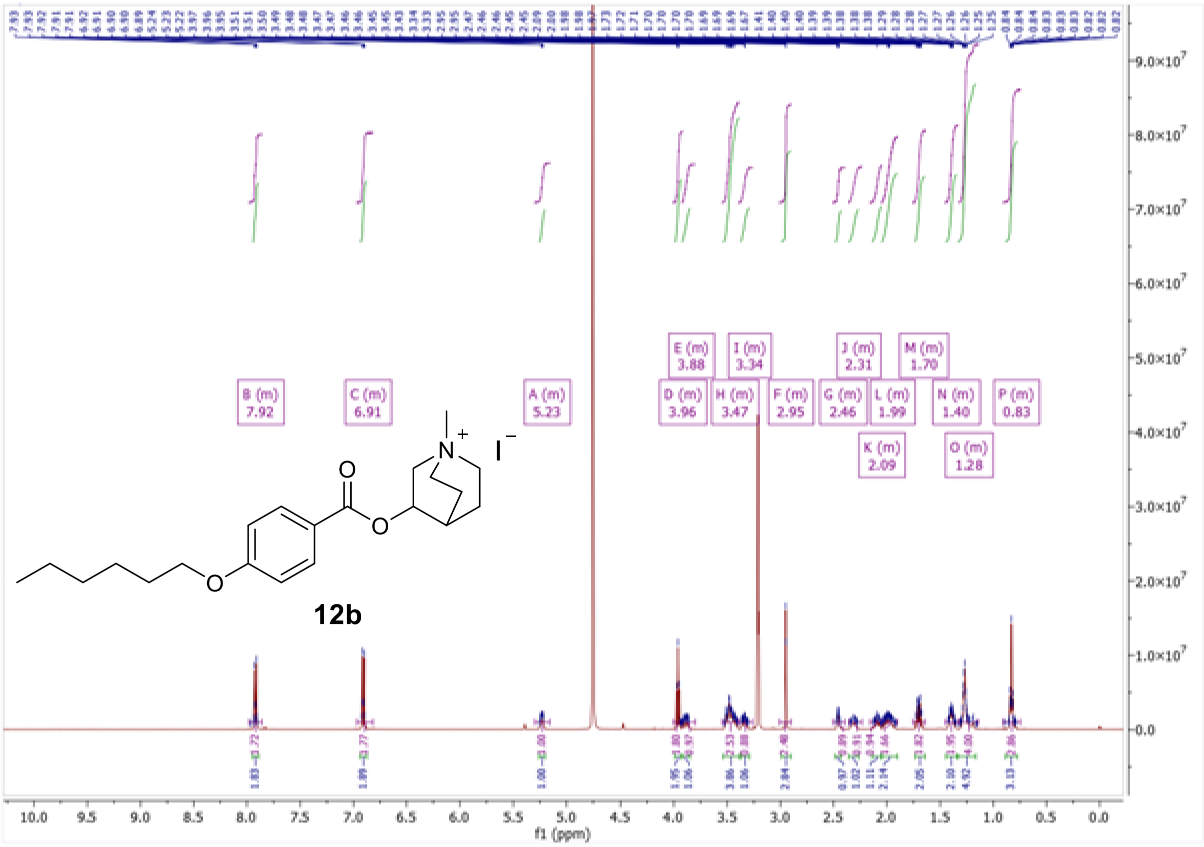

12b

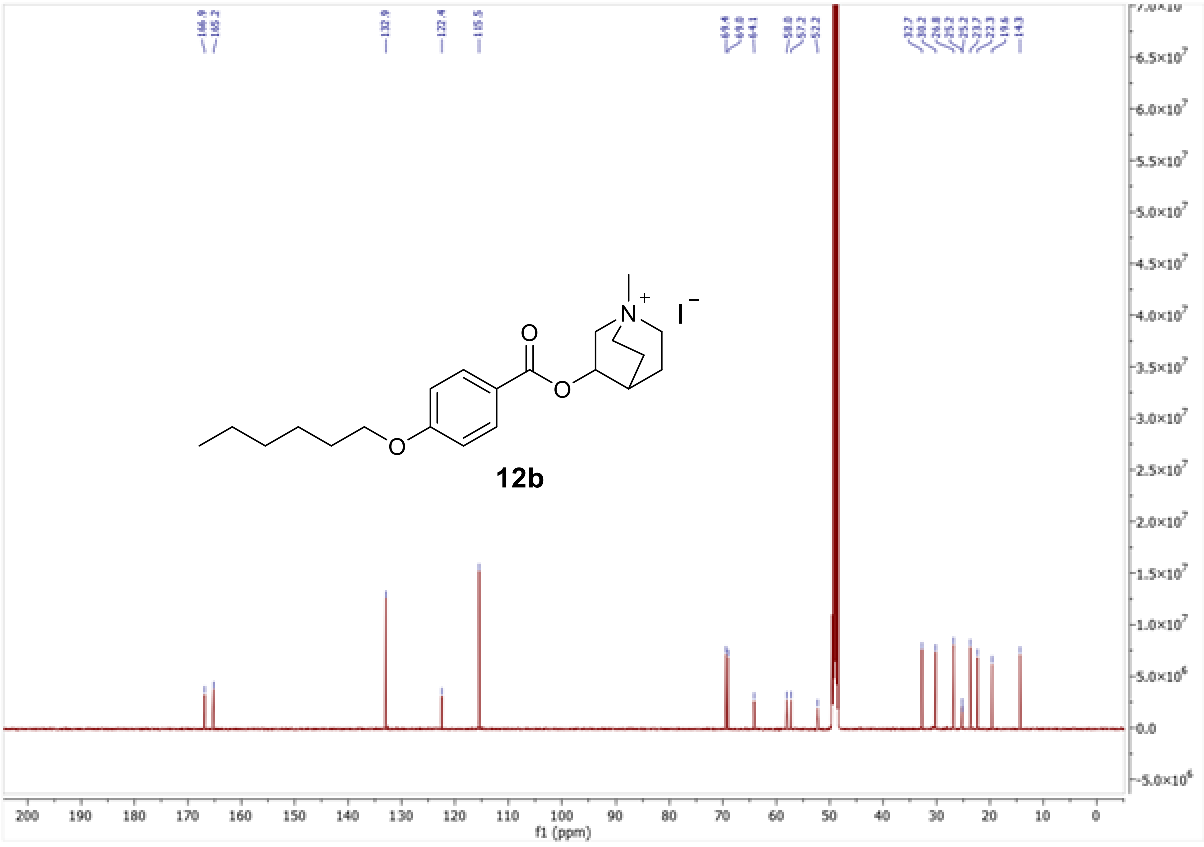

12b

<sup></sup>

[1]H and [13]C NMR spectra of quinuclidin-3-yl 4-butoxybenzoate hydrochloride (**13a**)

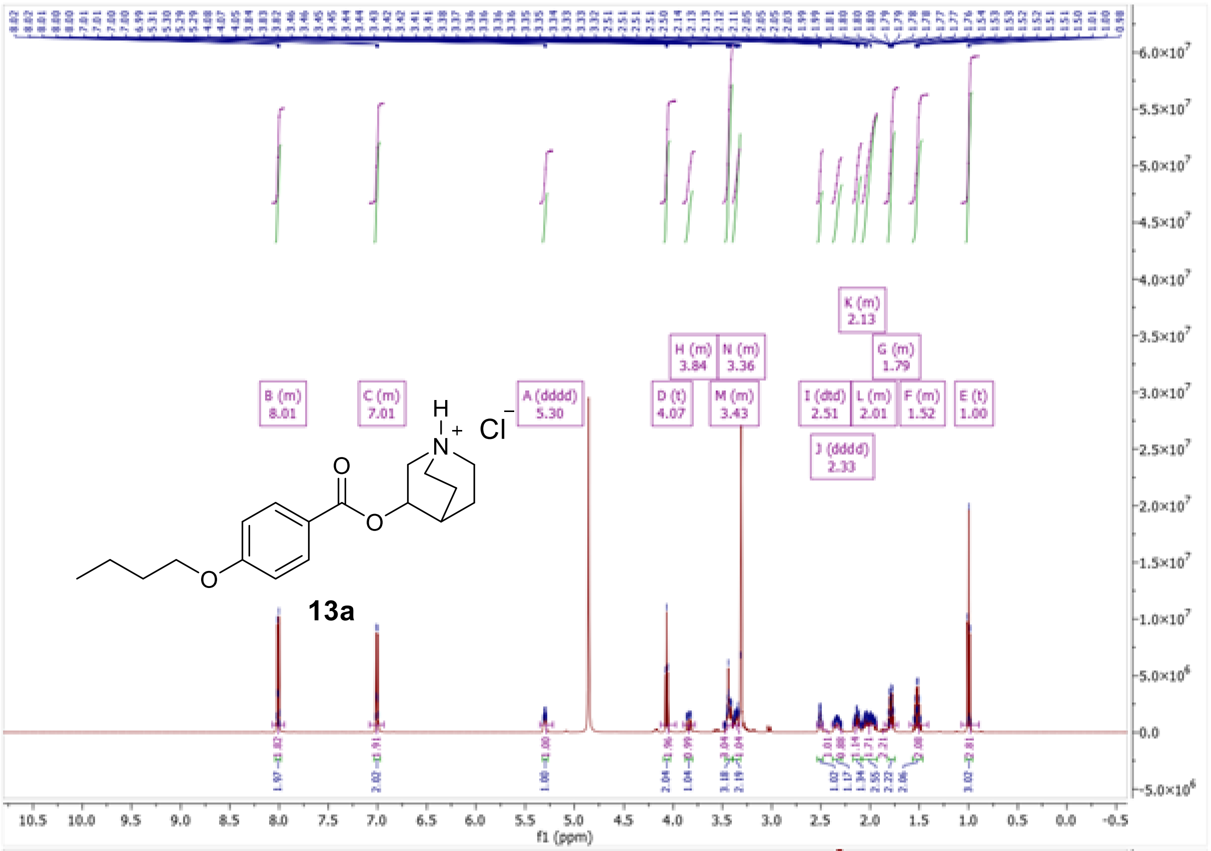

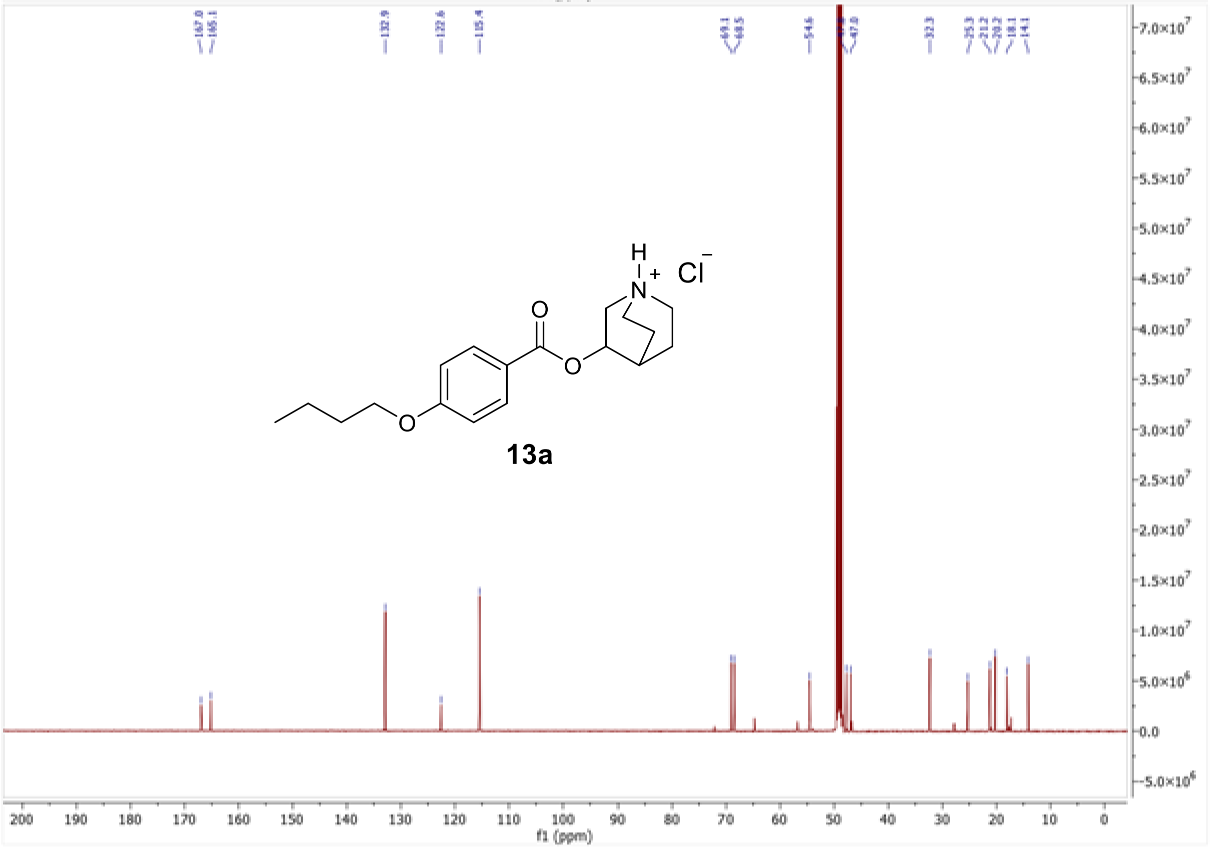

[1]H and [13]C NMR spectra of 1-methylquinuclidin-3-yl 4-butoxybenzoate iodide (**13b**)

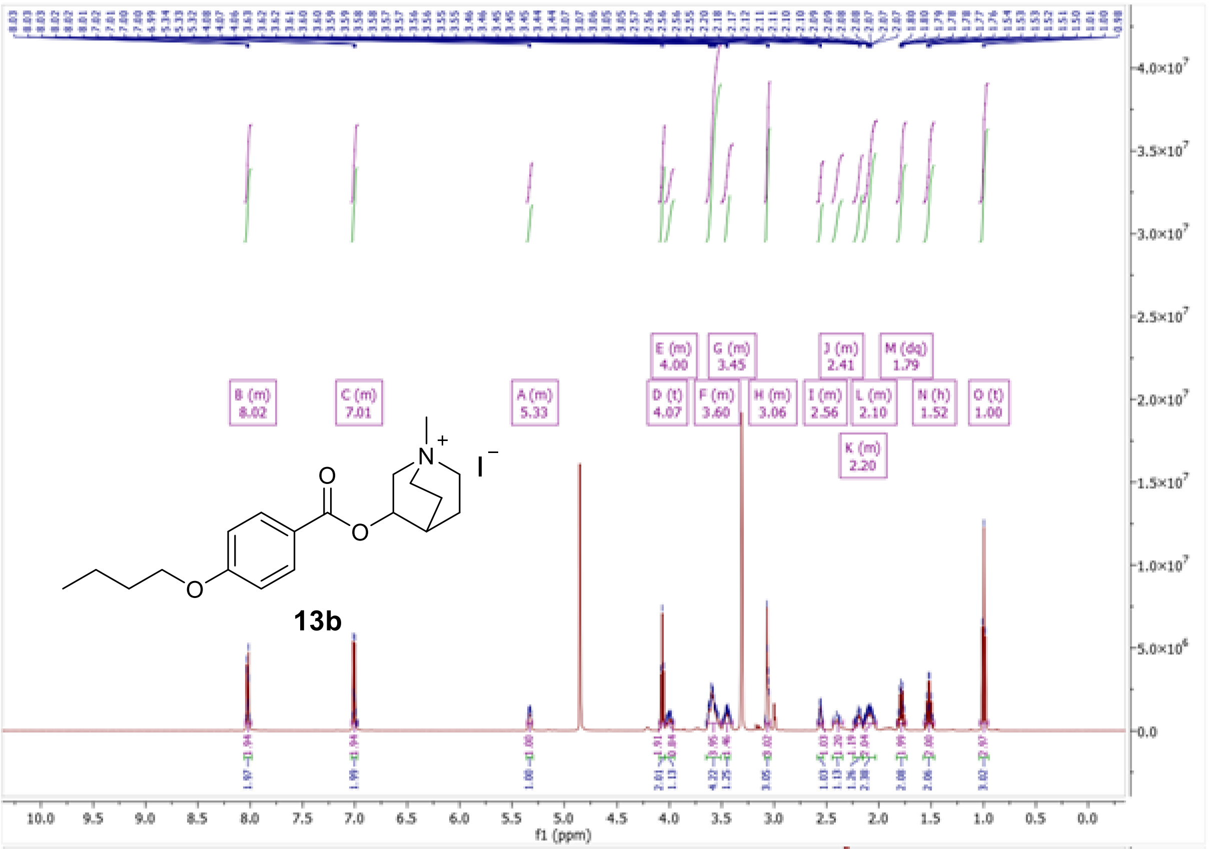

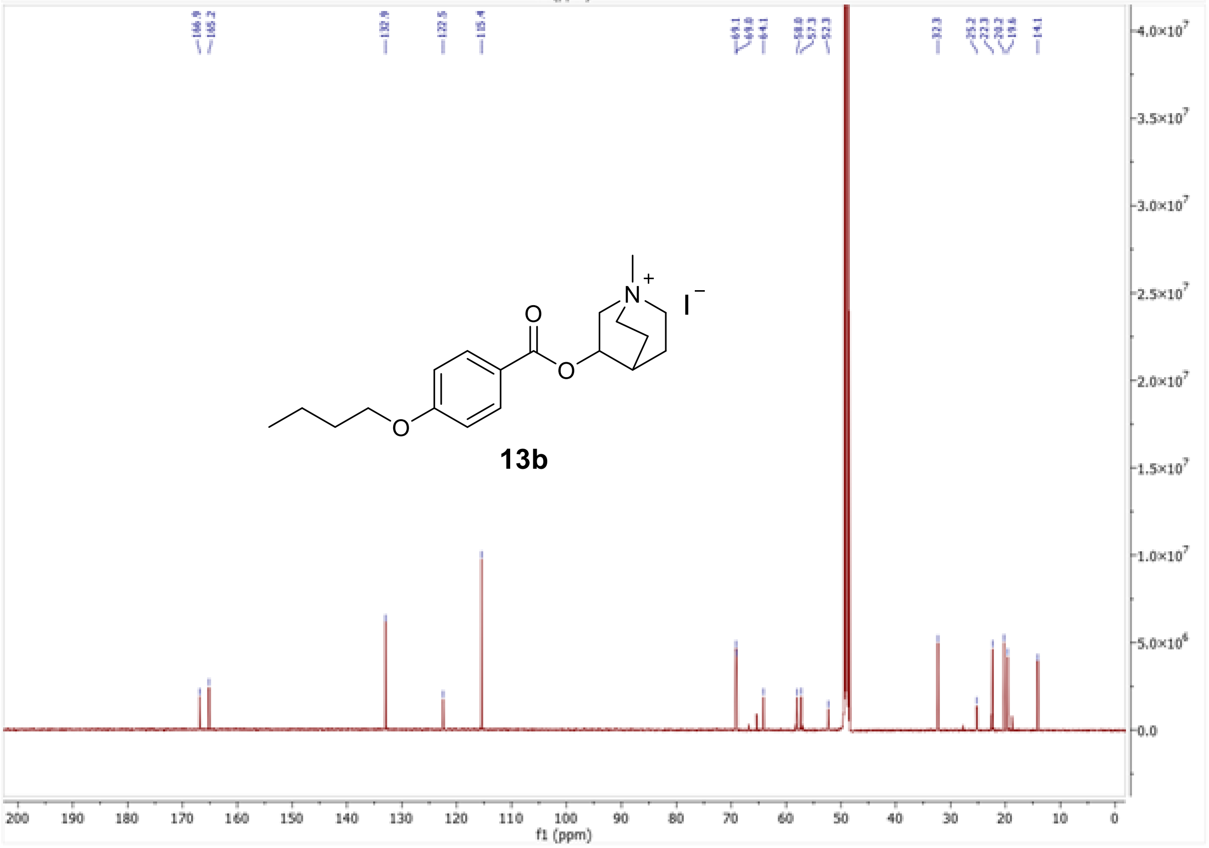

[1]H and [13]C NMR spectra of Quinuclidin-3-yl 4-methoxybenzoate hydrochloride (**14a**)

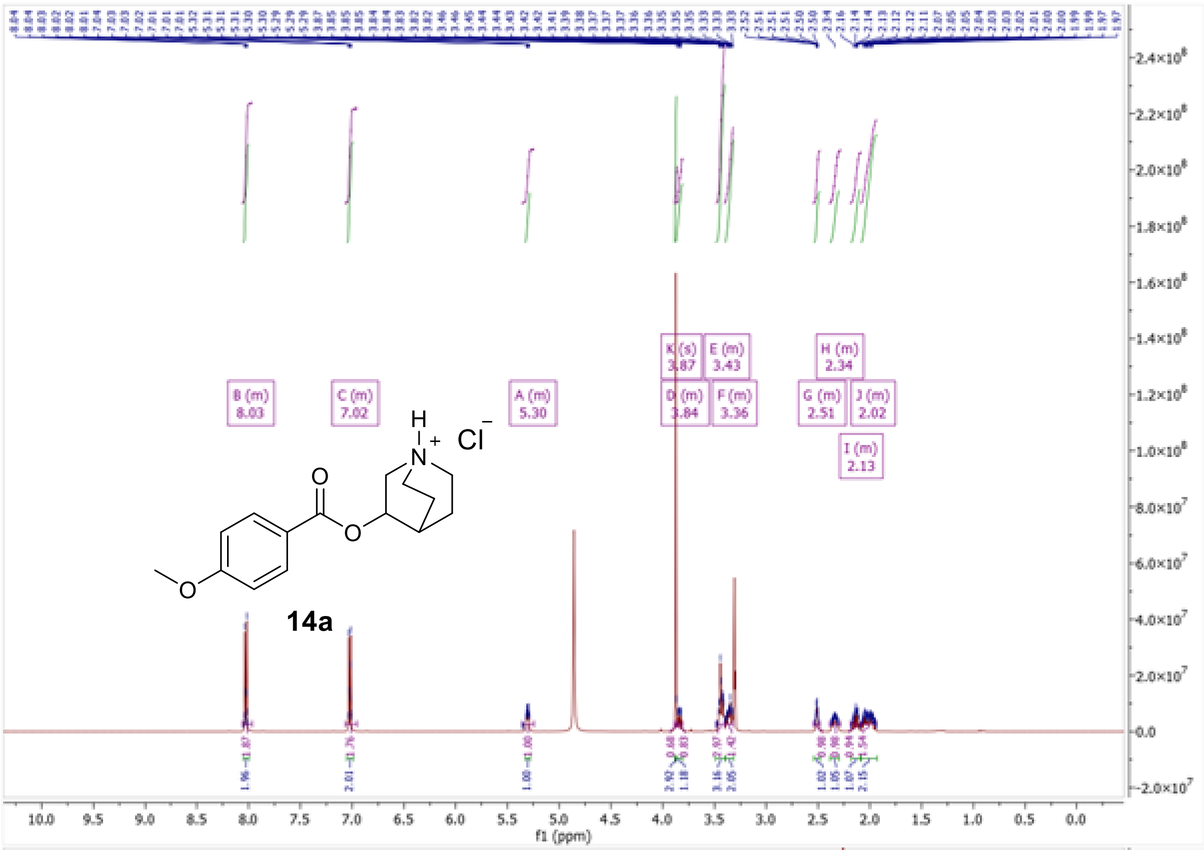

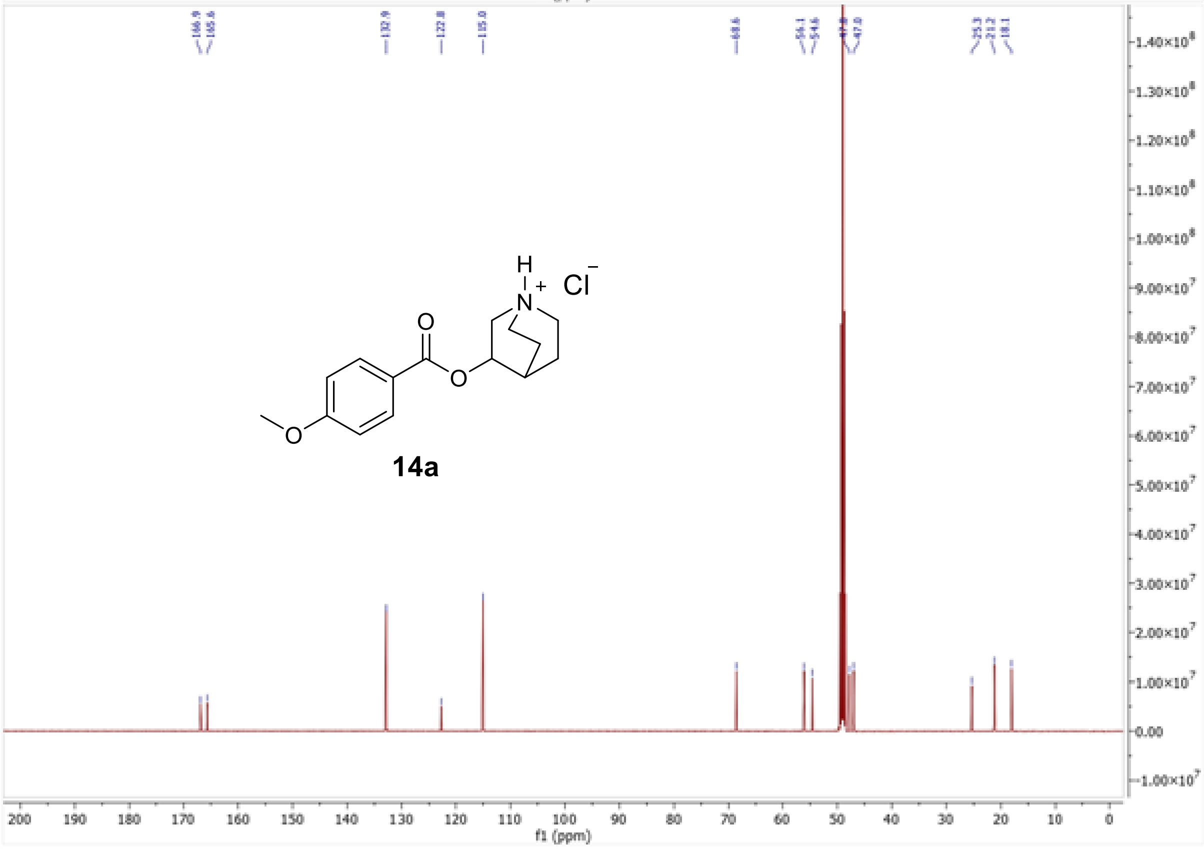

[1]H and [13]C NMR spectra of 1-methylquinuclidin-3-yl 4-methoxybenzoate iodide (**14b**)

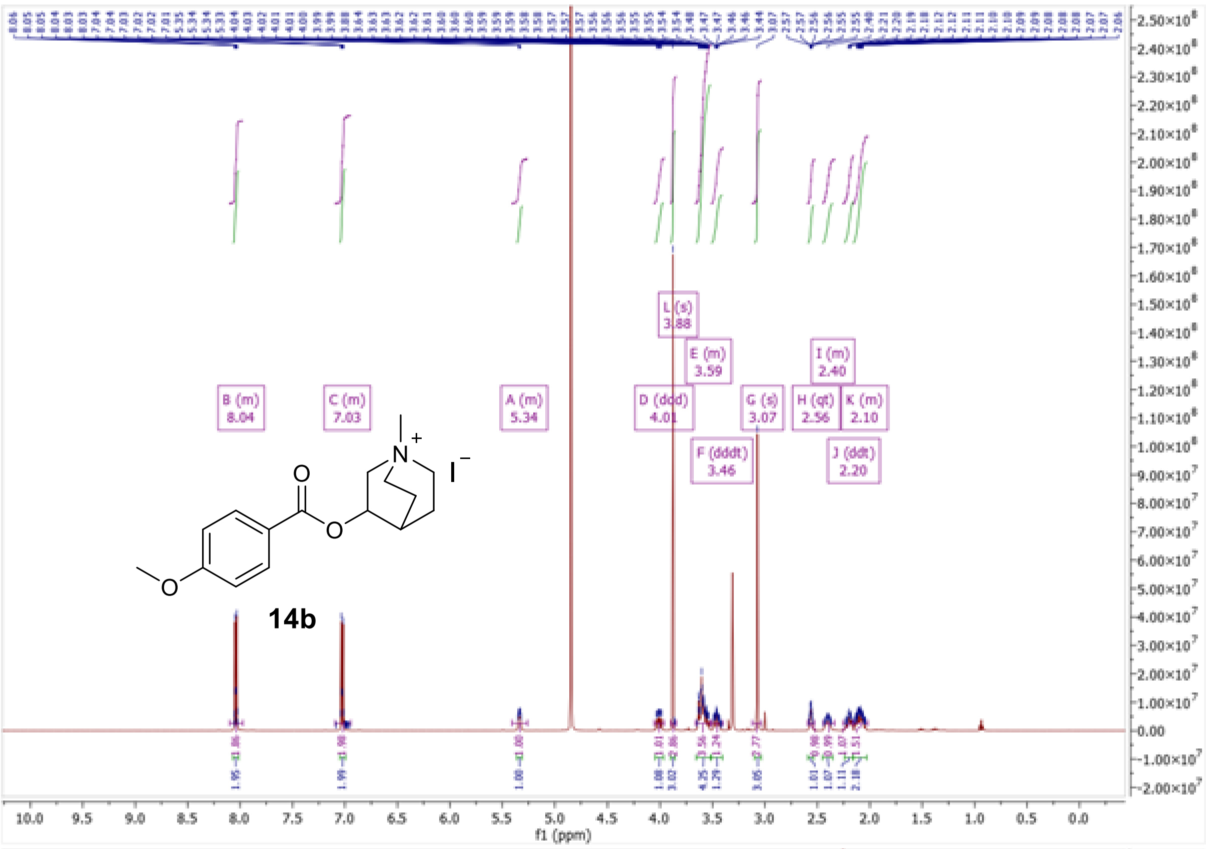

B (m)
8.04
C (m)
7.03
A (m)
5.34
D (ddd)
4.01
L (s)
3.88
E (m)
3.59
F (dddt)
3.46
G (s)
3.07
H (qt)
2.56
I (m)
2.40
J (ddt)
2.20
K (m)
2.10
14b
f1 (ppm)

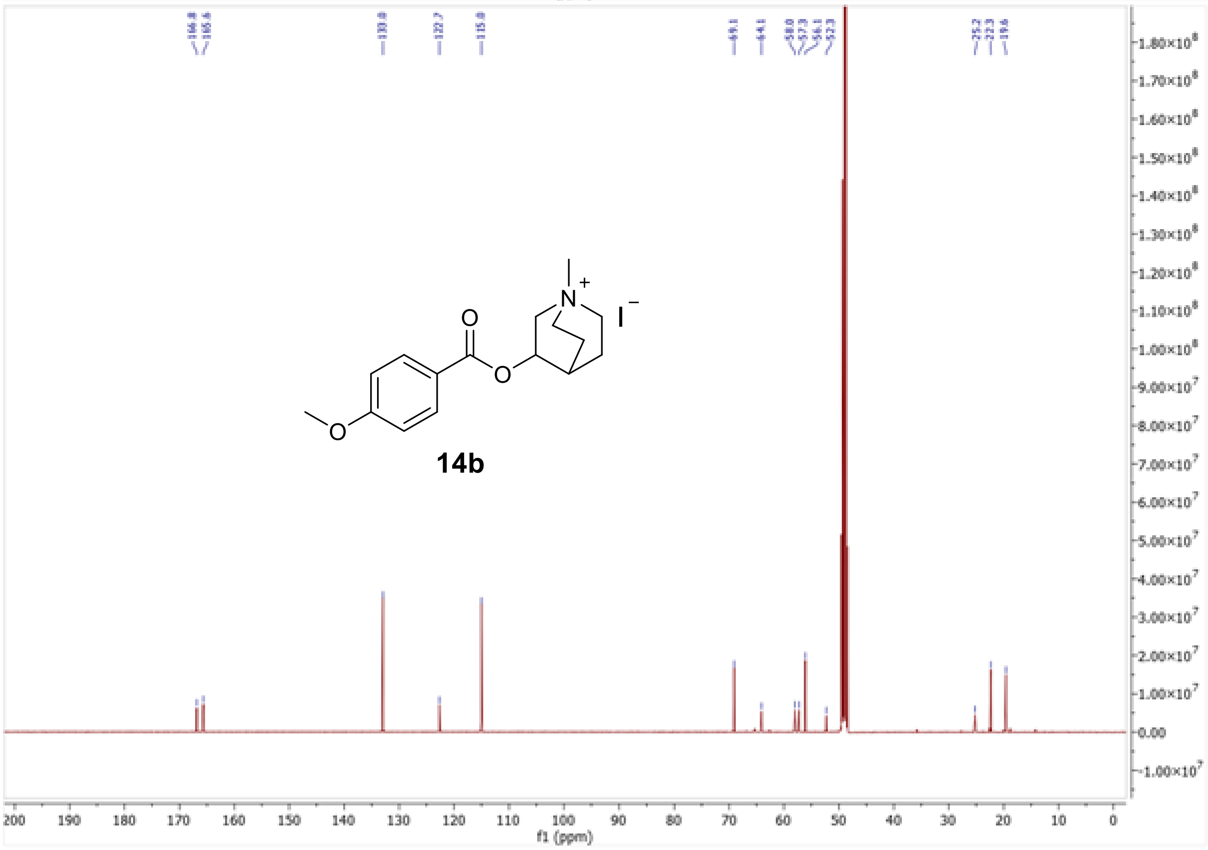

166.8
165.6
133.0
122.7
115.0
69.1
64.1
58.0
57.3
56.1
52.3
25.2
22.3
19.6
14b
f1 (ppm)

#### 3. HPLC Analysis and HRMS Records

High-performance liquid chromatography (HPLC) with mass spectrometry (MS) was used to determine high-resolution mass spectra (HRMS) and the purity of compounds.

*LC-MS Instrumentation*

The system used in this study was Dionex Ultimate 3000 UHPLC: RS LPG quaternary Pump, RS Column Compartment, RS Autosampler, Diode Array Detector, Chromeleon (version 7.2.9 build 11323) software with Q Exactive Plus Orbitrap mass spectrometer with Thermo Xcalibur (version 4.3.73.11) software (Thermo Fisher Scientific, Waltham, MA, USA). Settings of heated electrospray source were: Spray voltage 3.5 kV, Capillary temperature: 260 °C, Sheath gas: 50 arbitrary units, Auxiliary gas: 12.5 arbitrary units, Spray gas: 2.5 arbitrary units, Probe heater temperature: 300 °C, Max spray current: 100 µA, S-lens RF Level: 50.

*HRMS and purity determination*

High-resolution mass spectra and sample purities were obtained by HPLC gradient method with detection by UV and MS. A C18 column (Kinetex EVO C18, 2.1x50 mm, 1.7 µm, Phenomenex, Torrance, CA, USA) was used in this study. Mobile phase A was ultrapure water of ASTM I type (resistance 18.2 MΩ.cm at 25°C) prepared by Barnstead Smart2Pure 3 UV/UF apparatus (Thermo Fisher Scientific, Waltham, MA, USA) with 0.1 % (v/v) formic acid (LC-MS grade, VWR, Radnor, PA, USA); mobile phase B was acetonitrile (MS grade, VWR, Radnor, PA, USA) with 0.1 % (v/v) of formic acid. The flow was constant at 0.4 ml/min. Method started with 0.3 minutes of isocratic flow of 5 % B, then the gradient of B rose to 100 % B in 3 min and remained constant at 100 % B for 0.7 min. Then the composition went back to 5 % B and equilibrated for 3.5 min. Samples were dissolved in methanol (LC-MS grade, VWR, Radnor, PA, USA) at a concentration of 1 mg/mL and sample injection was 1 µL. Purity was determined from UV spectra measured at a wavelength of 254 nm. HRMS was determined in total ion current spectra from the mass spectrometer in positive mode.

HRMS spectrum of 2-(1,3-dioxo-2,3-dihydro-1H-isoindol-2-yl)ethyl 4-(pentyloxy)benzoate (**6a**)

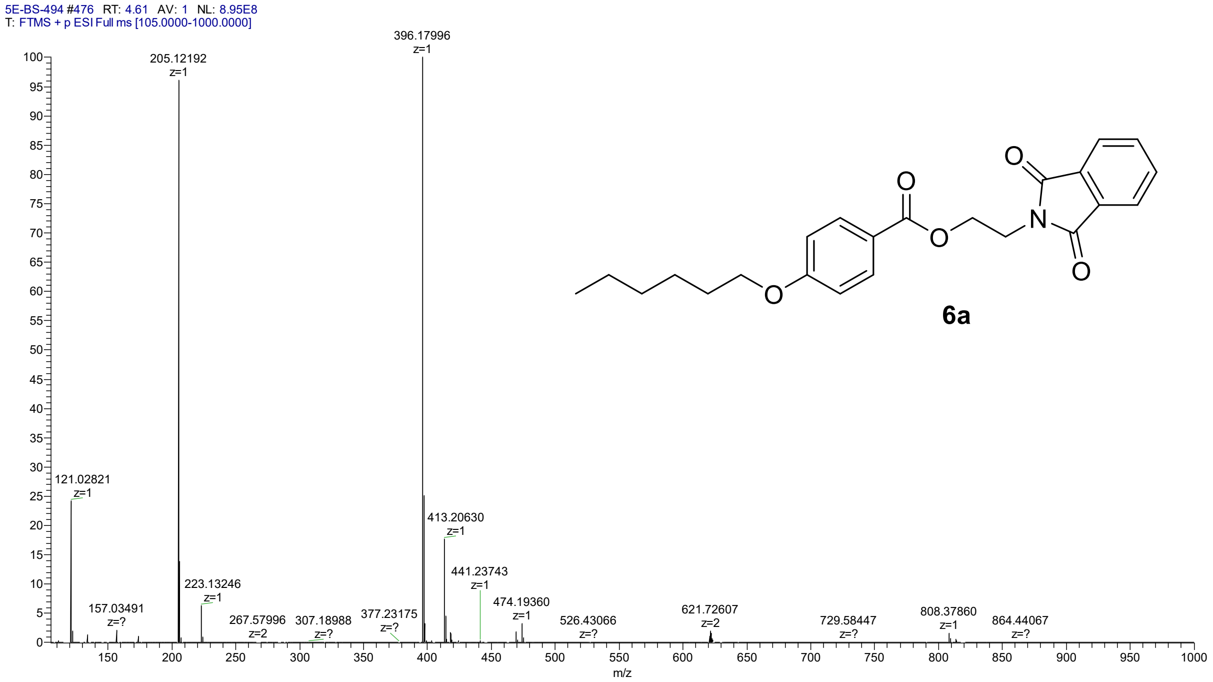

LC-UV chromatogram of 2-(1,3-dioxo-2,3-dihydro-1H-isoindol-2-yl)ethyl 4-(pentyloxy)benzoate (**6a**)

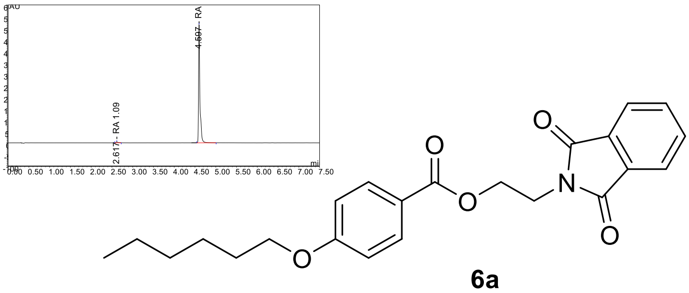

**6a**

HRMS spectrum of 1-{2-[4-(hexyloxy)benzoyloxy]ethyl}-1-azabicyclo[2.2.2]octan-1-ium (**6b**)

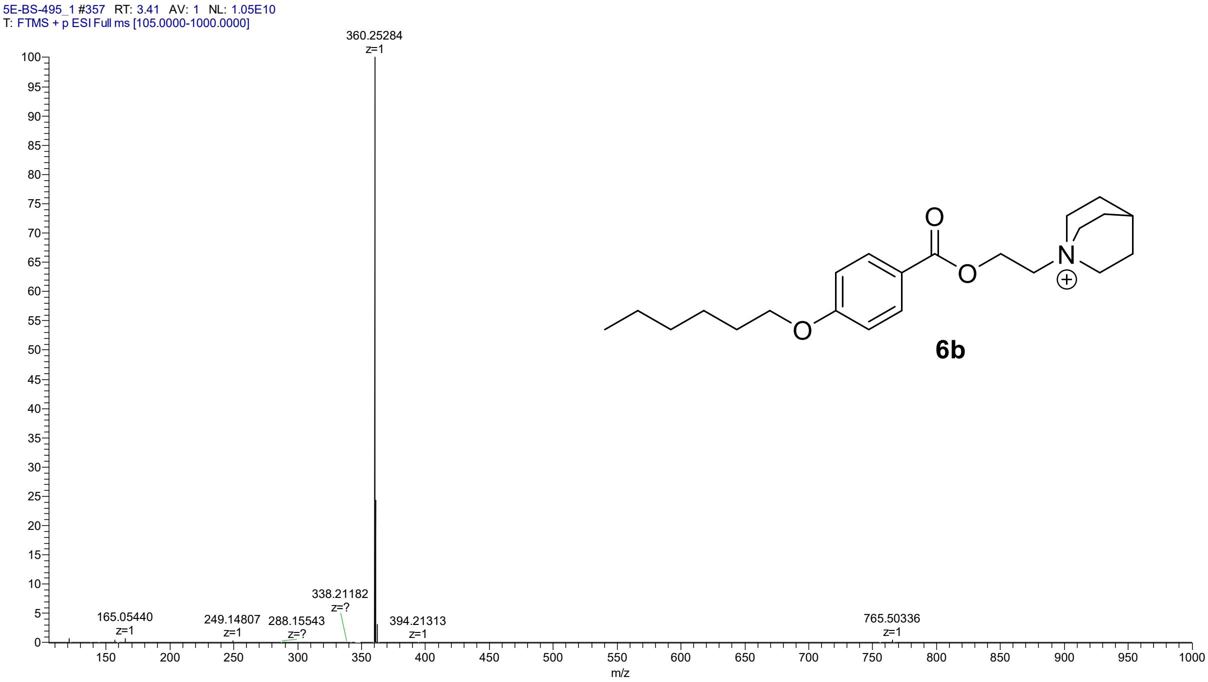

LC-UV chromatogram of 1-{2-[4-(hexyloxy)benzoyloxy]ethyl}-1-azabicyclo[2.2.2]octan-1-ium (**6b**)

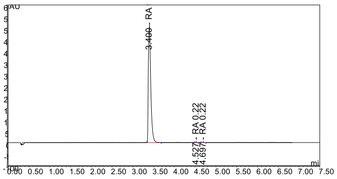

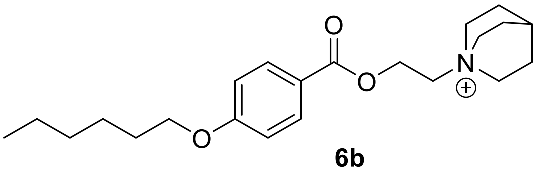

**6b**

HRMS spectrum of 2-(2,5-dioxopyrrolidin-1-yl)ethyl 4-(hexyloxy)benzoate (**6c**)

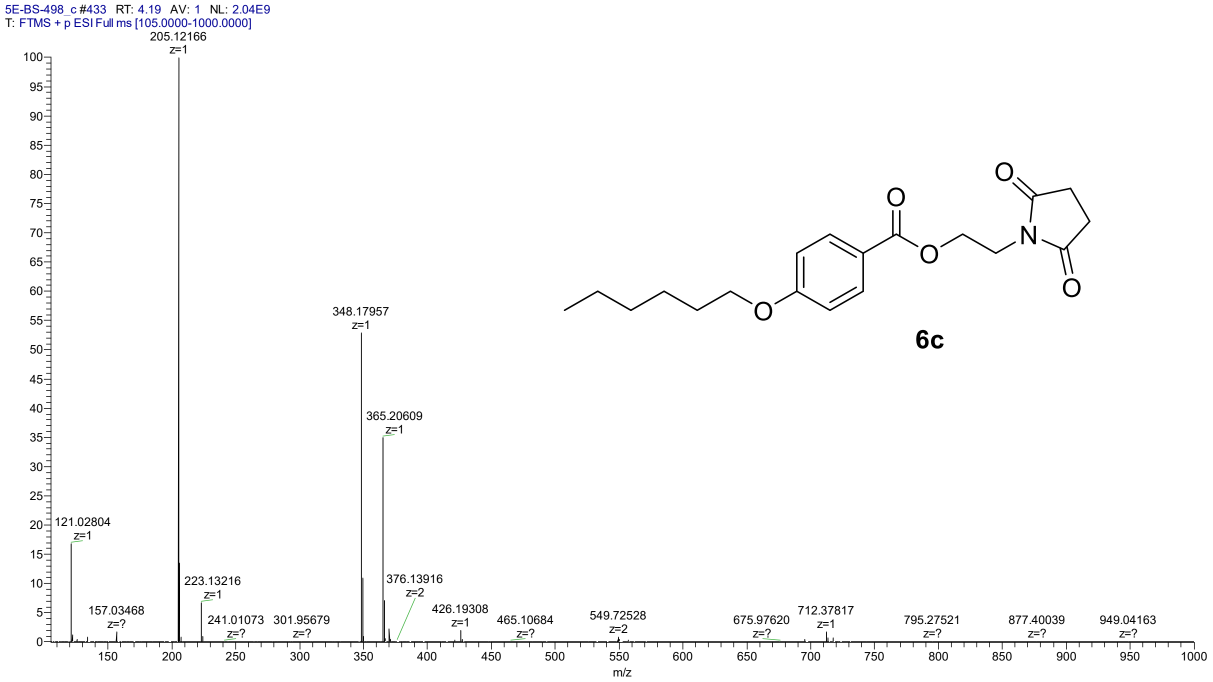

LC-UV chromatogram of 2-(2,5-dioxopyrrolidin-1-yl)ethyl 4-(hexyloxy)benzoate (**6c**)

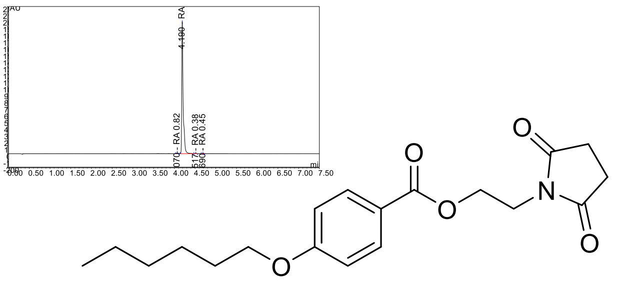

**6c**

HRMS spectrum of 2-{3,5-dioxo-10-oxa-4-azatricyclo[5.2.1.0$^{2,6}$]dec-8-en-4-yl}ethyl 4-(hexyloxy)benzoate (**6d**)

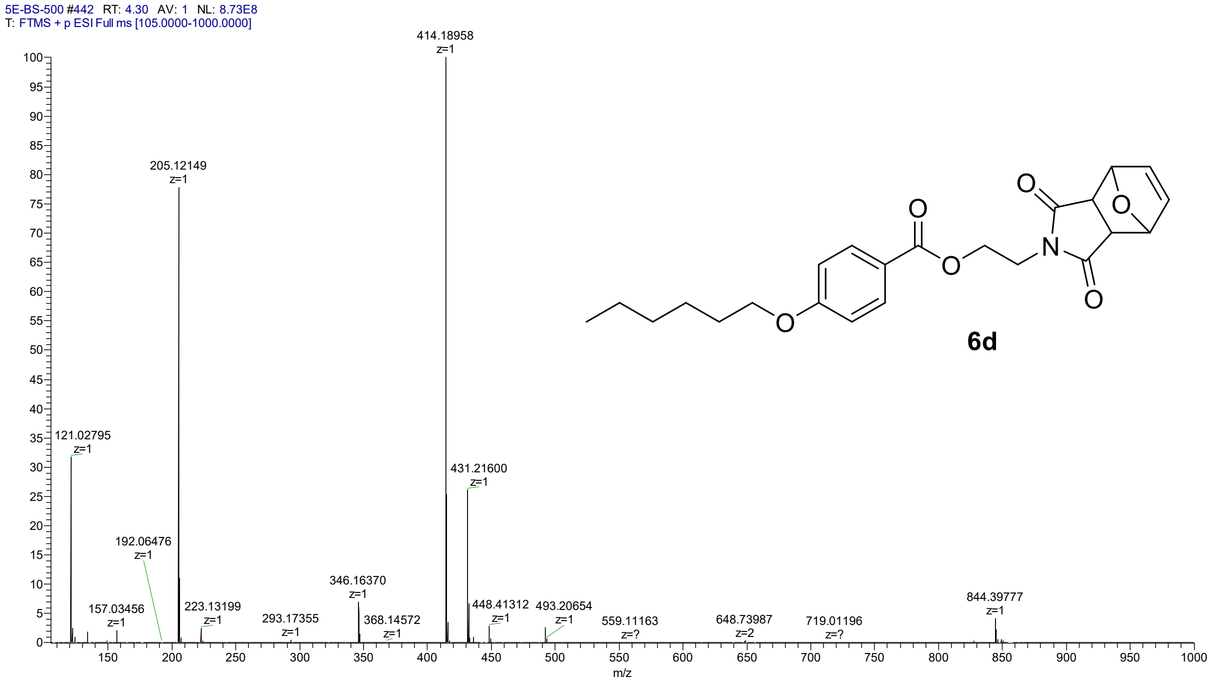

LC-UV chromatogram of 2-{3,5-dioxo-10-oxa-4-azatricyclo[5.2.1.0$^{2,6}$]dec-8-en-4-yl}ethyl 4-(hexyloxy)benzoate (**6d**)

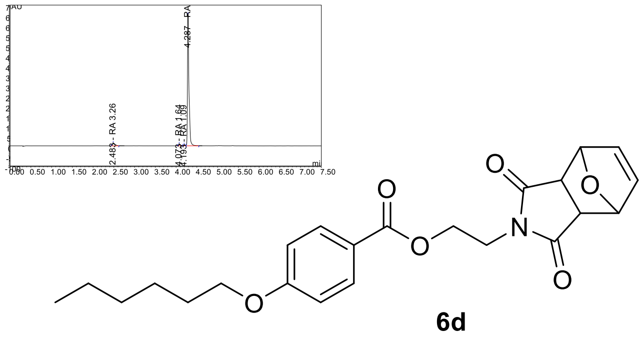

HRMS spectrum of 2-[(2*R*)-2-methylpiperidin-1-yl]ethyl 4-(hexyloxy)benzoate (**6e**)

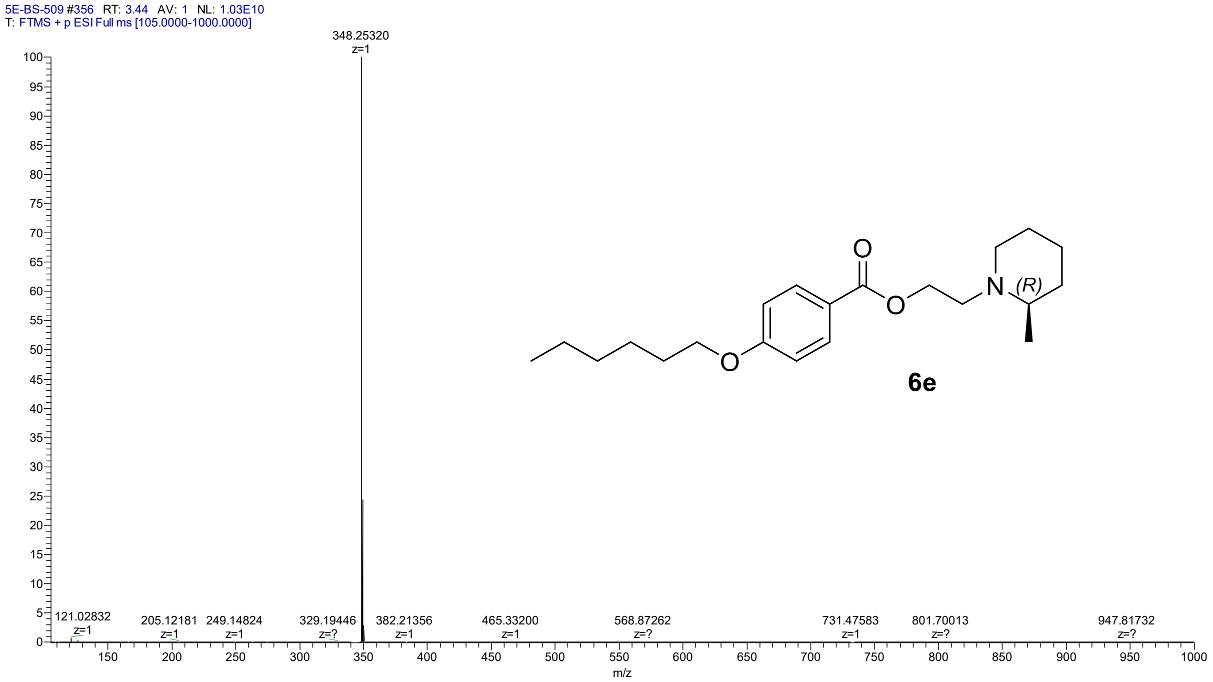

**6e**

LC-UV chromatogram of 2-[(2*R*)-2-methylpiperidin-1-yl]ethyl 4-(hexyloxy)benzoate (**6e**)

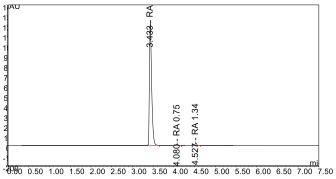

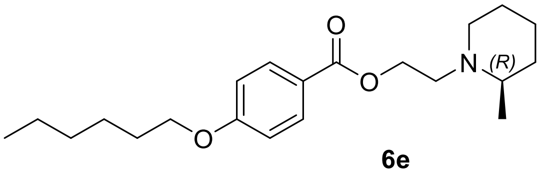

**6e**

HRMS spectrum of 2-[(2*S*)-2-methylpiperidin-1-yl]ethyl 4-(hexyloxy)benzoate (**6f**)

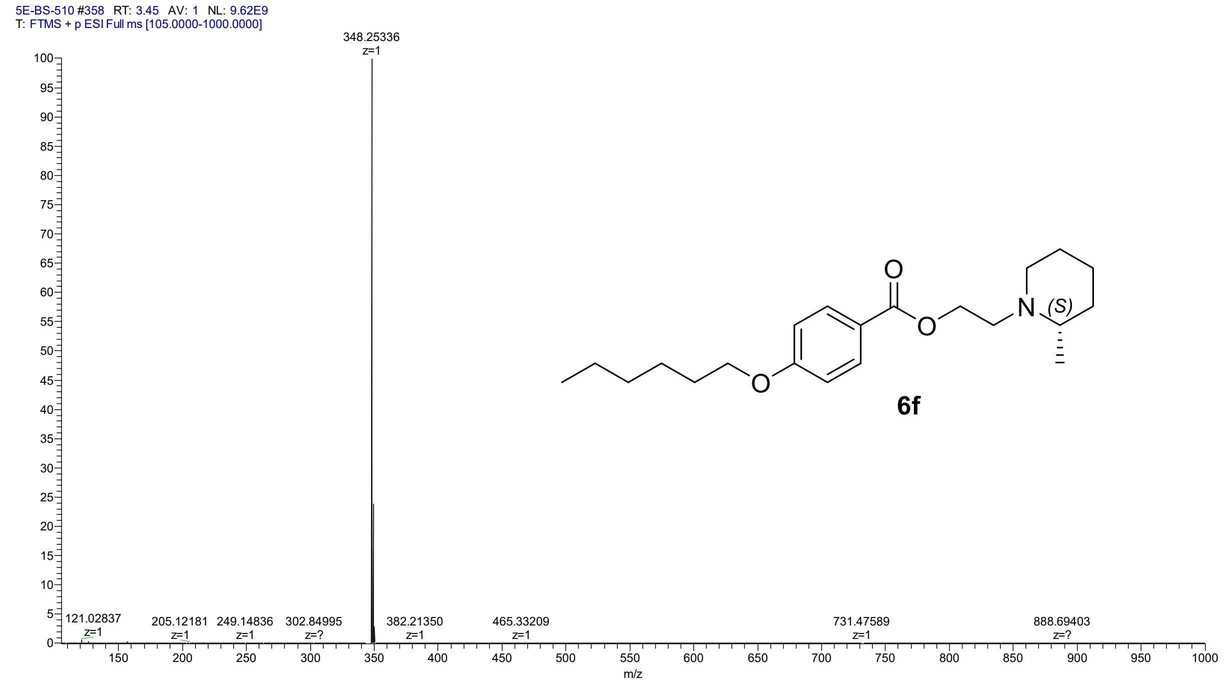

LC-UV

chromatogram of 2-[(2*S*)-2-methylpiperidin-1-yl]ethyl 4-(hexyloxy)benzoate (**6f**)

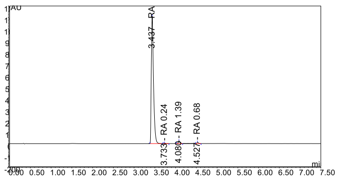

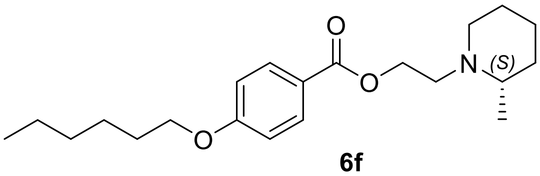

**6f**

HRMS spectrum of 2-{2-azabicyclo[2.2.2]octan-2-yl}ethyl 4-(hexyloxy)benzoate (**6g**)

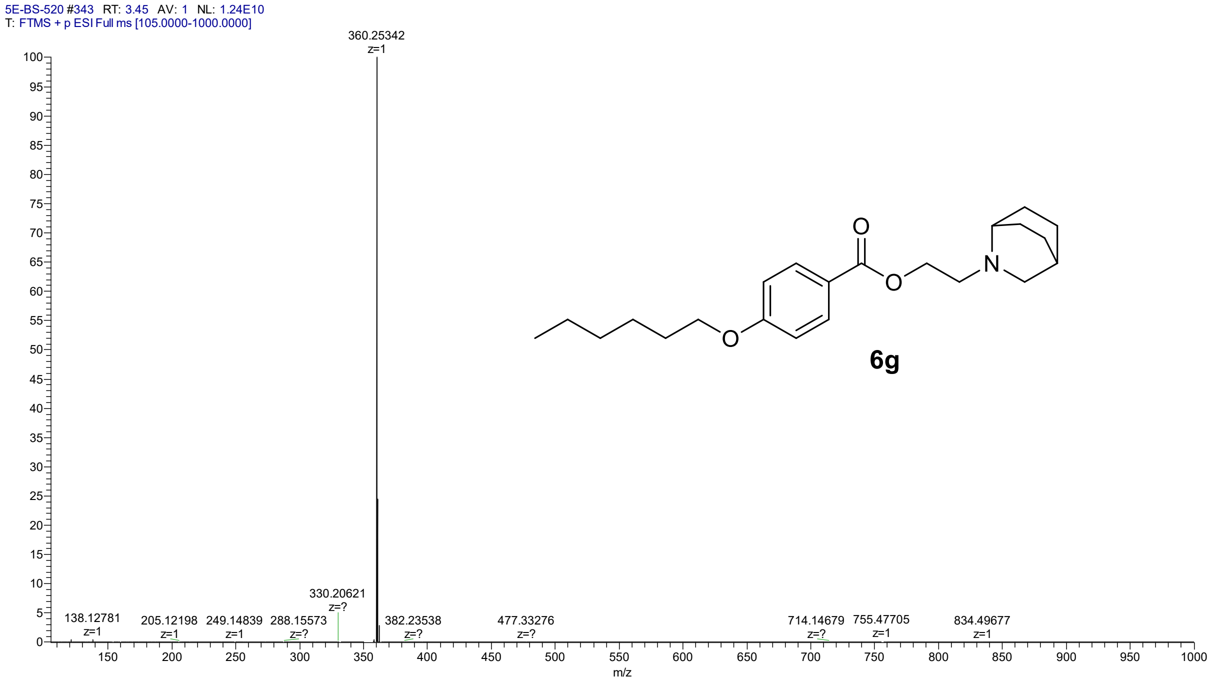

LC-UV chromatogram of 2-{2-azabicyclo[2.2.2]octan-2-yl}ethyl 4-(hexyloxy)benzoate (**6g**)

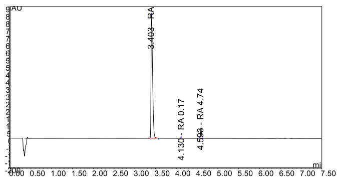

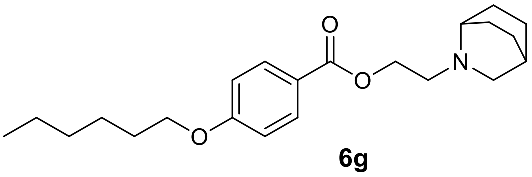

**6g**

#### HRMS spectrum of 2-(2,5-dihydro-1H-pyrrol-1-yl)ethyl 4-(hexyloxy)benzoate (**6h**)

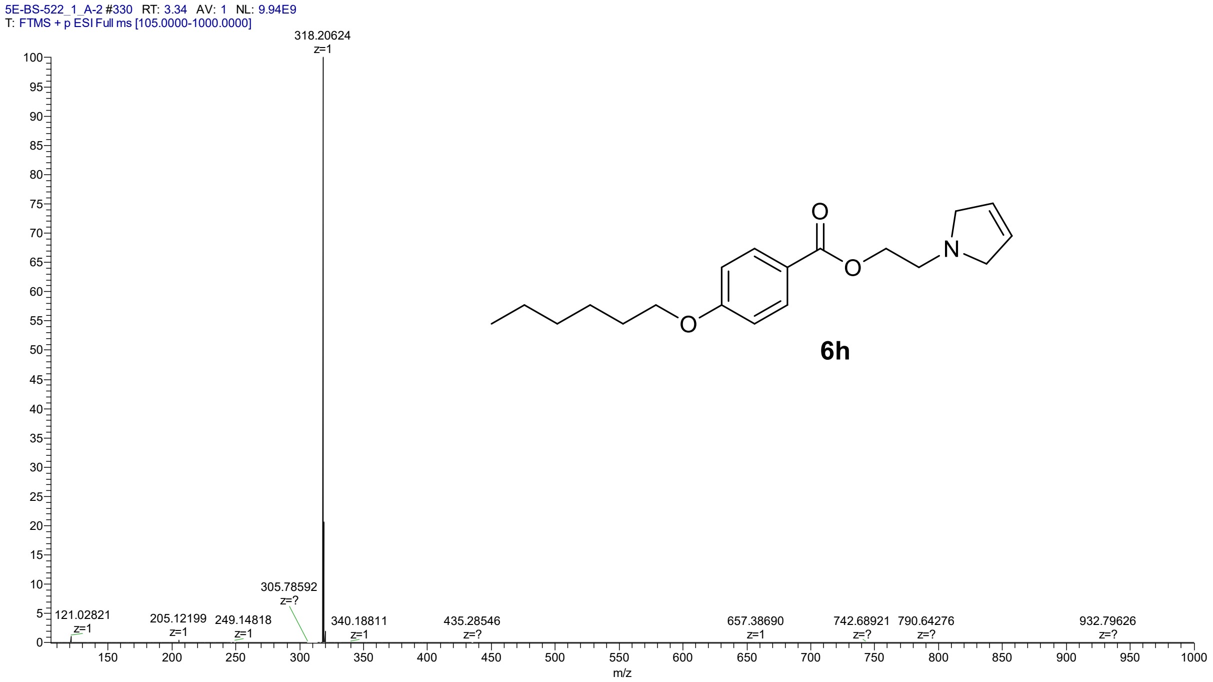

#### LC-UV chromatogram of 2-(2,5-dihydro-1H-pyrrol-1-yl)ethyl 4-(hexyloxy)benzoate (**6h)**

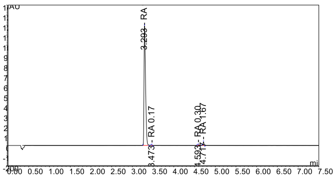

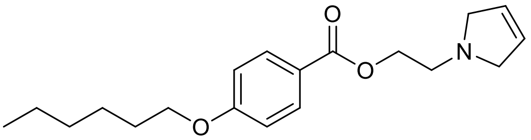

**6h**

HRMS spectrum of 2-(pyrrolidin-1-yl)ethyl 4-(hexyloxy)benzoate (**6j**)

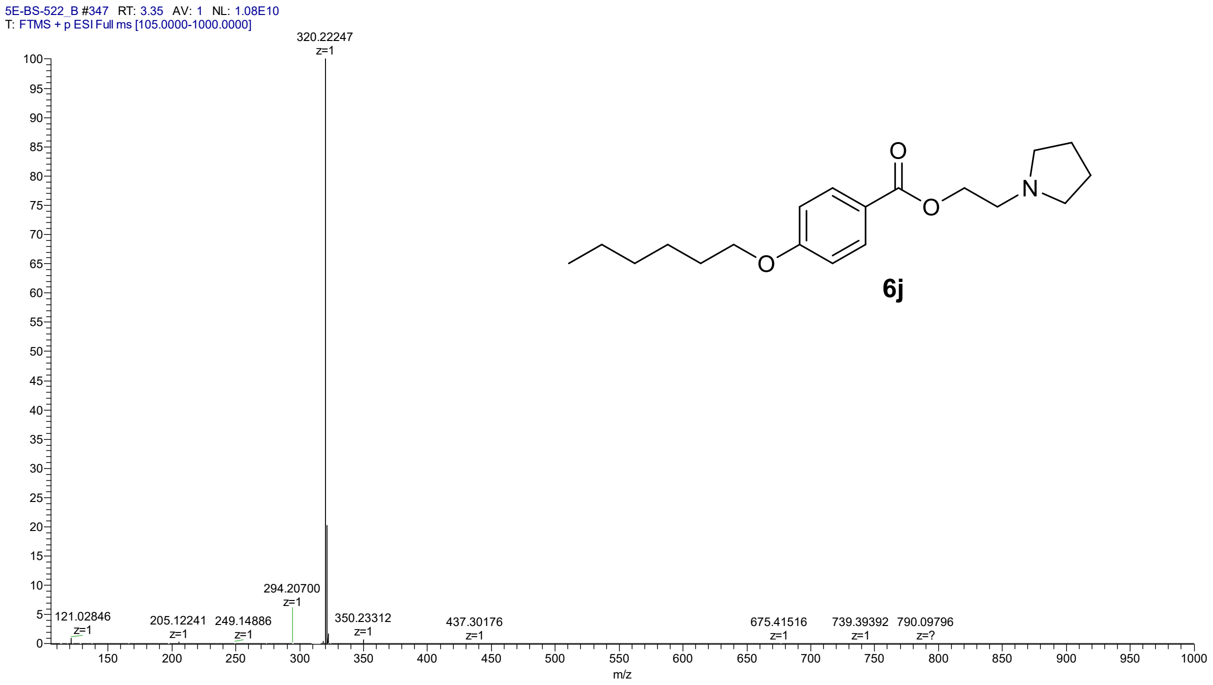

LC-UV chromatogram of 2-(pyrrolidin-1-yl)ethyl 4-(hexyloxy)benzoate (**6j**)

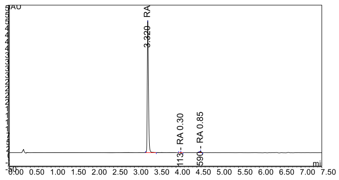

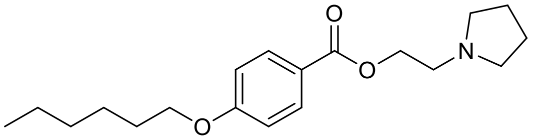

**6j**

HRMS spectrum of 2-(2,3-dihydro-1H-isoindol-2-yl)ethyl 4-(hexyloxy)benzoate (**6k**)

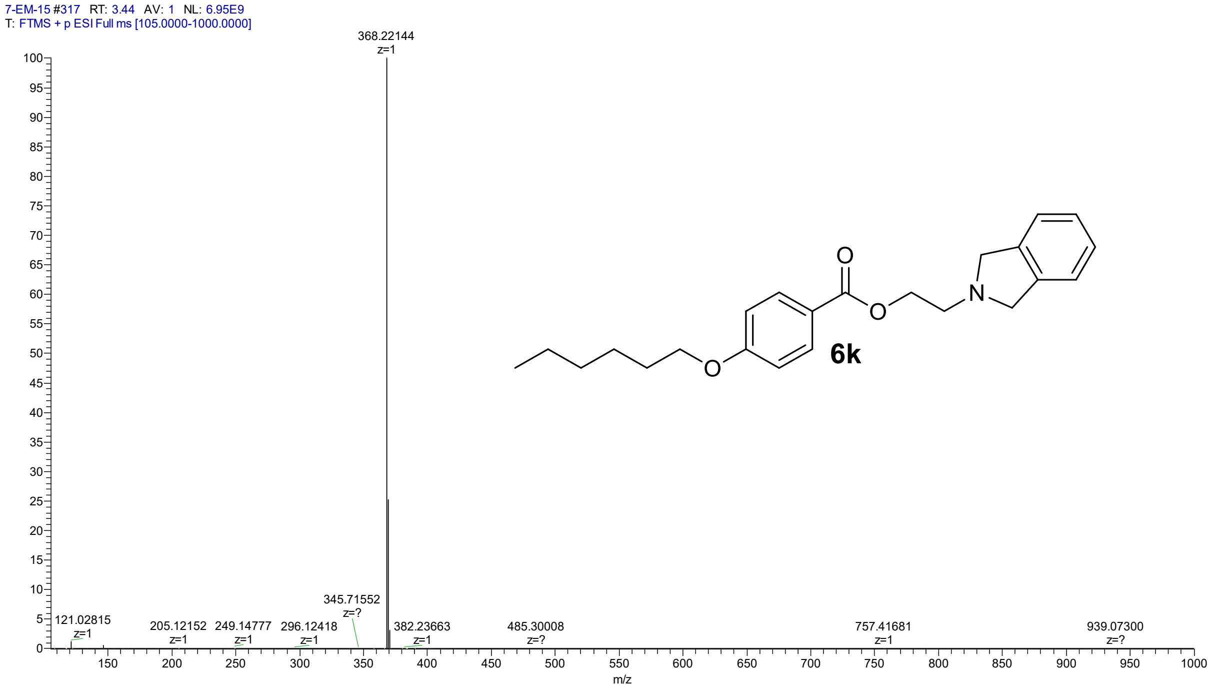

LC-UV chromatogram of 2-(2,3-dihydro-1H-isoindol-2-yl)ethyl 4-(hexyloxy)benzoate (**6k**)

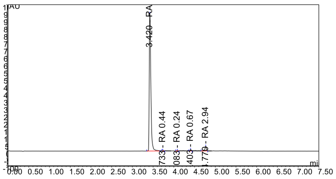

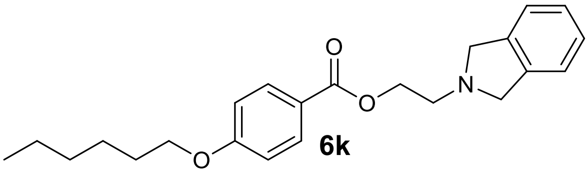

HMRS spectrum of 2-(1,2,3,4-tetrahydroquinolin-1-yl)ethyl 4-(hexyloxy)benzoate (**6l**)

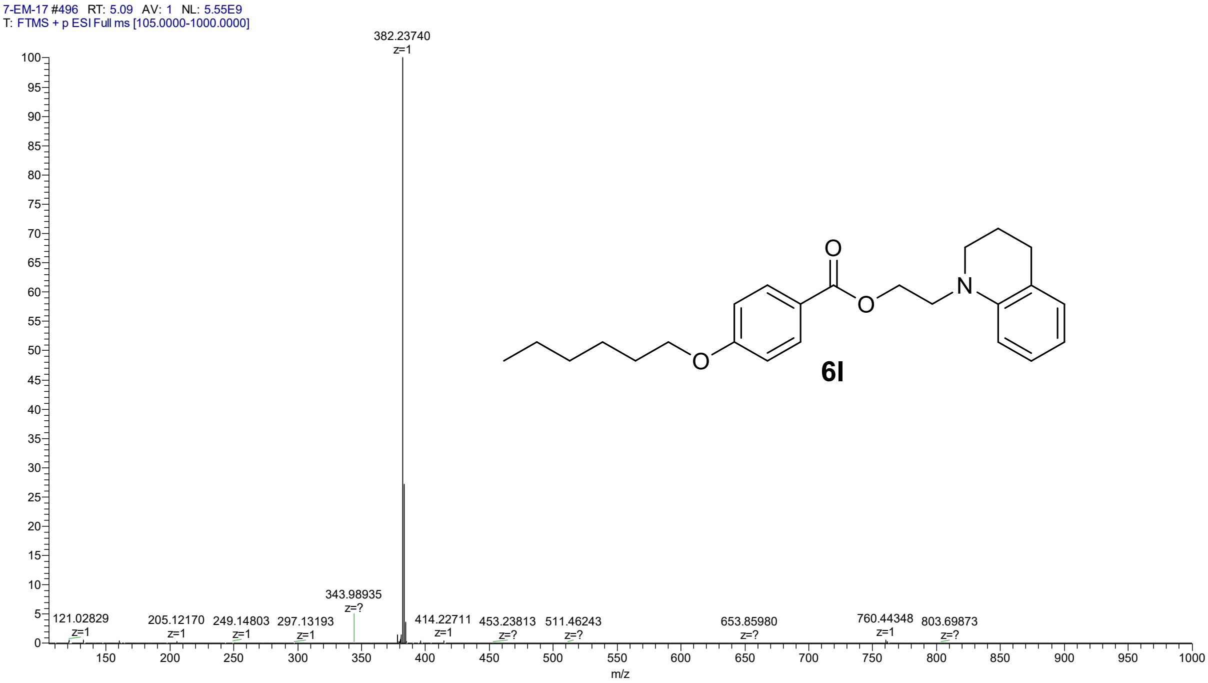

LC-UV chromatogram of 2-(1,2,3,4-tetrahydroquinolin-1-yl)ethyl 4-(hexyloxy)benzoate (**6l**)

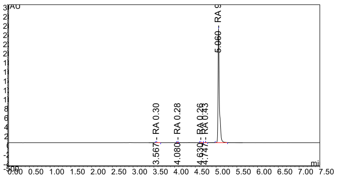

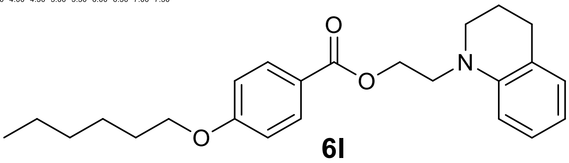

**6l**

HRMS spectrum of (1-methylpiperidin-2-yl)methyl 4-(hexyloxy)benzoate (**6m**)

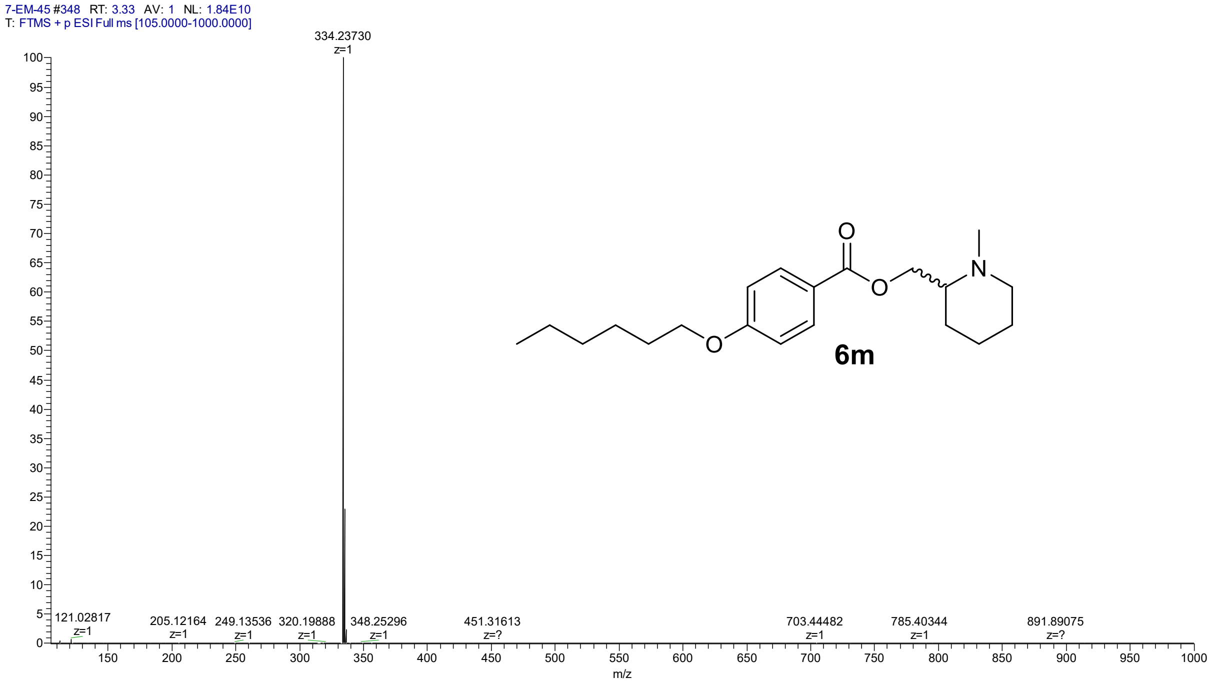

LC-UV chromatogram of (1-methylpiperidin-2-yl)methyl 4-(hexyloxy)benzoate (**6m**)

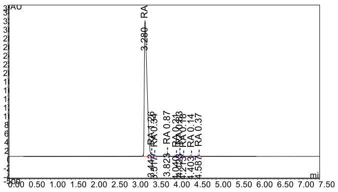

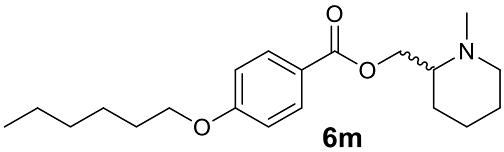

**6m**

HRMS spectrum of [(2*S*)-1-methylpiperidin-2-yl]methyl 4-(hexyloxy)benzoate (**6n**)

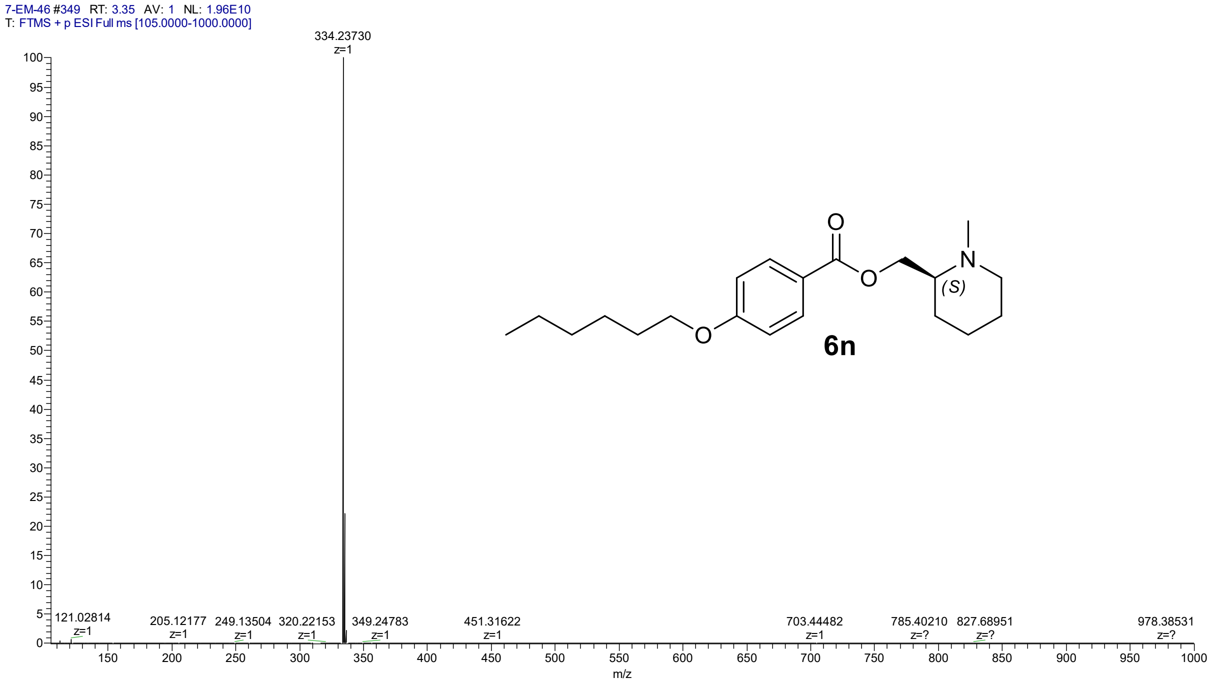

LC-UV chromatogram of [(2*S*)-1-methylpiperidin-2-yl]methyl 4-(hexyloxy)benzoate (**6n**)

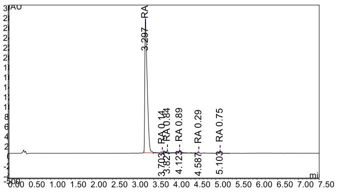

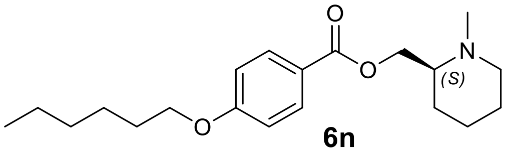

**6n**

HMRS spectrum of [(3*S*)-1-methylpiperidin-3-yl]methyl 4-(hexyloxy)benzoate (**6o**)

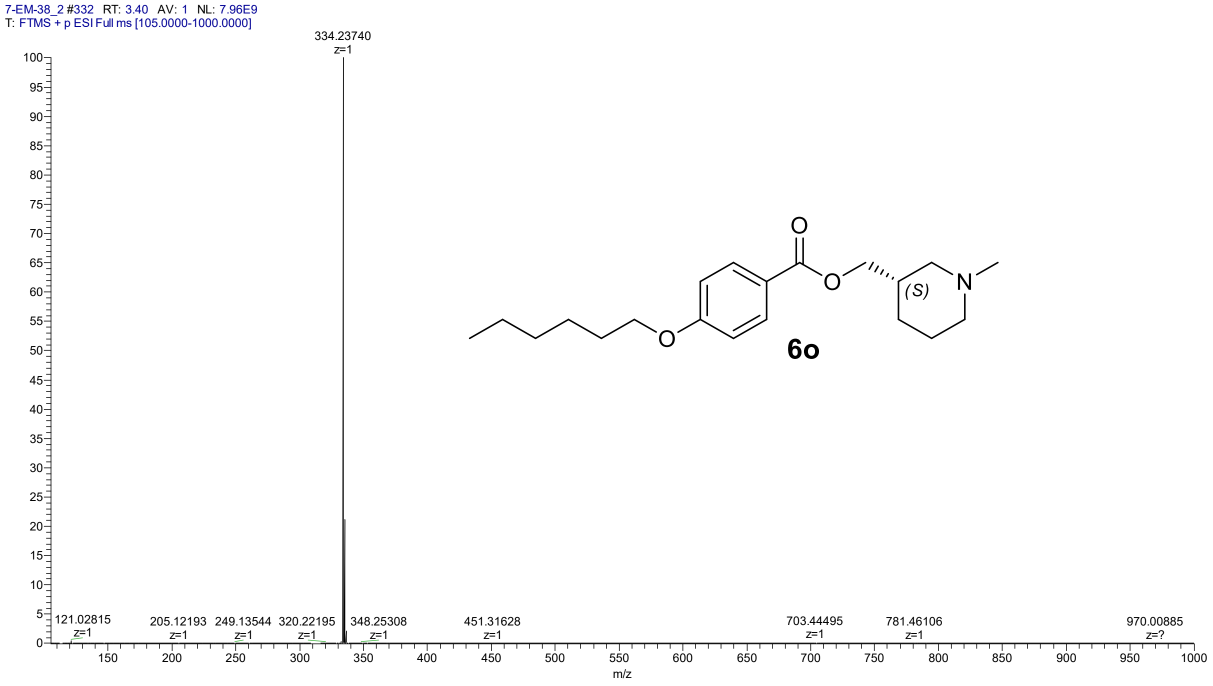

LC-UV chromatogram of [(3*S*)-1-methylpiperidin-3-yl]methyl 4-(hexyloxy)benzoate (**6o**)

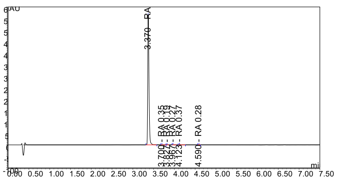

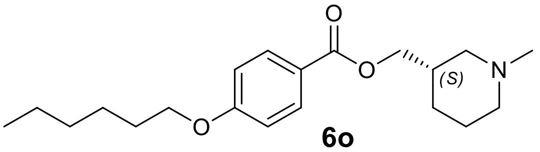

**6o**

HRMS spectrum of (1-methylpiperidin-3-yl)methyl 4-(hexyloxy)benzoate (**6p**)

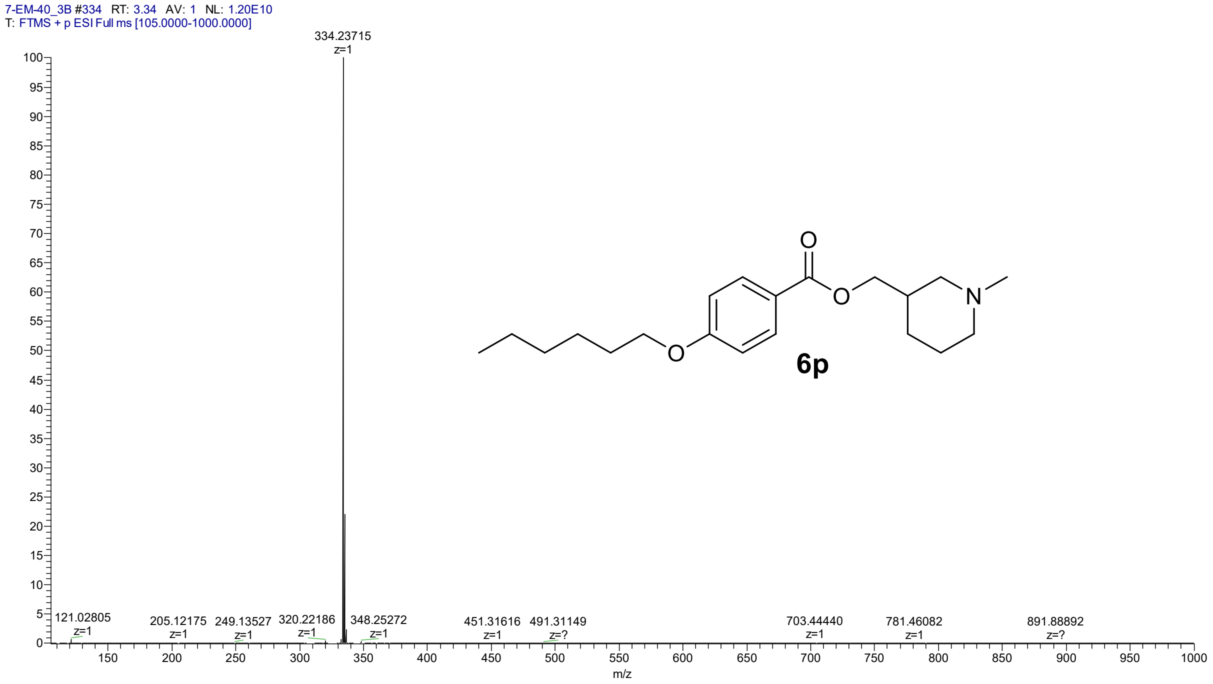

LC-UV chromatogram of (1-methylpiperidin-3-yl)methyl 4-(hexyloxy)benzoate (**6p**)

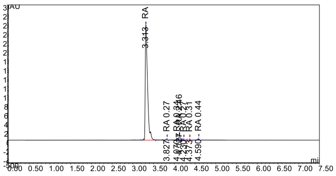

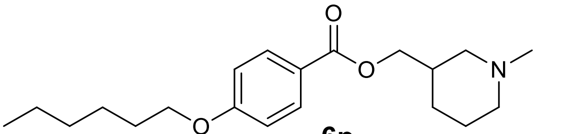

**6p**

#### HRMS spectrum of [(3*R*)-1-methylpiperidin-3-yl]methyl 4-(hexyloxy)benzoate (**6r**)

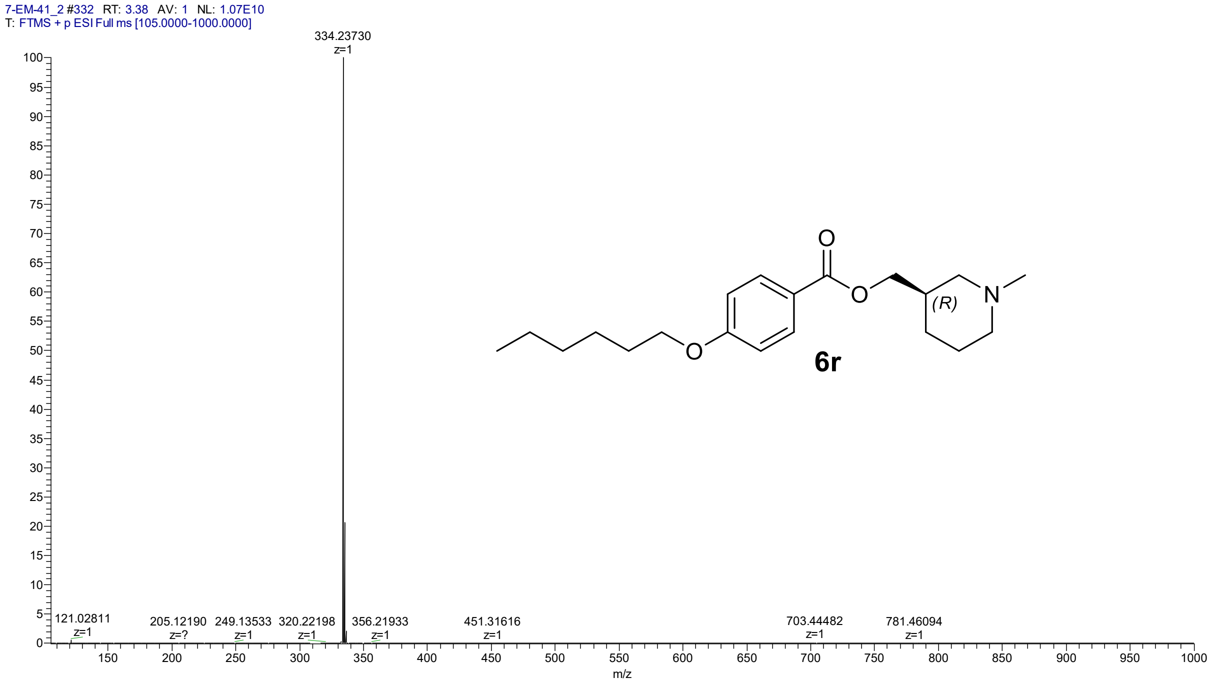

#### LC-UV chromatogram of [(3*R*)-1-methylpiperidin-3-yl]methyl 4-(hexyloxy)benzoate (**6r**)

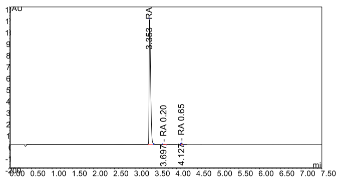

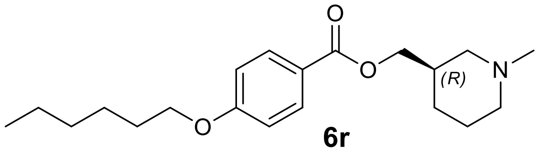

**6r**

HMRS spectrum of (1-methylpiperidin-4-yl)methyl 4-(hexyloxy)benzoate (**6s**)

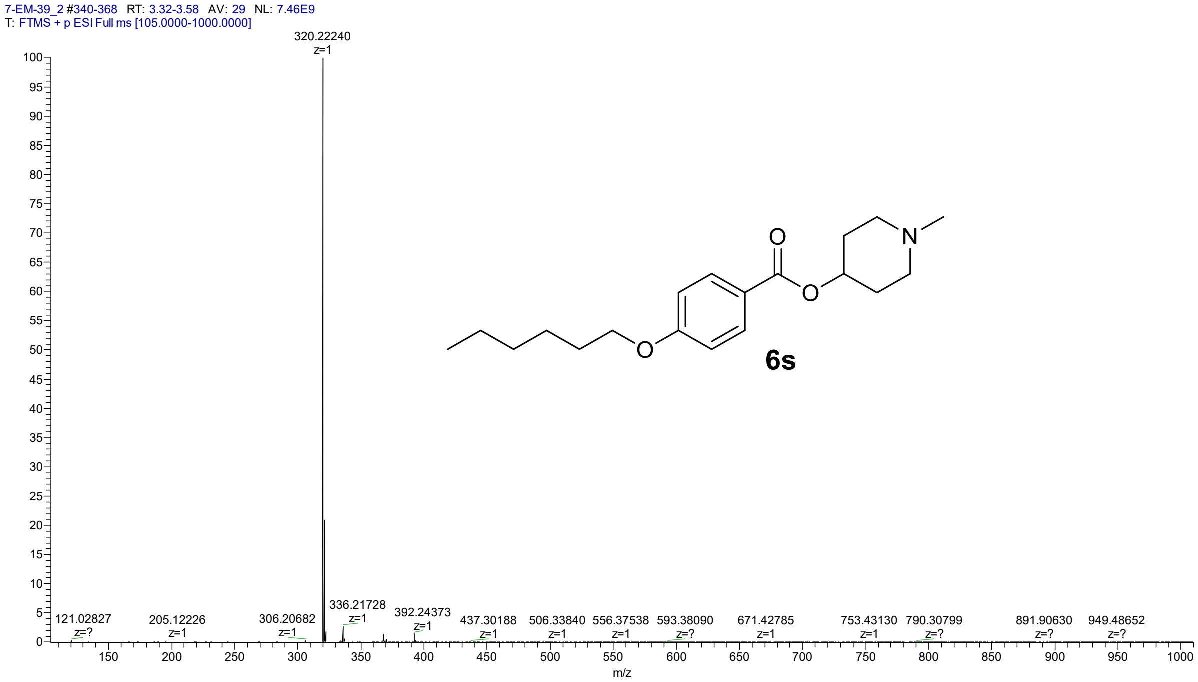

LC-UV chromatrogram of (1-methylpiperidin-4-yl)methyl 4-(hexyloxy)benzoate (**6s**)

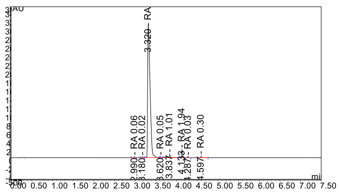

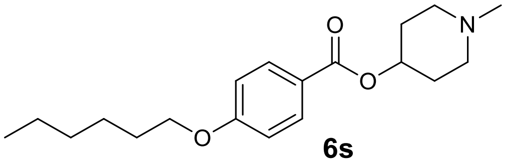

**6s**

HRMS spectrum of (1-methylpyrrolidin-2-yl)methyl 4-(hexyloxy)benzoate (**6t**)

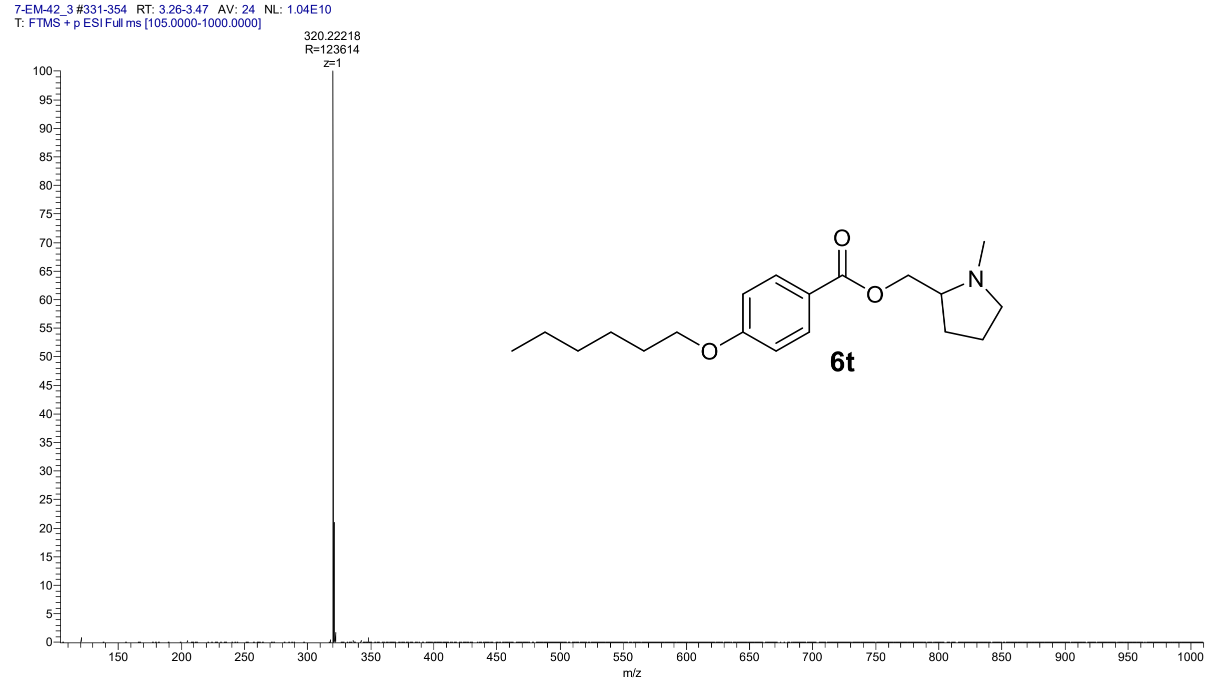

LC-UV chromatogram of (1-methylpyrrolidin-2-yl)methyl 4-(hexyloxy)benzoate (**6t**)

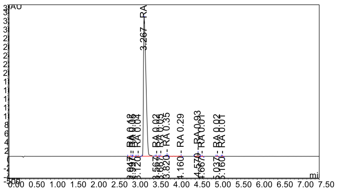

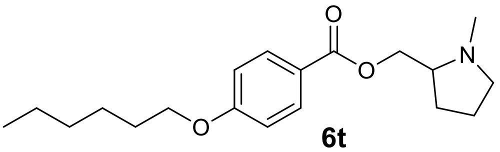

**6t**

HMRS spectrum of [(2*S*)-1-methylpyrrolidin-2-yl]methyl 4-(hexyloxy)benzoate (**6w**)

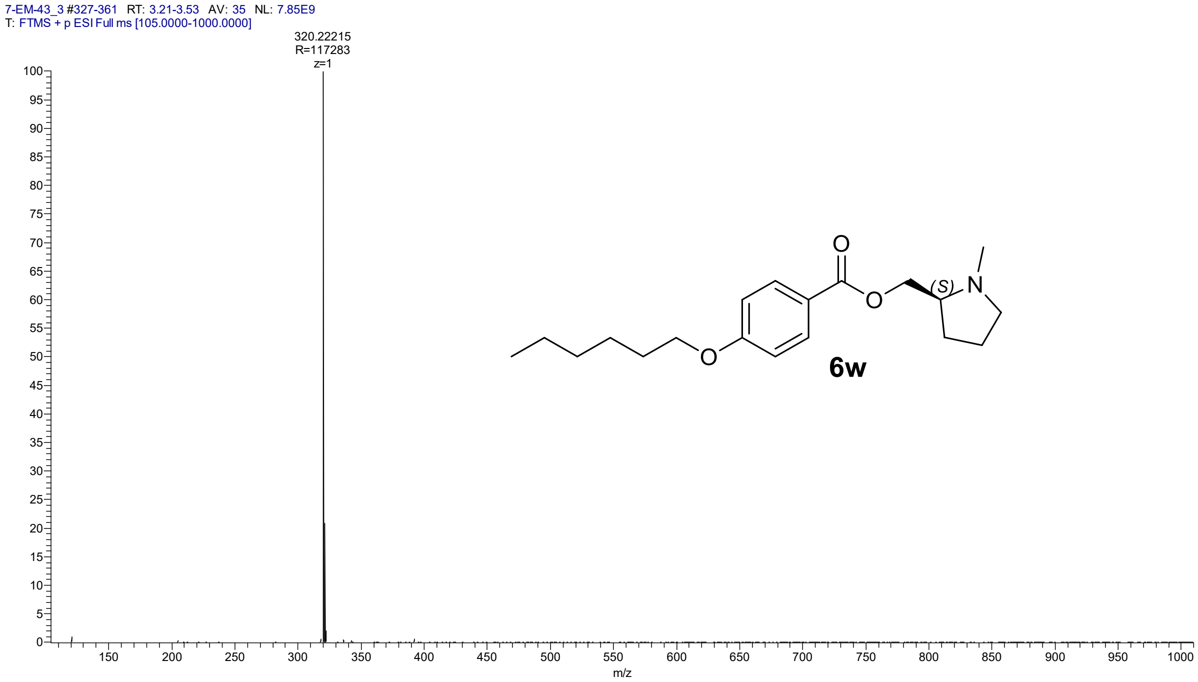

LC-UV chromatogram of [(2*S*)-1-methylpyrrolidin-2-yl]methyl 4-(hexyloxy)benzoate (**6w**)

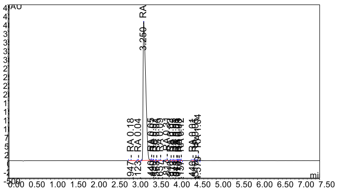

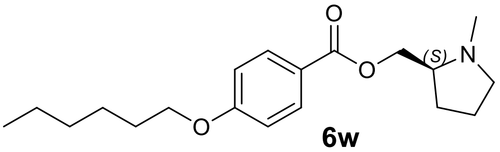

**6w**

HMRS spectrum of [(2*R*)-1-methylpyrrolidin-2-yl]methyl 4-(hexyloxy)benzoate (**6x**)

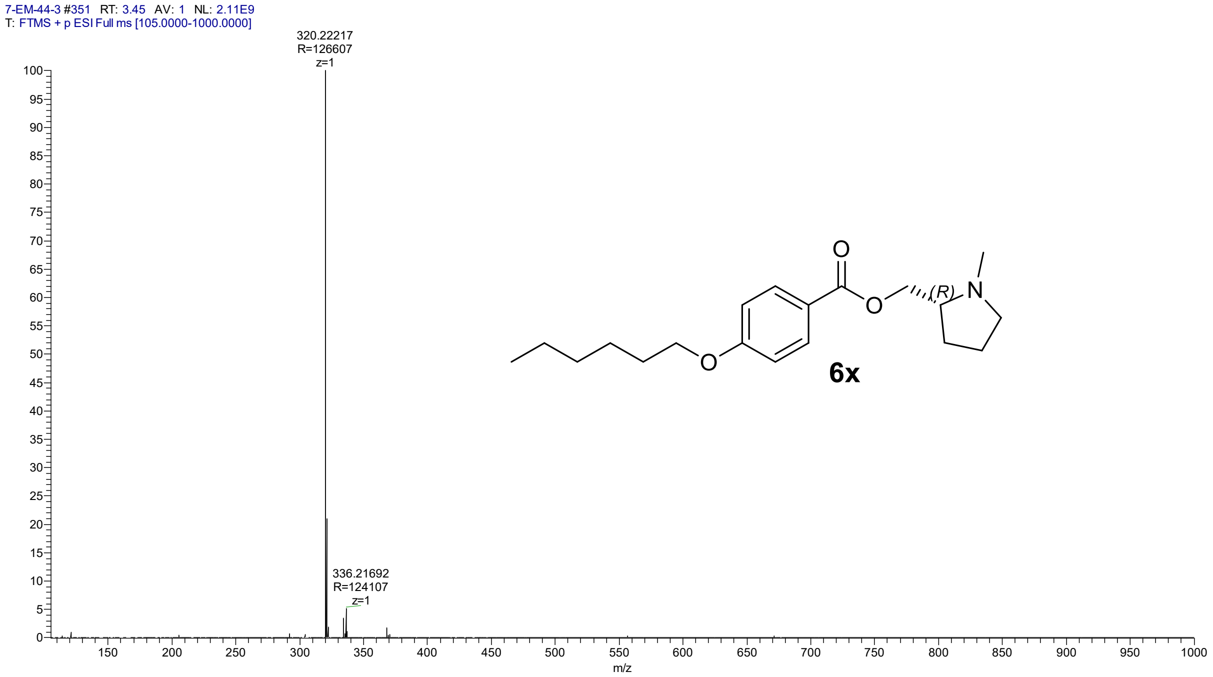

LC-UV chromatogram of [(2*R*)-1-methylpyrrolidin-2-yl]methyl 4-(hexyloxy)benzoate (**6x**)

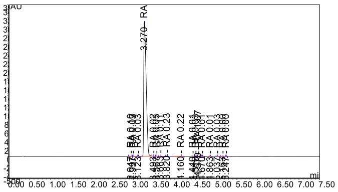

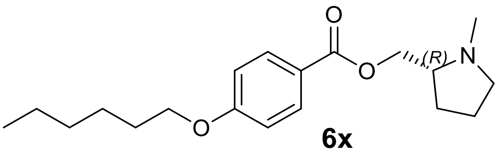

**6x**

HRMS spectrum of (1-methylpyrrolidin-3-yl)methyl 4-(hexyloxy)benzoate (**6y**)

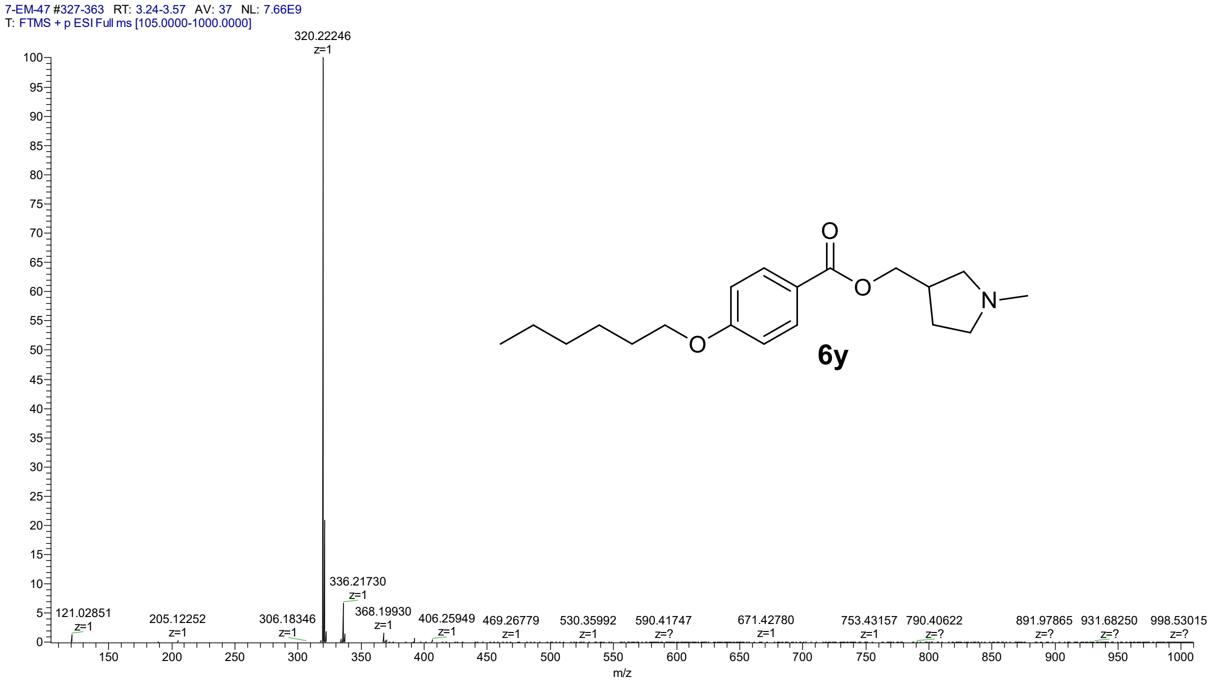

LC-UV chromatogram of (1-methylpyrrolidin-3-yl)methyl 4-(hexyloxy)benzoate (**6y**)

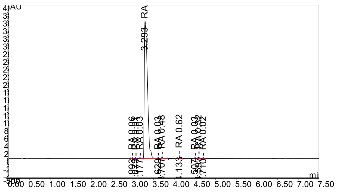

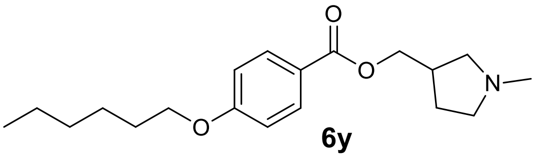

6y

HRMS spectrum of [(3*S*)-1-methylpyrrolidin-3-yl]methyl 4-(hexyloxy)benzoate (**6z**)

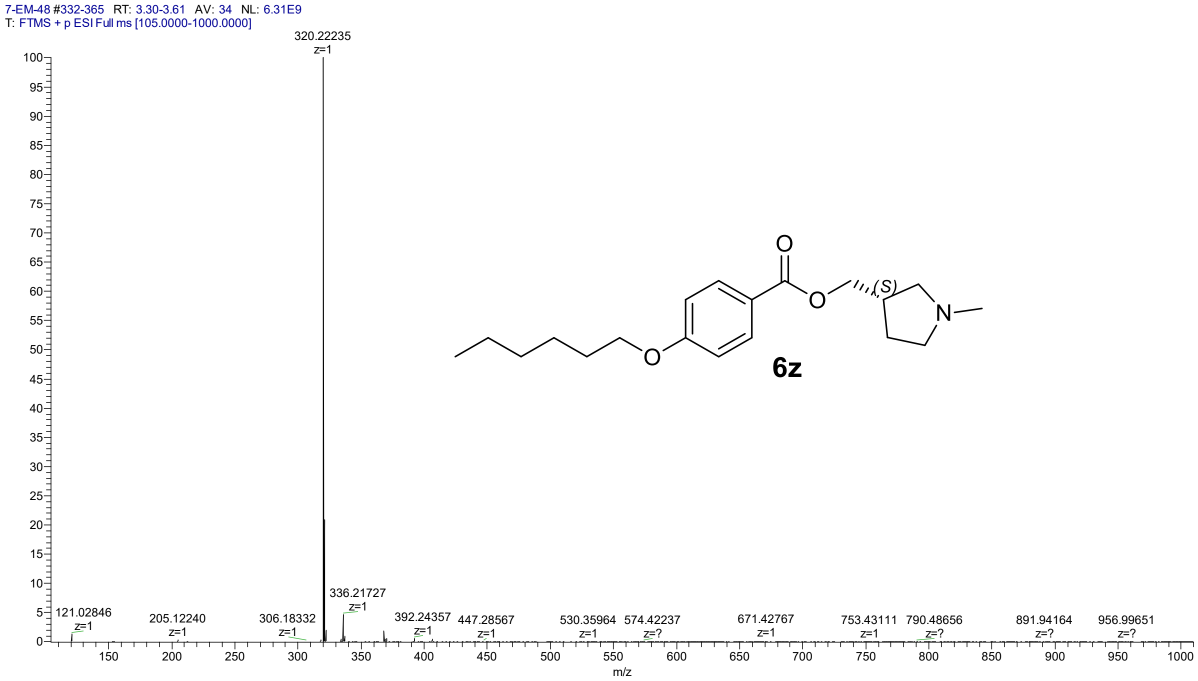

LC-UV chromatogram of [(3*S*)-1-methylpyrrolidin-3-yl]methyl 4-(hexyloxy)benzoate (**6z**)

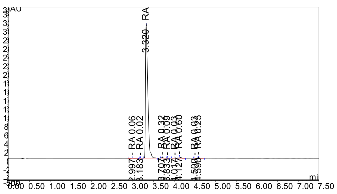

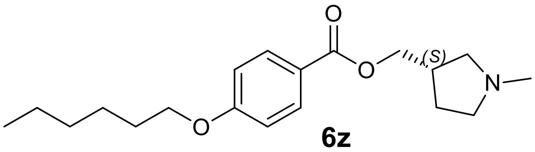

**6z**

HRMS spectrum of (2*R*)-1-{2-[4-(hexyloxy)benzoyloxy]ethyl}-1,2-dimethylpiperidin-1-ium iodide (**7e**)

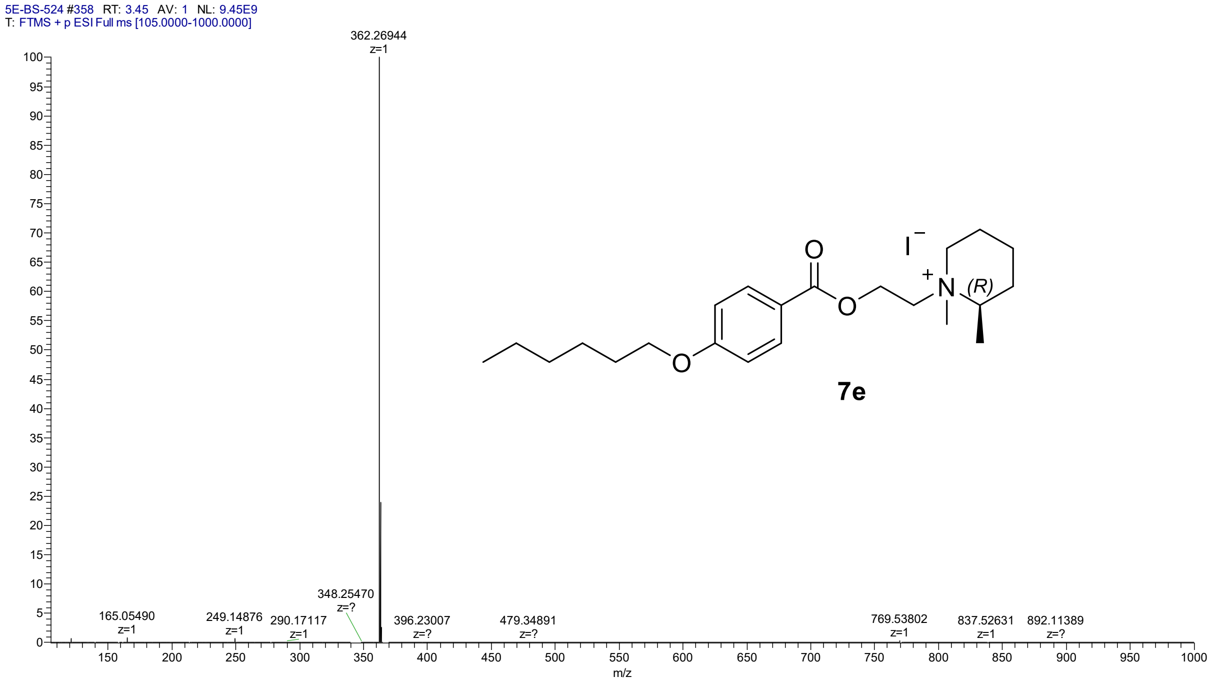

LC-UV chromatogram of (2*R*)-1-{2-[4-(hexyloxy)benzoyloxy]ethyl}-1,2-dimethylpiperidin-1-ium iodide (**7e**)

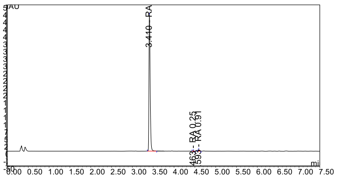

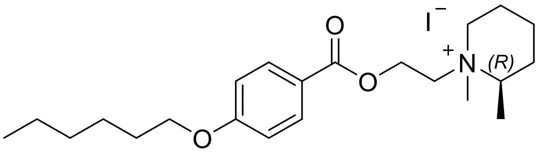

**7e**

HRMS spectrum of (2*S*)-1-{2-[4-(hexyloxy)benzoyloxy]ethyl}-1,2-dimethylpiperidin-1-ium iodide (**7f**)

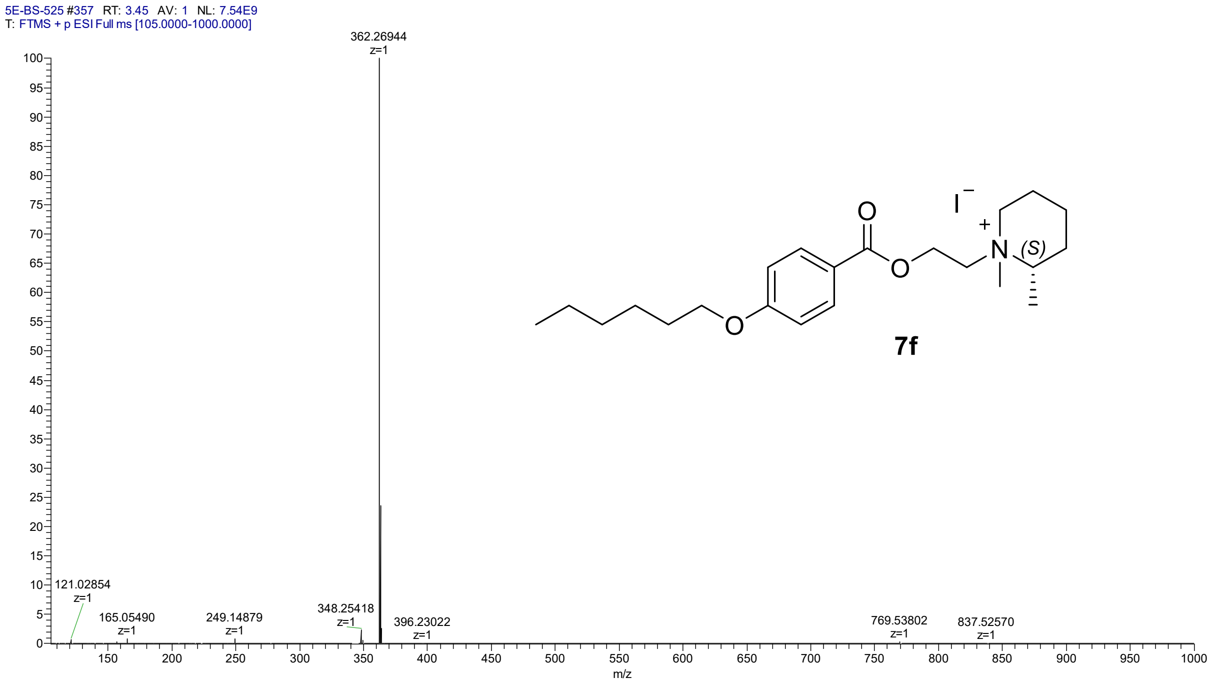

LC-UV chromatogram of (2*S*)-1-{2-[4-(hexyloxy)benzoyloxy]ethyl}-1,2-dimethylpiperidin-1-ium iodide (**7f**)

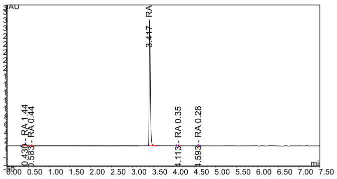

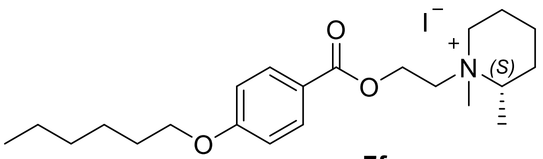

**7f**

HRMS spectrum of 2-{2-[4-(hexyloxy)benzoyloxy]ethyl}-2-methyl-2-azabicyclo[2.2.2]octan-2-ium iodide (**7g**)

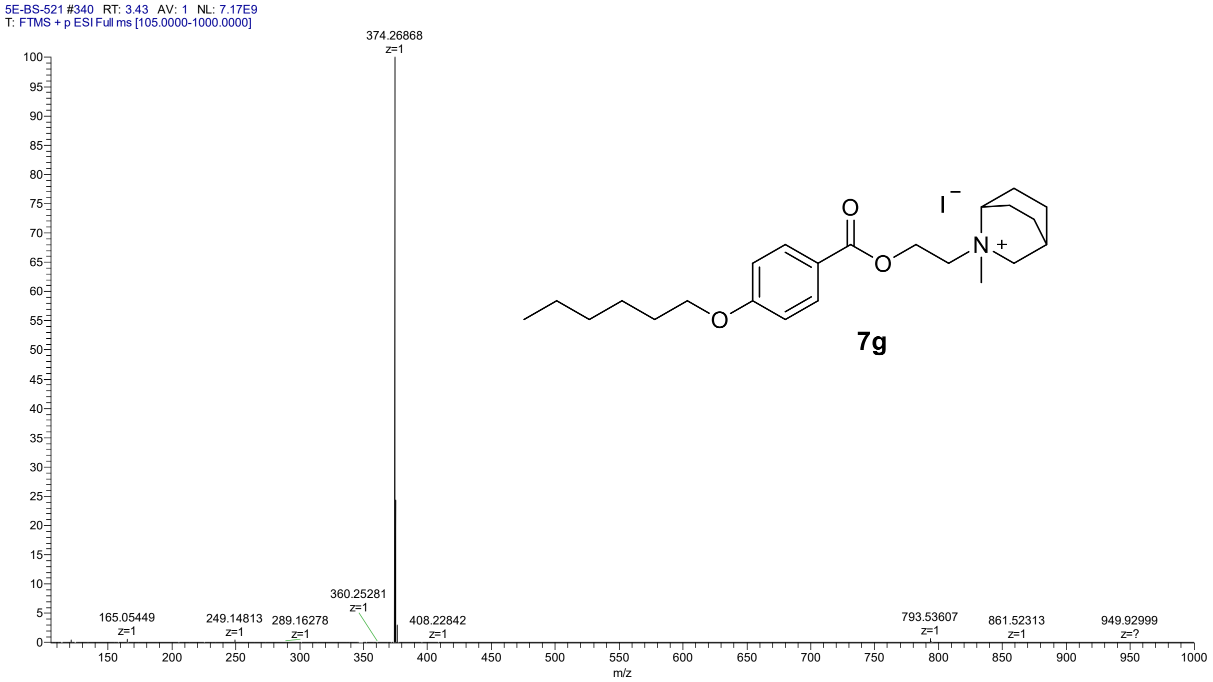

LC-UV chromatogram of 2-{2-[4-(hexyloxy)benzoyloxy]ethyl}-2-methyl-2-azabicyclo[2.2.2]octan-2-ium iodide (**7g**)

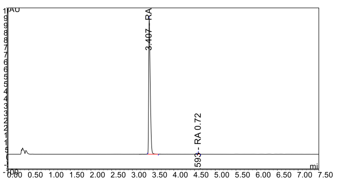

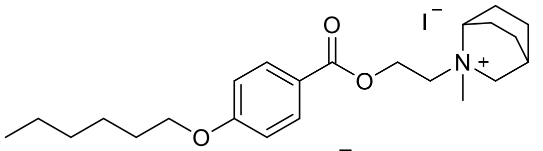

**7g**

HRMS spectrum of 1-{2-[4-(hexyloxy)benzoyloxy]ethyl}-1-methyl-2,5-dihydro-1H-pyrrol-1-ium iodide (**7h**)

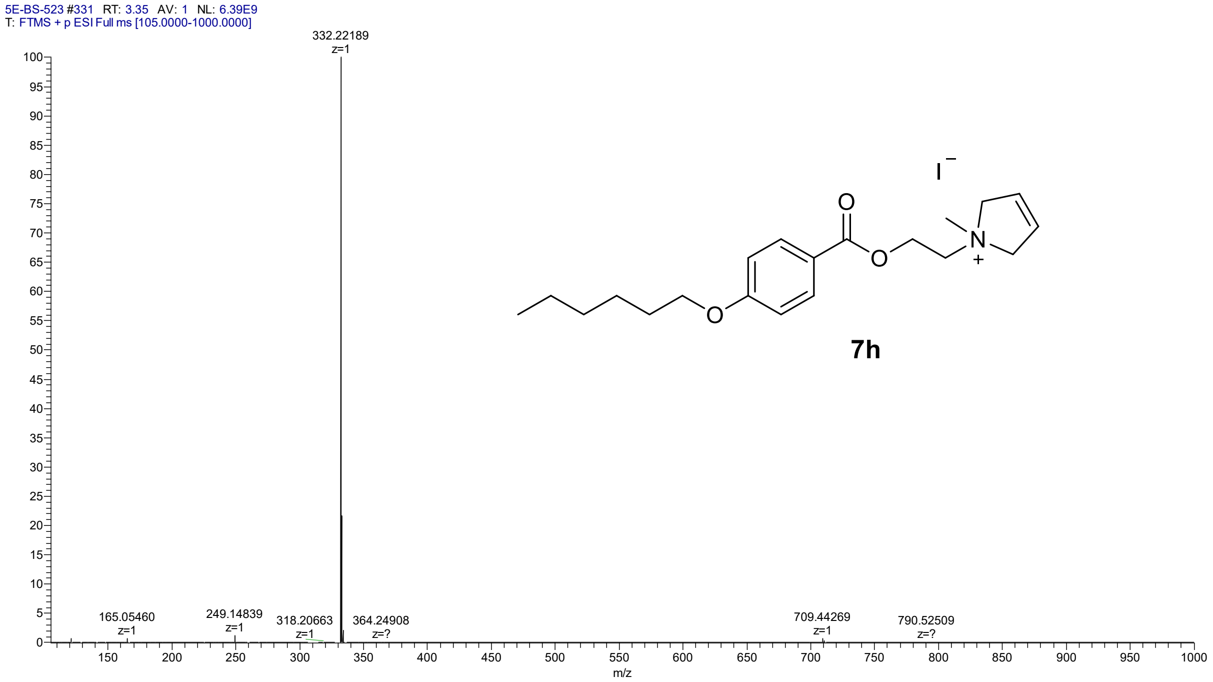

LC-UV chromatogram of 1-{2-[4-(hexyloxy)benzoyloxy]ethyl}-1-methyl-2,5-dihydro-1H-pyrrol-1-ium iodide (**7h**)

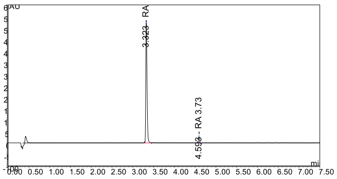

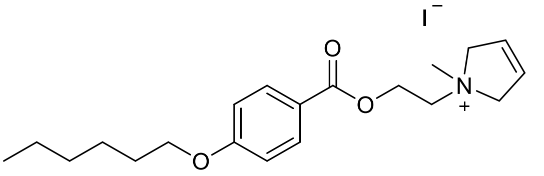

**7h**

HRMS spectrum of of 1-{2-[4-(hexyloxy)benzoyloxy]ethyl}-1-methylpyrrolidin-1-ium iodide (**7j**)

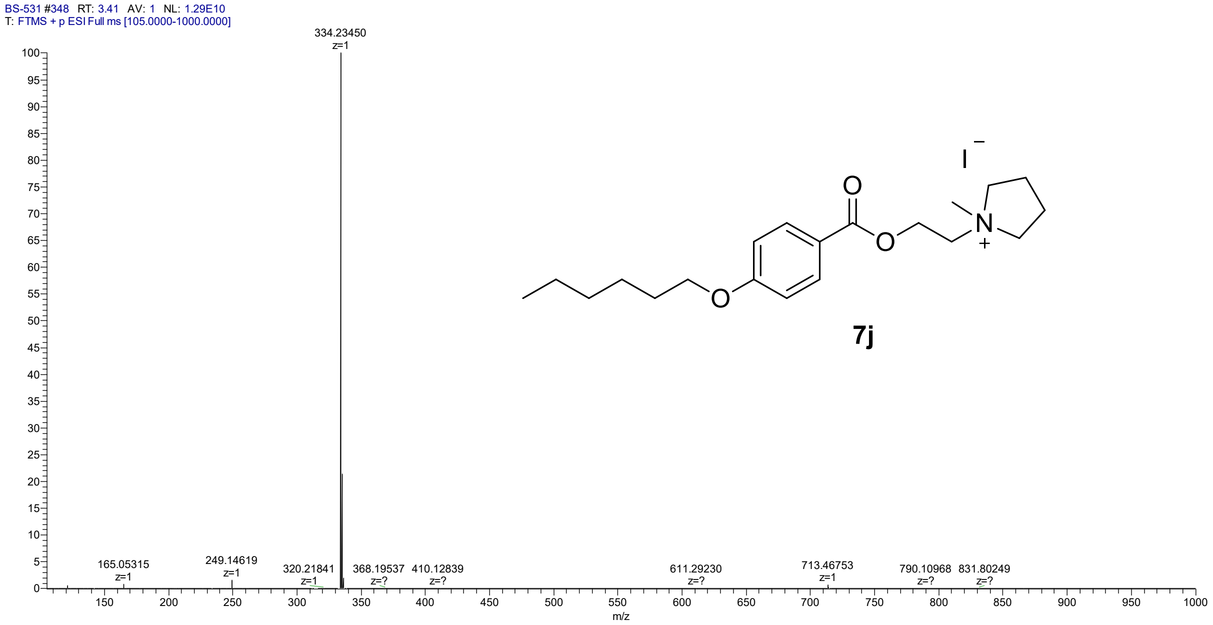

LC-UV chromatogram of of 1-{2-[4-(hexyloxy)benzoyloxy]ethyl}-1-methylpyrrolidin-1-ium iodide (**7j**)

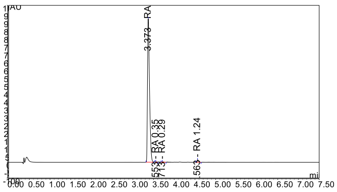

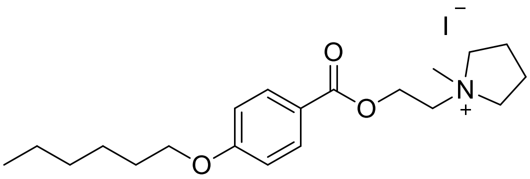

**7j**

HRMS spectrum of 2-{2-[4-(Hexyloxy)benzoyloxy]ethyl}-2-methyl-2,3-dihydro-1H-isoindol-2-ium iodide (**7k**)

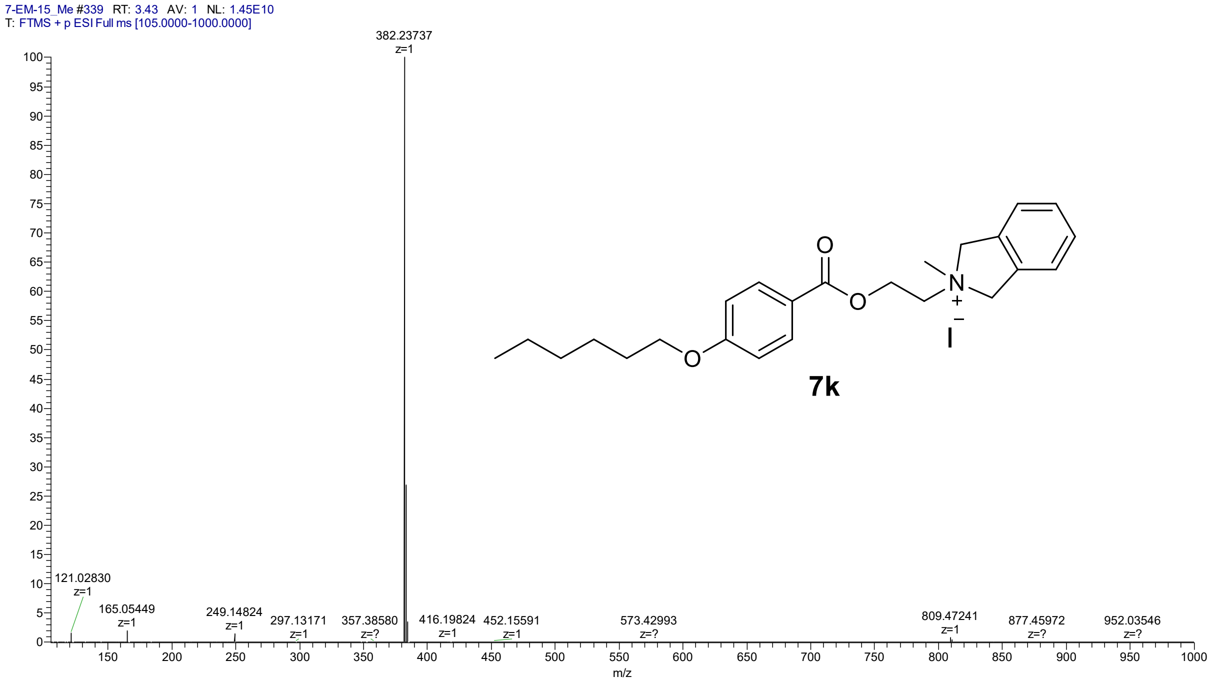

LC-UV chromatogram of 2-{2-[4-(Hexyloxy)benzoyloxy]ethyl}-2-methyl-2,3-dihydro-1H-isoindol-2-ium iodide (**7k**)

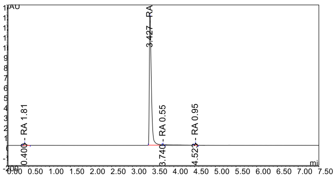

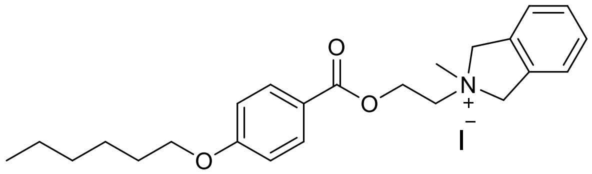

**7k**

HRMS spectrum of 2-{[4-(Hexyloxy)benzoyloxy]methyl}-1,1-dimethylpiperidin-1-ium iodide (**7m**)

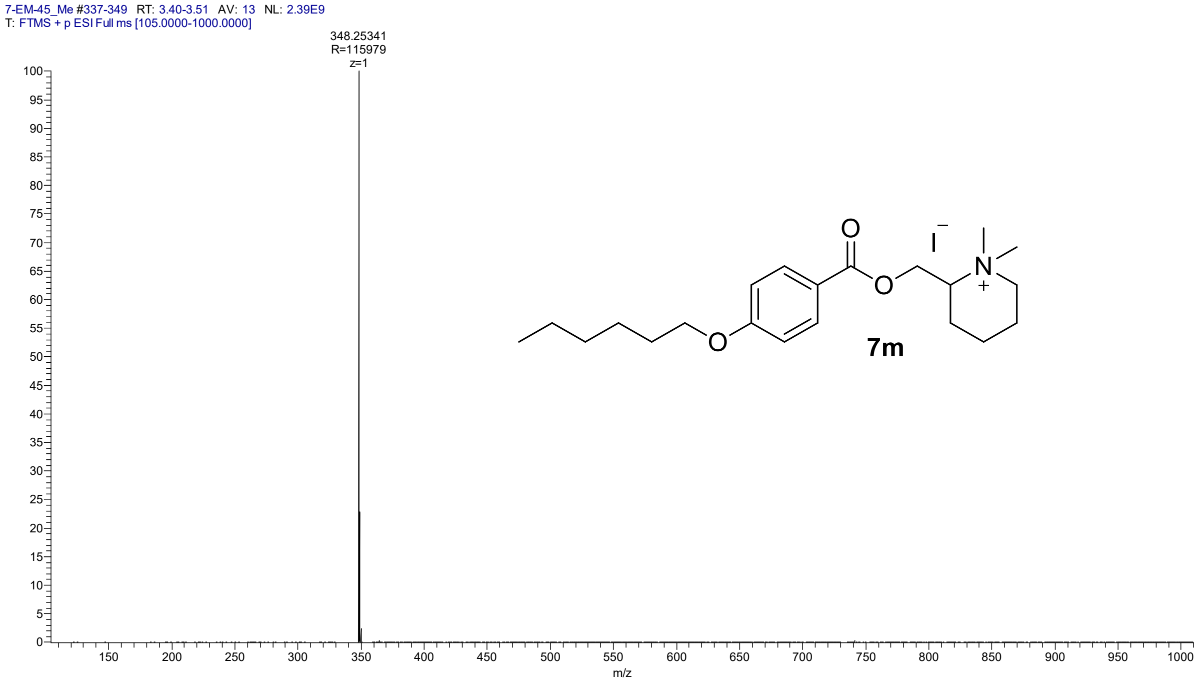

LC-UV chromatogram of 2-{[4-(Hexyloxy)benzoyloxy]methyl}-1,1-dimethylpiperidin-1-ium iodide (**7m**)

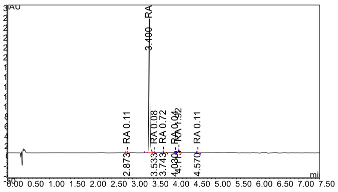

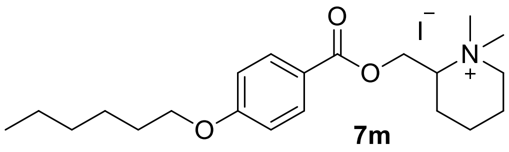

HRMS spectrum of (2*R*)-2-{[4-(Hexyloxy)benzoyloxy]methyl}-1,1-dimethylpiperidin-1-ium iodide (**7n**)

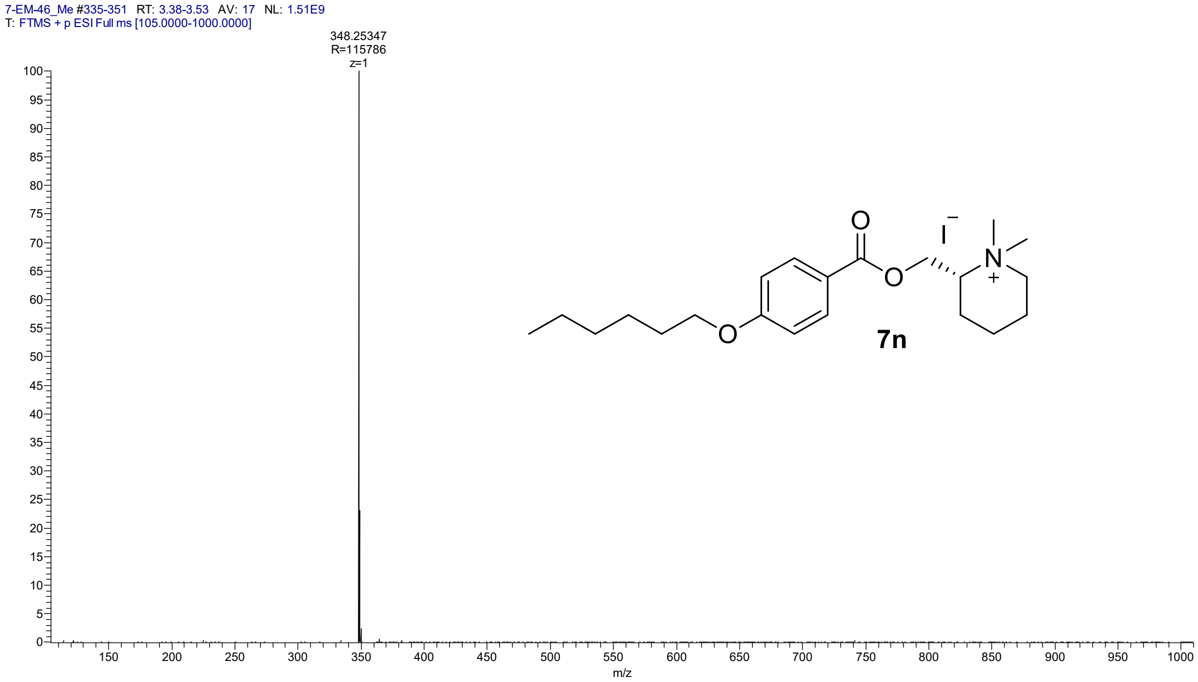

LC-UV chromatogram of (2*R*)-2-{[4-(Hexyloxy)benzoyloxy]methyl}-1,1-dimethylpiperidin-1-ium iodide (**7n**)

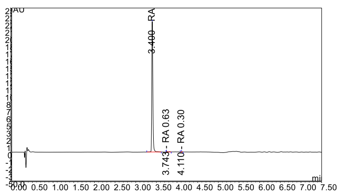

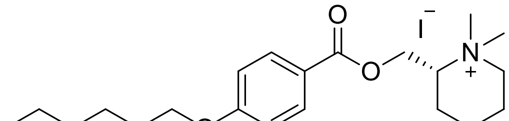

**7n**

HRMS spectrum of (3*S*)-3-{[4-(hexyloxy)benzoyloxy]methyl}-1,1-dimethylpiperidin-1-ium iodide (**7o**)

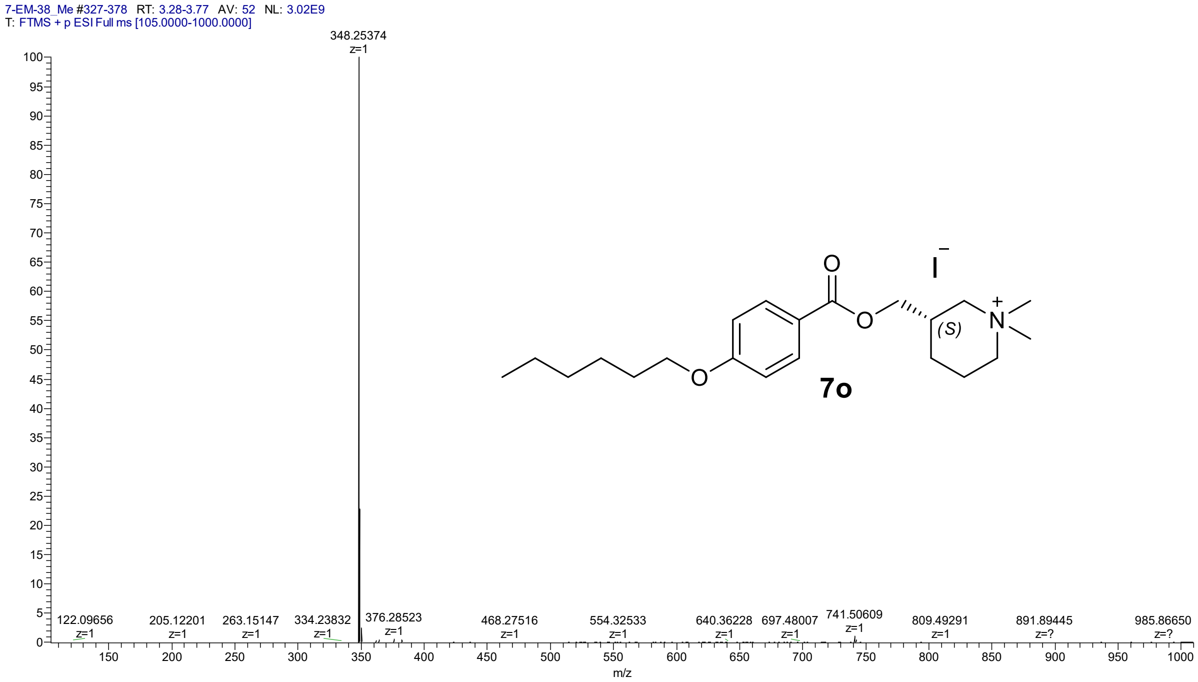

LC-UV chromatogram of (3*S*)-3-{[4-(hexyloxy)benzoyloxy]methyl}-1,1-dimethylpiperidin-1-ium iodide (**7o**)

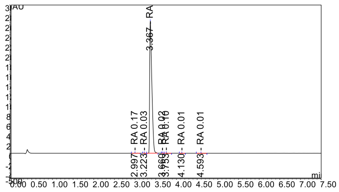

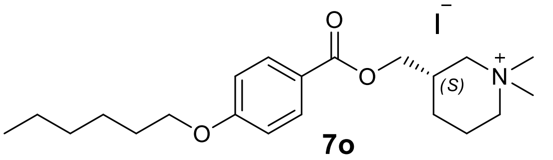

HMRS spectrum of 3-{[4-(hexyloxy)benzoyloxy]methyl}-1,1-dimethylpiperidin-1-ium iodide (**7p**)

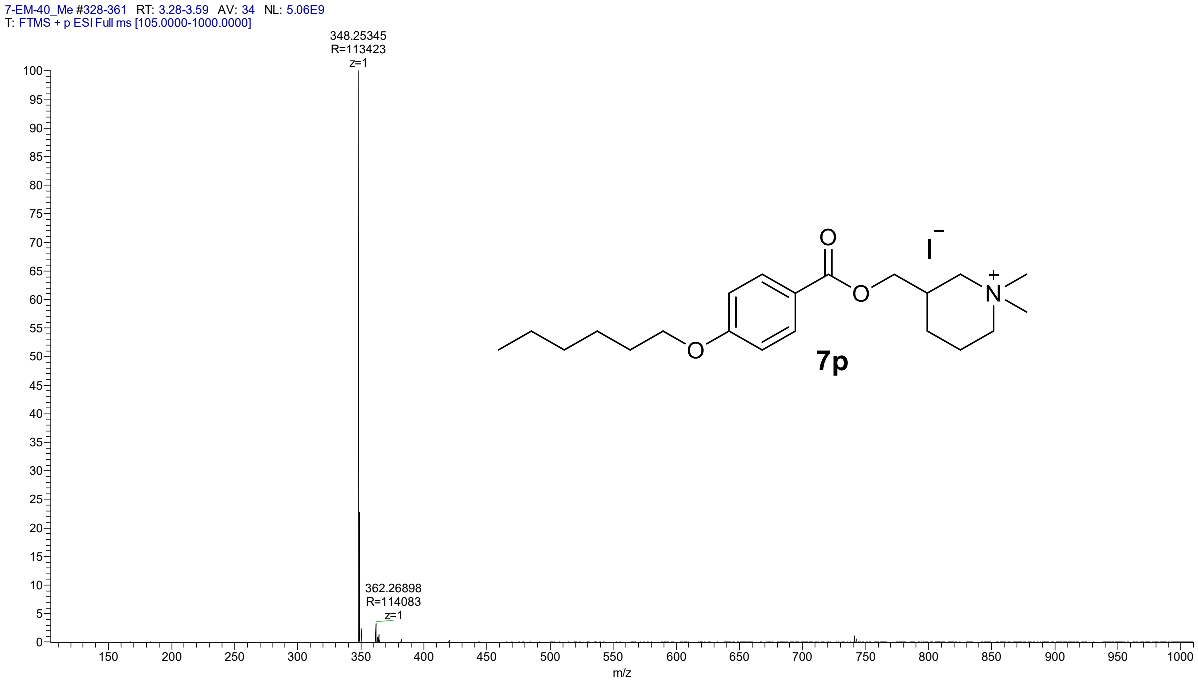

LC-UV chromatogram of 3-{[4-(hexyloxy)benzoyloxy]methyl}-1,1-dimethylpiperidin-1-ium iodide (**7p**)

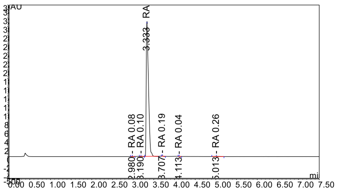

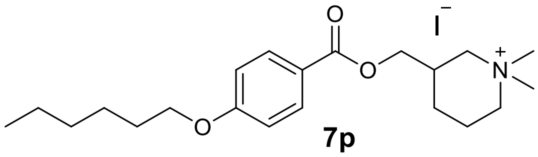

**7p**

HMRS spectrum of (3*R*)-3-{[4-(hexyloxy)benzoyloxy]methyl}-1,1-dimethylpiperidin-1-ium iodide (**7r**)

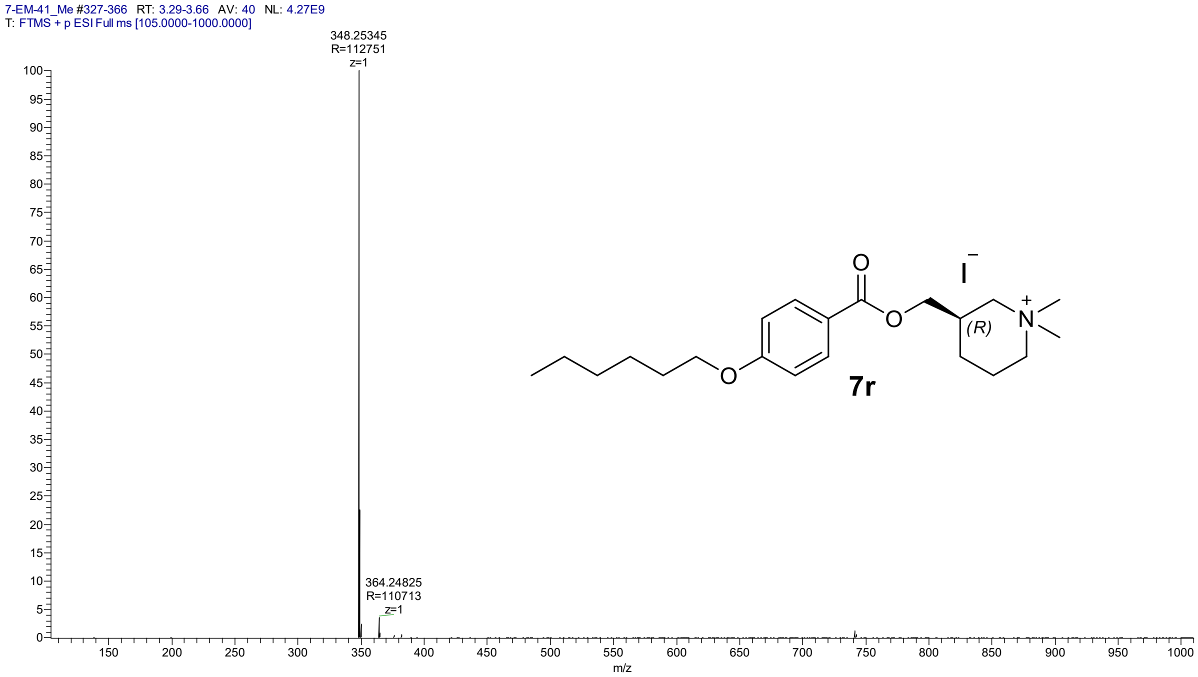

LC-UV chromatogram of (3*R*)-3-{[4-(hexyloxy)benzoyloxy]methyl}-1,1-dimethylpiperidin-1-ium iodide (**7r**)

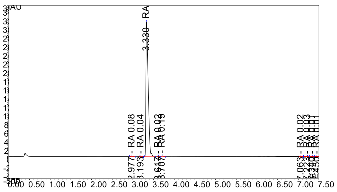

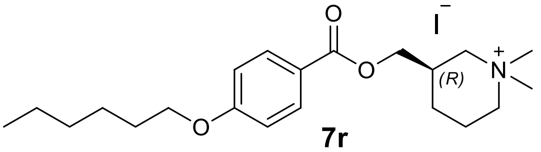

**7r**

HMRS spectrum of 4-[4-(hexyloxy)benzoyloxy]-1,1-dimethylpiperidin-1-ium iodide (**7s**)

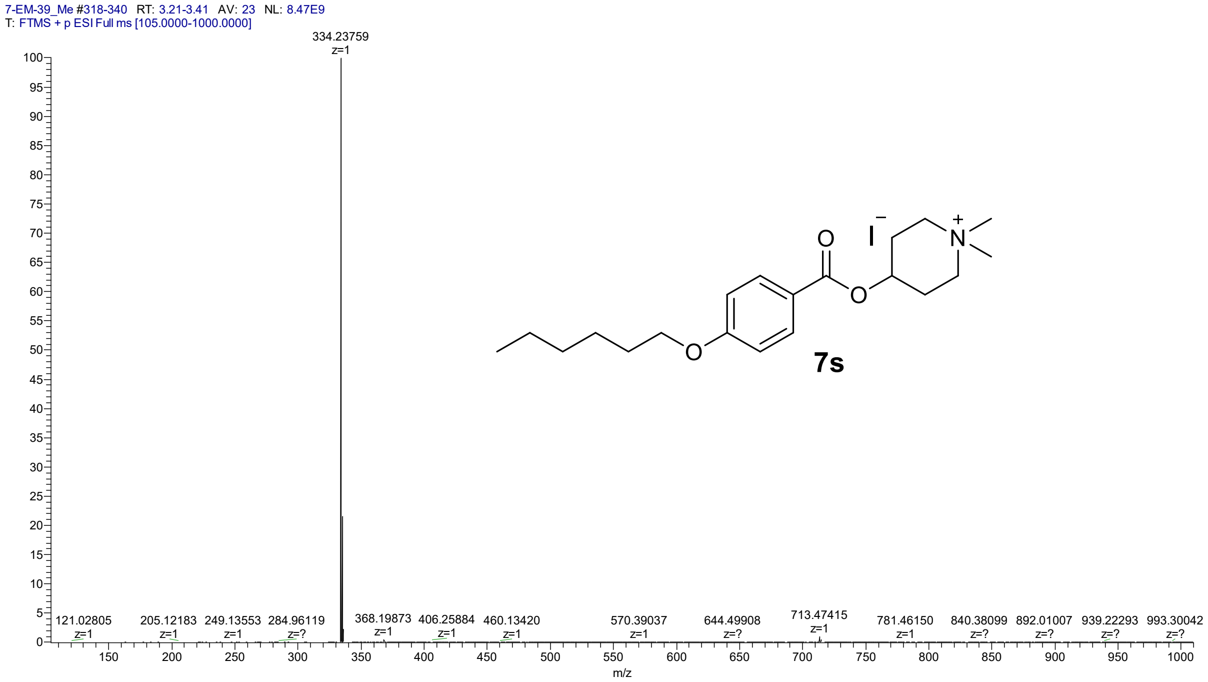

LC-UV chromatogram of 4-[4-(hexyloxy)benzoyloxy]-1,1-dimethylpiperidin-1-ium iodide (**7s**)

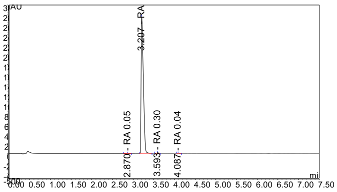

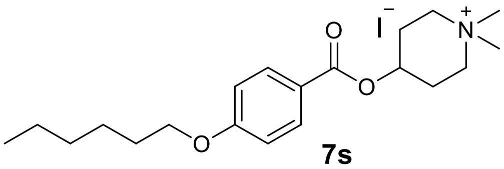

**7s**

HMRS spectrum of 2-{[4-(hexyloxy)benzoyloxy]methyl}-1,1-dimethylpyrrolidin-1-ium iodide (**7t**)

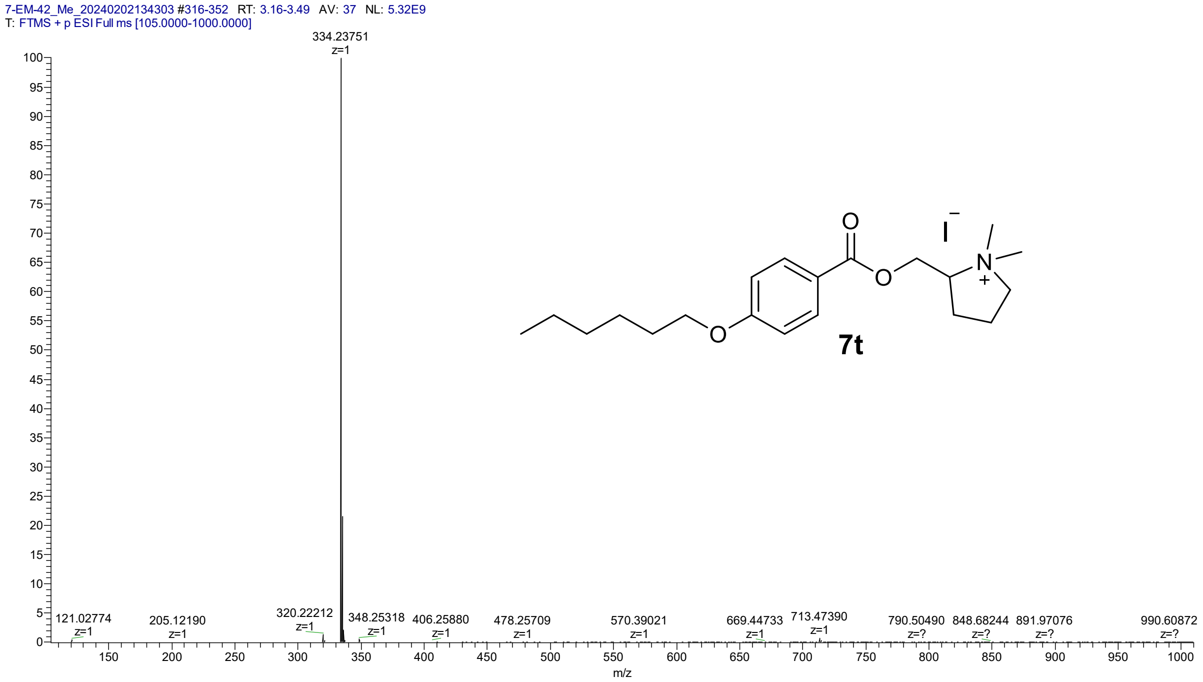

LC-UV chromatogram of 2-{[4-(hexyloxy)benzoyloxy]methyl}-1,1-dimethylpyrrolidin-1-ium iodide (**7t**)

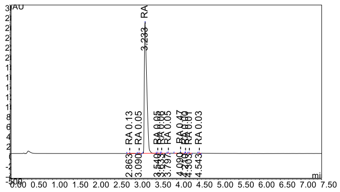

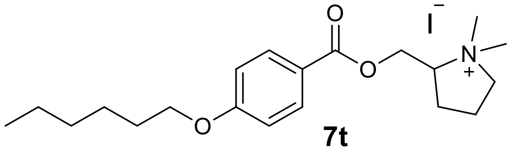

**7t**

HMRS spectrum of (2*S*)-2-{[4-(hexyloxy)benzoyloxy]methyl}-1,1-dimethylpyrrolidin-1-ium iodide (**7w**)

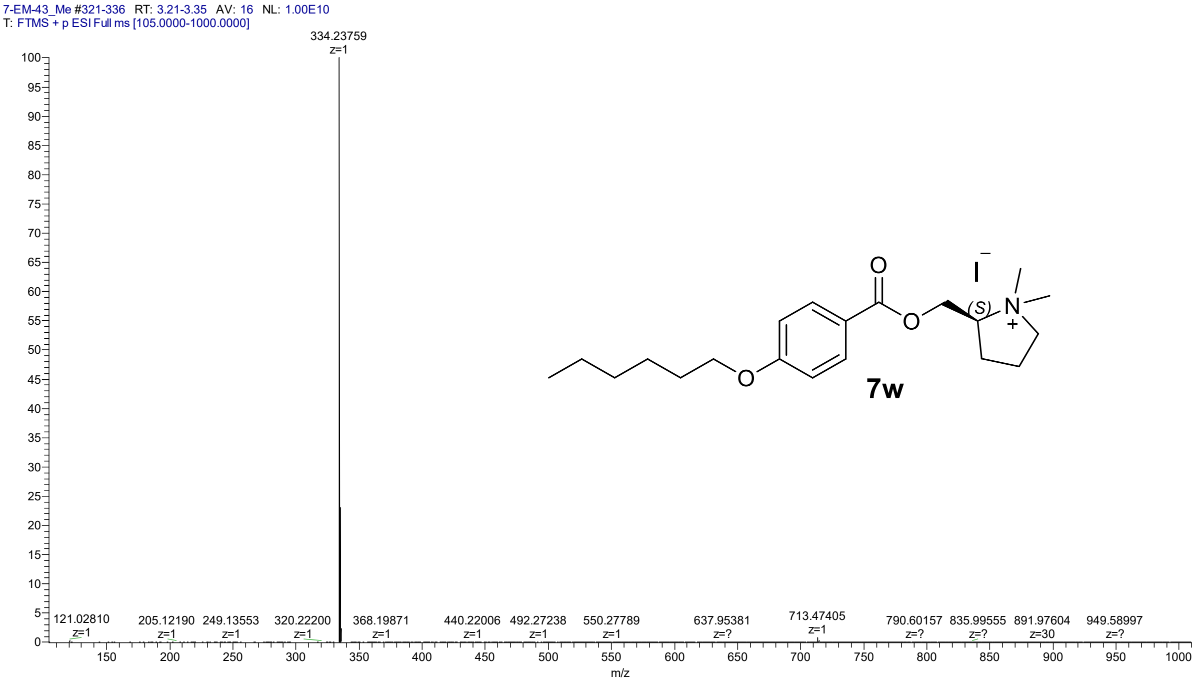

LC-UV chromatogram of (2*S*)-2-{[4-(hexyloxy)benzoyloxy]methyl}-1,1-dimethylpyrrolidin-1-ium iodide (**7w**)

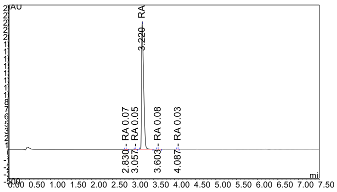

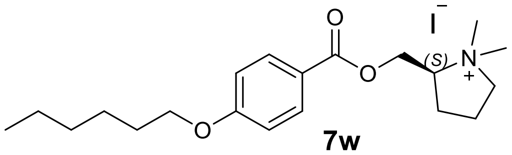

**7w**

HMRS spectrum of (2*R*)-2-{[4-(hexyloxy)benzoyloxy]methyl}-1,1-dimethylpyrrolidin-1-ium iodide (**7x**)

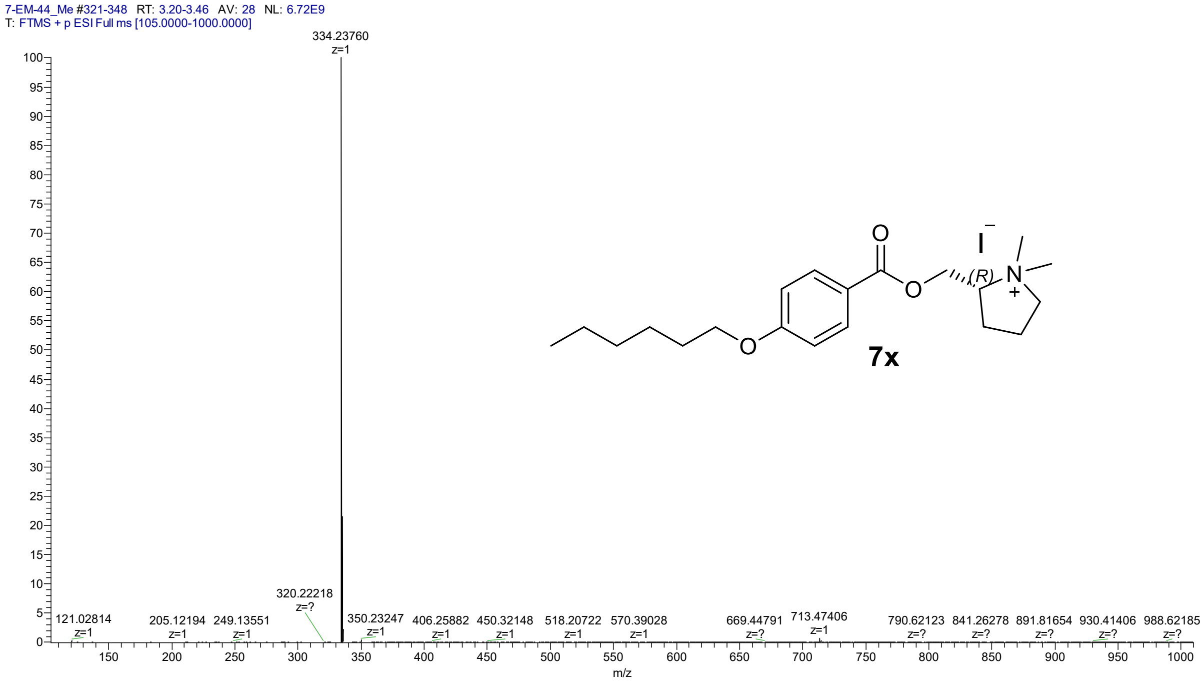

LC-UV chromatogram of (2*R*)-2-{[4-(hexyloxy)benzoyloxy]methyl}-1,1-dimethylpyrrolidin-1-ium iodide (**7x**)

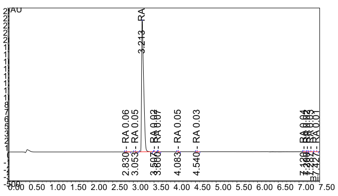

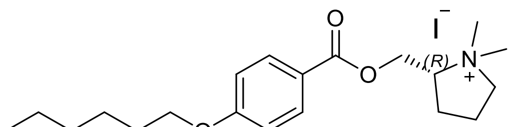

**7x**

HMRS spectrum of 3-{[4-(hexyloxy)benzoyloxy]methyl}-1,1-dimethylpyrrolidin-1-ium iodide (**7y**)

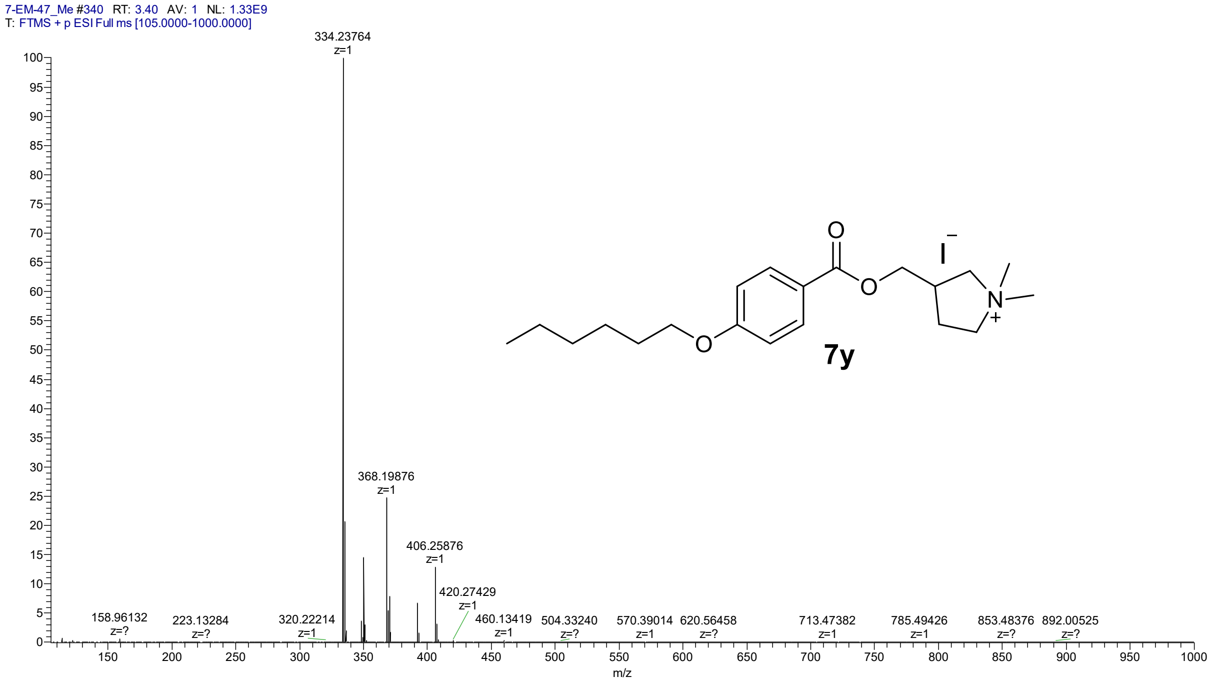

LC-UV chromatogram of 3-{[4-(hexyloxy)benzoyloxy]methyl}-1,1-dimethylpyrrolidin-1-ium iodide (**7y**)

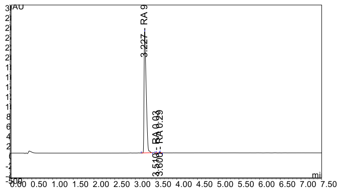

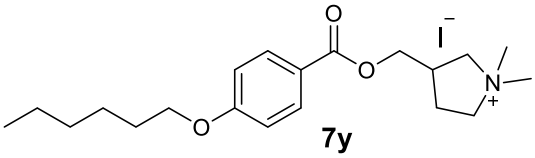

HRMS spectrum of (3*S*)-3-{[4-(hexyloxy)benzoyloxy]methyl}-1,1-dimethylpyrrolidin-1-ium iodide (**7z**)

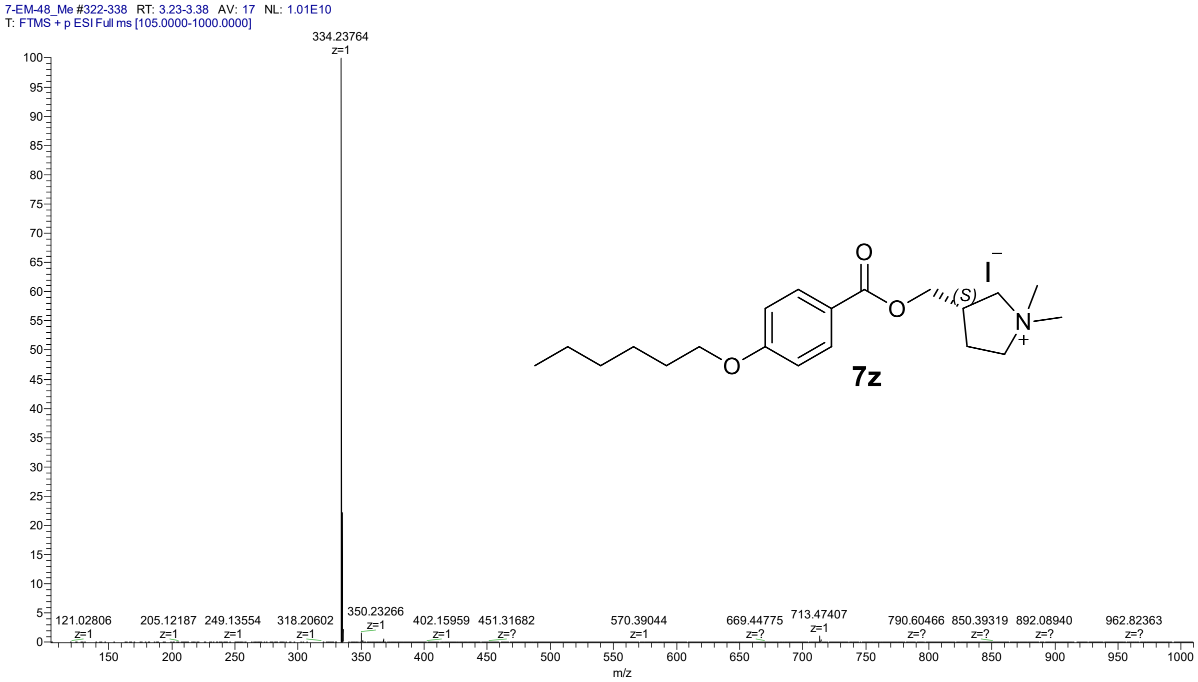

LC-UV chromatogram of (3*S*)-3-{[4-(hexyloxy)benzoyloxy]methyl}-1,1-dimethylpyrrolidin-1-ium iodide (**7z**)

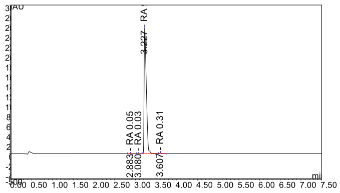

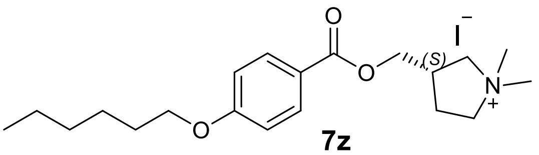

**7z**

HRMS spectrum of quinuclidin-3-yl 4-hexyloxybenzoate hydrochloride (**12a**)

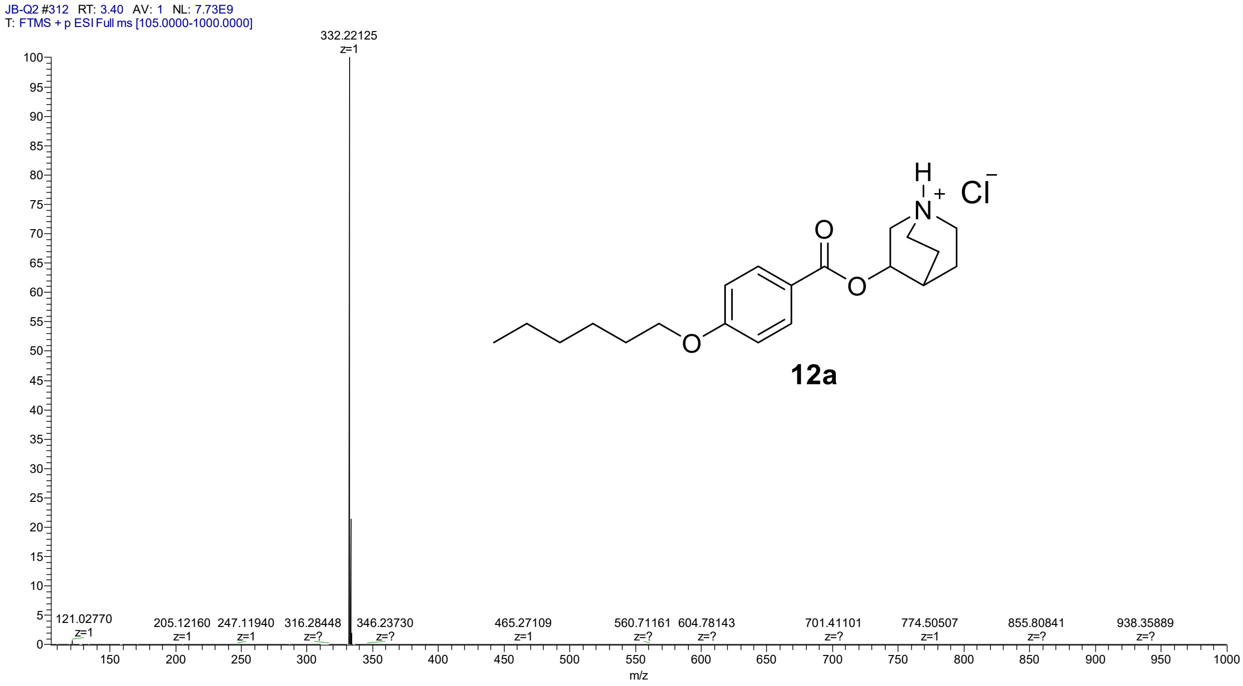

LC-UV chromatogram of quinuclidin-3-yl 4-hexyloxybenzoate hydrochloride (**12a**)

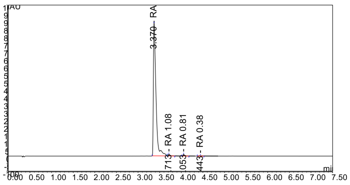

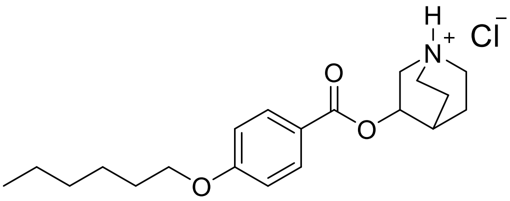

**12a**

HRMS spectrum of 1-methylquinuclidin-3-yl 4-hexyloxybenzoate iodide (**12b**):

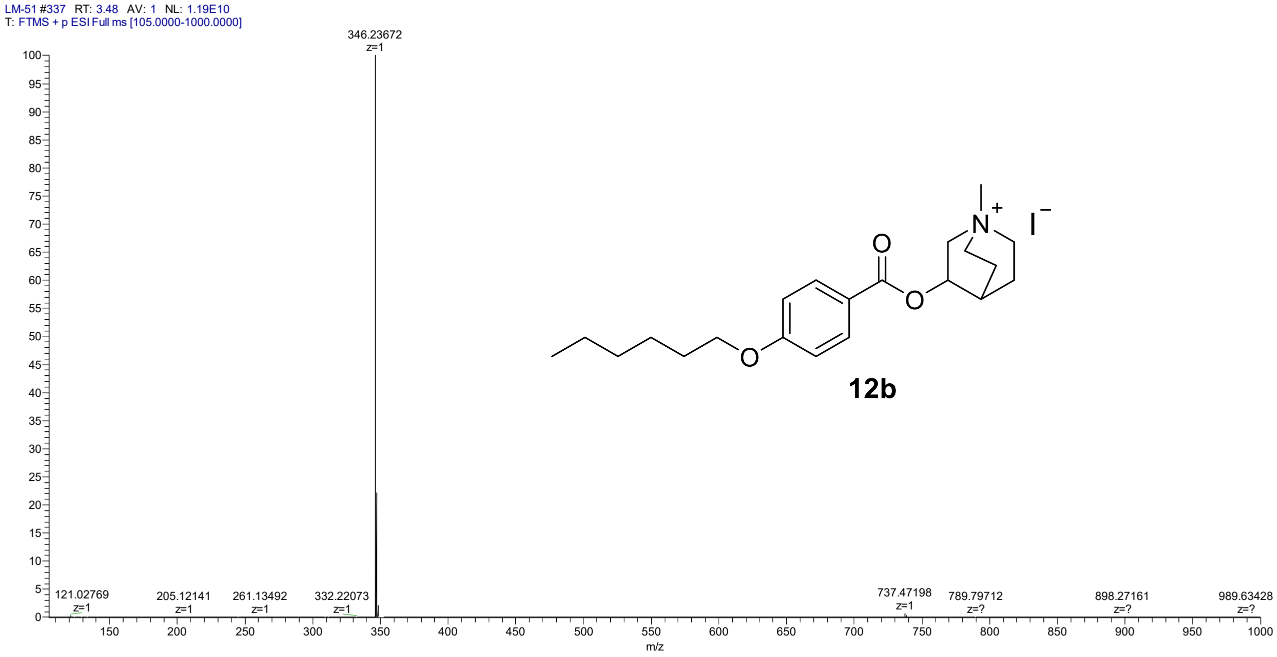

LC-UV chromatogram of 1-methylquinuclidin-3-yl 4-hexyloxybenzoate iodide (**12b**):

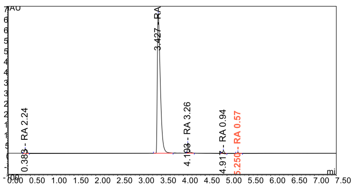

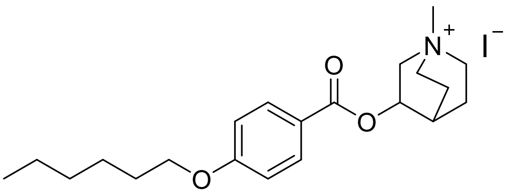

**12b**

HRMS spectrum of quinuclidin-3-yl 4-butoxybenzoate hydrochloride (**13a**)

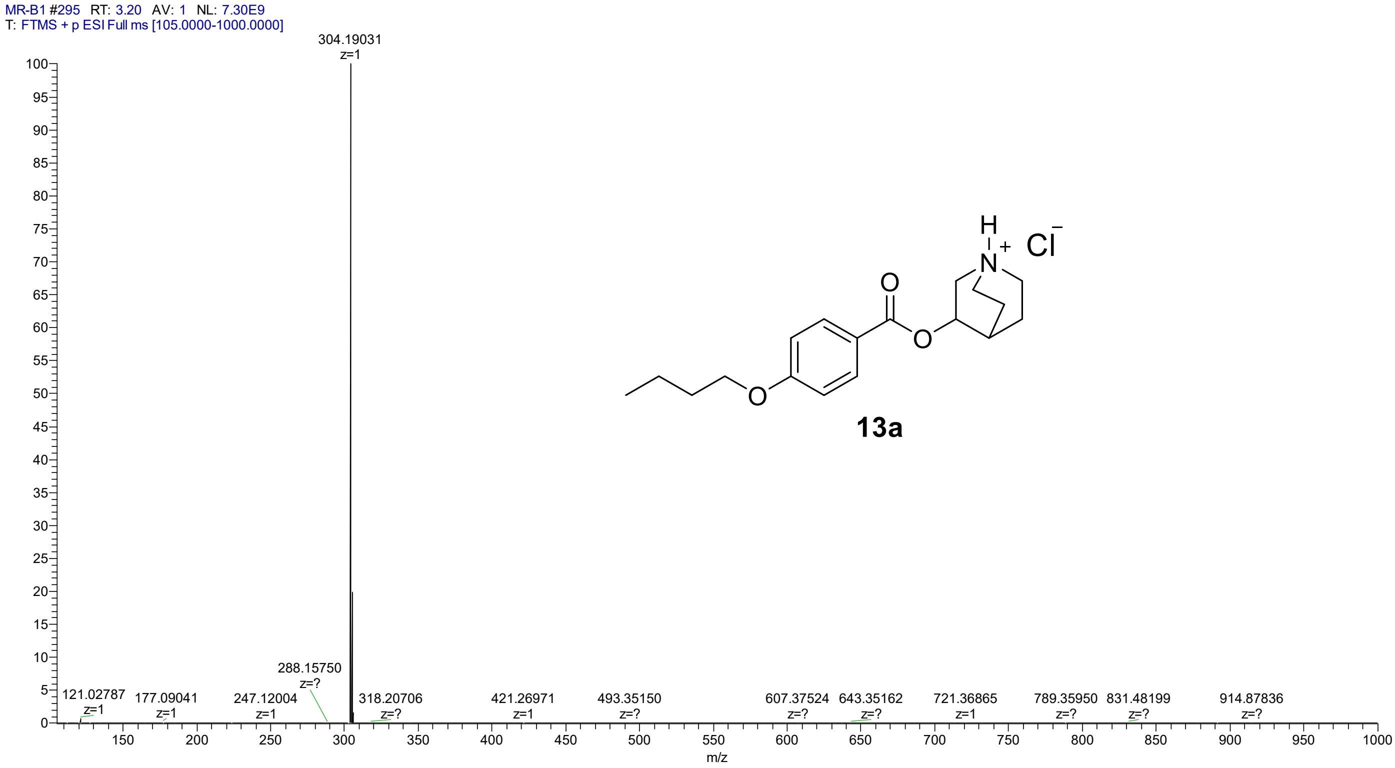

LC-UV chromatogram of quinuclidin-3-yl 4-butoxybenzoate hydrochloride (**13a**)

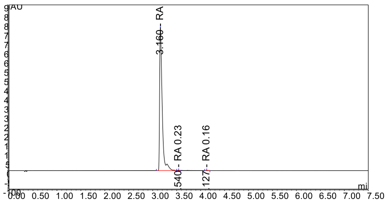

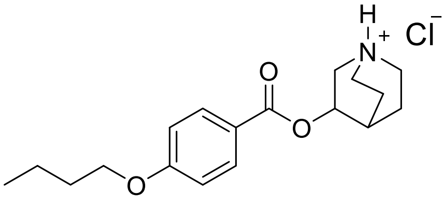

**13a**

HRMS spectrum of 1-methylquinuclidin-3-yl 4-butoxybenzoate iodide (**13b**)

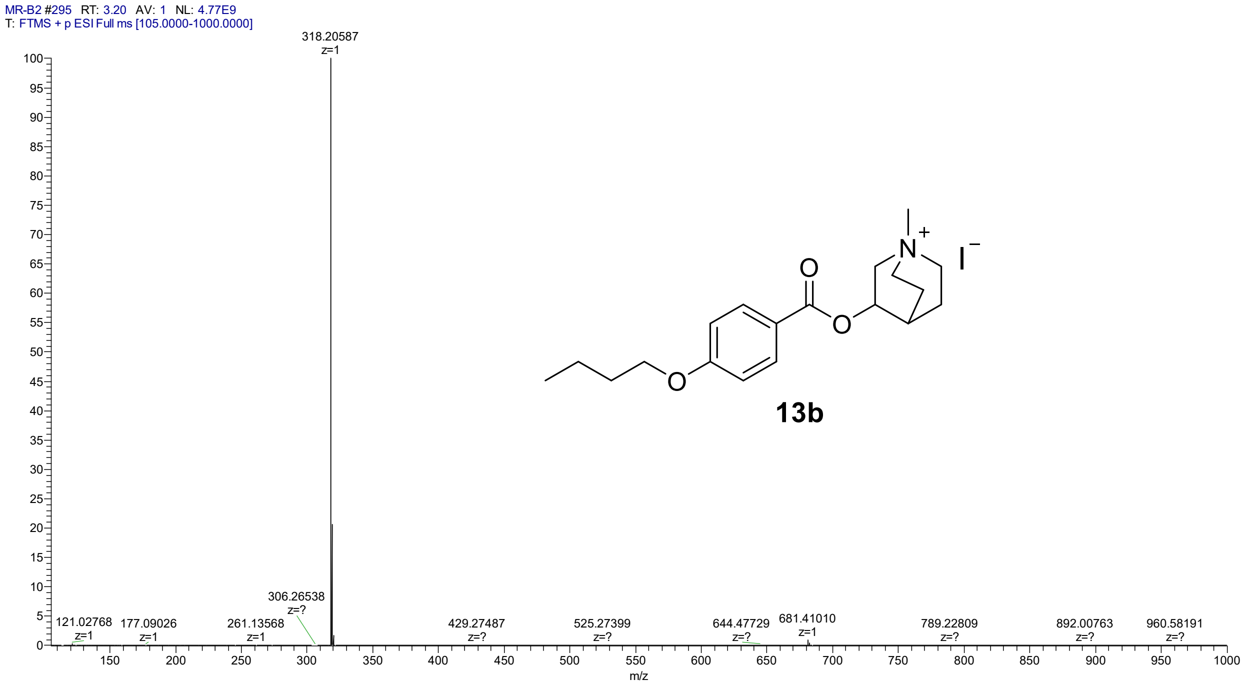

LC-UV chromatogram 1-methylquinuclidin-3-yl 4-butoxybenzoate iodide (**13b**)

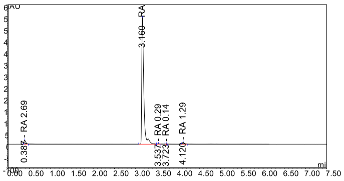

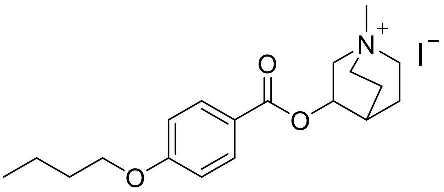

**13b**

HRMS spectrum of quinuclidine-3-yl 4-methoxybenzoate hydrochloride (**14a**):

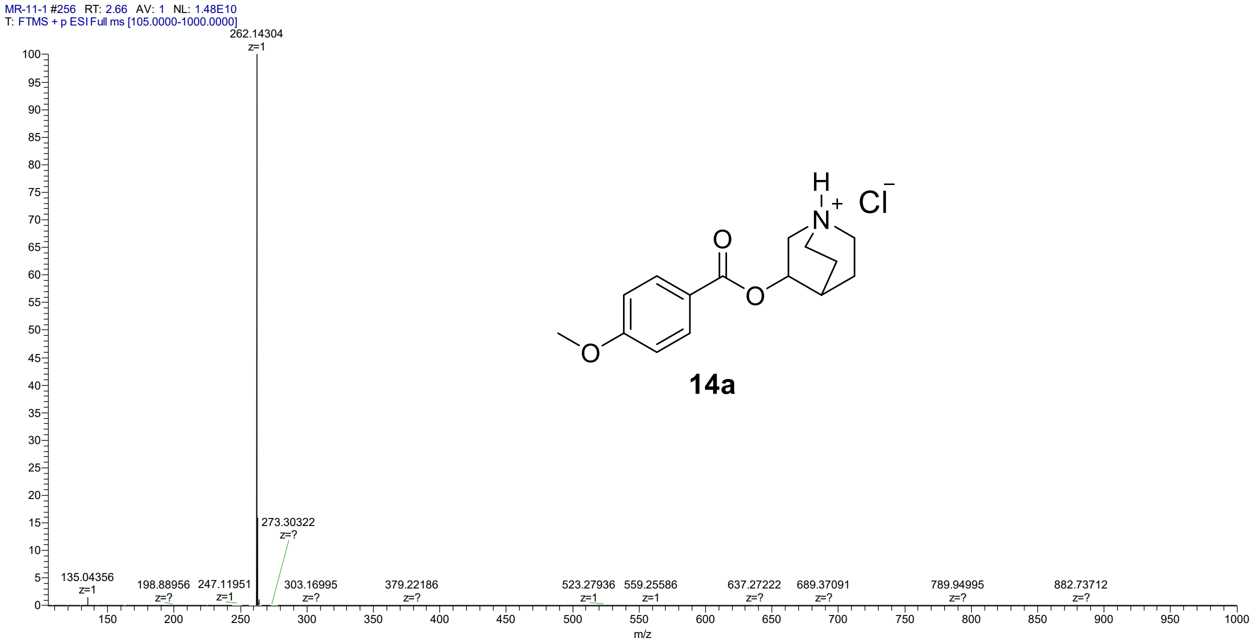

LC-UV chromatogram of quinuclidine-3-yl 4-methoxybenzoate hydrochloride (**14a**):

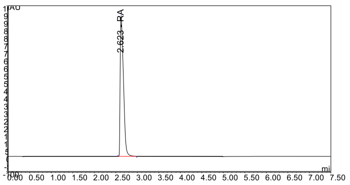

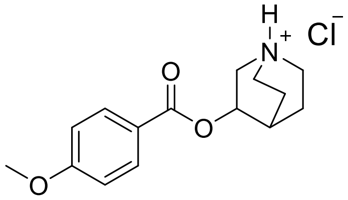

**14a**

HRMS spectrum of 1-methylquinuclidin-3-yl 4-methoxybenzoate iodide (**14b**):

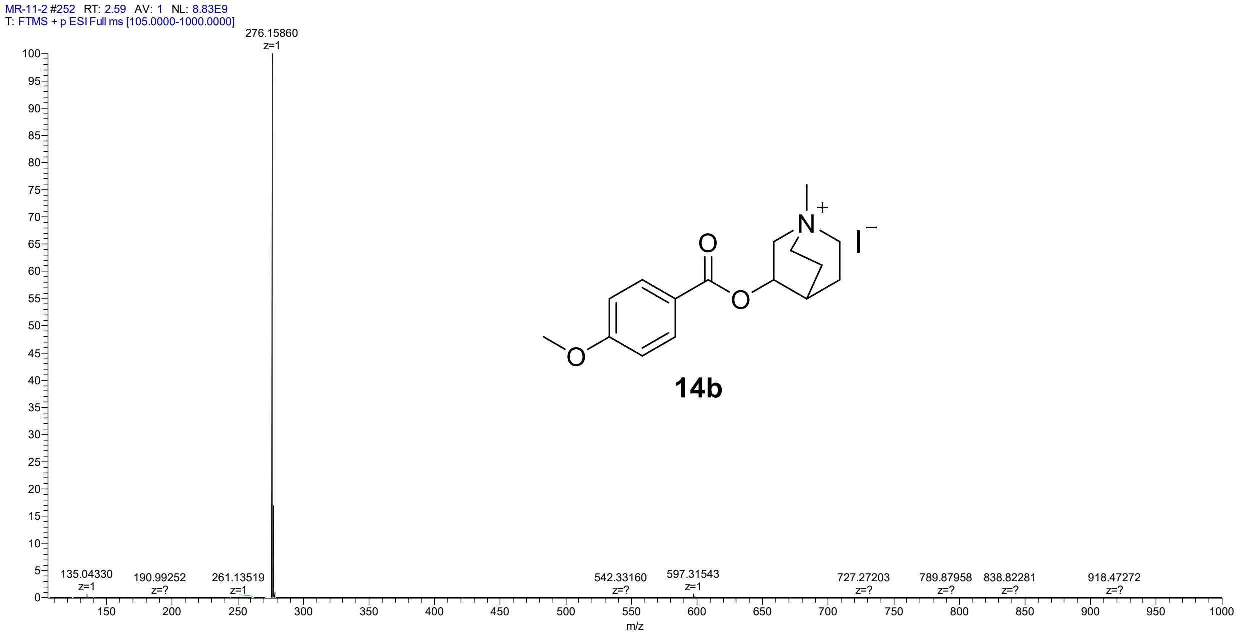

LC-UV chromatogram of 1-methylquinuclidin-3-yl 4-methoxybenzoate iodide (**14b**):

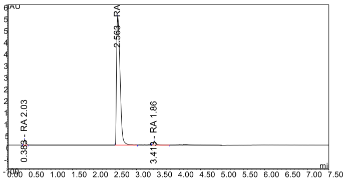

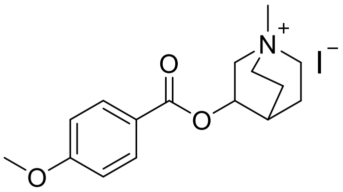

**14b**

#### 4. Figures

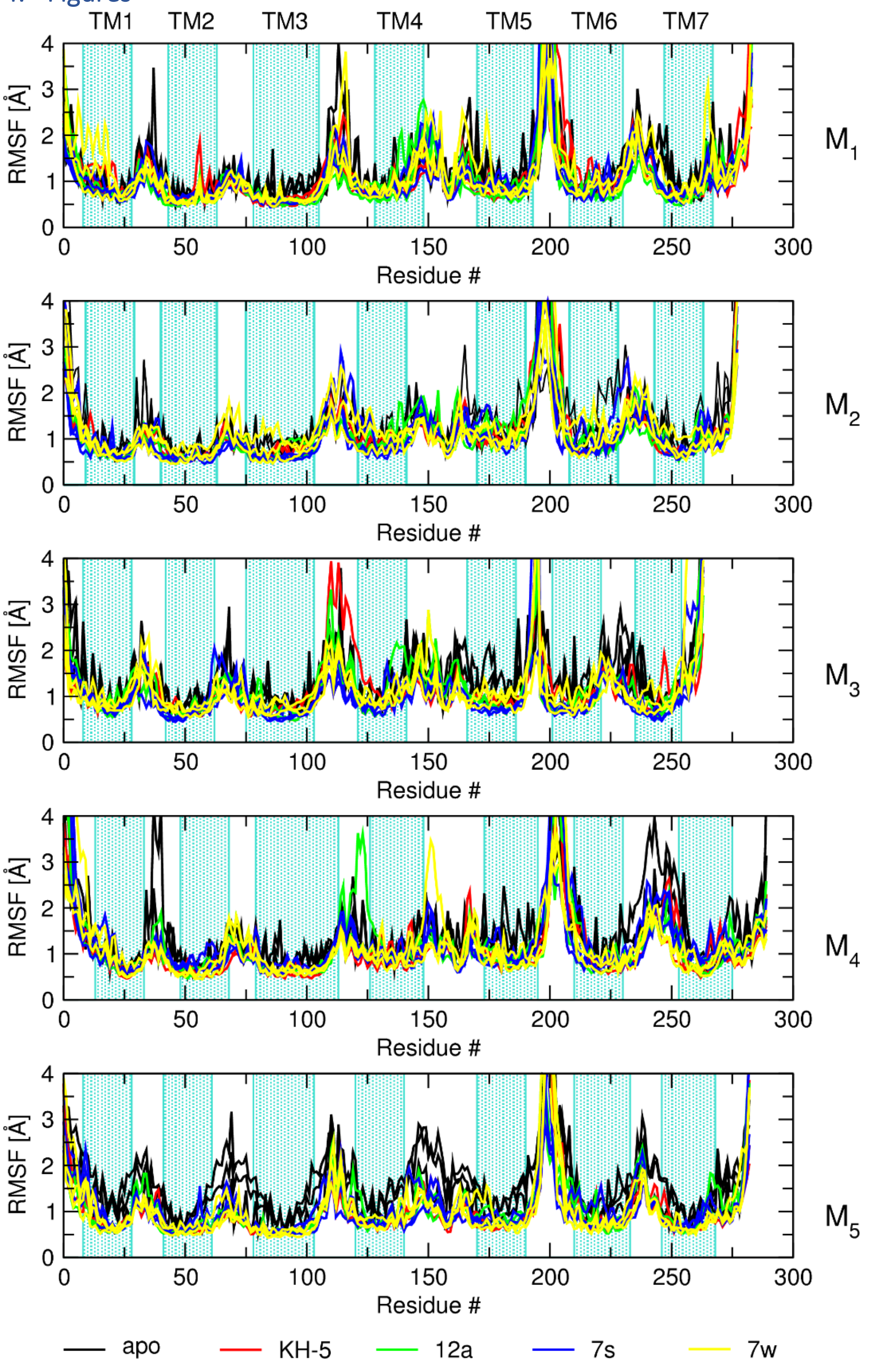

***Figure S1 Receptor RMSF***

*Root mean square of fluctuations (RMSF) of individual amino acid residues of ligand-receptor complexes or empty receptor (indicated in the legend in Å is plotted against residue number. Transmembrane helices are indicated by turquoise hatching. Traces are from 3 independent runs of MD.*

#### 5. Tables

***Table S1 Affinity ($pK_i$), potency ($pK_B$) and antagonism half-life ($t_{1/2}$) after washing of parental compounds.***
*Affinities are expressed as the negative decadic logarithm of inhibition constant ($pK_i$) determined from competition with [$^3$H]NMS. Potencies are expressed as negative decadic logarithms inhibition constant ($pK_B$) of the carbachol-induced functional response. Residence times are expressed as half-life ($t_{1/2}$) of the antagonism of the functional response of one-hour exposure to 10 µM compound, followed by washing. Data are means ± SD from 5 independent experiments. Data were published by Randakova et al. 2018.*

| compound | | $M_1$ | $M_2$ | $M_3$ | $M_4$ | $M_5$ |
|---|---|---|---|---|---|---|
| 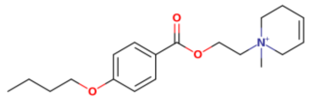 HD-4-2 | $pK_i$ | 5.70 ± 0.05 | 5.71 ± 0.05 | 5.23 ± 0.06 | 5.11 ± 0.07 | 5.47 ± 0.07 |
| | $pK_B$ | 7.41 ± 0.04 | 5.81 ± 0.03 | 5.50 ± 0.02 | 6.59 ± 0.03 | 5.79 ± 0.03 |
| | $t_{1/2}$ [h] | 0.4 ± 0.1 | 0.2 ± 0.1 | 0.4 ± 0.1 | 0.4 ± 0.1 | 0.5 ± 0.1 |
| 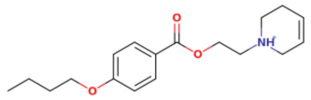 BK-23 | $pK_i$ | 5.19 ± 0.05 | 5.50 ± 0.05 | 5.11 ± 0.05 | 4.95 ± 0.04 | 5.20 ± 0.04 |
| | $pK_B$ | 4.40 ± 0.06 | 4.93 ± 0.05 | 4.53 ± 0.05 | 4.11 ± 0.06 | 4.71 ± 0.05 |
| | $t_{1/2}$ [h] | 0.4 ± 0.1 | 0.2 ± 0.1 | 0.2 ± 0.1 | 0.4 ± 0.1 | 0.3 ± 0.1 |
| 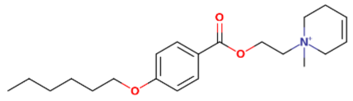 KH-5 | $pK_i$ | 6.50 ± 0.04 | 6.82 ± 0.02 | 6.38 ± 0.07 | 6.36 ± 0.05 | 6.79 ± 0.02 |
| | $pK_B$ | 7.68 ± 0.04 | 5.70 ± 0.03 | 7.10 ± 0.04 | 6.86 ± 0.03 | 7.31 ± 0.04 |
| | $t_{1/2}$ [h] | 4.5 ± 0.4 | 1.7 ± 0.2 | 4.1 ± 0.3 | 4.3 ± 0.4 | 5.1 ± 0.4 |
| 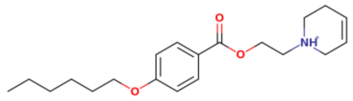 OT-231 | $pK_i$ | 5.37 ± 0.03 | 5.49 ± 0.06 | 5.24 ± 0.06 | 5.04 ± 0.05 | 5.18 ± 0.05 |
| | $pK_B$ | 7.48 ± 0.04 | 5.73 ± 0.03 | 5.39 ± 0.04 | 6.56 ± 0.03 | 5.68 ± 0.03 |
| | $t_{1/2}$ [h] | 4.4 ± 0.4 | 2.0 ± 0.2 | 2.7 ± 0.3 | 4.3 ± 0.4 | 2.6 ± 0.3 |

***Table S2 Binding affinities of stereoisomers 6x and 6w, and 7x and 7w.***
*Binding affinities are expressed in nM. Values are means with 95% confidence intervals in parentheses from 3 independent experiments performed in quadruplicates. *, higher than S-stereoisomer (P<0.05, one-tail t-test)*

| | $M_1$ | $M_2$ | $M_3$ | $M_4$ | $M_5$ |
|---|---|---|---|---|---|
| **6x** | 525 ( 447 - 617 ) | 457 ( 372 - 562 ) | 1122 ( 933 - 1349 ) | 776 ( 661 - 912 ) | 603 ( 490 - 741 ) |
| **6w** | 977 ( 813 - 1175 ) * | 617 ( 513 - 741 ) | 2818 ( 2399 - 3311 ) * | 2089 ( 1738 - 2512 ) * | 1585 ( 1288 - 1950 ) * |
| **7x** | 155 ( 126 - 191 ) | 141 ( 120 - 166 ) | 513 ( 490 - 537 ) | 372 ( 324 - 427 ) | 191 ( 182 - 200 ) |
| **7w** | 324 ( 309 - 339 ) * | 200 ( 166 - 240 ) | 933 ( 891 - 977 ) * | 794 ( 724 - 871 ) * | 550 ( 525 - 575 ) * |

***Table S3 Estimates of binding energies of docked compounds***
*Estimates of binding energies in kcal/mol of top poses of rescored docking poses of antagonists to the orthosteric binding sites of individual receptors in inactive conformations and presence of key interactions: salt bridge to $D^{3.32}$, π-π or cation-π to $Y^{3.33}$ and $Y^{7.39}$, hydrogen bond to $T^{5.42}$ and $N^{6.52}$ and cation-π to $W^{6.48}$, + meaning presence and – meaning the absence of the interaction.*

| compound | $M_1$ 5CXV | $M_2$ 3UON | $M_3$ 4DAJ | $M_4$ 5DSG | $M_5$ 6OL9 | $D^{3.32}$ | $Y^{3.33}$ | $T^{5.42}$ | $W^{6.48}$ | $N^{6.52}$ | $Y^{7.39}$ |
|---|---|---|---|---|---|---|---|---|---|---|---|
| **KH-5 (R)** | 6.95 | 5.85 | 7.74 | 6.34 | 7.10 | + | + | + | + | + | + |
| **KH-5 (S)** | 7.01 | 6.11 | 7.76 | 6.86 | 7.12 | + | + | + | + | + | + |
| **6a** | 9.09 | 8.45 | 8.53 | 8.93 | 8.97 | - | - | - | - | + | - |
| **6b** | 8.04 | 7.23 | 8.24 | 8.40 | 8.12 | - | + | - | + | + | - |
| **6c** | 7.49 | 7.06 | 7.36 | 7.50 | 7.44 | - | - | - | - | + | - |
| **6d** | 9.16 | 8.57 | 9.29 | 9.34 | 9.23 | - | + | - | + | + | - |
| **6e** | 8.03 | 7.22 | 7.29 | 7.48 | 7.83 | - | - | + | + | + | - |
| **6f** | 8.10 | 7.59 | 7.69 | 8.05 | 8.08 | - | - | + | + | + | - |
| **6g** | 7.93 | 7.61 | 7.69 | 7.53 | 8.51 | - | - | - | - | + | - |
| **6h** | 7.22 | 7.28 | 7.05 | 7.50 | 7.59 | - | +[a] | - | - | + | - |
| **6j** | 6.41 | 6.67 | 6.50 | 6.31 | 7.62 | - | - | + | - | + | - |
| **6k** | 8.69 | 7.92 | 8.31 | 8.87 | 8.80 | - | - | - | - | + | - |

| compound | $M_1$ 5CXV | $M_2$ 3UON | $M_3$ 4DAJ | $M_4$ 5DSG | $M_5$ 6OL9 | $D^{3.32}$ | $Y^{3.33}$ | $T^{5.42}$ | $W^{6.48}$ | $N^{6.52}$ | $Y^{7.39}$ |
|---|---|---|---|---|---|---|---|---|---|---|---|
| **6l** | 9.13 | 7.67 | 8.23 | 8.87 | 9.38 | - | + | - | + | + | - |
| **6m(S)** | 7.36 | 7.82 | 7.55 | 7.16 | 8.63 | - | - | + | - | + | - |
| **6n(R)** | 7.30 | 7.48 | 7.30 | 7.20 | 8.50 | - | - | + | - | + | - |
| **6o** | 7.24 | 7.86 | 7.15 | 6.87 | 8.07 | - | - | - | - | + | - |
| **6r** | 7.20 | 7.50 | 7.49 | 7.42 | 8.17 | - | - | - | - | + | - |
| **6s** | 7.50 | 7.31 | 7.55 | 8.21 | 8.17 | - | - | - | - | + | - |
| **6w** | 7.01 | 7.46 | 7.01 | 6.98 | 8.24 | - | - | + | - | + | - |
| **6x** | 7.21 | 7.39 | 7.39 | 6.71 | 7.83 | - | - | + | - | + | - |
| **6y(R)** | 7.21 | 7.53 | 7.31 | 7.44 | 8.01 | - | +[a] | + | - | + | - |
| **6z(S)** | 7.16 | 7.50 | 7.13 | 7.65 | 8.13 | - | +[a] | + | - | + | - |
| **7e** | 7.79 | 7.61 | 7.49 | 7.13 | 8.31 | - | + | + | + | + | + |
| **7f** | 7.55 | 7.38 | 7.85 | 7.62 | 8.46 | - | + | + | + | + | + |
| **7g** | 7.90 | 7.91 | 7.73 | 7.42 | 8.54 | - | + | - | + | + | - |
| **7h** | 7.10 | 7.09 | 7.16 | 6.97 | 7.74 | - | +[a] | - | - | + | - |
| **7j** | 6.90 | 7.20 | 7.01 | 6.90 | 8.23 | - | + | + | + | + | + |
| **7k** | 8.64 | 7.09 | 8.29 | 8.40 | 8.26 | - | + | - | - | + | - |
| **7m(S)** | 7.43 | 7.43 | 7.72 | 7.30 | 8.41 | + | + | + | + | + | + |
| **7n(R)** | 7.44 | 7.61 | 7.50 | 7.05 | 7.97 | + | + | + | + | + | + |
| **7o** | 7.21 | 7.32 | 7.49 | 7.11 | 8.26 | + | + | - | + | + | + |
| **7r** | 7.37 | 7.19 | 7.53 | 7.10 | 8.67 | + | + | - | + | + | + |
| **7s** | 7.37 | 7.50 | 7.74 | 7.07 | 8.56 | + | + | - | + | + | + |
| **7w** | 7.30 | 7.46 | 7.45 | 6.77 | 7.98 | + | + | + | + | + | + |
| **7x** | 7.21 | 7.39 | 7.39 | 6.71 | 7.83 | + | + | + | + | + | + |
| **7y(R)** | 7.33 | 7.52 | 7.62 | 7.62 | 8.01 | + | + | - | + | + | - |
| **7z(S)** | 7.15 | 7.41 | 7.33 | 7.58 | 8.34 | + | + | - | + | + | - |
| **12a** | 8.99 | 7.77 | 8.23 | 8.64 | 8.15 | + | + | - | + | + | + |
| **12b** | 8.92 | 8.93 | 9.12 | 8.51 | 8.55 | + | + | - | + | + | + |
| **13a** | 7.86 | 8.22 | 7.65 | 8.27 | 8.65 | + | + | - | - | + | + |
| **13b** | 7.69 | 8.42 | 7.87 | 7.95 | 8.75 | + | + | - | + | + | + |
| **14a** | 8.28 | 8.34 | 8.12 | 8.42 | 8.60 | + | + | - | + | + | + |
| **14b** | 8.43 | 8.52 | 8.39 | 8.54 | 8.59 | + | + | - | + | + | + |

[a], hydrogen bond to ether oxygen;

***Table S4 Interactions with key residues during conventional molecular dynamics***

*Frequency of ligand interactions with key residues ($D^{3.32}$, $Y^{3.33}$, $T^{5.42}$, $W^{6.48}$, $Y^{6.51}$, $N^{6.52}$, $Y^{7.39}$ and $Y^{7.43}$) over the course of cMD is expressed as a fraction of timeframes. Values are averages from 3 independent MD runs. Values are coloured in a blue-white-red gradient. Values over 1.0 are possible as some receptor residues may make multiple contacts with the ligand.*

| | $D^{3.32}$ | | | | | $Y^{3.33}$ | | | | | $T^{5.42}$ | | | | | $W^{6.48}$ | | | | | $Y^{6.51}$ | | | | | $N^{6.52}$ | | | | | $Y^{7.39}$ | | | | | $Y^{7.43}$ | | | | |
|---|---|---|---|---|---|---|---|---|---|---|---|---|---|---|---|---|---|---|---|---|---|---|---|---|---|---|---|---|---|---|---|---|---|---|---|---|---|---|---|---|
| | $M_1$ | $M_2$ | $M_3$ | $M_4$ | $M_5$ | $M_1$ | $M_2$ | $M_3$ | $M_4$ | $M_5$ | $M_1$ | $M_2$ | $M_3$ | $M_4$ | $M_5$ | $M_1$ | $M_2$ | $M_3$ | $M_4$ | $M_5$ | $M_1$ | $M_2$ | $M_3$ | $M_4$ | $M_5$ | $M_1$ | $M_2$ | $M_3$ | $M_4$ | $M_5$ | $M_1$ | $M_2$ | $M_3$ | $M_4$ | $M_5$ | $M_1$ | $M_2$ | $M_3$ | $M_4$ | $M_5$ |
| **KH-5 (R)** | 0,37 | 0,37 | 0,34 | 0,38 | 0,37 | 0,69 | 0,73 | 0,76 | 0,74 | 0,65 | 0,19 | 0,21 | 0,19 | 0,21 | 0,21 | 0,87 | 0,81 | 0,86 | 0,9 | 0,94 | 0,65 | 0,67 | 0,66 | 0,59 | 0,59 | 0,93 | 0,96 | 0,99 | 0,98 | 0,9 | 0,67 | 0,72 | 0,66 | 0,67 | 0,61 | 0,84 | 0,92 | 0,77 | 0,92 | 0,87 |
| **KH-5 (S)** | 0,62 | 0,59 | 0,67 | 0,62 | 0,56 | 0,38 | 0,39 | 0,39 | 0,4 | 0,36 | 0,47 | 0,47 | 0,46 | 0,47 | 0,47 | 0,92 | 0,93 | 0,93 | 0,92 | 0,87 | 0,89 | 0,97 | 0,94 | 0,92 | 0,86 | 0,59 | 0,58 | 0,57 | 0,55 | 0,6 | 0,52 | 0,52 | 0,55 | 0,51 | 0,52 | 0,99 | 1,03 | 1,08 | 1,06 | 1,04 |
| **6a** | 0,04 | 0,04 | 0,04 | 0,04 | 0,04 | 1,04 | 1,13 | 0,95 | 1,03 | 1,12 | 0,02 | 0,02 | 0,02 | 0,02 | 0,02 | 0,85 | 0,86 | 0,78 | 0,85 | 0,86 | 0,31 | 0,29 | 0,31 | 0,32 | 0,34 | 0,03 | 0,03 | 0,03 | 0,04 | 0,03 | 0,09 | 0,1 | 0,1 | 0,1 | 0,09 | 0,84 | 0,86 | 0,77 | 0,78 | 0,84 |
| **6b** | 0,34 | 0,34 | 0,31 | 0,33 | 0,34 | 0,97 | 1,01 | 0,88 | 1 | 0,98 | 0,22 | 0,21 | 0,24 | 0,22 | 0,22 | 0,96 | 0,88 | 0,95 | 0,89 | 0,92 | 0,36 | 0,33 | 0,34 | 0,39 | 0,36 | 0,67 | 0,7 | 0,68 | 0,72 | 0,63 | 0,87 | 0,85 | 0,78 | 0,8 | 0,87 | 0,57 | 0,58 | 0,61 | 0,52 | 0,56 |
| **6c** | 0,03 | 0,03 | 0,03 | 0,03 | 0,03 | 1,14 | 1,2 | 1,04 | 1,23 | 1,14 | 0 | 0 | 0 | 0 | 0 | 0,58 | 0,57 | 0,61 | 0,56 | 0,55 | 0,29 | 0,27 | 0,27 | 0,31 | 0,29 | 0,01 | 0,01 | 0,01 | 0,01 | 0,01 | 0,81 | 0,84 | 0,82 | 0,76 | 0,73 | 0,81 | 0,8 | 0,78 | 0,76 | 0,74 |
| **6d** | 0,1 | 0,1 | 0,1 | 0,11 | 0,1 | 0,99 | 1,03 | 1,02 | 0,99 | 1,05 | 0,06 | 0,06 | 0,07 | 0,06 | 0,06 | 0,95 | 0,9 | 1,02 | 0,95 | 0,86 | 0,92 | 0,98 | 0,89 | 0,94 | 0,89 | 0,07 | 0,07 | 0,07 | 0,07 | 0,08 | 0,92 | 1,01 | 0,84 | 0,86 | 0,83 | 0,47 | 0,46 | 0,45 | 0,49 | 0,51 |
| **6e** | 0,1 | 0,1 | 0,1 | 0,09 | 0,09 | 0,87 | 0,84 | 0,94 | 0,82 | 0,87 | 0,52 | 0,57 | 0,52 | 0,5 | 0,52 | 0,7 | 0,63 | 0,72 | 0,63 | 0,71 | 0,49 | 0,53 | 0,46 | 0,46 | 0,45 | 0,57 | 0,58 | 0,6 | 0,57 | 0,59 | 0,4 | 0,39 | 0,37 | 0,42 | 0,42 | 0,63 | 0,61 | 0,66 | 0,58 | 0,6 |
| **6f** | 0,12 | 0,11 | 0,12 | 0,11 | 0,12 | 1,01 | 0,95 | 1,07 | 1,08 | 0,95 | 0,43 | 0,39 | 0,42 | 0,46 | 0,44 | 0,85 | 0,86 | 0,83 | 0,87 | 0,93 | 0,43 | 0,42 | 0,47 | 0,45 | 0,47 | 0,35 | 0,34 | 0,39 | 0,36 | 0,37 | 0,9 | 0,98 | 0,92 | 0,85 | 0,98 | 0,84 | 0,79 | 0,86 | 0,79 | 0,86 |
| **6g** | 0 | 0 | 0 | 0 | 0 | 0,16 | 0,15 | 0,16 | 0,15 | 0,15 | 0,03 | 0,02 | 0,03 | 0,03 | 0,03 | 0,45 | 0,43 | 0,42 | 0,45 | 0,41 | 0,23 | 0,25 | 0,23 | 0,21 | 0,24 | 0,05 | 0,05 | 0,04 | 0,05 | 0,05 | 0,33 | 0,33 | 0,32 | 0,34 | 0,32 | 0,16 | 0,17 | 0,16 | 0,17 | 0,15 |
| **6h** | 0 | 0 | 0 | 0 | 0 | 0,59 | 0,54 | 0,57 | 0,56 | 0,57 | 0 | 0 | 0 | 0 | 0 | 0,44 | 0,46 | 0,45 | 0,43 | 0,45 | 0,13 | 0,13 | 0,13 | 0,12 | 0,12 | 0 | 0 | 0 | 0 | 0 | 0,37 | 0,39 | 0,38 | 0,37 | 0,36 | 0 | 0 | 0 | 0 | 0 |
| **6j** | 0,07 | 0,07 | 0,07 | 0,08 | 0,06 | 0,32 | 0,3 | 0,33 | 0,31 | 0,32 | 0,16 | 0,15 | 0,17 | 0,17 | 0,14 | 0,85 | 0,79 | 0,86 | 0,93 | 0,9 | 0,61 | 0,62 | 0,63 | 0,6 | 0,55 | 0,14 | 0,14 | 0,13 | 0,15 | 0,14 | 0,7 | 0,68 | 0,67 | 0,71 | 0,74 | 0,92 | 0,89 | 0,85 | 0,99 | 0,98 |
| **6k** | 0,1 | 0,1 | 0,1 | 0,1 | 0,1 | 1,02 | 1,1 | 1 | 1,06 | 0,98 | 0,08 | 0,08 | 0,08 | 0,07 | 0,08 | 0,6 | 0,61 | 0,64 | 0,62 | 0,58 | 0,56 | 0,52 | 0,56 | 0,61 | 0,59 | 0,04 | 0,04 | 0,04 | 0,04 | 0,04 | 0,76 | 0,74 | 0,74 | 0,71 | 0,76 | 0,91 | 0,99 | 0,84 | 0,9 | 0,87 |
| **6l** | 0,14 | 0,13 | 0,13 | 0,13 | 0,15 | 0,99 | 0,9 | 0,89 | 1,02 | 0,91 | 0 | 0 | 0 | 0 | 0 | 0,09 | 0,09 | 0,09 | 0,09 | 0,09 | 0,33 | 0,35 | 0,32 | 0,35 | 0,33 | 0 | 0 | 0 | 0 | 0 | 0,69 | 0,65 | 0,68 | 0,69 | 0,67 | 0,89 | 0,85 | 0,94 | 0,87 | 0,86 |
| **6m(S)** | 0 | 0 | 0 | 0 | 0 | 0,17 | 0,19 | 0,17 | 0,16 | 0,17 | 0,23 | 0,24 | 0,23 | 0,21 | 0,22 | 0,29 | 0,29 | 0,27 | 0,27 | 0,3 | 0,33 | 0,3 | 0,35 | 0,33 | 0,33 | 0,33 | 0,34 | 0,3 | 0,31 | 0,32 | 0,42 | 0,41 | 0,44 | 0,44 | 0,42 | 0,18 | 0,16 | 0,2 | 0,17 | 0,17 |
| **6n(R)** | 0 | 0 | 0 | 0 | 0 | 0,21 | 0,21 | 0,21 | 0,19 | 0,22 | 0,16 | 0,16 | 0,15 | 0,14 | 0,17 | 0,23 | 0,22 | 0,23 | 0,21 | 0,21 | 0,39 | 0,39 | 0,42 | 0,37 | 0,4 | 0,52 | 0,51 | 0,56 | 0,55 | 0,55 | 0,25 | 0,25 | 0,23 | 0,26 | 0,27 | 0 | 0 | 0 | 0 | 0 |
| **6o** | 0 | 0 | 0 | 0 | 0 | 0,1 | 0,1 | 0,1 | 0,09 | 0,09 | 0,33 | 0,36 | 0,32 | 0,31 | 0,33 | 0,56 | 0,53 | 0,54 | 0,51 | 0,53 | 0,3 | 0,33 | 0,33 | 0,3 | 0,28 | 0,65 | 0,63 | 0,68 | 0,6 | 0,59 | 0,3 | 0,29 | 0,31 | 0,3 | 0,33 | 0,39 | 0,42 | 0,42 | 0,41 | 0,42 |
| **6p** | 0 | 0 | 0 | 0 | 0 | 0,11 | 0,11 | 0,11 | 0,11 | 0,11 | 0,17 | 0,18 | 0,17 | 0,16 | 0,17 | 0,59 | 0,54 | 0,54 | 0,64 | 0,65 | 0,48 | 0,44 | 0,5 | 0,45 | 0,46 | 0,69 | 0,74 | 0,72 | 0,75 | 0,65 | 0,33 | 0,32 | 0,36 | 0,32 | 0,32 | 0,21 | 0,19 | 0,2 | 0,23 | 0,2 |
| **6r** | 0 | 0 | 0 | 0 | 0 | 0,05 | 0,05 | 0,05 | 0,05 | 0,05 | 0,16 | 0,17 | 0,17 | 0,16 | 0,16 | 0,53 | 0,57 | 0,55 | 0,47 | 0,52 | 0,39 | 0,39 | 0,42 | 0,41 | 0,42 | 0,78 | 0,84 | 0,8 | 0,74 | 0,83 | 0,24 | 0,25 | 0,26 | 0,23 | 0,25 | 0,23 | 0,22 | 0,23 | 0,25 | 0,22 |
| **6s** | 0 | 0 | 0 | 0 | 0 | 0,17 | 0,18 | 0,18 | 0,17 | 0,19 | 0,24 | 0,26 | 0,24 | 0,25 | 0,25 | 0,45 | 0,49 | 0,42 | 0,49 | 0,46 | 0,25 | 0,22 | 0,24 | 0,23 | 0,26 | 0,72 | 0,75 | 0,76 | 0,72 | 0,67 | 0,21 | 0,2 | 0,21 | 0,21 | 0,23 | 0,32 | 0,34 | 0,33 | 0,35 | 0,33 |
| **6t** | 0 | 0 | 0 | 0 | 0 | 0,24 | 0,27 | 0,22 | 0,23 | 0,23 | 0,08 | 0,09 | 0,08 | 0,07 | 0,07 | 0,35 | 0,33 | 0,32 | 0,37 | 0,33 | 0,27 | 0,25 | 0,27 | 0,28 | 0,26 | 0,25 | 0,27 | 0,27 | 0,25 | 0,25 | 0,25 | 0,26 | 0,25 | 0,23 | 0,27 | 0,11 | 0,1 | 0,11 | 0,1 | 0,12 |
| **6w** | 0 | 0 | 0 | 0 | 0 | 0,25 | 0,23 | 0,23 | 0,26 | 0,23 | 0,11 | 0,1 | 0,11 | 0,11 | 0,12 | 0,26 | 0,27 | 0,25 | 0,26 | 0,25 | 0,24 | 0,25 | 0,23 | 0,22 | 0,23 | 0,43 | 0,4 | 0,39 | 0,44 | 0,44 | 0,3 | 0,27 | 0,29 | 0,28 | 0,29 | 0,27 | 0,29 | 0,27 | 0,29 | 0,26 |
| **6x** | 0 | 0 | 0 | 0 | 0 | 0,89 | 0,93 | 0,96 | 0,96 | 0,95 | 0,2 | 0,22 | 0,2 | 0,22 | 0,18 | 0,84 | 0,91 | 0,83 | 0,86 | 0,78 | 0,59 | 0,64 | 0,56 | 0,65 | 0,62 | 0,83 | 0,83 | 0,9 | 0,78 | 0,77 | 0,66 | 0,71 | 0,63 | 0,61 | 0,67 | 0,81 | 0,78 | 0,88 | 0,78 | 0,79 |
| **6y(R)** | 0 | 0 | 0 | 0 | 0 | 0,35 | 0,36 | 0,34 | 0,32 | 0,36 | 0,08 | 0,08 | 0,07 | 0,08 | 0,08 | 0,55 | 0,56 | 0,56 | 0,6 | 0,58 | 0,35 | 0,37 | 0,33 | 0,36 | 0,34 | 0,68 | 0,66 | 0,7 | 0,71 | 0,63 | 0,28 | 0,3 | 0,26 | 0,3 | 0,31 | 0,29 | 0,27 | 0,28 | 0,29 | 0,31 |
| **6z(S)** | 0 | 0 | 0 | 0 | 0 | 0,18 | 0,19 | 0,17 | 0,18 | 0,18 | 0,3 | 0,32 | 0,33 | 0,29 | 0,27 | 0,29 | 0,29 | 0,28 | 0,3 | 0,3 | 0,43 | 0,4 | 0,46 | 0,44 | 0,46 | 0,51 | 0,5 | 0,54 | 0,5 | 0,5 | 0,19 | 0,19 | 0,2 | 0,2 | 0,19 | 0,2 | 0,21 | 0,22 | 0,19 | 0,18 |
| **7e** | 0,31 | 0,33 | 0,29 | 0,33 | 0,31 | 0,65 | 0,63 | 0,67 | 0,62 | 0,64 | 0,18 | 0,18 | 0,18 | 0,17 | 0,19 | 0,75 | 0,69 | 0,72 | 0,78 | 0,74 | 0,35 | 0,33 | 0,32 | 0,33 | 0,33 | 0,39 | 0,41 | 0,42 | 0,36 | 0,35 | 0,66 | 0,59 | 0,64 | 0,7 | 0,63 | 0,81 | 0,87 | 0,8 | 0,82 | 0,84 |
| **7f** | 0,38 | 0,4 | 0,41 | 0,34 | 0,38 | 0,79 | 0,76 | 0,71 | 0,72 | 0,81 | 0,27 | 0,28 | 0,29 | 0,26 | 0,26 | 0,78 | 0,82 | 0,71 | 0,82 | 0,73 | 0,42 | 0,43 | 0,43 | 0,46 | 0,46 | 0,64 | 0,65 | 0,64 | 0,6 | 0,62 | 0,64 | 0,61 | 0,69 | 0,68 | 0,69 | 0,6 | 0,58 | 0,59 | 0,59 | 0,64 |
| **7g** | 0 | 0 | 0 | 0 | 0 | 0,44 | 0,44 | 0,48 | 0,46 | 0,47 | 0,12 | 0,14 | 0,13 | 0,13 | 0,12 | 0,46 | 0,43 | 0,42 | 0,43 | 0,42 | 0,68 | 0,61 | 0,74 | 0,74 | 0,63 | 0,93 | 0,99 | 0,86 | 0,89 | 0,9 | 0,11 | 0,12 | 0,12 | 0,12 | 0,12 | 0,16 | 0,16 | 0,16 | 0,15 | 0,18 |
| **7h** | 0,27 | 0,29 | 0,27 | 0,28 | 0,25 | 0,4 | 0,41 | 0,43 | 0,38 | 0,42 | 0,31 | 0,29 | 0,29 | 0,34 | 0,33 | 0,83 | 0,87 | 0,85 | 0,81 | 0,79 | 0,51 | 0,51 | 0,51 | 0,55 | 0,52 | 0,57 | 0,55 | 0,52 | 0,52 | 0,53 | 0,47 | 0,46 | 0,5 | 0,43 | 0,44 | 0,94 | 0,97 | 0,85 | 0,87 | 0,94 |
| **7j** | 0,58 | 0,59 | 0,6 | 0,55 | 0,61 | 0,44 | 0,44 | 0,48 | 0,4 | 0,41 | 0,23 | 0,24 | 0,24 | 0,25 | 0,21 | 0,99 | 0,99 | 0,9 | 0,99 | 1,01 | 0,55 | 0,52 | 0,53 | 0,51 | 0,58 | 0,55 | 0,51 | 0,6 | 0,56 | 0,59 | 0,67 | 0,73 | 0,71 | 0,73 | 0,61 | 0,97 | 1,03 | 0,92 | 0,93 | 0,94 |
| **7k** | 0,1 | 0,11 | 0,09 | 0,11 | 0,11 | 0,77 | 0,75 | 0,79 | 0,84 | 0,83 | 0,56 | 0,55 | 0,57 | 0,55 | 0,56 | 0,48 | 0,47 | 0,44 | 0,48 | 0,5 | 0,66 | 0,6 | 0,6 | 0,65 | 0,72 | 0,19 | 0,18 | 0,2 | 0,2 | 0,2 | 0 | 0 | 0 | 0 | 0 | 0,17 | 0,19 | 0,18 | 0,17 | 0,16 |
| **7m(S)** | 0,91 | 0,9 | 0,84 | 0,88 | 0,99 | 0,54 | 0,53 | 0,53 | 0,57 | 0,58 | 0,3 | 0,31 | 0,28 | 0,28 | 0,33 | 0,95 | 0,94 | 0,96 | 1,03 | 0,86 | 0,63 | 0,63 | 0,59 | 0,63 | 0,57 | 0,81 | 0,86 | 0,79 | 0,87 | 0,79 | 0,69 | 0,72 | 0,73 | 0,65 | 0,66 | 1 | 1,07 | 1,06 | 1,08 | 1,05 |
| **7n(R)** | 0,9 | 0,98 | 0,81 | 0,95 | 0,94 | 0,7 | 0,73 | 0,64 | 0,65 | 0,64 | 0,05 | 0,05 | 0,05 | 0,05 | 0,05 | 0,76 | 0,8 | 0,71 | 0,72 | 0,72 | 0,37 | 0,35 | 0,36 | 0,36 | 0,4 | 0,94 | 0,97 | 0,99 | 0,88 | 0,89 | 0,88 | 0,88 | 0,84 | 0,82 | 0,85 | 1 | 0,9 | 1,06 | 1,02 | 1,03 |
| **7o** | 0,78 | 0,81 | 0,79 | 0,81 | 0,7 | 0,94 | 0,85 | 0,88 | 0,98 | 0,85 | 0,2 | 0,22 | 0,21 | 0,19 | 0,18 | 0,63 | 0,67 | 0,67 | 0,62 | 0,64 | 0,45 | 0,49 | 0,42 | 0,46 | 0,43 | 0,65 | 0,68 | 0,59 | 0,7 | 0,64 | 0,57 | 0,54 | 0,61 | 0,6 | 0,55 | 0,59 | 0,57 | 0,53 | 0,59 | 0,57 |
| **7p** | 0,95 | 1 | 0,92 | 0,96 | 0,97 | 0,68 | 0,74 | 0,64 | 0,64 | 0,68 | 0,21 | 0,19 | 0,2 | 0,2 | 0,19 | 0,79 | 0,78 | 0,76 | 0,86 | 0,82 | 0,71 | 0,68 | 0,75 | 0,64 | 0,69 | 0,78 | 0,75 | 0,79 | 0,8 | 0,75 | 0,59 | 0,58 | 0,62 | 0,54 | 0,63 | 0,72 | 0,71 | 0,7 | 0,69 | 0,65 |
| **7r** | 0,96 | 0,94 | 0,91 | 0,87 | 1,04 | 0,92 | 0,83 | 0,93 | 0,92 | 0,95 | 0,56 | 0,57 | 0,61 | 0,56 | 0,57 | 0,7 | 0,74 | 0,68 | 0,71 | 0,72 | 0,53 | 0,53 | 0,57 | 0,48 | 0,56 | 0,94 | 0,94 | 0,99 | 1,03 | 0,88 | 0,41 | 0,38 | 0,44 | 0,37 | 0,38 | 0,98 | 1,06 | 1,03 | 0,99 | 0,9 |
| **7s** | 0,7 | 0,7 | 0,68 | 0,7 | 0,75 | 0,71 | 0,77 | 0,67 | 0,75 | 0,67 | 0,08 | 0,08 | 0,08 | 0,09 | 0,08 | 0,7 | 0,7 | 0,76 | 0,69 | 0,68 | 0,51 | 0,55 | 0,53 | 0,52 | 0,46 | 0,31 | 0,34 | 0,34 | 0,31 | 0,32 | 0,65 | 0,68 | 0,68 | 0,64 | 0,64 | 0,75 | 0,79 | 0,79 | 0,82 | 0,78 |
| **7w** | 0,66 | 0,6 | 0,69 | 0,62 | 0,71 | 0,72 | 0,76 | 0,79 | 0,72 | 0,7 | 0,12 | 0,12 | 0,12 | 0,13 | 0,12 | 0,79 | 0,81 | 0,73 | 0,8 | 0,74 | 0,69 | 0,75 | 0,65 | 0,68 | 0,67 | 0,62 | 0,62 | 0,6 | 0,67 | 0,59 | 0,6 | 0,66 | 0,59 | 0,62 | 0,59 | 0,99 | 0,95 | 0,94 | 0,96 | 1,03 |
| **7x** | 0,55 | 0,5 | 0,6 | 0,53 | 0,58 | 0,36 | 0,35 | 0,36 | 0,38 | 0,33 | 0 | 0 | 0 | 0 | 0 | 0,82 | 0,85 | 0,85 | 0,83 | 0,75 | 0,59 | 0,55 | 0,54 | 0,56 | 0,61 | 0,74 | 0,71 | 0,79 | 0,7 | 0,75 | 0,64 | 0,7 | 0,63 | 0,62 | 0,67 | 0,84 | 0,83 | 0,8 | 0,82 | 0,84 |
| **7y(R)** | 0,99 | 0,97 | 1,05 | 1,08 | 0,96 | 0,72 | 0,76 | 0,68 | 0,65 | 0,73 | 0,25 | 0,27 | 0,26 | 0,24 | 0,23 | 0,81 | 0,86 | 0,8 | 0,86 | 0,88 | 0,86 | 0,92 | 0,82 | 0,9 | 0,89 | 0,98 | 0,94 | 0,98 | 1,06 | 0,91 | 0,53 | 0,48 | 0,51 | 0,48 | 0,57 | 0,98 | 0,91 | 0,89 | 1 | 1,02 |
| **7z(S)** | 0,81 | 0,82 | 0,87 | 0,73 | 0,74 | 0,48 | 0,49 | 0,48 | 0,43 | 0,46 | 0,11 | 0,1 | 0,11 | 0,1 | 0,1 | 0,7 | 0,72 | 0,74 | 0,75 | 0,72 | 0,52 | 0,48 | 0,52 | 0,47 | 0,55 | 0,81 | 0,81 | 0,85 | 0,81 | 0,73 | 0,69 | 0,76 | 0,74 | 0,67 | 0,62 | 0,9 | 0,92 | 0,89 | 0,98 | 0,84 |
| **12a** | 0,97 | 0,99 | 0,94 | 0,97 | 1,05 | 0,82 | 0,76 | 0,84 | 0,8 | 0,88 | 0,24 | 0,25 | 0,23 | 0,26 | 0,25 | 0,41 | 0,43 | 0,38 | 0,45 | 0,39 | 0,78 | 0,73 | 0,71 | 0,86 | 0,83 | 0,87 | 0,85 | 0,79 | 0,8 | 0,88 | 0,56 | 0,57 | 0,58 | 0,59 | 0,55 | 0,87 | 0,89 | 0,88 | 0,8 | 0,9 |
| **12b** | 0,42 | 0,45 | 0,44 | 0,4 | 0,38 | 0,97 | 1,06 | 0,96 | 1,02 | 0,95 | 0,25 | 0,27 | 0,24 | 0,27 | 0,26 | 0,37 | 0,34 | 0,37 | 0,35 | 0,4 | 0,22 | 0,23 | 0,24 | 0,24 | 0,22 | 0,88 | 0,85 | 0,94 | 0,82 | 0,81 | 0,91 | 0,95 | 0,9 | 0,92 | 0,94 | 0,81 | 0,74 | 0,74 | 0,84 | 0,74 |
| **13a** | 0,97 | 0,88 | 0,91 | 1,02 | 0,98 | 1,04 | 1,09 | 1,06 | 1,06 | 1,12 | 0,35 | 0,38 | 0,37 | 0,36 | 0,37 | 0,32 | 0,29 | 0,34 | 0,29 | 0,33 | 0,35 | 0,37 | 0,34 | 0,37 | 0,36 | 0,42 | 0,46 | 0,42 | 0,43 | 0,44 | 0,47 | 0,43 | 0,5 | 0,45 | 0,51 | 0,78 | 0,8 | 0,83 | 0,83 | 0,84 |
| **13b** | 0,7 | 0,7 | 0,73 | 0,66 | 0,68 | 0,76 | 0,78 | 0,79 | 0,7 | 0,82 | 0,32 | 0,29 | 0,31 | 0,34 | 0,34 | 0,67 | 0,74 | 0,68 | 0,63 | 0,61 | 0,44 | 0,46 | 0,44 | 0,48 | 0,45 | 0,42 | 0,46 | 0,45 | 0,41 | 0,46 | 0,46 | 0,47 | 0,5 | 0,49 | 0,48 | 0,7 | 0,75 | 0,75 | 0,67 | 0,76 |
| **14a** | 1 | 0,96 | 1,1 | 1,07 | 0,95 | 0,78 | 0,83 | 0,7 | 0,8 | 0,76 | 0,4 | 0,4 | 0,39 | 0,37 | 0,44 | 0,65 | 0,63 | 0,65 | 0,6 | 0,63 | 0,32 | 0,35 | 0,33 | 0,35 | 0,3 | 0,17 | 0,18 | 0,15 | 0,16 | 0,18 | 0,42 | 0,41 | 0,41 | 0,41 | 0,41 | 0,58 | 0,55 | 0,52 | 0,6 | 0,57 |
| **14b** | 0,79 | 0,75 | 0,72 | 0,82 | 0,86 | 0,8 | 0,85 | 0,74 | 0,8 | 0,81 | 0,59 | 0,57 | 0,59 | 0,61 | 0,54 | 0,73 | 0,8 | 0,77 | 0,74 | 0,67 | 0,52 | 0,49 | 0,52 | 0,56 | 0,57 | 0,63 | 0,67 | 0,65 | 0,65 | 0,58 | 0,64 | 0,69 | 0,7 | 0,61 | 0,63 | 0,93 | 0,85 | 0,87 | 0,88 | 0,87 |

***Table S5 Molecular descriptors of tested compounds***

*The distance between ketoxy oxygen and nitrogen (Distance), the improper dihedral angle (Angle), molecular volume (Mol. Volume), basic centre (BC) volume, radius and surface area and electrostatic potential (ESP) and net charge of molecule.*

| Compound | Distance [Å] | Angle [°] | Mol_volume [Å^3] | BC_volume [Å^3] | BC_radius [Å] | BC_surface [Å^2] | ESP [kcal/mol] | Net_charge |
|---|---|---|---|---|---|---|---|---|
| **6a** | 5.192 | 1.933 | 416.94 | 161.14 | 5.212 | 309.74 | -1.119 | 0 |
| **6b** | 5.306 | 0.996 | 422.89 | 167.54 | 4.482 | 303.63 | 1.837 | 0.205 |
| **6c** | 5.322 | 1.05 | 372.32 | 118.71 | 4.076 | 250.12 | -0.163 | 0 |
| **6d** | 5.272 | -1.04 | 418.14 | 163.22 | 5.07 | 311.5 | 1.436 | 0 |
| **6e** | 5.278 | 1.664 | 395.77 | 142.53 | 4.362 | 289.6 | 1.724 | 0 |
| **6f** | 5.279 | -0.256 | 392.82 | 141.69 | 4.375 | 284.89 | 1.772 | 0 |
| **6g** | 5.235 | 2.947 | 400.2 | 147.55 | 4.656 | 291.68 | -0.962 | 0 |
| **6h** | 5.916 | 4.678 | 357.79 | 104.28 | 3.996 | 237.24 | -1.227 | 0 |
| **6j** | 5.192 | 3.604 | 358.77 | 107.18 | 4.105 | 244.53 | -1.107 | 0 |
| **6k** | 5.186 | -0.683 | 410.79 | 172.65 | 5.818 | 331.92 | -1.779 | 0 |
| **6l** | 5.164 | 2.059 | 443.66 | 185.09 | 5.555 | 339.4 | -2.127 | 0 |
| **6m** | 4.903 | -64.339 | 377.5 | 142.23 | 4.369 | 266.2 | -1.075 | 0 |
| **6n** | 4.923 | 70.209 | 377.27 | 141.96 | 4.369 | 266.2 | -0.834 | 0 |
| **6o** | 6.575 | 11.321 | 379.66 | 142.49 | 4.199 | 261.3 | -2.453 | 0 |
| **6p** | 6.575 | 11.321 | 379.66 | 142.49 | 4.199 | 261.3 | -2.511 | 0 |
| **6r** | 6.591 | 2.356 | 377.45 | 142.4 | 4.199 | 261.3 | -2.511 | 0 |
| **6s** | 5.465 | -26.118 | 359.76 | 123.47 | 4.225 | 259.1 | -2.636 | 0 |
| **6t** | 4.856 | -76.728 | 359.9 | 123.83 | 4.088 | 244.19 | -1.24 | 0 |
| **6w** | 4.856 | -76.728 | 359.9 | 123.83 | 4.088 | 244.19 | -1.24 | 0 |
| **6x** | 5.18 | 6.11 | 358.83 | 125.6 | 4.088 | 244.19 | 0.922 | 0 |
| **6y** | 5.841 | -28.102 | 362.08 | 124.85 | 4.108 | 242.94 | -2.706 | 0 |
| **6z** | 5.852 | -27.979 | 360.67 | 125.36 | 4.108 | 242.94 | -2.708 | 0 |
| **7e** | 5.296 | 6.045 | 414.47 | 180.21 | 4.342 | 298.74 | 1.841 | 0.205 |
| **7f** | 5.309 | 6.625 | 413.65 | 161.45 | 4.374 | 301.21 | 1.825 | 0.205 |
| **7g** | 5.304 | 6.782 | 420.31 | 168.98 | 4.477 | 301.87 | 1.362 | 0.205 |
| **7h** | 5.292 | 5.895 | 375.73 | 122.22 | 4.246 | 261.89 | 1.541 | 0.205 |
| **7j** | 5.292 | 7.013 | 382.1 | 127.15 | 4.019 | 263.91 | 1.479 | 0.205 |
| **7k** | 5.288 | -0.984 | 428.76 | 155.46 | 5.178 | 307.16 | 1.387 | 0.205 |
| **7m (S)** | 5.279 | 0.37 | 395.63 | 160.83 | 4.328 | 282.24 | 0.327 | 0.205 |
| **7n (R)** | 5.281 | 10.379 | 398.79 | 161.19 | 4.328 | 282.24 | 0.339 | 0.205 |
| **7o** | 6.652 | 4.684 | 394.82 | 160.25 | 4.286 | 281.97 | 0.534 | 0.205 |
| **7p** | 6.639 | 9.144 | 395.59 | 160.75 | 4.286 | 281.97 | 0.534 | 0.205 |
| **7r** | 6.639 | 9.144 | 395.59 | 160.75 | 4.286 | 281.97 | 0.534 | 0.205 |
| **7s** | 5.515 | -23.538 | 375.42 | 144.81 | 4.352 | 282.29 | 0.009 | 0.205 |
| **7w** | 5.288 | -1.433 | 377.9 | 143.73 | 4.007 | 265.18 | 0.922 | 0.205 |
| **7x** | 5.282 | 6.358 | 380.09 | 145.12 | 4.007 | 265.18 | 0.922 | 0.205 |
| **7y (R)** | 4.971 | -58.758 | 382.92 | 144.82 | 4.039 | 268.37 | 1.217 | 0.205 |
| **7z (S)** | 5.967 | -25.172 | 378.28 | 145.88 | 4.039 | 268.37 | 0.285 | 0.205 |
| **12a** | 5.308 | 32.732 | 365.52 | 130.59 | 3.744 | 266.55 | 2.507 | 0.205 |
| **12b** | 5.316 | 33.935 | 384.62 | 150.47 | 4.284 | 287.45 | 1.712 | 0.205 |
| **13a** | 5.308 | 32.732 | 333.42 | 130.59 | 3.744 | 266.55 | 2.507 | 0.205 |
| **13b** | 5.316 | 33.935 | 348.71 | 150.47 | 4.284 | 287.45 | 1.712 | 0.205 |
| **14a** | 5.308 | 32.732 | 279.12 | 130.59 | 3.744 | 266.55 | 2.507 | 0.205 |
| **14b** | 5.316 | 33.935 | 294.01 | 150.47 | 4.284 | 287.45 | 1.712 | 0.205 |

***Table S6 Receptor RMSF of individual transmembrane helices and loops***

*RMSF is in Å. Data means ± SD from 3 independent runs of MD. *, lower than apo (P<0.05 one-tailed T-test with Bonferroni correction for multiple comparisons).*

| $M_1$ | apo | KH-5 | 12a | 7s | 7w |
|---|---|---|---|---|---|
| N-terminus | 1.76 ± 0.52 | 1.71 ± 0.33 | 1.76 ± 0.67 | 1.47 ± 0.41 | 1.94 ± 0.68 |
| TM1 | 0.86 ± 0.21 | 0.93 ± 0.26 | 0.79 ± 0.13 | 0.81 ± 0.17 | 1.05 ± 0.46 |
| ICL1 | 1.38 ± 0.49 | 1.06 ± 0.28 | 0.97 ± 0.23 | 1.06 ± 0.31 | 1.04 ± 0.21 |
| TM2 | 0.75 ± 0.16 | 0.68 ± 0.24 | 0.60 ± 0.11 | 0.66 ± 0.09 | 0.61 ± 0.12 |
| ECL1 | 1.02 ± 0.21 | 0.87 ± 0.14 | 0.99 ± 0.18 | 0.95 ± 0.18 | 0.92 ± 0.14 |
| TM3 | 0.84 ± 0.23 | 0.60 ± 0.12 | 0.58 ± 0.07 | 0.58 ± 0.08 | 0.62 ± 0.09 |
| ICL2 | 1.48 ± 0.76 | 1.17 ± 0.45 | 0.98 ± 0.31 | 1.11 ± 0.38 | 1.21 ± 0.60 |
| TM4 | 1.12 ± 0.36 | 0.93 ± 0.28 | 1.19 ± 0.61 | 1.00 ± 0.37 | 1.05 ± 0.35 |
| ECL2 | 1.38 ± 0.41 | 1.10 ± 0.32 | 1.36 ± 0.49 | 1.16 ± 0.41 | 1.35 ± 0.51 |
| TM5 | 1.21 ± 0.37 | 0.95 ± 0.32 | 0.89 ± 0.29 | 0.97 ± 0.30 | 0.93 ± 0.30 |
| ICL3 | 2.62 ± 1.34 | 3.14 ± 1.87 | 2.62 ± 1.67 | 2.71 ± 2.12 | 2.41 ± 1.28 |
| TM6 | 1.04 ± 0.29 | 1.00 ± 0.31 | 0.79 ± 0.15 | 0.88 ± 0.21 | 0.90 ± 0.19 |
| ECL3 | 1.80 ± 0.39 | 1.38 ± 0.42 | 1.14 ± 0.29 | 1.37 ± 0.34 | 1.44 ± 0.47 |
| TM7 | 1.09 ± 0.47 | 0.81 ± 0.23 | 0.79 ± 0.30 | 0.85 ± 0.26 | 0.98 ± 0.49 |
| C-terminus | 1.81 ± 0.98 | 1.60 ± 0.88 | 1.61 ± 0.62 | 1.49 ± 0.69 | 1.54 ± 0.76 |

| $M_2$ | apo | KH-5 | 12a | 7s | 7w |
|---|---|---|---|---|---|
| N-terminus | 2.62 ± 1.23 | 2.01 ± 0.92 | 1.70 ± 0.61 | 1.75 ± 0.93 | 2.04 ± 0.86 |
| TM1 | 0.93 ± 0.28 | 0.77 ± 0.21 | 0.69 ± 0.13 | 0.77 ± 0.17 | 0.77 ± 0.17 |
| ICL1 | 1.55 ± 0.55 | 0.99 ± 0.14 | 0.97 ± 0.19 | 0.97 ± 0.24 | 1.03 ± 0.21 |
| TM2 | 0.84 ± 0.19 | 0.65 ± 0.11 | 0.63 ± 0.12 | 0.62 ± 0.12 | 0.71 ± 0.17 |
| ECL1 | 1.04 ± 0.23 | 0.92 ± 0.15 | 0.95 ± 0.15 | 0.88 ± 0.12 | 1.18 ± 0.31 |
| TM3 | 0.92 ± 0.20 | 0.70 ± 0.10 | 0.67 ± 0.16 | 0.64 ± 0.10 | 0.81 ± 0.22 |
| ICL2 | 1.48 ± 0.35 | 1.27 ± 0.27 | 1.18 ± 0.22 | 1.45 ± 0.51 | 1.47 ± 0.44 |
| TM4 | 1.06 ± 0.25 | 0.90 ± 0.13 | 0.91 ± 0.24 | 0.85 ± 0.22 | 1.03 ± 0.24 |
| ECL2 | 1.39 ± 0.40 | 1.18 ± 0.28 | 1.17 ± 0.34 | 1.17 ± 0.27 | 1.11 ± 0.24 |
| TM5 | 1.17 ± 0.30 | 1.16 ± 0.23 | 1.13 ± 0.24 | 1.03 ± 0.23 | 1.09 ± 0.18 |
| ICL3 | 2.65 ± 1.71 | 2.83 ± 1.24 | 2.68 ± 1.32 | 2.26 ± 1.14 | 2.51 ± 1.29 |
| TM6 | 1.22 ± 0.46 | 0.95 ± 0.29 | 0.88 ± 0.20 | 0.94 ± 0.28 | 1.10 ± 0.39 |
| ECL3 | 1.67 ± 0.39 | 1.55 ± 0.40 | 1.43 ± 0.33 | 1.60 ± 0.38 | 1.64 ± 0.38 |
| TM7 | 1.09 ± 0.32 | 0.78 ± 0.15 | 0.82 ± 0.23 | 0.86 ± 0.24 | 0.92 ± 0.22 |
| C-terminus | 1.80 ± 0.80 | 1.39 ± 0.86 | 1.32 ± 0.77 | 1.32 ± 0.67 | 1.30 ± 0.71 |

| $M_3$ | apo | KH-5 | 12a | 7s | 7w |
|---|---|---|---|---|---|
| N-terminus | 2.48 ± 1.26 | 1.80 ± 0.75 | 1.87 ± 0.91 | 1.77 ± 0.69 | 1.54 ± 0.85 |
| TM1 | 1.10 ± 0.41 | 0.89 ± 0.14 | 0.89 ± 0.20 | 0.85 ± 0.14 | 0.90 ± 0.18 |
| ICL1 | 1.40 ± 0.41 | 1.23 ± 0.33 | 1.15 ± 0.30 | 1.10 ± 0.39 | 1.34 ± 0.41 |
| TM2 | 0.91 ± 0.21 | 0.70 ± 0.12 | 0.69 ± 0.13 | 0.66 ± 0.26 | 0.75 ± 0.14 |
| ECL1 | 1.56 ± 0.48 | 1.11 ± 0.19 | 1.23 ± 0.17 | 1.22 ± 0.29 | 1.07 ± 0.18 |
| TM3 | 0.95 ± 0.25 | 0.69 ± 0.17 | 0.76 ± 0.20 | 0.65 ± 0.14 | 0.78 ± 0.14 |
| ICL2 | 1.69 ± 0.58 | 1.79 ± 0.82 | 1.68 ± 0.48 | 1.23 ± 0.28 | 1.50 ± 0.34 |
| TM4 | 1.14 ± 0.38 | 0.92 ± 0.27 | 1.00 ± 0.40 | 0.91 ± 0.24 | 0.99 ± 0.17 |
| ECL2 | 1.62 ± 0.39 | 1.23 ± 0.30 | 1.47 ± 0.39 | 1.16 ± 0.22 | 1.24 ± 0.42 |
| TM5 | 1.23 ± 0.41 | 0.83 ± 0.14 | 0.84 ± 0.12 | 0.74 ± 0.07 * | 0.95 ± 0.15 |
| ICL3 | 2.00 ± 0.94 | 1.80 ± 1.19 | 1.74 ± 1.15 | 1.87 ± 1.83 | 1.68 ± 1.12 |
| TM6 | 1.25 ± 0.32 | 0.87 ± 0.21 | 0.86 ± 0.22 | 0.80 ± 0.24 | 0.96 ± 0.29 |
| ECL3 | 1.79 ± 0.45 | 1.29 ± 0.25 | 1.26 ± 0.23 | 1.24 ± 0.31 | 1.22 ± 0.27 |
| TM7 | 1.09 ± 0.34 | 0.88 ± 0.28 | 0.83 ± 0.24 | 0.71 ± 0.17 | 0.92 ± 0.26 |
| C-terminus | 2.47 ± 1.19 | 2.30 ± 1.13 | 2.03 ± 1.08 | 2.24 ± 1.09 | 3.37 ± 2.66 |

| $M_4$ | apo | KH-5 | 12a | 7s | 7w |
|---|---|---|---|---|---|
| N-terminus | 2.55 ± 1.38 | 2.57 ± 1.83 | 2.45 ± 1.47 | 2.62 ± 1.94 | 2.70 ± 1.73 |
| TM1 | 0.91 ± 0.23 | 0.79 ± 0.20 | 0.79 ± 0.23 | 0.91 ± 0.32 | 0.89 ± 0.32 |
| ICL1 | 1.56 ± 0.88 | 0.89 ± 0.21 | 0.99 ± 0.31 | 0.95 ± 0.17 | 0.90 ± 0.23 |
| TM2 | 0.87 ± 0.24 | 0.76 ± 0.22 | 0.72 ± 0.24 | 0.84 ± 0.18 | 0.73 ± 0.30 |
| ECL1 | 1.24 ± 0.22 | 1.00 ± 0.13 | 1.15 ± 0.26 | 1.03 ± 0.15 | 1.27 ± 0.27 |
| TM3 | 0.99 ± 0.26 | 0.75 ± 0.21 | 0.71 ± 0.19 | 0.79 ± 0.22 | 0.73 ± 0.18 |
| ICL2 | 1.50 ± 0.45 | 1.05 ± 0.25 | 1.65 ± 0.78 | 1.28 ± 0.34 | 1.13 ± 0.18 |
| TM4 | 1.12 ± 0.32 | 0.95 ± 0.22 | 0.99 ± 0.22 | 0.95 ± 0.15 | 0.98 ± 0.23 |
| ECL2 | 1.43 ± 0.32 | 1.15 ± 0.35 | 1.14 ± 0.29 | 1.12 ± 0.33 | 1.33 ± 0.58 |
| TM5 | 1.28 ± 0.36 | 0.90 ± 0.17 | 0.92 ± 0.22 | 0.97 ± 0.20 | 0.91 ± 0.19 |
| ICL3 | 2.79 ± 1.15 | 2.52 ± 1.18 | 2.73 ± 1.16 | 2.88 ± 1.43 | 2.45 ± 1.12 |
| TM6 | 1.27 ± 0.49 | 0.92 ± 0.34 | 0.88 ± 0.31 | 0.94 ± 0.34 | 0.90 ± 0.25 |
| ECL3 | 2.06 ± 0.74 | 1.32 ± 0.47 | 1.34 ± 0.41 | 1.48 ± 0.53 | 1.29 ± 0.36 |
| TM7 | 1.13 ± 0.47 | 0.87 ± 0.24 | 0.85 ± 0.27 | 1.04 ± 0.22 | 0.87 ± 0.23 |
| C-terminus | 1.65 ± 0.67 | 1.15 ± 0.35 | 1.32 ± 0.37 | 1.25 ± 0.38 | 1.20 ± 0.32 |

| $M_5$ | apo | KH-5 | 12a | 7s | 7w |
|---|---|---|---|---|---|
| N-terminus | 3.16 ± 0.89 | 1.91 ± 0.70 | 1.71 ± 0.58 | 1.98 ± 0.65 | 1.89 ± 0.76 |
| TM1 | 1.61 ± 0.64 | 0.84 ± 0.28 | 0.79 ± 0.25 | 0.93 ± 0.38 | 0.82 ± 0.29 |
| ICL1 | 1.69 ± 0.41 | 1.09 ± 0.16 * | 1.12 ± 0.29 | 1.05 ± 0.18 * | 1.03 ± 0.18 * |
| TM2 | 1.18 ± 0.42 | 0.61 ± 0.07 * | 0.64 ± 0.09 * | 0.69 ± 0.15 | 0.61 ± 0.10 * |
| ECL1 | 2.02 ± 0.86 | 0.84 ± 0.16 * | 0.84 ± 0.14 * | 1.00 ± 0.23 | 0.89 ± 0.23 * |
| TM3 | 1.27 ± 0.48 | 0.60 ± 0.09 * | 0.61 ± 0.07 * | 0.62 ± 0.11 * | 0.57 ± 0.08 * |
| ICL2 | 2.43 ± 0.75 | 1.20 ± 0.32 * | 1.38 ± 0.49 | 1.23 ± 0.39 * | 1.33 ± 0.42 |
| TM4 | 1.53 ± 0.52 | 0.76 ± 0.11 * | 0.84 ± 0.18 | 0.78 ± 0.13 * | 0.79 ± 0.14 * |
| ECL2 | 2.29 ± 0.76 | 1.01 ± 0.26 * | 1.03 ± 0.25 * | 1.10 ± 0.31 * | 1.04 ± 0.24 * |
| TM5 | 1.49 ± 0.47 | 0.75 ± 0.11 * | 0.82 ± 0.12 * | 0.78 ± 0.11 * | 0.77 ± 0.22 * |
| ICL3 | 3.67 ± 1.71 | 1.86 ± 0.88 | 2.23 ± 1.25 | 1.69 ± 0.84 | 2.38 ± 1.75 |
| TM6 | 1.49 ± 0.44 | 0.90 ± 0.24 | 0.84 ± 0.24 | 0.88 ± 0.22 | 0.74 ± 0.15 * |
| ECL3 | 2.33 ± 0.79 | 1.35 ± 0.33 | 1.47 ± 0.38 | 1.37 ± 0.38 | 1.28 ± 0.32 |
| TM7 | 1.33 ± 0.50 | 0.75 ± 0.20 | 0.73 ± 0.22 | 0.78 ± 0.17 | 0.68 ± 0.11 * |
| C-terminus | 2.35 ± 1.00 | 1.20 ± 0.61 | 1.48 ± 0.83 | 1.28 ± 0.75 | 1.41 ± 0.81 |
